# A high throughput lipidomics method using *scheduled* multiple reaction monitoring

**DOI:** 10.1101/2021.01.08.425875

**Authors:** Akash Kumar Bhaskar, Salwa Naushin, Arjun Ray, Shalini Pradhan, Khushboo Adlakha, Towfida Jahan Siddiqua, Dipankar Malakar, Shantanu Sengupta

## Abstract

Lipid compositions of cells, tissues and bio-fluids are complex, with varying concentrations and structural diversity, which makes their identification challenging. Newer methods for comprehensive analysis of lipids are thus necessary. Herein, we propose a targeted-mass spectrometry based method for large-scale lipidomics using a combination of variable retention time window and relative dwell time weightage. Using this, we detected more than 1000 lipid species, including structural isomers. The limit of detection varied from femtomolar to nanomolar range and the coefficient of variance <30% for 849 lipid species. We used this method to identify lipids altered due to Vitamin B_12_ deficiency and found that the levels of lipids with ω-3 fatty acid chains decreased while those with ω-6 increased. This method enables identification of by far the largest number of lipid species with structural isomers in a single experiment and would significantly advance our understanding of the role of lipids in biological processes.

## Introduction

Lipid constitutes highly diverse biomolecules, which play an important role in the normal functioning of the body, maintaining the cellular homeostasis, cell signaling and energy storage^1–5^. Dysregulation of lipid homeostasis is associated with a large number of pathologies such as obesity and diabetes^6,7^, cardiovascular disease^8^, cancer^9^ and other metabolic dieases^10^. Lipid compositions of cells, tissues and bio-fluids are complex, reflecting a wide range of concentrations of different lipid classes and structural diversity within lipid species^11,12^. Although the exact number of distinct lipids present in cells is not exactly known, it is believed that the cellular lipidome consists of more than 1000 different lipid species each with several structural isomers^4,13–15^.

Identification of lipids using traditional methods like thin layer chromatography (TLC), nuclear magnetic resonance (NMR), and soft ionization techniques (field desorption, chemical ionization or fast atom bombardment) are limited by their lower sensitivity and accuracy, hence is not suitable for comprehensive lipidomics studies^16,17^. Recent advances in electrospray ionization-mass spectrometry (ESI-MS) based lipidomics have enabled accurate identification of a large number of lipid species from various biological sources^18,19^. Analysis of lipids in both positive and negative ion modes in a single mass spectrometric scan using untargeted or targeted approaches have been used for greater coverage with increasing sensitivity and specificity^20,21^. The untargeted lipidomics approach however has some major challenges especially with respect to specific identification (without standards) and characterization of the lipid species, time required to process large quantity of raw data and the bias towards the detection of lipids with high-abundance^19,22^. These problems are greatly reduced in a targeted approach using multiple reaction monitoring (MRM), since defined groups of chemically characterized and annotated lipid species are analyzed^22,23^. The use of MRM enables simultaneous identification of numerous lipid species, including those with low abundance^24,25^. The number of lipid species identified could be further increased by using scheduled MRM, where the MRM transitions are monitored only around the expected retention time of the eluting lipid species^21,26,27^. This enables monitoring of greater number of MRM transitions in a single MS acquisition. Using scheduled MRM, Takeda et. al., and other groups, were able to identify/ quantify 413 of lipid species including isomers of phospholipids (PLs) and diacylglycerol (DAG) in a single targeted scan^21,28,29^. However, identification of triacylglycerols (TAG) was based on pseudo-transitions (the precursor and product ion are same) as identifying different species of TAG is challenging^21,30,31^.

In *scheduled* MRM, the retention time window assigned is primarily of fixed width. However, as the retention time window width varies for each lipid species, a variable window width for each lipid species could reduce the time necessary to develop high throughput targeted methods. There are a few reports where variable retention time window (dynamic MRM) has been used in various applications, including identifying lipids of a specific class^32–38^. However, none of these studies involved comprehensive lipidome analysis. Further, in these studies, the dwell time for each peak was automatically fixed based on the RT window width chosen. The quality of peaks can be improved by varying dwell time weightage for each transition without compromising with the cycle time (https://https://sciex.com/). Assigning a low dwell time weightage to high abundant compounds and high dwell time weightage to less abundant compounds, irrespective of the elution window, may help in accommodating large number of transitions in a single run with improved data quality.

Leveraging the combinatorial optimization of *scheduled*-MRM, variable RT window and dwell time weightage, we report a rapid and sensitive targeted lipidomics method capable of identifying more than 1000 lipid species, including isomers of triglycerides, diglycerides, and phospholipids in a single MS run-time of 24 minutes. To the best of our knowledge, this is the largest number of lipid species identified till date in a single experiment.

We further, exploited this method to quantitate isomer specific different lipid classes in vitamin B_12_ deficiency in the context of Indian population. Previously, we have shown that vitamin B12 deficiency alters the lipid metabolism to drive cardiometabolic phenotype in rats^39^. This study clearly demonstrates the effects of vitamin B_12_ deficiency with changes in the specific lipidomic isomers, laying the foundation to understand the development of highly prevalent cardio-metabolic diseases in a strictly vegetarian diet adhered country like India.

## Results

We developed a *scheduled*-MRM method that can identify more than 1000 lipid species in a single mass spectrometric acquisition using a combination of variable-RTW and relative-DTW for each lipid species along with an optimized LC-gradient. Initially, we generated a theoretical MRM library using LIPIDMAPS (http://www.lipidmaps.org/) which consisted of 1224 lipid species and 12 internal standards, belonging to the 18 lipid classes. The total ion chromatogram is shown in figure 1a. The 18 classes of lipids were analyzed in the positive or negative ion modes. In the positive ion mode, the M+H precursor ions were used for SM, Cer, CE, while for neutral lipids (TAG, DAG, and MAG) [M+NH4] precursor ions were considered. Phospholipids (PL’s) were identified in negative ion mode, forming [M-H] precursor ion except LPC’s and PC’s, for which [M+CH3COO]-were considered.

**Figure 1.**
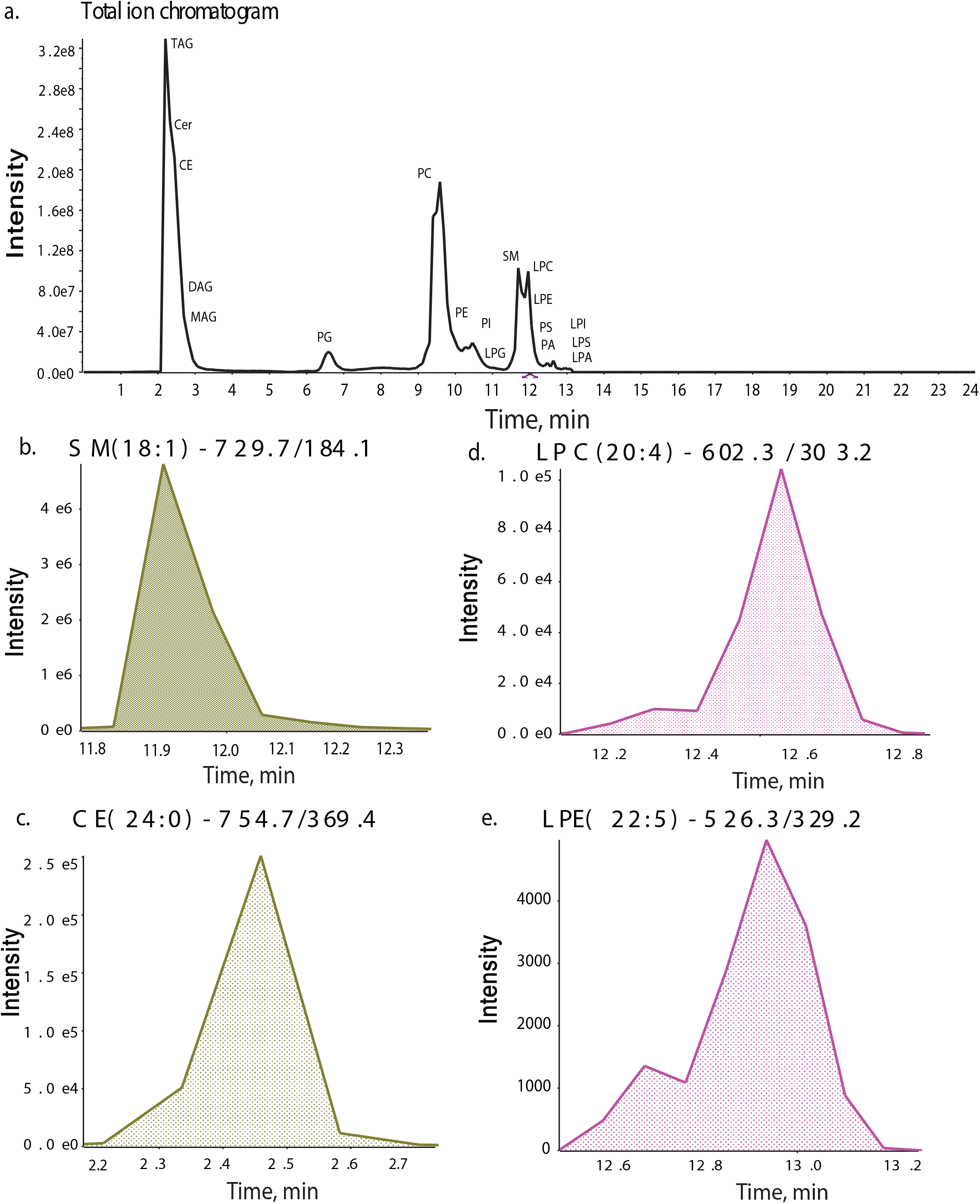
Chromatograms of the *scheduled* MRM method with variable-RTW and relative-DTW. (**a)** A total ion chromatogram of method consisting of 1224 lipid species and 12 internal standards from 18 lipid classes in positive or negative mode. **b, c** In positive ion mode, SM (18:1)-729/184.1 has elution window of 36.1 seconds with dwell weight 1 (**b**) and CE (24:0)-754.7/369.4 has elution window of 32.5 seconds with dwell weight 3.01 (**c)**. **d, e** In negative ion mode, LPC (20:4)-602.3/303.2 and LPE (22:5)-526.3/329.2 has equal elution window (40.2 seconds) but LPE (22:5) has higher dwell weight (1.15) (**d**) compared to LPC (20:4) dwell weight (1) (**e**).

The variable-RTW and relative-DTW for different species was determined based on the intensity and width of the peaks obtained for each lipid species. For instance, in positive ion mode, SM (18:1) had a broader elution window (36.1 seconds) compared to CE (24:0) (32.5 seconds), but the signal intensity of CE (24:0) was lower as compared to SM (18:1). Thus, to collect sufficient number of data points, higher dwell time weight of 3.01 was applied for CE (24:0) as compared to 1.00 for SM (18:1) (figure 1b and 1c). Furthermore, LPC (20:4) and LPE (22:5), had the same elution window of 40.2 seconds but a dwell time weightage of 1 was applied for LPC (20:4) as compared to 1.15 for LPE (22:5; figure 1d and 1e). A complete list of all parameters for each lipid species along with retention window and dwell weightage is given in supplementary table 1.

### Identification of isomers within lipid classes

In an attempt to identify different lipid isomers, we used customized-approaches for various lipid classes. For TAGs, instead of using pseudo-transitions, we identified different isomers of TAG species on the basis of *sn*-position by selecting a unique parent ion/ daughter ion (Q1/Q3) combination, which is based on neutral loss of one of the *sn-*position fatty acyl chain (RCOOH) and NH_3_ from parent ion [M+NH_4_]+. For instance, the parent ion (Q1) for TAG 52:6 is 868.8 while the product ion (Q3) was derived from the remaining mass of TAG after loss of fatty acid present at one of the *sn*-position like m/z 595.5 for TAG (52:6/FA16:0) as shown in figure 2. Using this approach, we found 9 isomers for TAG species (52:6) based on composition of fatty acid present at one of the *sn*-position (figure 2a). Furthermore, MS/MS through EPI scan confirmed six of the 9 isomers of TAG 52:6 unambiguously (supplementary figure 1). The MRM library used, consists of 445 TAG species which belongs to 96 different categories of TAG based on total chain length and unsaturation. Further validation of Q3 in MS/MS experiment through IDA-EPI scan confirmed the structural characterization of Q3 ion with MS/MS spectrum for 349 putative TAG species. Using this method, we were able to identify total of 415 TAG species from 90 different categories of TAG (figure 3a). Among these 90 TAG’s, we found TAG (52:3) was the most abundant form in human plasma (figure 3b and supplementary table 2). We identified 11 isomers of TAG (52:3) among which TAG (52:3/FA16:0) was the most abundant in human plasma (figure 3b and supplementary table 3).

**Figure 2.**
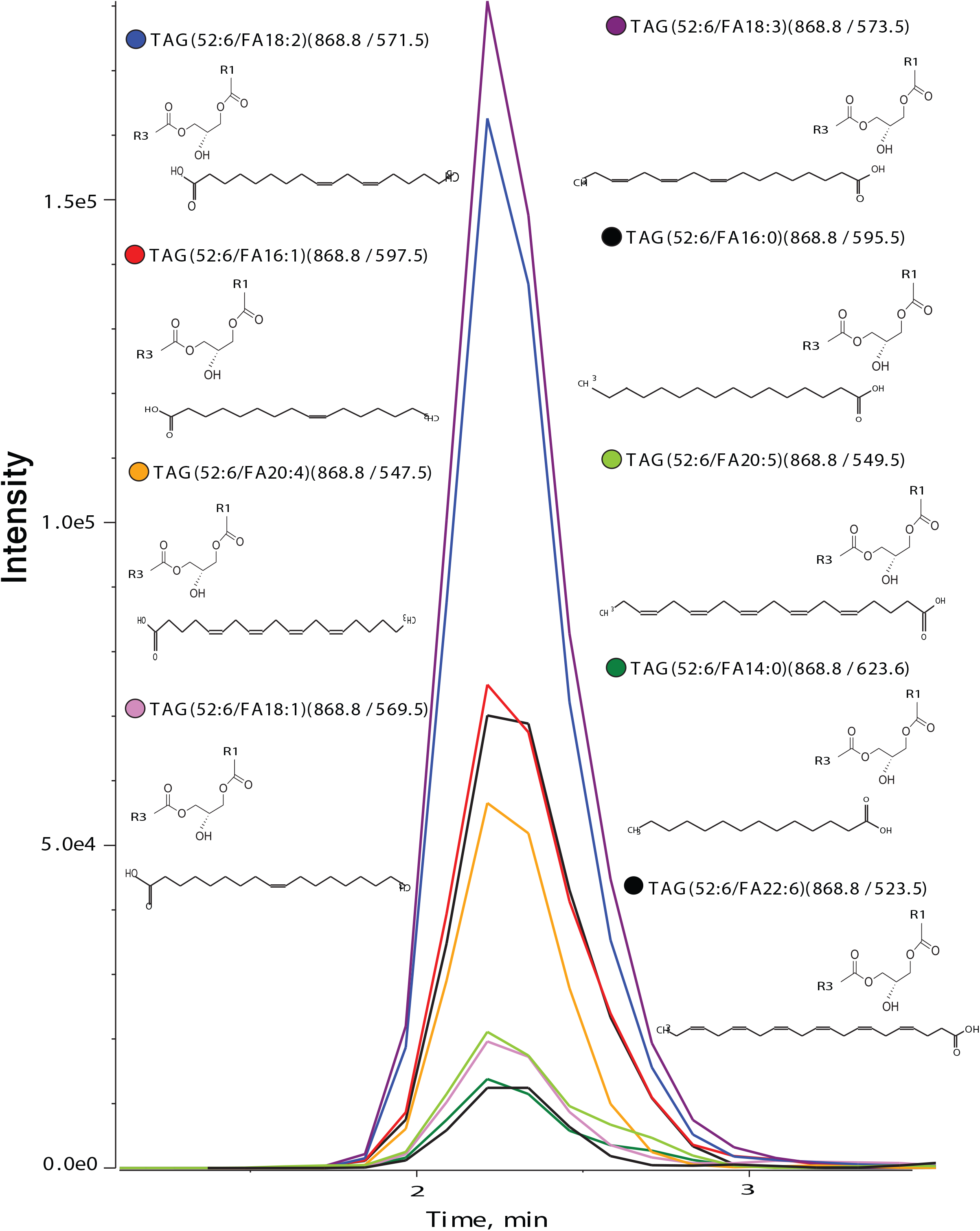
XIC (extracted ion chromatogram) of nine isomers of TAG (52:6). Parent m/z for all was 868.8 while the product m/z was derived from the remaining mass (R1+R2 with glycerol backbone) after the loss of fatty acid released from the parent ion. R1+R2 can be any composition of fatty acid which sum-up to give product ion. Different color of dot represents different isomers confirmed through IDA-EPI experiment (refer to supplementary figure 1).

**Figure 3.**
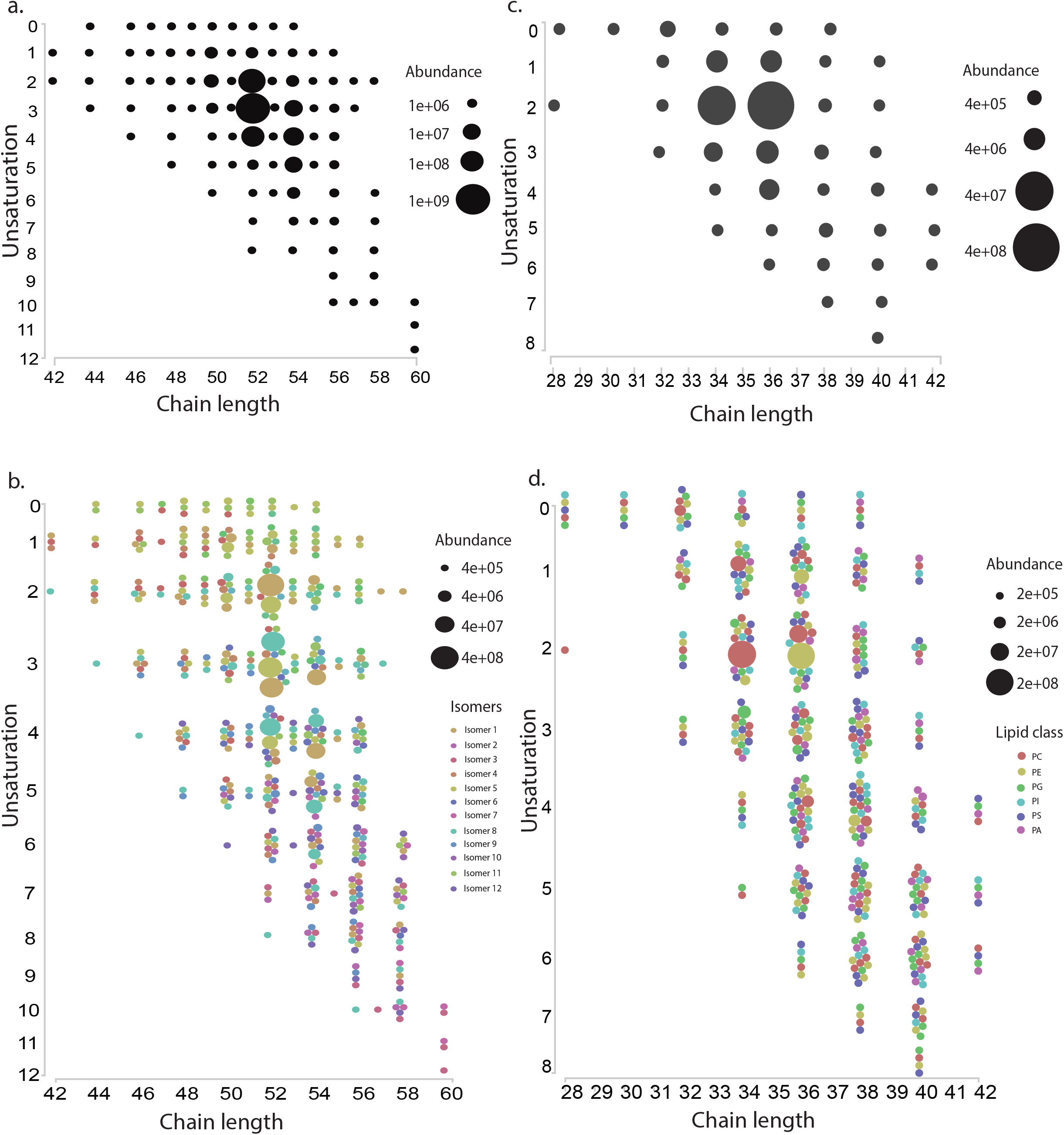
Abundance of different lipids. **a** Abundance of different TAGs on the basis of total chain length (as a function of main-chain carbon atoms) and unsaturation. **b** 415 TAG isomers were detected from 90 different categories of TAG. **c** Abundance of different phospholipids on the basis of total chain length and unsaturation. **d** Abundance of 385 phospholipids belonging to 6 classes (PC, PE, PG, PI, PS, and PA), different dots of same color represent isomers. The abundance of the difference lipids/isomers is represented by the varying size of the bubble in all the panels.

For phospholipids (PC, PE, PG, PS, PI, and PA), instead of the conventional method of using the head group loss in positive ion mode (e.g.: PC-38:4, 868.607/184.4), we used a modified approach using negative ion mode via the loss of fatty acid to identify the phospholipids at the fatty acid composition level. Using this approach, we were able to identify isomers of phospholipids within a class, like PC16:0‒22:5, PC 18:0‒20:5, PC 18:1‒20:4 and PC 18:2‒20:3 for PC 38:5 (supplementary figure 2a). Further, EPI scan for MSMS confirmed the fragmented daughter ions for the identification of three PC (38:5) isomers (supplementary figure 2b,2c and 2d). From the analysis of 455 phospholipids belonging to 6 phospholipid classes (PC, PE, PG, PI, PS, and PA) in the library, we were able to identify 385 phospholipid species. Among them, phospholipid (PC, PE, PG, PI, PS, PA) with chain length 36 with 2 unsaturation had the highest abundance (figure 3c and supplementary table 2). Within PLs, PC 34:2 has highest abundance (supplementary figure 3 and supplementary table 4). We observed three isomers of PC 34:2, among which PC (16:0/18:2) was the most abundant (figure 3d (supplementary table 3). We were also able to identify isomers of DAG (e.g. DAG 16:1/20:2 and DAG 18:1/18:2) (supplementary table 3). A list of all lipid species with their isomers and abundance in terms of area under the chromatogram is given in (supplementary table 3).

## Method validation

### Limit of blank (LoB), limit of detection (LoD), limit of quantitation (LoQ), and linear range

The raw analytical signal in blank was considered for establishing the LoB, which was determined from the area under the chromatogram for the selected transition of each lipid standards (supplementary table 5). The LoD and LoQ were obtained from the raw analytical signal (area under the chromatogram) obtained by progressively diluting the lipid standards. The LoD and LoQ were based on the average values obtained in 3 replicates, reflecting inter day variability as mentioned in the materials and methods section. A representative graph of LoD and LoQ for SM (positive mode) and PC (negative mode) is shown in figure 4a and 4b and table 1, while the values of LoD and LoQ for all the species are provided in table 1. The LoDs for all lipid classes were in range of 0.245 pmol/L – 41.961 pmol/L except for DAG (1 nmol/L). Detection limit for SM, LPC, PE, and PG were found to be in femtomolar range, while the rest were in picomolar range. The lowest LoQ was detected for PG- 0.291 pmol/L and highest for DAG- 2 nmol/L.

**Table 1.** Analytical validation of the method with lipid standards. Table1.

| Lipid<br>clas<br>s | Ion<br>mode | Number<br>of lipid<br>species | Internal standard | LoD<br>Conc.<br>(pmol/L) | LoQ<br>Conc.<br>(pmol/L) | Coefficient of<br>determination<br>(R<sup>2</sup>) |
| --- | --- | --- | --- | --- | --- | --- |
| SM | ESI+ | 12 | SM (d18:1-18:1(d9)) | 0.319 | 0.639 | 0.99 |
| CE | ESI+ | 21 | Ceramide (17:0) | 6.082 | 12.164 | 0.99 |
| Cer | ESI+ | 62 |  |  |  |  |
| TAG | ESI+ | 445 | TAG (15:0-18:1(d7)-15:0) | 17.233 | 34.466 | 0.99 |
| DAG | ESI+ | 50 | DAG (15:0-18:1(d7)) | 999.184 | 1998.367 | 0.99 |
| MA<br>G | ESI+ | 17 |  |  |  |  |
| LPC | ESI- | 16 | LPC (18:1(d7)) | 0.368 | 5.887 | 0.99 |
| PC | ESI- | 79 | PC (15:0-18:1(d7)) | 13.024 | 26.048 | 0.98 |
| LPE | ESI- | 16 | LPE (18:1(d7)) | 1.329 | 5.318 | 0.99 |
| PE | ESI- | 142 | PE (15:0-18:1(d7)) | 0.245 | 0.979 | 0.99 |
| LPG | ESI- | 16 | PG (15:0-18:1(d7)) | 0.291 | 0.291 | 0.99 |
| PG | ESI- | 78 |  |  |  |  |
| LPI | ESI- | 16 | PI (15:0-18:1(d7)) | 2.639 | 10.557 | 0.98 |
| PI | ESI- | 77 |  |  |  |  |
| LPS | ESI- | 16 | PS (15:0-18:1(d7)) | 41.961 | 167.846 | 0.99 |
| PS | ESI- | 78 |  |  |  |  |
| LPA | ESI- | 6 | PA (15:0-18:1(d7)) | 41.897 | 167.587 | 0.97 |
| PA | ESI- | 77 |  |  |  |  |

**Figure 4.**
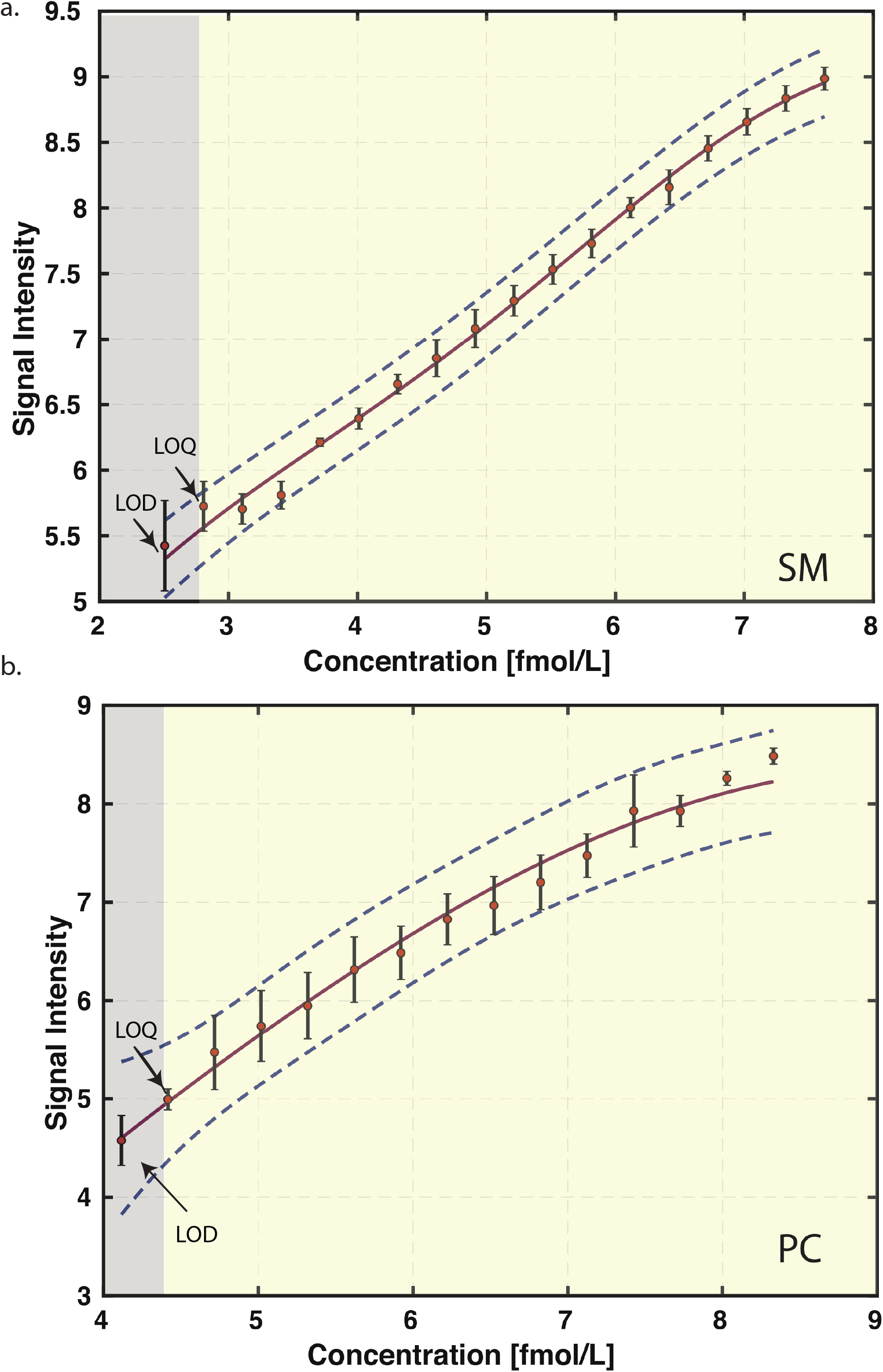
Representative graphs from positive and negative ion mode showing LoD, LoQ and coefficient of determination, x and y-axis was log transformed. **a** SM from positive ion mode and **b** PC from negative ion mode. The grey area represent the concentration below the linear range while the yellow region is indicative of linear range. The error bar represents the variance/standard deviation obtained in 3 replicates, reflecting inter day variability

The linearity of the method was checked by defining the relationship between raw values of analytical signal for each lipid standard and its concentration in presence of matrix (plasma). The linear range was determined by checking the performance limit from LoQ to the highest end of the concentration; based on the coefficient of determination (R^2^) value (table 1).

### Spike and recovery and coefficient of variation

To determine the percent recovery of all the lipid species, a known amount of lipid standards were added to plasma (matrix) before or after (spike) extraction of the lipids from the plasma. The raw area signals obtained from these two conditions were compared to determine the percentage recovery. These experiments were performed on three different days and the average percent recovery of the lipid standards is provided in figure 5a and supplementary table 6.

**Figure 5.**
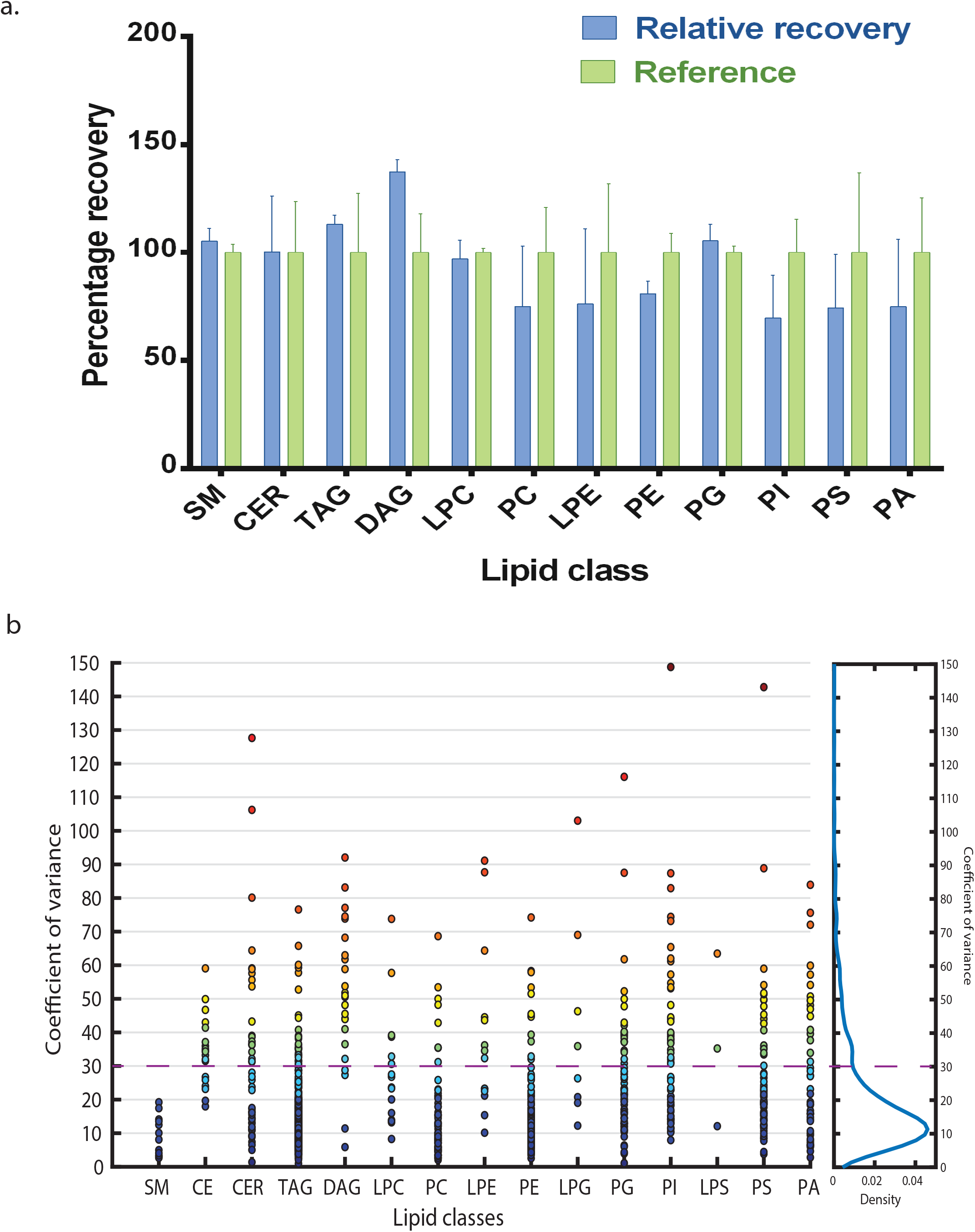
Validation of the method. **a** Spike and recovery of different lipid classes where blue bars represent the recovery of lipids when known concentration of lipid standards was spiked during extraction and green bars represents the reference (same concentration of lipid standard spiked after extraction). **b** Coefficient of variance on day 1 where 1018 lipid species from 15 lipid classes were detected (n=5). The color scale of the bubble is based on the function of coefficient of variance in the increasing order. The right hand panel represents the density function w.r.t. coefficient of variance.

To determine the coefficient of variation of all the lipid species, we extracted lipids from plasma pooled from 5 individuals. For intra batch variations, the same sample was subjected to mass spectrometric analysis 5 times. The coefficient of variation was calculated after sum normalization of raw values obtained within each class. To obtain the inter day variability; lipids were extracted from the same sample on 3 different days. A total of 1018, 952, 986 lipid species were detected on day 1, day 2, and day 3 respectively. The median CV of all the identified lipids on three different days was 15.1%, 15.5%, and 14.7% respectively. On day 1 out of 1018 lipid species, we observed 809 lipid species with CV below 30%. Of these, 259 had CV <10% and 665 had CV < 20% (figure 5b). We observed 737 and 773 lipids species on day 2 and day 3 respectively with CV<30%. In total we identified 849 lipid species with CV<30% in either of the three days, out of which 586 lipid species has been consistently detected in all days with CV<30%. The detailed table with CV for individual lipid species observed on 3 different days is given in supplementary table 7.

### Lipidomics study in normal and vitamin B12 deficient human plasma

Vitamin B_12_, is a micronutrient mainly sourced from animal products, deficiency of which has been reported to result in lipid imbalance^39^. Using this method, we attempted to identify lipid species that are altered due to vitamin B12 deficiency. There was no significant alteration in any of the lipid classes when taken as a whole between the two groups (supplementary table 8). Importantly, when individual lipid species within the classes were compared, we found that lipid species containing one of the types of omega 3 fatty acid (FA 20:5) was significantly low in plasma of vitamin B12 deficient individuals (figure 6a). This highlights the use of such a sensitive MS based-method to uncover subtle differences. In total 6 lipid species containing 20:5 fatty acids were down-regulated significantly, two of TAG and PC, one each from PE and PA. Additionally, lipid species containing a omega 6 fatty acid (FA 18:2) was significantly high in vitamin B12 deficient condition (figure 6b, supplementary table 9). These results hint at the possibility of lower ω-3: ω-6 ratio in vitamin B12 deficient individuals.

**Figure 6.**
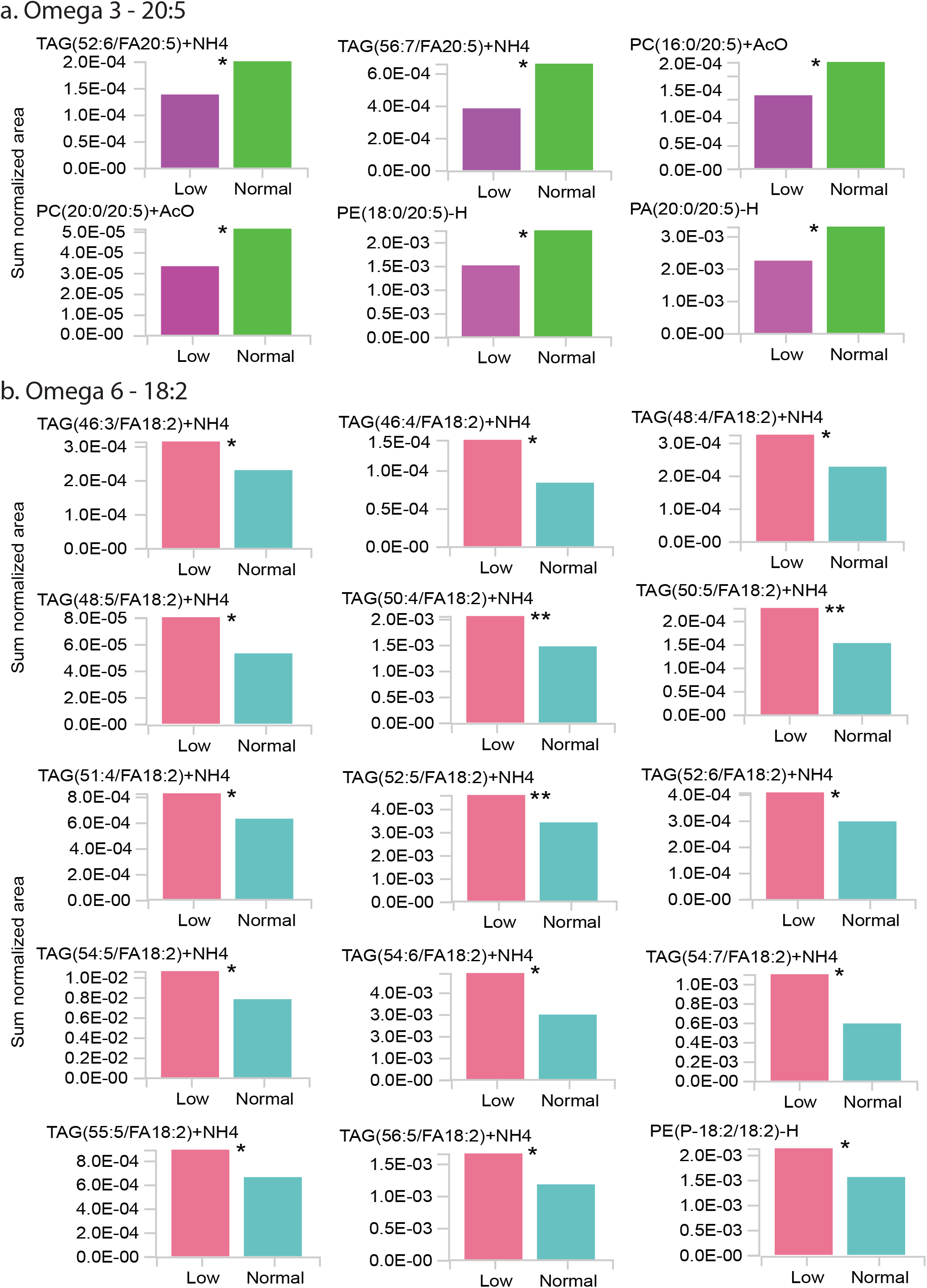
Significantly altered lipid species in vitamin B_12_ deficiency. **a** Significantly down-regulated Omega 3 fatty acid 20:5 in vitamin B12 deficiency. **b** Significantly upregulated Omega 6 fatty acid 18:2 in vitamin B12 deficient condition.

## Discussion

Lipids in general are known to be associated with the pathogenesis of various complex diseases^10^. However, the exact role played by each lipid species has not been studied in detail majorly due to the limitation in identifying individual lipid species in a large scale approach. We report a single extraction, targeted mass spectrometric method using Amide-HILIC-chromatography (scheduled MRM with variable-RTW and relative-DTW) which detects more than 1000 lipid species from 18 lipid classes including various isomers in a single MS run-time of just 24 minutes per sample injection. This method covers most of the lipid species present in human plasma with 14-22 carbons atoms and 0-6 double bonds in fatty acid chain, could enabled us to identify considerably higher number of lipid species than those reported in previous large-scale lipidomics studies^14,21,40–42^.

In this method, the MRM transitions were monitored in a particular time segment, rather than performing scans for all the lipid species during the entire run. This strategy reduces the time required for identification of the multiple transitions. We improved the coverage by additionally optimizing the assigned dwell time weightage for each lipid species, which is required especially for medium and low abundant lipid species. The dwell time for each lipid species was customized and the dwell weightage was optimized based on lipid species abundance without affecting the target scan time in each cycle. This improved peak quality with good reproducibility.

Current methods for large-scale lipid analysis can only identify the lipid classes and total fatty acyl composition of lipid species but the structure specificity is critical for studying the biological function of lipid species. Finding the composition of fatty acyl chain with respect to *sn*-position is a major limitation in large scale lipidomics studies^21,30^. Using pseudo-transitions for identifying TAG has its own disadvantages^21^. Firstly, it is based on same Q1 and Q3 m/z value (eg: 868.8/868.8), other compound which has same parent mass (Q1) and similar polarity, will also be eluted at same time and MS cannot differentiate between two compounds. So scanning unique pair of Q1/Q3 transition, where Q1 is parent ion and Q3 is characteristic daughter ion, for that compound is essential. Secondly, isomers cannot be detected as Q3 is same as Q1. Recently using a combination of photochemical reaction (Ozone-induced dissociation and ultraviolet photo-dissociation) with tandem MS, Cao et al. reported the identification of isomers for TAGs and PLs on the basis of *sn*-position and carbon-carbon double bond (C=C)^43^. Their identification also revealed the sequential loss of different fatty acyl chain based on *sn*-position, disclosing identification of different positional isomers^43^. However, a single step identification of TAG isomers in large scale studies remains a challenge due to the three fatty acyl chains with glycerol backbone, bearing no easily ionizable moiety^21,30^. We have focused on identification of structural isomers based on *sn*-position using LC-MS platform, without adding extra step to burden the analysis time and effort. We were able to detect structural isomers with respect to fatty acyl chain at *sn*-position where the neutral loss of one of the *sn-position* fatty acyl chain (RCOOH) and NH_3_ from parent ion (M+NH_4_+) makes their detection possible. Detection was purely based on assigning a unique combination of Q1/Q3 for structural isomer of TAG species (figure 2a); however, one of the limitations of this method is the inability to assign fatty acyl group (*sn1*, *sn2*, or *sn3*) to their respective *sn*-position. Hence, the three fatty acyl chains are represented by the adding the number of carbon atoms and unsaturation level (e.g., TAG (52:6) and the identified fatty acid at one of the *sn*-position (e.g., FA-14:0) is represented by TAG (52:6/FA14:0).

The LoD for various lipid species in our method was between 0.245 fmol/L – 41.96 pmol/L which was better than or similar to previously reported LoD utilizing different LC-MS platforms^21,27,31,41,42^ and similar to a previously reported large scale lipidomics method using supercritical fluid-scheduled MRM (5‒1,000 fmol/L)^21^. The LoQ in previously reported methods were in between nmol to μmol/L range while we have observed much lower LoQs (0.291 pmol/L to 167.84 pmol/L)^21^. Apart from this, the calculation of limits was based on mean raw analytical signal and SD, which gives better idea about the method, without any false detection hope (or lower detection limits). In our method, DAG has highest LoD and LoQ of 1 nmol/L and 2nmol/L respectively, which was still lower as compared to the previously reported methods for targeted analysis^21^. The linearity of our method was found to be comparable to previous lipidomics methods^21,27,41^.

The recovery of lipid species in our method was in the range of 69.75 % - 113.19 %, except DAG - 137.5%, which were within the generally accepted range for quantification and is comparable with other lipidomics studies^21,27^.

A major challenge in lipidomics experiments have been the high variability in the signals and even the “shared reference materials harmonize lipidomics across MS-based detection platforms and laboratories” have shown that most lipid species showed large variability (CV) between 30% to 80%^44^. However variability for endogenous lipid species that were normalized to the corresponding stable isotope-labelled analogue were lower than 30%^40,44^. In this method, we used sum normalization (although we are not addressing batch effect in this study) and found that 849 lipid species had a CV <30%^40^. Overall, the median CV of our method (15.1%, 15.5%, and 14.7%), was similar to or better than the previous reports^21,27,31^. In addition, we have also reported species-specific CV. It should be noted that most of the large scale lipidomics studies previously done reports the median or average CV of the method but not the species-specific CV^14,21,27,31^.

### Lipidomics study in normal and vitamin B12 deficient human plasma-

Using the method developed we identified lipid species that are altered in individuals with vitamin B_12_ deficiency. Vitamin B_12_ is a cofactor of methyl malonyl CoA mutase and controls the transfer of long-chain fatty acyl-CoA into the mitochondria^45^. Deficiency of vitamin B_12_ results in accumulation of methylmalonyl CoA increasing lipogenesis via inhibition of beta-oxidation.

In the last decade, several studies revealed that vitamin B_12_ deficiency causes alteration in the lipid profile through changes in lipid metabolism, either by modulating their synthesis or its transport^46^. In particular, the effects of vitamin B_12_ on omega 3 fatty acid and phospholipid metabolism have received much attention. Khaire A et al., found that vitamin B_12_ deficiency increased cholesterol levels but reduced docosahexaenoic acid (DHA-omega 3)^47^. An imbalance in maternal micronutrients (folic acid, vitamin B_12_) in Wistar rats increased maternal oxidative stress, decreases placental and pup brain DHA levels, and decreases placental global methylation levels^48,49^. Although various studies have shown that B_12_ deficiency results in adverse lipid profile as well as pathophysiological changes linked to CAD, type 2 diabetes mellitus and atherosclerosis, very few studies have independently investigated the effect of vitamin B_12_ status on changes in human plasma lipid among apparently healthy population^50–52^. Importantly the lipid species that are altered because of the vitamin deficiency are still not yet well understood.

To our knowledge, this is the first study to identify lipids with a significantly decreased ω-3 fatty acid (20:5) chains and increased ω-6 (18:2) chains, which might alter/increased ω-6 to ω-3 fatty acid ratio in human plasma in relation to vitamin B_12_ deficiency and may promote development of many chronic diseases. Most importantly we found that although there was no significant alteration in the lipid classes, individual lipid species varied in vitamin B_12_ deficient individuals clearly demonstrating the utility of identifying lipid species.

The application of scheduled MRM with variable-RTW and relative-DTW enabled large-scale quantification of lipid species in a single-run as compared to unscheduled/scheduled/dynamic MRM. With this combinatorial approach, we were able to detect more than 1000 lipid species in plasma, including isomers of TAG, DAG and PL’s. Additionally we validated the retention time through MSMS analysis in IDA-EPI scan mode by matching fragmented daughter ion from MSMS spectrum to putative lipid species structure. It should be noted that the MRMs currently used were specific for plasma and may not be ideal for other biological systems. Therefore, for developing a separate MRM panel may be required for each system. To the best of our knowledge this is the largest number of lipid species identified till date in a single experiment. A comprehensive identification of structural isomers in large-scale lipid method proves to be critical for studying the important biological functions of lipids.

## Acknowledgement

The authors would like to thank Dr. Mainak Dutta from BITS Dubai, Mrs. Akanksha Singh and Dr. Christei Hunter of Sciex for their invaluable inputs and suggestions in shaping this study. Akash Kumar Bhaskar and Salwa Naushin would like to thank CSIR for their fellowship. The study was funded by Council of Scientific and Industrial Research (CARDIOMED MLP 0122 and MLP 1811).

## Materials and Methods

### Chemicals and reagents

MS-grade acetonitrile, methanol, water, 2-propanol (IPA) and HPLC-grade dichloromethane (DCM), were purchased from Biosolve (Dieuze, France); ammonium acetate and ethanol were obtained from Merck (Merck & Co. Inc., Kenilworth, NJ, USA). Lipid internal standards used in the study : SM (d18:1-18:1(d9)), TAG (15:0-18:1(d7)-15:0), DAG (15:0-18:1(d7)), LPC (18:1(d7)), PC (15:0-18:1(d7)), LPE (18:1(d7)), PE (15:0-18:1(d7)), PG (15:0-18:1(d7)), PI (15:0-18:1(d7)), PS (15:0-18:1(d7), PA (15:0-18:1(d7)) in the form of Splash mix and ceramide (17:0) were purchased from Avanti polar (Alabaster, Alabama, USA).

### Lipid extraction from human plasma

We used *a* modified *Bligh and Dyer method* using Dichloromethane/methanol/water *(2:2:1 v/v)*. The study was approved by institutional ethical committee of CSIR-IGIB. Human plasma (10 μL) was mixed with 490 μL of water (in glass tube) and incubated on ice for 10 minutes. Lipid internal standard mixes (5 μL, consisting of splashmix and ceramide) was added to a mixture of methanol (2 mL) and dichloromethane (1 mL); the mixture was vortexed and allowed to incubate for 30 minutes at room temperature. After incubation, 500 μL water and 1 mL dichloromethane was added to the solution and vortexed for 5 seconds. The mixture was centrifuged at 300 g for 10 minutes when there was a phase separation. The lower organic layer was collected into a fresh glass tube. 2 mL dichloromethane was added to remaining mixture in extraction tube and centrifuged again to collect the lower layer. The previous step was repeated one more time. Solvent was evaporated in vacuum dryer at 25 °C and the lipids were resuspended in 100μl of ethanol; vortexed for 5 minutes, sonicated for 10 minutes and again vortexed for 5 minutes. The suspension was transferred to polypropylene auto sampler vials and subjected to LC-MS run.

### Liquid chromatography-Mass spectrometry

We used an Exion LC system with a Waters AQUITY UPLC BEH HILIC XBridge Amide column (3.5 μm, 4.6 x 150 mm) for chromatographic separation. The oven temperature was set at 35°C and the auto sampler was set at 4°C. Lipids were separated using buffer A (95% acetonitrile with 10mM ammonium acetate, pH-8.2) and buffer B (50% acetonitrile with 10mM ammonium acetate, pH-8.2) with following gradient: with a flow rate of 0.7 ml/minute, buffer B was increased from 0.1% to 6% in 6 minutes, increased to 25% buffer B in next 4 minutes. In the next 1 minute buffer B was ramped up to 98%, further increased to 100% in the next 2 minutes, and held at the same concentration and flow rate for 30 seconds. Flow rate was increased from 0.7 ml/min to 1.5 ml/min and 100% buffer B was maintained for the next 5.1 minutes. Buffer B was brought to initial 0.1% concentration in 0.1 minute and column was equilibrated at the same concentration and flow for 4.3 minutes before flow rate was brought to initial 0.7 ml/minute in next 30 seconds and maintained at the same till the end of 24 minutes gradient. Additionally the separation system was equilibrated for 3 minute for subsequent runs.

Sciex QTRAP 6500+ LC/MS/MS system in low mass range, Turbo source with Electrospray Ionization (ESI) probe was used with the following parameters; curtain gas (CUR): 35 psi, temperature (TEM): 500 degree, source gas 1(GS1): 50 and source gas 2 (GS2): 60 psi, ionization voltage (IS): 5500 for positive mode and IS: −4500 for negative mode, target scan time: 0.5 sec, scan speed: 10 Da/s, settling time: 5.0000 msec and MR pause: 5.0070 msec. Acquisition was done using Analyst 1.6.3 software.

### Method development

For identification and relative quantification of all the lipid species, theoretical MRM library were generated using LIPIDMAPS (https://www.lipidmaps.org/). Using internal standards from different lipid classes, the MRM parameters (collision energy, declusturing potential, cell exit potential, and entrance potential) were optimized for 1224 lipid species which belonged to 18 lipid classes - sphingomyelin (SM), ceramide (Cer), cholesterol ester (CE), Monoacylglycerol (MAG), diacylglycerol (DAG), Triacylglycerol (TAG), lysophosphatidic acid (LPA), phosphotidic acid (PA), lysophosphatidylcholine (LPC), phosphatidylcholine (PC), lysophosphatidylethanolamine (LPE), phosphatidylethanolamine (PE), lysophosphatidylinositol (LPI), phosphatidylinositol (PI), lysophosphatidylglycerol (LPG), phosphatidylglycerol (PG), lysophosphatidylserine, and (LPS), phosphatidylserine (PS) (supplementary table-1).

The MRM library consisted of 1236 transitions including 12 internal standards, of which 611 species were identified in positive mode (SM, CE, Cer, TAG, DAG, MAG) and 625 identified in negative mode (Phospholipids and lysophospholipids). The current MRM panel covers major lipid classes and categories having fatty acids with 14-22 carbons and 0-6 double bonds per fatty acyl chain. Transitions were distributed into multiple unscheduled MRM method and the relative retention time of each transition was determined with respect to their respective internal standards through Amide-HILIC column. Furthermore, the retention time validation was done by performing MS/MS experiment using Information dependent acquisition (IDA) with enhanced product ion scan (EPI) of specific ions in unscheduled MRM for each lipid class. MS/MS analysis in EPI mode was based on the conventional triple quadrupole ion path property of an ion-trap for the third quadrupole. The basic parameters were kept the same as mentioned in MRM experiment. MS/MS spectra were compared with MS/MS information from LIPID MAPS (http://www.lipidmaps.org/) to verify the structures of the putative lipid species and predicting the structure from MS/MS spectra based on specific cleavage rules for lipids.

### Retention time window and Dwell time weightage

Using *sMRM* Builder (https:// https://sciex.com/), an Excel based tool from Sciex, the variable retention time window and variable dwell time weightage for all transitions were optimized. The principle on which the tool works is based on the width and intensity of the chromatographic peak. With variable retention time window width, each MRM transition can have its own RT window. Wider windows are assigned to analytes that show higher run to run variation or have broader peak widths. Variable dwell times were assigned to improve the signal to noise ratio of MRM transitions based on the abundance of the analyte in the sample- higher dwell time weightage assigned for analytes with low abundance (supplementary table 1). Dwell time for each species were assigned based on this weight which maintains the cycle time and optimizes the signal to noise ratio for low abundant peaks. Detailed for optimized parameters is given in supplementary table 1.

### Limit of Detection and Quantitation

The limits of detection and quantitation were derived from peak area of known amounts of lipid internal standards added to lipid extract from human plasma (matrix):

The master mix of lipid internal standards was prepared from splashmix and ceramide (17:0) having following concentrations: SM (41.86 nmol), Cer (24.91 nmol), TAG (70.59 nmol), DAG (15.99 nmol), LPC (48.23 nmol), PC (213.38 μmol), LPE (10.89 nmol), PE (8.02 nmol), PG (38.09 nmol), PI (5.40 nmol), PS (10.74 nmol), PA (10.73 nmol).

Limit of Blank- was defined as the average (based on triplicate experiments) signal found only in matrix (without internal standards; blank). LoB was calculated using mean and standard deviation from plasma matrix:

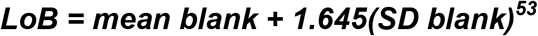

The raw analytical signal obtained for standards from plasma lipid extract (spiked with standards) was used to estimate the LoD and LoQ, using the following formula:

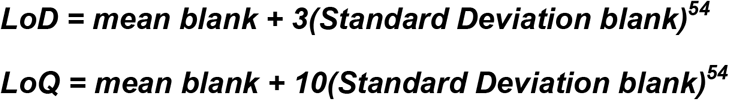

The standard solution was diluted serially with matrix and the lipid standards were run in the following concentration ranges: 319.39 fmol- 41.86 nmol for SM, 190.06 fmol- 24.91 nmol for Cer, 538.53 fmol-70.59 nmol for TAG, 121.97 fmol- 15.99 nmol for DAG, 367.9633086 fmol- 48.23 nmol for LPC, 1.63 pmol-213.38 μmol for PC, 83.09 fmol-10.89 nmol for LPE, 61.16 fmol- 8.02 nmol for PE, 290.59 fmol- 38.09 nmol for PG, 41.24 fmol- 5.40 nmol for PI, 81.96 fmol- 10.74 nmol for PS, 81.83 fmol- 10.73 nmol for PA. The lowest concentration which has signal more than the estimated method limits (based on above formula) was considered as LoD and LoQ. The mean and standard deviation was calculated from 3 replicates. Linearity was represented by R^2^, where LoQ was taken as the lowest calibrator concentration for each lipid standards.

### Spike and recovery and coefficient of variance

Extraction recovery for the method was measured by comparing the peak area of matrix extract spiked with standards before and after extraction. For this, 5uL of lipid internal standard mix (standard mix: lipid extract resuspension volume :: *1:20 v/v*) was used. The percentage recovery and relative standard deviation was calculated from 3 biological replicates.

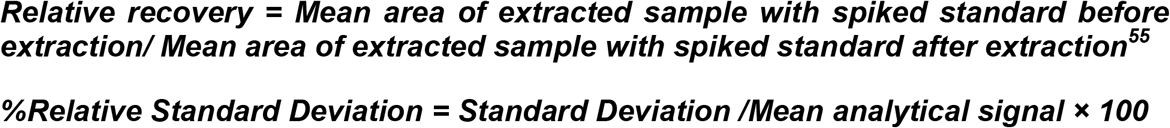

Coefficient of variance (CV) of the method was determined by observing individual lipid species variation within batch. The intra-batch variation was assessed by analyzing 5 technical replicates of lipids extracted from pooled plasma. CV values were only calculated for those lipid species which has carry over less than 20% and present in at least 3 replicates^56^. Inter day variability for each lipid species was determined by analyzing lipids on 3 different days from a stock of pooled plasma. The CV values were reported for 3 different days (n=5, technical replicates) after sum-normalization within lipid class.

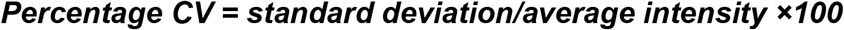

### Alteration of plasma lipids due to vitamin B12 deficiency

#### Study population

The study (which was a part of a larger study), was designed to identify plasma lipids that were altered due to vitamin B12 deficiency. Apparently healthy individuals were classified in two groups based on their plasma vitamin B12 levels. An informed consent was obtained from the participants. The study was approved by institutional ethical committee of CSIR-IGIB. Individuals with vitamin B12 values less than 150 pg/ml, were considered to be vitamin B12 deficient and those with levels between 400-800 pg/ml were considered be in the normal range. Lipids from plasma were extracted as described above. For this study, plasma of 95 individuals (48 with B12 deficiency and 47 with normal plasma vitamin B12 levels) were used. Lipids that had a CV<30% and that were altered by more than 1.3 folds with p<0.05 were considered to be significantly altered between the two groups.

#### Data analysis

The .wiff files for relative quantitation were processed in MultiQuant 3.0.2 and for the identification of different lipid species; MS/MS spectrum matching with the structure of putative lipid species using .mol file was done using Peakview 2.0.1. Statistical analysis was done using Excel. Figures were drawn using MATLAB (MATLAB, 2010. *version 7.10.0 (R2010a)*, Natick, Massachusetts: The MathWorks Inc.), Raw graph (https://rawgraphs.io) and GraphPad Prism version 6.0.

## References

1 Smilowitz, J. T. et al. Nutritional lipidomics: molecular metabolism, analytics, and diagnostics. Molecular nutrition & food research 57, 1319–1335 (2013).

2 Muro, E., Atilla-Gokcumen, G. E. & Eggert, U. S. Lipids in cell biology: how can we understand them better? Molecular biology of the cell 25, 1819–1823 (2014).

3 Yáñez-Mó, M. et al. Biological properties of extracellular vesicles and their physiological functions. Journal of extracellular vesicles 4, 27066 (2015).

4 Van Meer, G., Voelker, D. R. & Feigenson, G. W. Membrane lipids: where they are and how they behave. Nature reviews Molecular cell biology 9, 112–124 (2008).

5 Glomset, J. A. Protein-lipid interactions on the surfaces of cell membranes. Curr. Opin. Struct. Biol 9, 425–427 (1999).

6 Ye, R., Onodera, T. & Scherer, P. E. Lipotoxicity and β cell maintenance in obesity and type 2 diabetes. Journal of the Endocrine Society 3, 617–631 (2019).

7 Fu, S. et al. Aberrant lipid metabolism disrupts calcium homeostasis causing liver endoplasmic reticulum stress in obesity. Nature 473, 528–531 (2011).

8 Yang, M., Zhang, Y. & Ren, J. Autophagic regulation of lipid homeostasis in cardiometabolic syndrome. Frontiers in cardiovascular medicine 5, 38 (2018).

9 Beloribi-Djefaflia, S., Vasseur, S. & Guillaumond, F. Lipid metabolic reprogramming in cancer cells. Oncogenesis 5, e189–e189 (2016).

10 Wymann, M. P. & Schneiter, R. Lipid signalling in disease. Nature reviews Molecular cell biology 9, 162–176 (2008).

11 Quehenberger, O. & Dennis, E. A. The human plasma lipidome. New England Journal of Medicine 365, 1812–1823 (2011).

12 Shevchenko, A. & Simons, K. Lipidomics: coming to grips with lipid diversity. Nature reviews Molecular cell biology 11, 593–598 (2010).

13 Sud, M. et al. Lmsd: Lipid maps structure database. Nucleic acids research 35, D527–D532 (2007).

14 Pradas, I. et al. Lipidomics reveals a tissue-specific fingerprint. Frontiers in physiology 9, 1165 (2018).

15 van Meer, G. Cellular lipidomics. The EMBO journal 24, 3159–3165 (2005).

16 Brügger, B., Erben, G., Sandhoff, R., Wieland, F. T. & Lehmann, W. D. Quantitative analysis of biological membrane lipids at the low picomole level by nano-electrospray ionization tandem mass spectrometry. Proceedings of the National Academy of Sciences 94, 2339–2344 (1997).

17 Wu, Z., Shon, J. C. & Liu, K.-H. Mass spectrometry-based lipidomics and its application to biomedical research. Journal of lifestyle medicine 4, 17 (2014).

18 Wenk, M. R. The emerging field of lipidomics. Nature reviews Drug discovery 4, 594–610 (2005).

19 Han, X. & Gross, R. W. Global analyses of cellular lipidomes directly from crude extracts of biological samples by ESI mass spectrometry a bridge to lipidomics. Journal of lipid research 44, 1071–1079 (2003).

20 Kirkwood, J. S., Maier, C. & Stevens, J. F. Simultaneous, untargeted metabolic profiling of polar and nonpolar metabolites by LC-Q-TOF Mass Spectrometry. Current protocols in toxicology 56, 4.39. 31–34.39. 12 (2013).

21 Takeda, H. et al. Widely-targeted quantitative lipidomics method by supercritical fluid chromatography triple quadrupole mass spectrometry. Journal of lipid research 59, 1283–1293 (2018).

22 Contrepois, K. et al. Cross-platform comparison of untargeted and targeted lipidomics approaches on aging mouse plasma. Scientific reports 8, 1–9 (2018).

23 Khan, M. J. et al. Evaluating a targeted multiple reaction monitoring approach to global untargeted lipidomic analyses of human plasma. Rapid Communications in Mass Spectrometry 34, e8911 (2020).

24 Dekker, B. Reduce complexity by choosing your reactions. Nature Methods 12, 16–16 (2015).

25 Mao, C. et al. Cloning and Characterization of a Mouse Endoplasmic Reticulum Alkaline Ceramidase AN ENZYME THAT PREFERENTIALLY REGULATES METABOLISM OF VERY LONG CHAIN CERAMIDES. Journal of Biological Chemistry 278, 31184–31191 (2003).

26 Song, J. et al. A highly efficient, high-throughput lipidomics platform for the quantitative detection of eicosanoids in human whole blood. Analytical biochemistry 433, 181–188 (2013).

27 Weir, J. M. et al. Plasma lipid profiling in a large population-based cohort. Journal of lipid research 54, 2898–2908 (2013).

28 Zhang, W. et al. Online photochemical derivatization enables comprehensive mass spectrometric analysis of unsaturated phospholipid isomers. Nature communications 10, 1–9 (2019).

29 Thomas, M. C., Mitchell, T. W. & Blanksby, S. J. Ozonolysis of phospholipid double bonds during electrospray ionization: A new tool for structure determination. Journal of the American Chemical Society 128, 58–59 (2006).

30 Baba, T., Campbell, J. L., Le Blanc, J. Y. & Baker, P. R. Structural identification of triacylglycerol isomers using electron impact excitation of ions from organics (EIEIO). Journal of lipid research 57, 2015–2027 (2016).

31 Tabassum, R. et al. Genetic architecture of human plasma lipidome and its link to cardiovascular disease. Nature communications 10, 1–14 (2019).

32 Li, J. et al. Large-scaled human serum sphingolipid profiling by using reversed-phase liquid chromatography coupled with dynamic multiple reaction monitoring of mass spectrometry: method development and application in hepatocellular carcinoma. Journal of chromatography A 1320, 103–110 (2013).

33 Liang, J. et al. A dynamic multiple reaction monitoring method for the multiple components quantification of complex traditional Chinese medicine preparations: Niuhuang Shangqing pill as an example. Journal of Chromatography a 1294, 58–69 (2013).

34 Rao, Z. et al. Development of a dynamic multiple reaction monitoring method for determination of digoxin and six active components of Ginkgo biloba leaf extract in rat plasma. Journal of Chromatography B 959, 27–35 (2014).

35 Andrade, G. et al. Liquid chromatography–electrospray ionization tandem mass spectrometry and dynamic multiple reaction monitoring method for determining multiple pesticide residues in tomato. Food chemistry 175, 57–65 (2015).

36 Jia, Z.-X., Zhang, J.-L., Shen, C.-P. & Ma, L. Profile and quantification of human stratum corneum ceramides by normal-phase liquid chromatography coupled with dynamic multiple reaction monitoring of mass spectrometry: development of targeted lipidomic method and application to human stratum corneum of different age groups. Analytical and bioanalytical chemistry 408, 6623–6636 (2016).

37 Shah, I., Petroczi, A., Uvacsek, M., Ránky, M. & Naughton, D. P. Hair-based rapid analyses for multiple drugs in forensics and doping: application of dynamic multiple reaction monitoring with LC-MS/MS. Chemistry Central Journal 8, 73 (2014).

38 Xu, G., Amicucci, M. J., Cheng, Z., Galermo, A. G. & Lebrilla, C. B. Revisiting monosaccharide analysis–quantitation of a comprehensive set of monosaccharides using dynamic multiple reaction monitoring. Analyst 143, 200–207 (2018).

39 Kumar, K. A. et al. Maternal dietary folate and/or vitamin B12 restrictions alter body composition (adiposity) and lipid metabolism in Wistar rat offspring. The Journal of nutritional biochemistry 24, 25–31 (2013).

40 Medina, J. et al. Single-Step Extraction Coupled with Targeted HILIC-MS/MS Approach for Comprehensive Analysis of Human Plasma Lipidome and Polar Metabolome. Metabolites 10, 495 (2020).

41 Rampler, E. et al. Simultaneous non-polar and polar lipid analysis by on-line combination of HILIC, RP and high resolution MS. Analyst 143, 1250–1258 (2018).

42 Schoeny, H. et al. Preparative supercritical fluid chromatography for lipid class fractionation—a novel strategy in high-resolution mass spectrometry based lipidomics. Analytical and bioanalytical chemistry, 1–10 (2020).

43 Cao, W. et al. Large-scale lipid analysis with C= C location and sn-position isomer resolving power. Nature communications 11, 1–11 (2020).

44 Triebl, A. et al. Shared reference materials harmonize lipidomics across MS-based detection platforms and laboratories. Journal of lipid research 61, 105–115 (2020).

45 Green, R. et al. Vitamin B 12 deficiency. Nature reviews Disease primers 3, 1–20 (2017).

46 Saraswathy, K. N., Joshi, S., Yadav, S. & Garg, P. R. Metabolic distress in lipid & one carbon metabolic pathway through low vitamin B-12: a population based study from North India. Lipids in health and disease 17, 96 (2018).

47 Khaire, A., Rathod, R., Kale, A. & Joshi, S. Vitamin B12 and omega-3 fatty acids together regulate lipid metabolism in Wistar rats. Prostaglandins, Leukotrienes and Essential Fatty Acids 99, 7–17 (2015).

48 Kulkarni, A. et al. Effects of altered maternal folic acid, vitamin B 12 and docosahexaenoic acid on placental global DNA methylation patterns in Wistar rats. PLoS One 6, e17706 (2011).

49 Roy, S. et al. Maternal micronutrients (folic acid and vitamin B12) and omega 3 fatty acids: implications for neurodevelopmental risk in the rat offspring. Brain and Development 34, 64–71 (2012).

50 Adaikalakoteswari, A. et al. Vitamin B12 deficiency is associated with adverse lipid profile in Europeans and Indians with type 2 diabetes. Cardiovascular diabetology 13, 129 (2014).

51 Kumar, J. et al. Vitamin B12 deficiency is associated with coronary artery disease in an Indian population. Clinical Chemistry and Laboratory Medicine (CCLM) 47, 334–338 (2009).

52 Mahalle, N., Kulkarni, M. V., Garg, M. K. & Naik, S. S. Vitamin B12 deficiency and hyperhomocysteinemia as correlates of cardiovascular risk factors in Indian subjects with coronary artery disease. Journal of cardiology 61, 289–294 (2013).

53 Armbruster, D. A. & Pry, T. Limit of blank, limit of detection and limit of quantitation. The clinical biochemist reviews 29, S49 (2008).

54 Armbruster, D. A., Tillman, M. D. & Hubbs, L. M. Limit of detection (LQD)/limit of quantitation (LOQ): comparison of the empirical and the statistical methods exemplified with GC-MS assays of abused drugs. Clinical chemistry 40, 1233–1238 (1994).

55 Rower, J. E., Bushman, L. R., Hammond, K. P., Kadam, R. S. & Aquilante, C. L. Validation of an LC/MS method for the determination of gemfibrozil in human plasma and its application to a pharmacokinetic study. Biomedical Chromatography 24, 1300–1308 (2010).

56 van Amsterdam, P. et al. The European Bioanalysis Forum community’s evaluation, interpretation and implementation of the European Medicines Agency guideline on Bioanalytical Method Validation. Bioanalysis 5, 645–659 (2013).

