## Supplementary- table 1 for "A high throughput lipidomics method using *scheduled* multiple reaction monitoring"

**Supplementary figure and table:**

**Supplementary Figure 1 MS/MS through IDA-EPI experiment confirms six of the nine isomers TAG (52:6).**

**Supplementary Figure 2 Isomers of PC (38:5). A(i)** XIC of four isomers of PC (38:5). **A(ii)** MS/MS of PC (16:0/22:5), **A(iii)** MSMS of PC (18:0/20:5) and **A(iv)** MSMS of PC (18:1/20:4).

**Supplementary Figure 3** Lipid class wise abundance of phospholipids on the basis of total chain length and un-saturation showing PC 34:2 has highest abundance.

**Supplementary- table 1** Optimized MRM parameters for 1224 transitions and 12 internal standards.

**Supplementary- table 2** TAG and PL abundance (total chain length and unsaturation wise).

**Supplementary- table 3** All lipid species and their abundance.

**Supplementary- table 4** Class-wise abundance of six classes of phospholipids based on total chain length and un-saturation.

**Supplementary- table 5** Limit of blank for different lipid classes.

**Supplementary-table 6** Spike and recovery.

**Supplementary table 7** Detailed CV(n=5) table for 3 different days.

**Supplementary table 8** Fold change and p-value of different lipid classes in vitamin $B^{12}$ deficient study.

**Supplementary table 9** Fold change and p value for individual lipid species in vitamin $B^{12}$ deficiency.

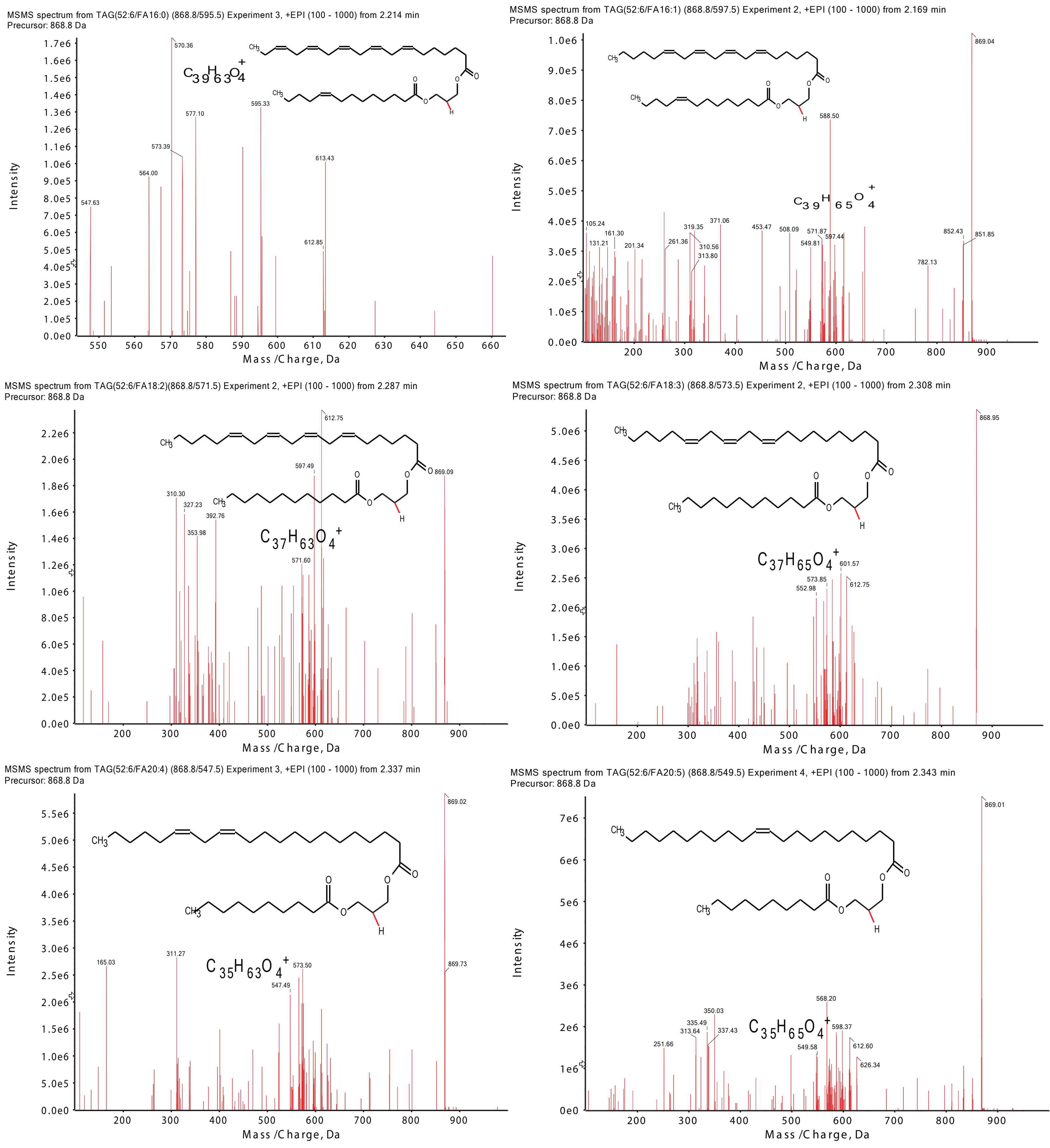

Supplementary figure 1

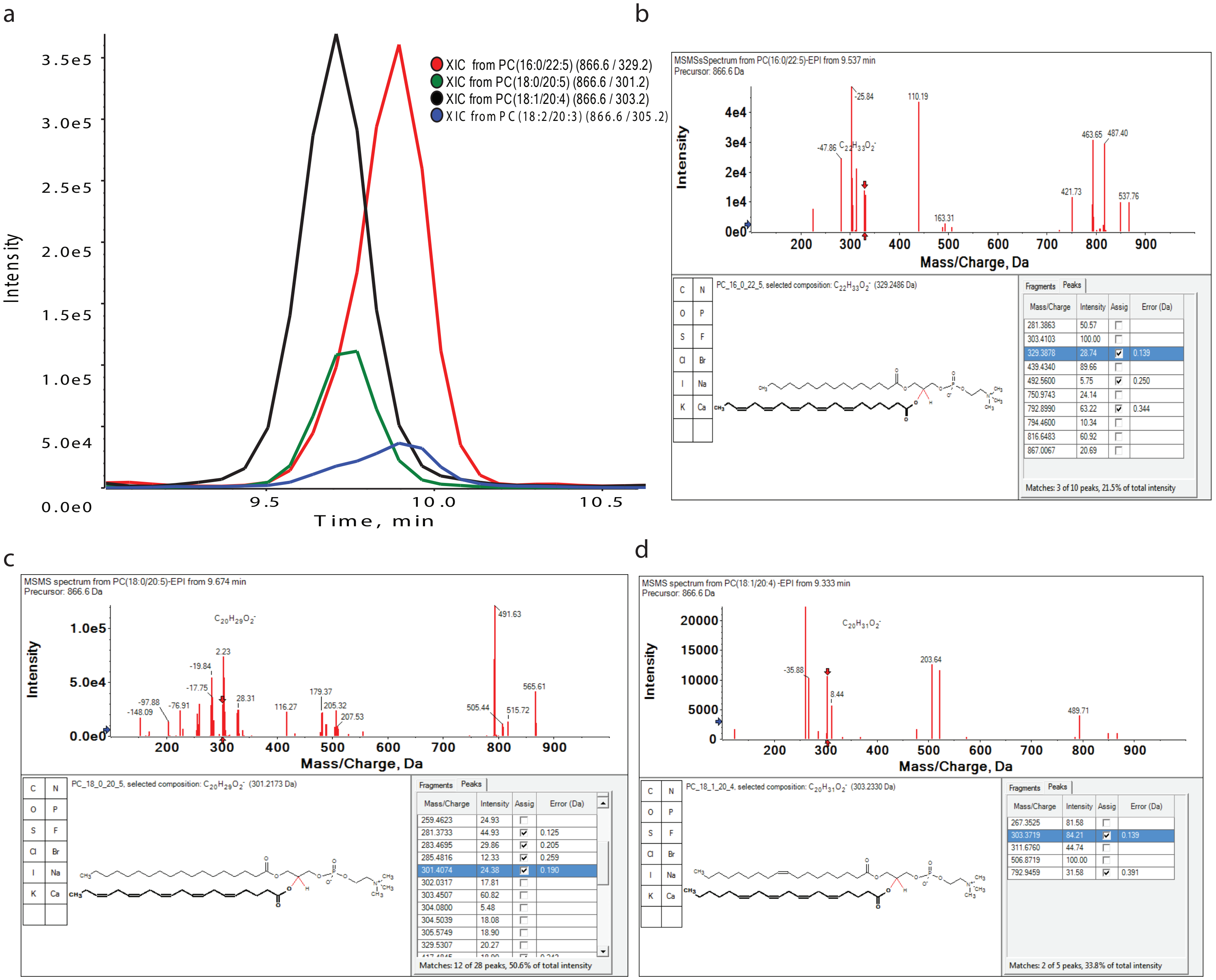

Supplementary figure 2

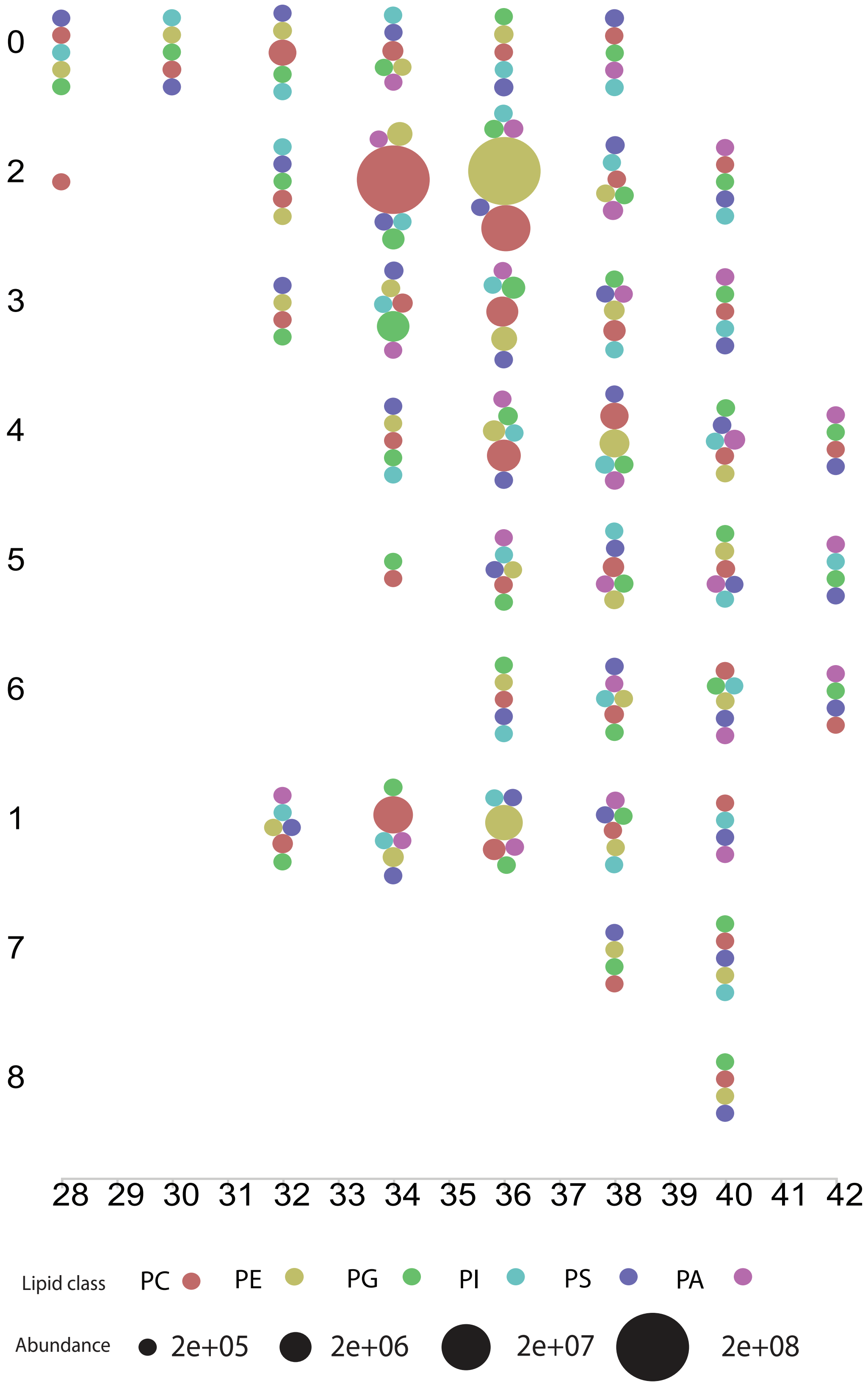

Supplementary figure 3

Supplementary- table 1:

| S.No. | Q1 | Q3 | Retention time | ID | DP | EP | CE | CXP | WINDOW (SEC) | DWELL WEGHT |
|---|---|---|---|---|---|---|---|---|---|---|
| 1 | 675.5 | 184.1 | 12.03 | SM(14:0)+H | 80 | 10 | 43 | 15 | 34.3 | 1 |
| 2 | 703.6 | 184.1 | 11.96 | SM(16:0)+H | 80 | 10 | 43 | 15 | 942.9 | 9 |
| 3 | 731.6 | 184.1 | 11.87 | SM(18:0)+H | 80 | 10 | 43 | 15 | 36.8 | 1 |
| 4 | 729.6 | 184.1 | 11.88 | SM(18:1)+H | 80 | 10 | 43 | 15 | 36.1 | 1 |
| 5 | 759.6 | 184.1 | 9.83 | SM(20:0)+H | 80 | 10 | 43 | 15 | 36 | 1 |
| 6 | 757.6 | 184.1 | 10.52 | SM(20:1)+H | 80 | 10 | 43 | 15 | 49.7 | 1 |
| 7 | 787.7 | 184.1 | 9.53 | SM(22:0)+H | 80 | 10 | 43 | 15 | 73.2 | 1 |
| 8 | 785.7 | 184.1 | 9.4 | SM(22:1)+H | 80 | 10 | 43 | 15 | 55.9 | 1 |
| 9 | 815.7 | 184.1 | 11.66 | SM(24:0)+H | 80 | 10 | 43 | 15 | 40.8 | 1 |
| 10 | 813.7 | 184.1 | 11.67 | SM(24:1)+H | 80 | 10 | 43 | 15 | 39.6 | 1 |
| 11 | 843.7 | 184.1 | 10.47 | SM(26:0)+H | 80 | 10 | 43 | 15 | 31.9 | 1 |
| 12 | 841.7 | 184.1 | 11.61 | SM(26:1)+H | 80 | 10 | 25 | 15 | 174.8 | 1 |
| 13 | 754.7 | 369.4 | 2.31 | CE(24:0)+H | 80 | 10 | 25 | 15 | 32.5 | 3.01 |
| 14 | 714.6 | 369.4 | 2.24 | CE(22:6)+H | 80 | 10 | 25 | 15 | 61.1 | 1 |
| 15 | 698.7 | 369.4 | 2.24 | CE(20:0)+H | 80 | 10 | 25 | 15 | 43.8 | 1 |
| 16 | 696.7 | 369.4 | 2.17 | CE(20:1)+H | 80 | 10 | 25 | 15 | 40.2 | 1 |
| 17 | 716.6 | 369.4 | 2.28 | CE(22:5)+H | 80 | 10 | 25 | 15 | 43.2 | 1 |
| 18 | 614.6 | 369.4 | 2.28 | CE(14:0)+H | 80 | 10 | 25 | 15 | 80.5 | 1.03 |
| 19 | 642.6 | 369.4 | 2.36 | CE(16:0)+H | 80 | 10 | 25 | 15 | 96.5 | 1.24 |
| 20 | 640.6 | 369.4 | 2.38 | CE(16:1)+H | 80 | 10 | 25 | 15 | 191.4 | 1.1 |
| 21 | 670.6 | 369.4 | 2.24 | CE(18:0)+H | 80 | 10 | 25 | 15 | 209 | 1.12 |
| 22 | 668.6 | 369.4 | 2.25 | CE(18:1)+H | 80 | 10 | 25 | 15 | 63.4 | 1 |
| 23 | 666.6 | 369.4 | 2.24 | CE(18:2)+H | 80 | 10 | 25 | 15 | 55.2 | 1.01 |
| 24 | 664.6 | 369.4 | 2.24 | CE(18:3)+H | 80 | 10 | 25 | 15 | 58.5 | 1 |
| 25 | 694.6 | 369.4 | 2.27 | CE(20:2)+H | 80 | 10 | 25 | 15 | 35.4 | 1 |
| 26 | 692.6 | 369.4 | 2.29 | CE(20:3)+H | 80 | 10 | 25 | 15 | 64.8 | 1.01 |
| 27 | 690.6 | 369.4 | 2.25 | CE(20:4)+H | 80 | 10 | 25 | 15 | 110.3 | 1.07 |
| 28 | 688.6 | 369.4 | 2.25 | CE(20:5)+H | 80 | 10 | 25 | 15 | 56.7 | 1.02 |
| 29 | 726.7 | 369.4 | 2.28 | CE(22:0)+H | 80 | 10 | 25 | 15 | 45.7 | 1 |
| 30 | 724.7 | 369.4 | 2.25 | CE(22:1)+H | 80 | 10 | 25 | 15 | 90.8 | 1.05 |
| 31 | 722.7 | 369.4 | 1.9 | CE(22:2)+H | 80 | 10 | 25 | 15 | 33.5 | 1 |
| 32 | 718.6 | 369.4 | 2.3 | CE(22:4)+H | 80 | 10 | 25 | 15 | 47.2 | 1 |
| 33 | 752.7 | 369.4 | 2.32 | CE(24:1)+H | 80 | 10 | 43 | 15 | 96.2 | 1 |
| 34 | 510.6 | 264.4 | 2.54 | CER(14:0)+H | 80 | 10 | 43 | 15 | 31.1 | 1 |
| 35 | 538.6 | 264.4 | 2.54 | CER(16:0)+H | 80 | 10 | 43 | 15 | 140 | 1.26 |
| 36 | 566.7 | 264.4 | 2.49 | CER(18:0)+H | 80 | 10 | 43 | 15 | 31.8 | 1 |
| 37 | 564.8 | 264.4 | 2.58 | CER(18:1)+H | 80 | 10 | 43 | 15 | 56 | 1 |
| 38 | 594.6 | 264.4 | 2.48 | CER(20:0)+H | 80 | 10 | 43 | 15 | 154.6 | 1.04 |
| 39 | 592.6 | 264.4 | 2.31 | CER(20:1)+H | 80 | 10 | 43 | 15 | 45.3 | 1.01 |
| 40 | 622.7 | 264.4 | 2.48 | CER(22:0)+H | 80 | 10 | 43 | 15 | 67.5 | 1.03 |
| 41 | 620.7 | 264.4 | 2.48 | CER(22:1)+H | 80 | 10 | 43 | 15 | 61.2 | 1 |
| 42 | 650.8 | 264.4 | 2.49 | CER(24:0)+H | 80 | 10 | 43 | 15 | 56 | 1 |
| 43 | 648.8 | 264.4 | 2.5 | CER(24:1)+H | 80 | 10 | 43 | 15 | 41.7 | 1 |

| | | | | | | | | | | |
|---|---|---|---|---|---|---|---|---|---|---|
| 44 | 678.9 | 264.4 | 2.55 | CER(26:0)+H | 80 | 10 | 43 | 15 | 50.8 | 1 |
| 45 | 676.9 | 264.4 | 0.69 | CER(26:1)+H | 80 | 10 | 43 | 15 | 177.9 | 1.02 |
| 46 | 512.6 | 266.4 | 2.23 | DCER(14:0)+H | 80 | 10 | 43 | 15 | 63.4 | 1 |
| 47 | 540.6 | 266.4 | 2.48 | DCER(16:0)+H | 80 | 10 | 43 | 15 | 197.2 | 1.25 |
| 48 | 568.7 | 266.4 | 2.49 | DCER(18:0)+H | 80 | 10 | 43 | 15 | 55.9 | 1 |
| 49 | 566.8 | 266.4 | 2.54 | DCER(18:1)+H | 80 | 10 | 43 | 15 | 141.3 | 1.04 |
| 50 | 596.7 | 266.4 | 2.48 | DCER(20:0)+H | 80 | 10 | 43 | 15 | 92.8 | 1.29 |
| 51 | 594.4 | 266.4 | 2.34 | DCER(20:1)+H | 80 | 10 | 43 | 15 | 77.9 | 1.05 |
| 52 | 624.8 | 266.4 | 2.49 | DCER(22:0)+H | 80 | 10 | 43 | 15 | 141.3 | 1.35 |
| 53 | 620.4 | 266.4 | 2.48 | DCER(22:1)+H | 80 | 10 | 43 | 15 | 42.2 | 1 |
| 54 | 652.9 | 266.4 | 2.49 | DCER(24:0)+H | 80 | 10 | 43 | 15 | 105.5 | 1 |
| 55 | 650.9 | 266.4 | 2.48 | DCER(24:1)+H | 80 | 10 | 43 | 15 | 43 | 1 |
| 56 | 680.5 | 266.4 | 2.54 | DCER(26:0)+H | 80 | 10 | 43 | 15 | 84.1 | 1 |
| 57 | 678.5 | 266.4 | 3.73 | DCER(26:1)+H | 80 | 10 | 43 | 15 | 141.3 | 1.14 |
| 58 | 672.5 | 264.4 | 7.36 | HCER(14:0)+H | 80 | 10 | 43 | 15 | 64 | 1.09 |
| 59 | 700.7 | 264.4 | 6.44 | HCER(16:0)+H | 80 | 10 | 43 | 15 | 47.1 | 1.18 |
| 60 | 728.8 | 264.4 | 6.14 | HCER(18:0)+H | 80 | 10 | 43 | 15 | 41.7 | 1 |
| 61 | 726.7 | 264.4 | 7.74 | HCER(18:1)+H | 80 | 10 | 43 | 15 | 55.5 | 1.01 |
| 62 | 756.7 | 264.4 | 5.74 | HCER(20:0)+H | 80 | 10 | 43 | 15 | 69.2 | 1 |
| 63 | 754.7 | 264.4 | 7.65 | HCER(20:1)+H | 80 | 10 | 43 | 15 | 63.9 | 1 |
| 64 | 784.9 | 264.4 | 5.95 | HCER(22:0)+H | 80 | 10 | 43 | 15 | 101.1 | 1 |
| 65 | 782.8 | 264.4 | 5.08 | HCER(22:1)+H | 80 | 10 | 43 | 15 | 83.9 | 1 |
| 66 | 812.9 | 264.4 | 5.11 | HCER(24:0)+H | 80 | 10 | 43 | 15 | 93.1 | 1 |
| 67 | 810.9 | 264.4 | 5.93 | HCER(24:1)+H | 80 | 10 | 43 | 15 | 138.6 | 1 |
| 68 | 840.9 | 264.4 | 5.44 | HCER(26:0)+H | 80 | 10 | 43 | 15 | 56 | 1 |
| 69 | 838.9 | 264.4 | 6.05 | HCER(26:1)+H | 80 | 10 | 43 | 15 | 643.7 | 1.07 |
| 70 | 730.8 | 266.4 | 9.95 | HCER(d18:0/18:0)+H | 80 | 10 | 43 | 15 | 206.2 | 1.07 |
| 71 | 758.7 | 266.4 | 6.01 | HCER(d18:0/20:0)+H | 80 | 10 | 43 | 15 | 334.1 | 1.12 |
| 72 | 786.9 | 266.4 | 5.98 | HCER(d18:0/22:0)+H | 80 | 10 | 43 | 15 | 643.7 | 1.87 |
| 73 | 814.9 | 266.4 | 5.95 | HCER(d18:0/24:0)+H | 80 | 10 | 43 | 15 | 119.8 | 1.1 |
| 74 | 812.9 | 266.4 | 5.93 | HCER(d18:0/24:1)+H | 80 | 10 | 43 | 15 | 197.6 | 1.06 |
| 75 | 842.9 | 266.4 | 9.28 | HCER(d18:0/26:0)+H | 80 | 10 | 43 | 15 | 79.7 | 1.03 |
| 76 | 840.9 | 266.4 | 9.31 | HCER(d18:0/26:1)+H | 80 | 10 | 43 | 15 | 146.6 | 1.14 |
| 77 | 834.9 | 264.4 | 2.27 | LCER(14:0)+H | 80 | 10 | 43 | 15 | 77.3 | 1.07 |
| 78 | 862.9 | 264.4 | 2.22 | LCER(16:0)+H | 80 | 10 | 43 | 15 | 83.3 | 1.11 |
| 79 | 890.2 | 264.4 | 2.3 | LCER(18:0)+H | 80 | 10 | 43 | 15 | 92.3 | 1.09 |
| 80 | 888.2 | 264.4 | 2.27 | LCER(18:1)+H | 80 | 10 | 43 | 15 | 86.3 | 1.07 |
| 81 | 918.2 | 264.4 | 2.21 | LCER(20:0)+H | 80 | 10 | 43 | 15 | 93.5 | 1.04 |
| 82 | 916.2 | 264.4 | 2.27 | LCER(20:1)+H | 80 | 10 | 43 | 15 | 66.4 | 1.04 |
| 83 | 946.2 | 264.4 | 2.34 | LCER(22:0)+H | 80 | 10 | 43 | 15 | 125.9 | 1.06 |
| 84 | 944.2 | 264.4 | 2.29 | LCER(22:1)+H | 80 | 10 | 43 | 15 | 136.7 | 1.13 |
| 85 | 974.8 | 264.4 | 2.39 | LCER(24:0)+H | 80 | 10 | 43 | 15 | 96.4 | 1.14 |
| 86 | 972.9 | 264.4 | 2.41 | LCER(24:1)+H | 80 | 10 | 43 | 15 | 615.1 | 1.34 |
| 87 | 1002.4 | 264.4 | 2.48 | LCER(26:0)+H | 80 | 10 | 43 | 15 | 128.1 | 1.26 |
| 88 | 1000.4 | 264.4 | 2.32 | LCER(26:1)+H | 80 | 10 | 38 | 15 | 615.1 | 1.7 |
| 89 | 892.7 | 266.4 | 2.23 | LCER(d18:0/18:0)+H | 80 | 10 | 43 | 15 | 125.6 | 2.23 |

| | | | | | | | | | | |
|---|---|---|---|---|---|---|---|---|---|---|
| 90 | 920.7 | 266.4 | 2.28 | LCER(d18:0/20:0)+H | 80 | 10 | 43 | 15 | 157.7 | 1.15 |
| 91 | 948.7 | 266.4 | 2.29 | LCER(d18:0/22:0)+H | 80 | 10 | 43 | 15 | 86.8 | 1.22 |
| 92 | 976.8 | 266.4 | 2.15 | LCER(d18:0/24:0)+H | 80 | 10 | 43 | 15 | 615.1 | 1.68 |
| 93 | 974.7 | 266.4 | 2.74 | LCER(d18:0/24:1)+H | 80 | 10 | 43 | 15 | 615.1 | 2.38 |
| 94 | 1004.9 | 266.4 | 2.24 | LCER(d18:0/26:0)+H | 80 | 10 | 43 | 15 | 615.1 | 1.41 |
| 95 | 1002.9 | 266.4 | 2.76 | LCER(d18:0/26:1)+H | 80 | 10 | 43 | 15 | 615.1 | 2.58 |
| 96 | 712.645 | 467.409 | 2.2 | TAG(40:0/FA14:0)+NH4 | 80 | 10 | 38 | 15 | 95.7 | 1 |
| 97 | 712.645 | 439.378 | 2.2 | TAG(40:0/FA16:0)+NH4 | 80 | 10 | 38 | 15 | 136.1 | 1.04 |
| 98 | 740.676 | 495.441 | 2.2 | TAG(42:0/FA14:0)+NH4 | 80 | 10 | 38 | 15 | 115 | 1.01 |
| 99 | 740.676 | 467.409 | 2.19 | TAG(42:0/FA16:0)+NH4 | 80 | 10 | 38 | 15 | 94.6 | 1.01 |
| 100 | 738.661 | 493.425 | 2.19 | TAG(42:1/FA14:0)+NH4 | 80 | 10 | 38 | 15 | 90.5 | 1 |
| 101 | 738.661 | 465.394 | 2.19 | TAG(42:1/FA16:0)+NH4 | 80 | 10 | 38 | 15 | 93.3 | 1.05 |
| 102 | 738.661 | 467.409 | 2.2 | TAG(42:1/FA16:1)+NH4 | 80 | 10 | 38 | 15 | 130.6 | 1.04 |
| 103 | 738.661 | 439.378 | 2.2 | TAG(42:1/FA18:1)+NH4 | 80 | 10 | 38 | 15 | 137.5 | 1.11 |
| 104 | 736.645 | 439.378 | 2.19 | TAG(42:2/FA18:2)+NH4 | 80 | 10 | 38 | 15 | 89.5 | 1 |
| 105 | 768.708 | 523.472 | 2.2 | TAG(44:0/FA14:0)+NH4 | 80 | 10 | 38 | 15 | 88 | 1 |
| 106 | 768.708 | 495.441 | 2.2 | TAG(44:0/FA16:0)+NH4 | 80 | 10 | 38 | 15 | 90.4 | 1 |
| 107 | 768.708 | 467.409 | 2.21 | TAG(44:0/FA18:0)+NH4 | 80 | 10 | 38 | 15 | 104.5 | 1 |
| 108 | 766.692 | 521.456 | 2.19 | TAG(44:1/FA14:0)+NH4 | 80 | 10 | 38 | 15 | 301.2 | 1 |
| 109 | 766.692 | 493.425 | 2.2 | TAG(44:1/FA16:0)+NH4 | 80 | 10 | 38 | 15 | 101.4 | 1 |
| 110 | 766.692 | 495.441 | 2.19 | TAG(44:1/FA16:1)+NH4 | 80 | 10 | 38 | 15 | 103.3 | 1 |
| 111 | 766.692 | 467.409 | 2.2 | TAG(44:1/FA18:1)+NH4 | 80 | 10 | 38 | 15 | 125.8 | 1.02 |
| 112 | 764.676 | 519.441 | 2.19 | TAG(44:2/FA14:0)+NH4 | 80 | 10 | 38 | 15 | 82 | 1 |
| 113 | 764.676 | 491.409 | 2.19 | TAG(44:2/FA16:0)+NH4 | 80 | 10 | 38 | 15 | 93.6 | 1 |
| 114 | 764.676 | 493.425 | 2.21 | TAG(44:2/FA16:1)+NH4 | 80 | 10 | 38 | 15 | 97.1 | 1 |
| 115 | 764.676 | 465.394 | 2.2 | TAG(44:2/FA18:1)+NH4 | 80 | 10 | 38 | 15 | 97.3 | 1.09 |
| 116 | 764.676 | 467.409 | 0.9 | TAG(44:2/FA18:2)+NH4 | 80 | 10 | 38 | 15 | 68.7 | 1.02 |
| 117 | 762.661 | 465.394 | 2.2 | TAG(44:3/FA18:2)+NH4 | 80 | 10 | 38 | 15 | 100 | 1 |
| 118 | 782.723 | 537.488 | 2.21 | TAG(45:0/FA14:0)+NH4 | 80 | 10 | 38 | 15 | 82.8 | 1.03 |
| 119 | 782.723 | 509.456 | 2.28 | TAG(45:0/FA16:0)+NH4 | 80 | 10 | 38 | 15 | 93.4 | 1.04 |
| 120 | 780.708 | 507.4 | 2.21 | TAG(45:1/FA16:0)+NH4 | 80 | 10 | 38 | 15 | 94.3 | 1 |
| 121 | 780.708 | 481.4 | 2.21 | TAG(45:1/FA18:1)+NH4 | 80 | 10 | 38 | 15 | 90.6 | 1.02 |
| 122 | 796.7 | 551.503 | 2.23 | TAG(46:0/FA14:0)+NH4 | 80 | 10 | 38 | 15 | 67.1 | 1.03 |
| 123 | 796.7 | 523.472 | 2.25 | TAG(46:0/FA16:0)+NH4 | 80 | 10 | 38 | 15 | 100.7 | 1 |
| 124 | 796.7 | 495.441 | 2.24 | TAG(46:0/FA18:0)+NH4 | 80 | 10 | 38 | 15 | 88.9 | 1 |
| 125 | 794.7 | 549.5 | 2.2 | TAG(46:1/FA14:0)+NH4 | 80 | 10 | 38 | 15 | 89.6 | 1 |
| 126 | 794.7 | 521.4 | 2.21 | TAG(46:1/FA16:0)+NH4 | 80 | 10 | 38 | 15 | 90.1 | 1 |
| 127 | 794.7 | 523.472 | 2.21 | TAG(46:1/FA16:1)+NH4 | 80 | 10 | 38 | 15 | 97.4 | 1 |
| 128 | 794.7 | 493.425 | 2.23 | TAG(46:1/FA18:0)+NH4 | 80 | 10 | 38 | 15 | 92.3 | 1 |
| 129 | 794.7 | 495.441 | 2.2 | TAG(46:1/FA18:1)+NH4 | 80 | 10 | 38 | 15 | 90.4 | 1.02 |
| 130 | 792.7 | 547.5 | 2.2 | TAG(46:2/FA14:0)+NH4 | 80 | 10 | 38 | 15 | 85.5 | 1 |
| 131 | 792.7 | 519.441 | 2.19 | TAG(46:2/FA16:0)+NH4 | 80 | 10 | 38 | 15 | 77.2 | 1 |
| 132 | 792.7 | 521.4 | 2.2 | TAG(46:2/FA16:1)+NH4 | 80 | 10 | 38 | 15 | 95.5 | 1 |
| 133 | 792.7 | 493.425 | 2.19 | TAG(46:2/FA18:1)+NH4 | 80 | 10 | 38 | 15 | 96 | 1.01 |
| 134 | 792.7 | 495.441 | 2.19 | TAG(46:2/FA18:2)+NH4 | 80 | 10 | 38 | 15 | 78.2 | 1 |
| 135 | 790.7 | 545.5 | 2.2 | TAG(46:3/FA14:0)+NH4 | 80 | 10 | 38 | 15 | 82 | 1 |

| | | | | | | | | | | |
|---|---|---|---|---|---|---|---|---|---|---|
| 136 | 790.7 | 517.4 | 2.19 | TAG(46:3/FA16:0)+NH4 | 80 | 10 | 38 | 15 | 110.9 | 1.03 |
| 137 | 790.7 | 519.441 | 2.19 | TAG(46:3/FA16:1)+NH4 | 80 | 10 | 38 | 15 | 76.2 | 1 |
| 138 | 790.7 | 491.409 | 2.2 | TAG(46:3/FA18:1)+NH4 | 80 | 10 | 38 | 15 | 62.6 | 1.02 |
| 139 | 790.7 | 493.425 | 2.2 | TAG(46:3/FA18:2)+NH4 | 80 | 10 | 38 | 15 | 65.1 | 1 |
| 140 | 790.7 | 495.441 | 2.19 | TAG(46:3/FA18:3)+NH4 | 80 | 10 | 38 | 15 | 78.6 | 1 |
| 141 | 788.7 | 491.409 | 2.2 | TAG(46:4/FA18:2)+NH4 | 80 | 10 | 38 | 15 | 91.4 | 1 |
| 142 | 810.7 | 565.5 | 2.25 | TAG(47:0/FA14:0)+NH4 | 80 | 10 | 38 | 15 | 65 | 1 |
| 143 | 810.7 | 537.4 | 2.27 | TAG(47:0/FA16:0)+NH4 | 80 | 10 | 38 | 15 | 99 | 1.01 |
| 144 | 810.7 | 523.472 | 2.26 | TAG(47:0/FA17:0)+NH4 | 80 | 10 | 38 | 15 | 75.3 | 1 |
| 145 | 808.7 | 563.5 | 2.2 | TAG(47:1/FA14:0)+NH4 | 80 | 10 | 38 | 15 | 129 | 1.01 |
| 146 | 808.7 | 535.4 | 2.2 | TAG(47:1/FA16:0)+NH4 | 80 | 10 | 38 | 15 | 104.6 | 1.01 |
| 147 | 808.7 | 537.4 | 2.21 | TAG(47:1/FA16:1)+NH4 | 80 | 10 | 38 | 15 | 82.4 | 1 |
| 148 | 808.7 | 521.4 | 2.22 | TAG(47:1/FA17:0)+NH4 | 80 | 10 | 38 | 15 | 136.1 | 1.01 |
| 149 | 808.7 | 509.4 | 2.29 | TAG(47:1/FA18:1)+NH4 | 80 | 10 | 38 | 15 | 136.1 | 1.05 |
| 150 | 806.7 | 561.5 | 2.19 | TAG(47:2/FA14:0)+NH4 | 80 | 10 | 38 | 15 | 74.6 | 1 |
| 151 | 806.7 | 535.4 | 2.23 | TAG(47:2/FA16:1)+NH4 | 80 | 10 | 38 | 15 | 106.9 | 1.02 |
| 152 | 806.7 | 507.4 | 2.22 | TAG(47:2/FA18:1)+NH4 | 80 | 10 | 38 | 15 | 136.1 | 1.01 |
| 153 | 806.7 | 509.4 | 2.22 | TAG(47:2/FA18:2)+NH4 | 80 | 10 | 38 | 15 | 80.3 | 1.01 |
| 154 | 824.7 | 579.5 | 2.29 | TAG(48:0/FA14:0)+NH4 | 80 | 10 | 38 | 15 | 99.4 | 1 |
| 155 | 824.7 | 551.4 | 2.3 | TAG(48:0/FA16:0)+NH4 | 80 | 10 | 38 | 15 | 80.6 | 1 |
| 156 | 824.7 | 523.472 | 2.28 | TAG(48:0/FA18:0)+NH4 | 80 | 10 | 38 | 15 | 93.5 | 1 |
| 157 | 822.7 | 577.5 | 2.23 | TAG(48:1/FA14:0)+NH4 | 80 | 10 | 38 | 15 | 89 | 1 |
| 158 | 822.7 | 549.4 | 2.19 | TAG(48:1/FA16:0)+NH4 | 80 | 10 | 38 | 15 | 80.5 | 1 |
| 159 | 822.7 | 551.4 | 2.25 | TAG(48:1/FA16:1)+NH4 | 80 | 10 | 38 | 15 | 79.5 | 1 |
| 160 | 822.7 | 521.4 | 2.2 | TAG(48:1/FA18:0)+NH4 | 80 | 10 | 38 | 15 | 100.2 | 1 |
| 161 | 822.7 | 523.472 | 2.22 | TAG(48:1/FA18:1)+NH4 | 80 | 10 | 38 | 15 | 90.2 | 1 |
| 162 | 820.7 | 575.5 | 2.19 | TAG(48:2/FA14:0)+NH4 | 80 | 10 | 38 | 15 | 81.9 | 1 |
| 163 | 820.7 | 547.4 | 2.2 | TAG(48:2/FA16:0)+NH4 | 80 | 10 | 38 | 15 | 84.1 | 1 |
| 164 | 820.7 | 549.4 | 2.19 | TAG(48:2/FA16:1)+NH4 | 80 | 10 | 38 | 15 | 78.9 | 1 |
| 165 | 820.7 | 519.441 | 2.2 | TAG(48:2/FA18:0)+NH4 | 80 | 10 | 38 | 15 | 82.5 | 1 |
| 166 | 820.7 | 521.4 | 2.19 | TAG(48:2/FA18:1)+NH4 | 80 | 10 | 38 | 15 | 91 | 1 |
| 167 | 820.7 | 523.472 | 2.2 | TAG(48:2/FA18:2)+NH4 | 80 | 10 | 38 | 15 | 92.4 | 1 |
| 168 | 818.7 | 573.5 | 2.19 | TAG(48:3/FA14:0)+NH4 | 80 | 10 | 38 | 15 | 89.1 | 1 |
| 169 | 818.7 | 545.4 | 2.19 | TAG(48:3/FA16:0)+NH4 | 80 | 10 | 38 | 15 | 87.5 | 1 |
| 170 | 818.7 | 547.4 | 2.19 | TAG(48:3/FA16:1)+NH4 | 80 | 10 | 38 | 15 | 80.5 | 1 |
| 171 | 818.7 | 519.441 | 2.2 | TAG(48:3/FA18:1)+NH4 | 80 | 10 | 38 | 15 | 89.4 | 1 |
| 172 | 818.7 | 521.4 | 2.19 | TAG(48:3/FA18:2)+NH4 | 80 | 10 | 38 | 15 | 86.9 | 1 |
| 173 | 818.7 | 523.472 | 2.19 | TAG(48:3/FA18:3)+NH4 | 80 | 10 | 38 | 15 | 87.9 | 1 |
| 174 | 816.7 | 571.5 | 2.19 | TAG(48:4/FA14:0)+NH4 | 80 | 10 | 38 | 15 | 88.5 | 1 |
| 175 | 816.7 | 543.4 | 2.21 | TAG(48:4/FA16:0)+NH4 | 80 | 10 | 38 | 15 | 121.2 | 1.05 |
| 176 | 816.7 | 545.4 | 2.2 | TAG(48:4/FA16:1)+NH4 | 80 | 10 | 38 | 15 | 92.4 | 1.04 |
| 177 | 816.7 | 517.4 | 2.19 | TAG(48:4/FA18:1)+NH4 | 80 | 10 | 38 | 15 | 97.6 | 1.04 |
| 178 | 816.7 | 519.441 | 2.19 | TAG(48:4/FA18:2)+NH4 | 80 | 10 | 38 | 15 | 95.2 | 1 |
| 179 | 816.7 | 521.4 | 2.19 | TAG(48:4/FA18:3)+NH4 | 80 | 10 | 38 | 15 | 88.8 | 1 |
| 180 | 816.7 | 495.441 | 2.21 | TAG(48:4/FA20:4)+NH4 | 80 | 10 | 38 | 15 | 69.7 | 1 |
| 181 | 814.7 | 517.4 | 2.2 | TAG(48:5/FA18:2)+NH4 | 80 | 10 | 38 | 15 | 132.4 | 1.05 |

| | | | | | | | | | | |
|---|---|---|---|---|---|---|---|---|---|---|
| 182 | 814.7 | 519.441 | 2.19 | TAG(48:5/FA18:3)+NH4 | 80 | 10 | 38 | 15 | 80.8 | 1 |
| 183 | 838.8 | 565.5 | 2.28 | TAG(49:0/FA16:0)+NH4 | 80 | 10 | 38 | 15 | 77.8 | 1 |
| 184 | 838.8 | 551.503 | 2.28 | TAG(49:0/FA17:0)+NH4 | 80 | 10 | 38 | 15 | 70.1 | 1 |
| 185 | 838.8 | 537.5 | 2.29 | TAG(49:0/FA18:0)+NH4 | 80 | 10 | 38 | 15 | 83.5 | 1 |
| 186 | 836.8 | 591.6 | 2.2 | TAG(49:1/FA14:0)+NH4 | 80 | 10 | 38 | 15 | 82.9 | 1 |
| 187 | 836.8 | 563.5 | 2.25 | TAG(49:1/FA16:0)+NH4 | 80 | 10 | 38 | 15 | 70 | 1 |
| 188 | 836.8 | 565.5 | 2.25 | TAG(49:1/FA16:1)+NH4 | 80 | 10 | 38 | 15 | 76.6 | 1 |
| 189 | 836.8 | 549.5 | 2.22 | TAG(49:1/FA17:0)+NH4 | 80 | 10 | 38 | 15 | 136.1 | 1 |
| 190 | 836.8 | 537.5 | 2.25 | TAG(49:1/FA18:1)+NH4 | 80 | 10 | 38 | 15 | 85.3 | 1 |
| 191 | 834.8 | 589.6 | 2.19 | TAG(49:2/FA14:0)+NH4 | 80 | 10 | 38 | 15 | 88.9 | 1 |
| 192 | 834.8 | 561.5 | 2.2 | TAG(49:2/FA16:0)+NH4 | 80 | 10 | 38 | 15 | 115.4 | 1 |
| 193 | 834.8 | 563.5 | 2.21 | TAG(49:2/FA16:1)+NH4 | 80 | 10 | 38 | 15 | 100.4 | 1 |
| 194 | 834.8 | 547.5 | 2.2 | TAG(49:2/FA17:0)+NH4 | 80 | 10 | 38 | 15 | 77.9 | 1 |
| 195 | 834.8 | 535.5 | 2.2 | TAG(49:2/FA18:1)+NH4 | 80 | 10 | 38 | 15 | 121.4 | 1.01 |
| 196 | 834.8 | 537.5 | 2.2 | TAG(49:2/FA18:2)+NH4 | 80 | 10 | 38 | 15 | 78 | 1 |
| 197 | 832.8 | 559.5 | 2.2 | TAG(49:3/FA16:0)+NH4 | 80 | 10 | 38 | 15 | 98.3 | 1 |
| 198 | 832.8 | 561.5 | 2.21 | TAG(49:3/FA16:1)+NH4 | 80 | 10 | 38 | 15 | 109.6 | 1 |
| 199 | 832.8 | 535.5 | 2.2 | TAG(49:3/FA18:2)+NH4 | 80 | 10 | 38 | 15 | 91 | 1 |
| 200 | 832.8 | 537.5 | 2.19 | TAG(49:3/FA18:3)+NH4 | 80 | 10 | 38 | 15 | 78.5 | 1 |
| 201 | 852.8 | 607.6 | 2.31 | TAG(50:0/FA14:0)+NH4 | 80 | 10 | 38 | 15 | 118.9 | 1.01 |
| 202 | 852.8 | 579.5 | 2.33 | TAG(50:0/FA16:0)+NH4 | 80 | 10 | 38 | 15 | 79.6 | 1 |
| 203 | 852.8 | 551.503 | 2.34 | TAG(50:0/FA18:0)+NH4 | 80 | 10 | 38 | 15 | 79.4 | 1 |
| 204 | 850.8 | 605.6 | 2.26 | TAG(50:1/FA14:0)+NH4 | 80 | 10 | 38 | 15 | 77.7 | 1 |
| 205 | 850.8 | 577.5 | 2.27 | TAG(50:1/FA16:0)+NH4 | 80 | 10 | 38 | 15 | 92 | 1 |
| 206 | 850.8 | 579.5 | 2.26 | TAG(50:1/FA16:1)+NH4 | 80 | 10 | 38 | 15 | 77.2 | 1 |
| 207 | 850.8 | 549.5 | 2.27 | TAG(50:1/FA18:0)+NH4 | 80 | 10 | 38 | 15 | 84.5 | 1 |
| 208 | 850.8 | 551.503 | 2.28 | TAG(50:1/FA18:1)+NH4 | 80 | 10 | 38 | 15 | 83.1 | 1 |
| 209 | 850.8 | 523.472 | 2.21 | TAG(50:1/FA20:1)+NH4 | 80 | 10 | 38 | 15 | 81 | 1 |
| 210 | 848.8 | 603.6 | 2.22 | TAG(50:2/FA14:0)+NH4 | 80 | 10 | 38 | 15 | 68.4 | 1 |
| 211 | 848.8 | 575.5 | 2.22 | TAG(50:2/FA16:0)+NH4 | 80 | 10 | 38 | 15 | 90 | 1 |
| 212 | 848.8 | 577.5 | 2.21 | TAG(50:2/FA16:1)+NH4 | 80 | 10 | 38 | 15 | 93.1 | 1 |
| 213 | 848.8 | 547.5 | 2.22 | TAG(50:2/FA18:0)+NH4 | 80 | 10 | 38 | 15 | 95.4 | 1 |
| 214 | 848.8 | 549.5 | 2.22 | TAG(50:2/FA18:1)+NH4 | 80 | 10 | 38 | 15 | 97.5 | 1 |
| 215 | 848.8 | 551.503 | 2.23 | TAG(50:2/FA18:2)+NH4 | 80 | 10 | 38 | 15 | 85.3 | 1 |
| 216 | 848.8 | 523.472 | 2.22 | TAG(50:2/FA20:2)+NH4 | 80 | 10 | 38 | 15 | 92.5 | 1 |
| 217 | 846.8 | 601.6 | 2.19 | TAG(50:3/FA14:0)+NH4 | 80 | 10 | 38 | 15 | 86.7 | 1.02 |
| 218 | 846.8 | 573.5 | 2.2 | TAG(50:3/FA16:0)+NH4 | 80 | 10 | 38 | 15 | 93.8 | 1 |
| 219 | 846.8 | 575.5 | 2.2 | TAG(50:3/FA16:1)+NH4 | 80 | 10 | 38 | 15 | 90.8 | 1 |
| 220 | 846.8 | 545.5 | 2.21 | TAG(50:3/FA18:0)+NH4 | 80 | 10 | 38 | 15 | 94 | 1 |
| 221 | 846.8 | 547.5 | 2.19 | TAG(50:3/FA18:1)+NH4 | 80 | 10 | 38 | 15 | 108.9 | 1 |
| 222 | 846.8 | 549.5 | 2.2 | TAG(50:3/FA18:2)+NH4 | 80 | 10 | 38 | 15 | 100 | 1 |
| 223 | 846.8 | 551.503 | 2.2 | TAG(50:3/FA18:3)+NH4 | 80 | 10 | 38 | 15 | 90.8 | 1 |
| 224 | 846.8 | 523.472 | 2.19 | TAG(50:3/FA20:3)+NH4 | 80 | 10 | 38 | 15 | 96.1 | 1 |
| 225 | 844.6 | 599.4 | 2.2 | TAG(50:4/FA14:0)+NH4 | 80 | 10 | 38 | 15 | 112.1 | 1.01 |
| 226 | 844.6 | 571.3 | 2.19 | TAG(50:4/FA16:0)+NH4 | 80 | 10 | 38 | 15 | 100.3 | 1 |
| 227 | 844.6 | 573.3 | 2.19 | TAG(50:4/FA16:1)+NH4 | 80 | 10 | 38 | 15 | 101.8 | 1 |

| | | | | | | | | | | |
|---|---|---|---|---|---|---|---|---|---|---|
| 228 | 844.6 | 545.3 | 2.19 | TAG(50:4/FA18:1)+NH4 | 80 | 10 | 38 | 15 | 102.6 | 1 |
| 229 | 844.6 | 547.3 | 2.19 | TAG(50:4/FA18:2)+NH4 | 80 | 10 | 38 | 15 | 116.1 | 1 |
| 230 | 844.6 | 549.3 | 2.2 | TAG(50:4/FA18:3)+NH4 | 80 | 10 | 38 | 15 | 98.2 | 1 |
| 231 | 844.6 | 521.3 | 2.22 | TAG(50:4/FA20:3)+NH4 | 80 | 10 | 38 | 15 | 96.5 | 1 |
| 232 | 844.6 | 523.3 | 2.21 | TAG(50:4/FA20:4)+NH4 | 80 | 10 | 38 | 15 | 115.3 | 1.07 |
| 233 | 842.6 | 597.4 | 2.19 | TAG(50:5/FA14:0)+NH4 | 80 | 10 | 38 | 15 | 121.3 | 1.01 |
| 234 | 842.6 | 569.3 | 2.21 | TAG(50:5/FA16:0)+NH4 | 80 | 10 | 38 | 15 | 93.3 | 1 |
| 235 | 842.6 | 571.3 | 0.91 | TAG(50:5/FA16:1)+NH4 | 80 | 10 | 38 | 15 | 109 | 1.05 |
| 236 | 842.6 | 543.3 | 2.2 | TAG(50:5/FA18:1)+NH4 | 80 | 10 | 38 | 15 | 105.9 | 1.04 |
| 237 | 842.6 | 545.3 | 2.19 | TAG(50:5/FA18:2)+NH4 | 80 | 10 | 38 | 15 | 94.7 | 1.04 |
| 238 | 842.6 | 547.3 | 2.19 | TAG(50:5/FA18:3)+NH4 | 80 | 10 | 38 | 15 | 89.9 | 1 |
| 239 | 842.6 | 521.3 | 2.22 | TAG(50:5/FA20:4)+NH4 | 80 | 10 | 38 | 15 | 91.2 | 1 |
| 240 | 842.6 | 523.3 | 2.23 | TAG(50:5/FA20:5)+NH4 | 80 | 10 | 38 | 15 | 104.7 | 1.03 |
| 241 | 840.7 | 519.441 | 2.19 | TAG(50:6/FA20:4)+NH4 | 80 | 10 | 38 | 15 | 129.5 | 1.08 |
| 242 | 866.8 | 593.5 | 2.32 | TAG(51:0/FA16:0)+NH4 | 80 | 10 | 38 | 15 | 89.8 | 1.02 |
| 243 | 866.8 | 579.5 | 2.32 | TAG(51:0/FA17:0)+NH4 | 80 | 10 | 38 | 15 | 104.2 | 1 |
| 244 | 866.8 | 565.5 | 2.32 | TAG(51:0/FA18:0)+NH4 | 80 | 10 | 38 | 15 | 73.1 | 1 |
| 245 | 864.8 | 591.5 | 2.25 | TAG(51:1/FA16:0)+NH4 | 80 | 10 | 38 | 15 | 82.4 | 1 |
| 246 | 864.8 | 577.5 | 2.26 | TAG(51:1/FA17:0)+NH4 | 80 | 10 | 38 | 15 | 87.2 | 1 |
| 247 | 864.8 | 563.5 | 2.28 | TAG(51:1/FA18:0)+NH4 | 80 | 10 | 38 | 15 | 89.3 | 1 |
| 248 | 864.8 | 565.5 | 2.27 | TAG(51:1/FA18:1)+NH4 | 80 | 10 | 38 | 15 | 86.7 | 1 |
| 249 | 862.8 | 589.5 | 2.21 | TAG(51:2/FA16:0)+NH4 | 80 | 10 | 38 | 15 | 79.4 | 1 |
| 250 | 862.8 | 591.5 | 2.23 | TAG(51:2/FA16:1)+NH4 | 80 | 10 | 38 | 15 | 87.8 | 1 |
| 251 | 862.8 | 575.5 | 2.2 | TAG(51:2/FA17:0)+NH4 | 80 | 10 | 38 | 15 | 95.5 | 1 |
| 252 | 862.8 | 563.5 | 2.23 | TAG(51:2/FA18:1)+NH4 | 80 | 10 | 38 | 15 | 100.8 | 1 |
| 253 | 862.8 | 565.5 | 2.21 | TAG(51:2/FA18:2)+NH4 | 80 | 10 | 38 | 15 | 89.6 | 1 |
| 254 | 860.8 | 589.5 | 2.19 | TAG(51:3/FA16:1)+NH4 | 80 | 10 | 38 | 15 | 86.8 | 1 |
| 255 | 860.8 | 573.5 | 2.19 | TAG(51:3/FA17:0)+NH4 | 80 | 10 | 38 | 15 | 92.8 | 1 |
| 256 | 860.8 | 563.5 | 2.19 | TAG(51:3/FA18:2)+NH4 | 80 | 10 | 38 | 15 | 92.9 | 1 |
| 257 | 860.8 | 565.5 | 2.21 | TAG(51:3/FA18:3)+NH4 | 80 | 10 | 38 | 15 | 89.1 | 1 |
| 258 | 858.8 | 587.5 | 2.25 | TAG(51:4/FA16:1)+NH4 | 80 | 10 | 38 | 15 | 98.7 | 1 |
| 259 | 858.8 | 561.5 | 2.19 | TAG(51:4/FA18:2)+NH4 | 80 | 10 | 38 | 15 | 86.7 | 1.01 |
| 260 | 858.8 | 563.5 | 2.21 | TAG(51:4/FA18:3)+NH4 | 80 | 10 | 38 | 15 | 82.4 | 1 |
| 261 | 858.8 | 537.5 | 2.21 | TAG(51:4/FA20:4)+NH4 | 80 | 10 | 38 | 15 | 85.3 | 1 |
| 262 | 856.8 | 559.5 | 2.19 | TAG(51:5/FA18:2)+NH4 | 80 | 10 | 38 | 15 | 136.8 | 1.03 |
| 263 | 856.8 | 561.5 | 2.2 | TAG(51:5/FA18:3)+NH4 | 80 | 10 | 38 | 15 | 81 | 1 |
| 264 | 880.8 | 607.5 | 2.34 | TAG(52:0/FA16:0)+NH4 | 80 | 10 | 38 | 15 | 83.1 | 1 |
| 265 | 880.8 | 579.5 | 2.34 | TAG(52:0/FA18:0)+NH4 | 80 | 10 | 38 | 15 | 67 | 1 |
| 266 | 880.8 | 551.503 | 2.21 | TAG(52:0/FA20:0)+NH4 | 80 | 10 | 38 | 15 | 68.9 | 1 |
| 267 | 878.8 | 605.5 | 2.3 | TAG(52:1/FA16:0)+NH4 | 80 | 10 | 38 | 15 | 77.8 | 1 |
| 268 | 878.8 | 607.5 | 2.29 | TAG(52:1/FA16:1)+NH4 | 80 | 10 | 38 | 15 | 81.2 | 1 |
| 269 | 878.8 | 577.5 | 2.3 | TAG(52:1/FA18:0)+NH4 | 80 | 10 | 38 | 15 | 95.2 | 1 |
| 270 | 878.8 | 579.5 | 2.29 | TAG(52:1/FA18:1)+NH4 | 80 | 10 | 38 | 15 | 76.1 | 1 |
| 271 | 878.8 | 549.5 | 2.2 | TAG(52:1/FA20:0)+NH4 | 80 | 10 | 38 | 15 | 81.5 | 1 |
| 272 | 878.8 | 551.503 | 2.23 | TAG(52:1/FA20:1)+NH4 | 80 | 10 | 38 | 15 | 46.3 | 1 |
| 273 | 876.8 | 631.6 | 2.2 | TAG(52:2/FA14:0)+NH4 | 80 | 10 | 38 | 15 | 87.5 | 1 |

| | | | | | | | | | | |
|---|---|---|---|---|---|---|---|---|---|---|
| 274 | 876.8 | 603.5 | 2.28 | TAG(52:2/FA16:0)+NH4 | 80 | 10 | 38 | 15 | 73.9 | 1 |
| 275 | 876.8 | 605.5 | 2.24 | TAG(52:2/FA16:1)+NH4 | 80 | 10 | 38 | 15 | 85.2 | 1 |
| 276 | 876.8 | 575.5 | 2.27 | TAG(52:2/FA18:0)+NH4 | 80 | 10 | 38 | 15 | 86 | 1 |
| 277 | 876.8 | 577.5 | 2.27 | TAG(52:2/FA18:1)+NH4 | 80 | 10 | 38 | 15 | 92.1 | 1 |
| 278 | 876.8 | 579.5 | 2.25 | TAG(52:2/FA18:2)+NH4 | 80 | 10 | 38 | 15 | 98 | 1 |
| 279 | 876.8 | 547.5 | 2.2 | TAG(52:2/FA20:0)+NH4 | 80 | 10 | 38 | 15 | 98.6 | 1 |
| 280 | 876.8 | 549.5 | 2.2 | TAG(52:2/FA20:1)+NH4 | 80 | 10 | 38 | 15 | 71.4 | 1 |
| 281 | 876.8 | 551.503 | 2.25 | TAG(52:2/FA20:2)+NH4 | 80 | 10 | 38 | 15 | 67.1 | 1 |
| 282 | 874.8 | 629.6 | 2.19 | TAG(52:3/FA14:0)+NH4 | 80 | 10 | 38 | 15 | 90.9 | 1 |
| 283 | 874.8 | 601.5 | 2.24 | TAG(52:3/FA16:0)+NH4 | 80 | 10 | 38 | 15 | 77 | 1 |
| 284 | 874.8 | 603.5 | 2.21 | TAG(52:3/FA16:1)+NH4 | 80 | 10 | 38 | 15 | 87.6 | 1 |
| 285 | 874.8 | 573.5 | 2.24 | TAG(52:3/FA18:0)+NH4 | 80 | 10 | 38 | 15 | 86.9 | 1 |
| 286 | 874.8 | 575.5 | 2.22 | TAG(52:3/FA18:1)+NH4 | 80 | 10 | 38 | 15 | 92.3 | 1 |
| 287 | 874.8 | 577.5 | 2.24 | TAG(52:3/FA18:2)+NH4 | 80 | 10 | 38 | 15 | 84.4 | 1 |
| 288 | 874.8 | 579.5 | 2.21 | TAG(52:3/FA18:3)+NH4 | 80 | 10 | 38 | 15 | 89.8 | 1 |
| 289 | 874.8 | 545.5 | 2.21 | TAG(52:3/FA20:0)+NH4 | 80 | 10 | 38 | 15 | 79.2 | 1 |
| 290 | 874.8 | 547.5 | 2.2 | TAG(52:3/FA20:1)+NH4 | 80 | 10 | 38 | 15 | 81.9 | 1 |
| 291 | 874.8 | 549.5 | 2.22 | TAG(52:3/FA20:2)+NH4 | 80 | 10 | 38 | 15 | 83.3 | 1 |
| 292 | 874.8 | 551.503 | 2.23 | TAG(52:3/FA20:3)+NH4 | 80 | 10 | 38 | 15 | 85.7 | 1 |
| 293 | 872.8 | 627.6 | 2.22 | TAG(52:4/FA14:0)+NH4 | 80 | 10 | 38 | 15 | 94.9 | 1 |
| 294 | 872.8 | 599.5 | 2.21 | TAG(52:4/FA16:0)+NH4 | 80 | 10 | 38 | 15 | 85.1 | 1 |
| 295 | 872.8 | 601.5 | 2.2 | TAG(52:4/FA16:1)+NH4 | 80 | 10 | 38 | 15 | 52.4 | 1 |
| 296 | 872.8 | 571.5 | 2.19 | TAG(52:4/FA18:0)+NH4 | 80 | 10 | 38 | 15 | 55.1 | 1 |
| 297 | 872.8 | 573.5 | 2.2 | TAG(52:4/FA18:1)+NH4 | 80 | 10 | 38 | 15 | 94.6 | 1 |
| 298 | 872.8 | 575.5 | 2.21 | TAG(52:4/FA18:2)+NH4 | 80 | 10 | 38 | 15 | 60.5 | 1 |
| 299 | 872.8 | 577.5 | 2.19 | TAG(52:4/FA18:3)+NH4 | 80 | 10 | 38 | 15 | 50.7 | 1 |
| 300 | 872.8 | 543.5 | 2.21 | TAG(52:4/FA20:0)+NH4 | 80 | 10 | 38 | 15 | 68.9 | 1 |
| 301 | 872.8 | 547.5 | 2.22 | TAG(52:4/FA20:2)+NH4 | 80 | 10 | 38 | 15 | 49.3 | 1 |
| 302 | 872.8 | 549.5 | 2.22 | TAG(52:4/FA20:3)+NH4 | 80 | 10 | 38 | 15 | 89.4 | 1 |
| 303 | 872.8 | 551.503 | 2.21 | TAG(52:4/FA20:4)+NH4 | 80 | 10 | 38 | 15 | 136.1 | 1 |
| 304 | 872.8 | 523.472 | 2.21 | TAG(52:4/FA22:4)+NH4 | 80 | 10 | 38 | 15 | 75.4 | 1 |
| 305 | 870.8 | 625.6 | 2.19 | TAG(52:5/FA14:0)+NH4 | 80 | 10 | 38 | 15 | 90.9 | 1.01 |
| 306 | 870.8 | 597.5 | 2.2 | TAG(52:5/FA16:0)+NH4 | 80 | 10 | 38 | 15 | 95.6 | 1 |
| 307 | 870.8 | 599.5 | 2.19 | TAG(52:5/FA16:1)+NH4 | 80 | 10 | 38 | 15 | 55.4 | 1 |
| 308 | 870.8 | 571.5 | 2.2 | TAG(52:5/FA18:1)+NH4 | 80 | 10 | 38 | 15 | 56.8 | 1 |
| 309 | 870.8 | 573.5 | 2.19 | TAG(52:5/FA18:2)+NH4 | 80 | 10 | 38 | 15 | 55.1 | 1 |
| 310 | 870.8 | 575.5 | 2.2 | TAG(52:5/FA18:3)+NH4 | 80 | 10 | 38 | 15 | 53.7 | 1 |
| 311 | 870.8 | 547.5 | 2.19 | TAG(52:5/FA20:3)+NH4 | 80 | 10 | 38 | 15 | 57.4 | 1 |
| 312 | 870.8 | 549.5 | 2.2 | TAG(52:5/FA20:4)+NH4 | 80 | 10 | 38 | 15 | 67.6 | 1 |
| 313 | 870.8 | 551.503 | 2.2 | TAG(52:5/FA20:5)+NH4 | 80 | 10 | 38 | 15 | 79 | 1 |
| 314 | 870.8 | 523.472 | 2.19 | TAG(52:5/FA22:5)+NH4 | 80 | 10 | 38 | 15 | 63.8 | 1 |
| 315 | 868.8 | 623.6 | 2.2 | TAG(52:6/FA14:0)+NH4 | 80 | 10 | 38 | 15 | 74.6 | 1 |
| 316 | 868.8 | 595.5 | 2.22 | TAG(52:6/FA16:0)+NH4 | 80 | 10 | 38 | 15 | 71.1 | 1.02 |
| 317 | 868.8 | 597.5 | 2.19 | TAG(52:6/FA16:1)+NH4 | 80 | 10 | 38 | 15 | 51.5 | 1 |
| 318 | 868.8 | 569.5 | 2.19 | TAG(52:6/FA18:1)+NH4 | 80 | 10 | 38 | 15 | 54.9 | 1 |
| 319 | 868.8 | 571.5 | 2.19 | TAG(52:6/FA18:2)+NH4 | 80 | 10 | 38 | 15 | 87.6 | 1 |

| | | | | | | | | | | |
|---|---|---|---|---|---|---|---|---|---|---|
| 320 | 868.8 | 573.5 | 2.21 | TAG(52:6/FA18:3)+NH4 | 80 | 10 | 38 | 15 | 62.1 | 1 |
| 321 | 868.8 | 547.5 | 2.21 | TAG(52:6/FA20:4)+NH4 | 80 | 10 | 38 | 15 | 52.1 | 1 |
| 322 | 868.8 | 549.5 | 2.19 | TAG(52:6/FA20:5)+NH4 | 80 | 10 | 38 | 15 | 64.3 | 1 |
| 323 | 868.8 | 523.472 | 2.2 | TAG(52:6/FA22:6)+NH4 | 80 | 10 | 38 | 15 | 85.6 | 1 |
| 324 | 866.7 | 593.4 | 2.29 | TAG(52:7/FA16:0)+NH4 | 80 | 10 | 38 | 15 | 88.6 | 1.01 |
| 325 | 866.7 | 567.4 | 2.27 | TAG(52:7/FA18:1)+NH4 | 80 | 10 | 38 | 15 | 95.9 | 1 |
| 326 | 866.7 | 547.4 | 2.22 | TAG(52:7/FA20:5)+NH4 | 80 | 10 | 38 | 15 | 94.5 | 1 |
| 327 | 866.7 | 521.4 | 2.2 | TAG(52:7/FA22:6)+NH4 | 80 | 10 | 38 | 15 | 124.5 | 1.04 |
| 328 | 864.8 | 593.5 | 2.23 | TAG(52:8/FA16:1)+NH4 | 80 | 10 | 38 | 15 | 81.4 | 1.01 |
| 329 | 864.8 | 567.5 | 2.24 | TAG(52:8/FA18:2)+NH4 | 80 | 10 | 38 | 15 | 136.1 | 1.02 |
| 330 | 894.8 | 621.5 | 2.23 | TAG(53:0/FA16:0)+NH4 | 80 | 10 | 38 | 15 | 92 | 1 |
| 331 | 892.8 | 619.5 | 2.19 | TAG(53:1/FA16:0)+NH4 | 80 | 10 | 38 | 15 | 68.4 | 1 |
| 332 | 892.8 | 605.5 | 2.28 | TAG(53:1/FA17:0)+NH4 | 80 | 10 | 38 | 15 | 48.8 | 1 |
| 333 | 892.8 | 591.5 | 2.28 | TAG(53:1/FA18:0)+NH4 | 80 | 10 | 38 | 15 | 87.4 | 1 |
| 334 | 892.8 | 593.5 | 2.28 | TAG(53:1/FA18:1)+NH4 | 80 | 10 | 38 | 15 | 94 | 1 |
| 335 | 890.8 | 617.5 | 2.19 | TAG(53:2/FA16:0)+NH4 | 80 | 10 | 38 | 15 | 88.9 | 1 |
| 336 | 890.8 | 603.5 | 2.25 | TAG(53:2/FA17:0)+NH4 | 80 | 10 | 38 | 15 | 70.6 | 1 |
| 337 | 890.8 | 591.5 | 2.24 | TAG(53:2/FA18:1)+NH4 | 80 | 10 | 38 | 15 | 86.5 | 1 |
| 338 | 890.8 | 593.5 | 2.22 | TAG(53:2/FA18:2)+NH4 | 80 | 10 | 38 | 15 | 84.8 | 1 |
| 339 | 888.8 | 615.5 | 2.2 | TAG(53:3/FA16:0)+NH4 | 80 | 10 | 38 | 15 | 82.6 | 1 |
| 340 | 888.8 | 601.5 | 2.21 | TAG(53:3/FA17:0)+NH4 | 80 | 10 | 38 | 15 | 60 | 1 |
| 341 | 888.8 | 591.5 | 2.2 | TAG(53:3/FA18:2)+NH4 | 80 | 10 | 38 | 15 | 65 | 1 |
| 342 | 886.8 | 613.5 | 2.19 | TAG(53:4/FA16:0)+NH4 | 80 | 10 | 38 | 15 | 60.2 | 1 |
| 343 | 886.8 | 599.5 | 2.19 | TAG(53:4/FA17:0)+NH4 | 80 | 10 | 38 | 15 | 51.1 | 1 |
| 344 | 886.8 | 589.5 | 2.19 | TAG(53:4/FA18:2)+NH4 | 80 | 10 | 38 | 15 | 52.6 | 1 |
| 345 | 886.8 | 591.5 | 2.18 | TAG(53:4/FA18:3)+NH4 | 80 | 10 | 38 | 15 | 47.2 | 1 |
| 346 | 886.8 | 565.5 | 2.23 | TAG(53:4/FA20:4)+NH4 | 80 | 10 | 38 | 15 | 54.2 | 1 |
| 347 | 884.8 | 563.5 | 2.21 | TAG(53:5/FA20:4)+NH4 | 80 | 10 | 38 | 15 | 60.3 | 1.01 |
| 348 | 882.8 | 561.5 | 2.19 | TAG(53:6/FA20:4)+NH4 | 80 | 10 | 38 | 15 | 68 | 1 |
| 349 | 908.8 | 635.5 | 2.36 | TAG(54:0/FA16:0)+NH4 | 80 | 10 | 38 | 15 | 54.8 | 1.01 |
| 350 | 908.8 | 607.5 | 2.38 | TAG(54:0/FA18:0)+NH4 | 80 | 10 | 38 | 15 | 82.1 | 1 |
| 351 | 906.8 | 633.5 | 2.29 | TAG(54:1/FA16:0)+NH4 | 80 | 10 | 38 | 15 | 90.2 | 1 |
| 352 | 906.8 | 605.5 | 2.32 | TAG(54:1/FA18:0)+NH4 | 80 | 10 | 38 | 15 | 92.2 | 1 |
| 353 | 906.8 | 607.5 | 2.3 | TAG(54:1/FA18:1)+NH4 | 80 | 10 | 38 | 15 | 76 | 1 |
| 354 | 906.8 | 577.5 | 2.2 | TAG(54:1/FA20:0)+NH4 | 80 | 10 | 38 | 15 | 80.6 | 1 |
| 355 | 906.8 | 579.5 | 2.21 | TAG(54:1/FA20:1)+NH4 | 80 | 10 | 38 | 15 | 69.9 | 1 |
| 356 | 904.8 | 631.5 | 2.22 | TAG(54:2/FA16:0)+NH4 | 80 | 10 | 38 | 15 | 74.3 | 1 |
| 357 | 904.8 | 603.5 | 2.27 | TAG(54:2/FA18:0)+NH4 | 80 | 10 | 38 | 15 | 64.6 | 1 |
| 358 | 904.8 | 605.5 | 2.27 | TAG(54:2/FA18:1)+NH4 | 80 | 10 | 38 | 15 | 82.2 | 1 |
| 359 | 904.8 | 607.5 | 2.26 | TAG(54:2/FA18:2)+NH4 | 80 | 10 | 38 | 15 | 89.6 | 1 |
| 360 | 904.8 | 575.5 | 2.19 | TAG(54:2/FA20:0)+NH4 | 80 | 10 | 38 | 15 | 93.4 | 1 |
| 361 | 904.8 | 577.5 | 2.2 | TAG(54:2/FA20:1)+NH4 | 80 | 10 | 38 | 15 | 64.4 | 1 |
| 362 | 904.8 | 579.5 | 2.21 | TAG(54:2/FA20:2)+NH4 | 80 | 10 | 38 | 15 | 65.4 | 1 |
| 363 | 902.8 | 629.5 | 2.21 | TAG(54:3/FA16:0)+NH4 | 80 | 10 | 38 | 15 | 92.4 | 1 |
| 364 | 902.8 | 631.5 | 2.19 | TAG(54:3/FA16:1)+NH4 | 80 | 10 | 38 | 15 | 64.1 | 1 |
| 365 | 902.8 | 601.5 | 2.21 | TAG(54:3/FA18:0)+NH4 | 80 | 10 | 38 | 15 | 34.6 | 1 |

| | | | | | | | | | | |
|---|---|---|---|---|---|---|---|---|---|---|
| 366 | 902.8 | 603.5 | 2.21 | TAG(54:3/FA18:1)+NH4 | 80 | 10 | 38 | 15 | 78 | 1 |
| 367 | 902.8 | 605.5 | 2.2 | TAG(54:3/FA18:2)+NH4 | 80 | 10 | 38 | 15 | 76.3 | 1 |
| 368 | 902.8 | 607.5 | 2.21 | TAG(54:3/FA18:3)+NH4 | 80 | 10 | 38 | 15 | 72.8 | 1 |
| 369 | 902.8 | 575.5 | 2.19 | TAG(54:3/FA20:1)+NH4 | 80 | 10 | 38 | 15 | 75.6 | 1 |
| 370 | 902.8 | 577.5 | 2.21 | TAG(54:3/FA20:2)+NH4 | 80 | 10 | 38 | 15 | 43.7 | 1 |
| 371 | 902.8 | 579.5 | 2.19 | TAG(54:3/FA20:3)+NH4 | 80 | 10 | 38 | 15 | 80.9 | 1 |
| 372 | 900.8 | 627.5 | 2.18 | TAG(54:4/FA16:0)+NH4 | 80 | 10 | 38 | 15 | 76.3 | 1 |
| 373 | 900.8 | 629.5 | 2.19 | TAG(54:4/FA16:1)+NH4 | 80 | 10 | 38 | 15 | 49 | 1 |
| 374 | 900.8 | 599.5 | 2.16 | TAG(54:4/FA18:0)+NH4 | 80 | 10 | 38 | 15 | 43.9 | 1 |
| 375 | 900.8 | 601.5 | 2.16 | TAG(54:4/FA18:1)+NH4 | 80 | 10 | 38 | 15 | 63.2 | 1 |
| 376 | 900.8 | 603.5 | 2.17 | TAG(54:4/FA18:2)+NH4 | 80 | 10 | 38 | 15 | 47.4 | 1 |
| 377 | 900.8 | 605.5 | 2.16 | TAG(54:4/FA18:3)+NH4 | 80 | 10 | 38 | 15 | 66.1 | 1 |
| 378 | 900.8 | 573.5 | 2.19 | TAG(54:4/FA20:1)+NH4 | 80 | 10 | 38 | 15 | 50.9 | 1 |
| 379 | 900.8 | 575.5 | 2.18 | TAG(54:4/FA20:2)+NH4 | 80 | 10 | 38 | 15 | 38.2 | 1 |
| 380 | 900.8 | 577.5 | 2.17 | TAG(54:4/FA20:3)+NH4 | 80 | 10 | 38 | 15 | 53.2 | 1 |
| 381 | 900.8 | 579.5 | 2.17 | TAG(54:4/FA20:4)+NH4 | 80 | 10 | 38 | 15 | 69.2 | 1 |
| 382 | 900.8 | 551.503 | 2.17 | TAG(54:4/FA22:4)+NH4 | 80 | 10 | 38 | 15 | 72.5 | 1 |
| 383 | 898.8 | 625.5 | 2.15 | TAG(54:5/FA16:0)+NH4 | 80 | 10 | 38 | 15 | 75.1 | 1 |
| 384 | 898.8 | 627.5 | 2.19 | TAG(54:5/FA16:1)+NH4 | 80 | 10 | 38 | 15 | 47.6 | 1 |
| 385 | 898.8 | 597.5 | 2.13 | TAG(54:5/FA18:0)+NH4 | 80 | 10 | 38 | 15 | 46.1 | 1 |
| 386 | 898.8 | 599.5 | 2.15 | TAG(54:5/FA18:1)+NH4 | 80 | 10 | 38 | 15 | 45.9 | 1 |
| 387 | 898.8 | 601.5 | 2.14 | TAG(54:5/FA18:2)+NH4 | 80 | 10 | 38 | 15 | 50.1 | 1 |
| 388 | 898.8 | 603.5 | 2.14 | TAG(54:5/FA18:3)+NH4 | 80 | 10 | 38 | 15 | 46.9 | 1 |
| 389 | 898.8 | 573.5 | 2.15 | TAG(54:5/FA20:2)+NH4 | 80 | 10 | 38 | 15 | 47.7 | 1 |
| 390 | 898.8 | 575.5 | 2.14 | TAG(54:5/FA20:3)+NH4 | 80 | 10 | 38 | 15 | 49.5 | 1 |
| 391 | 898.8 | 577.5 | 2.16 | TAG(54:5/FA20:4)+NH4 | 80 | 10 | 38 | 15 | 49.6 | 1 |
| 392 | 898.8 | 579.5 | 2.16 | TAG(54:5/FA20:5)+NH4 | 80 | 10 | 38 | 15 | 54.1 | 1 |
| 393 | 898.8 | 549.5 | 2.16 | TAG(54:5/FA22:4)+NH4 | 80 | 10 | 38 | 15 | 58.9 | 1 |
| 394 | 898.8 | 551.503 | 2.13 | TAG(54:5/FA22:5)+NH4 | 80 | 10 | 38 | 15 | 48.5 | 1 |
| 395 | 896.8 | 623.5 | 2.17 | TAG(54:6/FA16:0)+NH4 | 80 | 10 | 38 | 15 | 53.3 | 1 |
| 396 | 896.8 | 625.5 | 2.14 | TAG(54:6/FA16:1)+NH4 | 80 | 10 | 38 | 15 | 46.3 | 1 |
| 397 | 896.8 | 597.5 | 2.14 | TAG(54:6/FA18:1)+NH4 | 80 | 10 | 38 | 15 | 42.2 | 1 |
| 398 | 896.8 | 599.5 | 2.13 | TAG(54:6/FA18:2)+NH4 | 80 | 10 | 38 | 15 | 43.2 | 1 |
| 399 | 896.8 | 601.5 | 2.16 | TAG(54:6/FA18:3)+NH4 | 80 | 10 | 38 | 15 | 39.6 | 1 |
| 400 | 896.8 | 573.5 | 2.17 | TAG(54:6/FA20:3)+NH4 | 80 | 10 | 38 | 15 | 39.9 | 1 |
| 401 | 896.8 | 575.5 | 2.16 | TAG(54:6/FA20:4)+NH4 | 80 | 10 | 38 | 15 | 43.8 | 1 |
| 402 | 896.8 | 577.5 | 2.17 | TAG(54:6/FA20:5)+NH4 | 80 | 10 | 38 | 15 | 41.6 | 1 |
| 403 | 896.8 | 549.5 | 2.16 | TAG(54:6/FA22:5)+NH4 | 80 | 10 | 38 | 15 | 39.7 | 1 |
| 404 | 896.8 | 551.503 | 2.14 | TAG(54:6/FA22:6)+NH4 | 80 | 10 | 38 | 15 | 46 | 1 |
| 405 | 894.8 | 623.5 | 2.19 | TAG(54:7/FA16:1)+NH4 | 80 | 10 | 38 | 15 | 50.2 | 1 |
| 406 | 894.8 | 595.5 | 2.17 | TAG(54:7/FA18:1)+NH4 | 80 | 10 | 38 | 15 | 41.3 | 1 |
| 407 | 894.8 | 597.5 | 2.15 | TAG(54:7/FA18:2)+NH4 | 80 | 10 | 38 | 15 | 42.6 | 1 |
| 408 | 894.8 | 599.5 | 2.15 | TAG(54:7/FA18:3)+NH4 | 80 | 10 | 38 | 15 | 41.3 | 1 |
| 409 | 894.8 | 573.5 | 2.16 | TAG(54:7/FA20:4)+NH4 | 80 | 10 | 38 | 15 | 40.9 | 1 |
| 410 | 894.8 | 575.5 | 2.16 | TAG(54:7/FA20:5)+NH4 | 80 | 10 | 38 | 15 | 39.7 | 1 |
| 411 | 894.8 | 547.5 | 2.17 | TAG(54:7/FA22:5)+NH4 | 80 | 10 | 38 | 15 | 38.1 | 1 |

| | | | | | | | | | | |
|---|---|---|---|---|---|---|---|---|---|---|
| 412 | 894.8 | 549.5 | 2.18 | TAG(54:7/FA22:6)+NH4 | 80 | 10 | 38 | 15 | 65.3 | 1 |
| 413 | 892.8 | 595.5 | 2.16 | TAG(54:8/FA18:2)+NH4 | 80 | 10 | 38 | 15 | 60.5 | 1 |
| 414 | 892.8 | 597.5 | 2.15 | TAG(54:8/FA18:3)+NH4 | 80 | 10 | 38 | 15 | 43.6 | 1 |
| 415 | 892.8 | 571.5 | 2.17 | TAG(54:8/FA20:4)+NH4 | 80 | 10 | 38 | 15 | 37 | 1 |
| 416 | 892.8 | 573.5 | 2.17 | TAG(54:8/FA20:5)+NH4 | 80 | 10 | 38 | 15 | 60.8 | 1.03 |
| 417 | 892.8 | 547.5 | 2.2 | TAG(54:8/FA22:6)+NH4 | 80 | 10 | 38 | 15 | 43.4 | 1.01 |
| 418 | 920.9 | 647.6 | 2.16 | TAG(55:1/FA16:0)+NH4 | 80 | 10 | 38 | 15 | 69.9 | 1 |
| 419 | 920.9 | 621.6 | 2.19 | TAG(55:1/FA18:1)+NH4 | 80 | 10 | 38 | 15 | 47.4 | 1 |
| 420 | 918.8 | 619.5 | 2.17 | TAG(55:2/FA18:1)+NH4 | 80 | 10 | 38 | 15 | 56.6 | 1 |
| 421 | 918.8 | 621.5 | 2.18 | TAG(55:2/FA18:2)+NH4 | 80 | 10 | 38 | 15 | 37.1 | 1 |
| 422 | 916.8 | 617.5 | 2.17 | TAG(55:3/FA18:1)+NH4 | 80 | 10 | 38 | 15 | 43.4 | 1 |
| 423 | 916.8 | 619.5 | 2.16 | TAG(55:3/FA18:2)+NH4 | 80 | 10 | 38 | 15 | 36.1 | 1 |
| 424 | 914.8 | 615.5 | 2.17 | TAG(55:4/FA18:1)+NH4 | 80 | 10 | 38 | 15 | 34.9 | 1 |
| 425 | 914.8 | 617.5 | 2.15 | TAG(55:4/FA18:2)+NH4 | 80 | 10 | 38 | 15 | 35.8 | 1 |
| 426 | 912.8 | 613.5 | 2.18 | TAG(55:5/FA18:1)+NH4 | 80 | 10 | 38 | 15 | 34.9 | 1 |
| 427 | 912.8 | 615.5 | 2.16 | TAG(55:5/FA18:2)+NH4 | 80 | 10 | 38 | 15 | 43.4 | 1 |
| 428 | 912.8 | 591.5 | 2.16 | TAG(55:5/FA20:4)+NH4 | 80 | 10 | 38 | 15 | 36.3 | 1 |
| 429 | 908.8 | 563.5 | 2.17 | TAG(55:7/FA22:6)+NH4 | 80 | 10 | 38 | 15 | 55.2 | 1 |
| 430 | 916.7 | 619.4 | 2.16 | TAG(56:10/FA18:2)+NH4 | 80 | 10 | 38 | 15 | 69.1 | 1.02 |
| 431 | 934.9 | 661.6 | 2.33 | TAG(56:1/FA16:0)+NH4 | 80 | 10 | 38 | 15 | 36.9 | 1 |
| 432 | 934.9 | 635.6 | 2.33 | TAG(56:1/FA18:1)+NH4 | 80 | 10 | 38 | 15 | 79.2 | 1 |
| 433 | 932.9 | 659.6 | 2.28 | TAG(56:2/FA16:0)+NH4 | 80 | 10 | 38 | 15 | 81.1 | 1 |
| 434 | 932.9 | 631.6 | 2.19 | TAG(56:2/FA18:0)+NH4 | 80 | 10 | 38 | 15 | 83.8 | 1 |
| 435 | 932.9 | 603.6 | 2.22 | TAG(56:2/FA20:0)+NH4 | 80 | 10 | 38 | 15 | 50.6 | 1 |
| 436 | 932.9 | 605.6 | 2.17 | TAG(56:2/FA20:1)+NH4 | 80 | 10 | 38 | 15 | 59.8 | 1 |
| 437 | 930.8 | 657.5 | 2.21 | TAG(56:3/FA16:0)+NH4 | 80 | 10 | 38 | 15 | 46 | 1 |
| 438 | 930.8 | 629.5 | 2.16 | TAG(56:3/FA18:0)+NH4 | 80 | 10 | 38 | 15 | 66.6 | 1 |
| 439 | 930.8 | 631.5 | 2.16 | TAG(56:3/FA18:1)+NH4 | 80 | 10 | 38 | 15 | 49.7 | 1 |
| 440 | 930.8 | 633.5 | 2.19 | TAG(56:3/FA18:2)+NH4 | 80 | 10 | 38 | 15 | 41.5 | 1 |
| 441 | 930.8 | 601.5 | 2.22 | TAG(56:3/FA20:0)+NH4 | 80 | 10 | 38 | 15 | 67.8 | 1 |
| 442 | 930.8 | 603.5 | 2.17 | TAG(56:3/FA20:1)+NH4 | 80 | 10 | 38 | 15 | 62.3 | 1 |
| 443 | 930.8 | 605.5 | 2.19 | TAG(56:3/FA20:2)+NH4 | 80 | 10 | 38 | 15 | 39.7 | 1 |
| 444 | 928.8 | 655.5 | 2.16 | TAG(56:4/FA16:0)+NH4 | 80 | 10 | 38 | 15 | 49.7 | 1 |
| 445 | 928.8 | 627.5 | 2.19 | TAG(56:4/FA18:0)+NH4 | 80 | 10 | 38 | 15 | 50.7 | 1 |
| 446 | 928.8 | 629.5 | 2.16 | TAG(56:4/FA18:1)+NH4 | 80 | 10 | 38 | 15 | 64.9 | 1 |
| 447 | 928.8 | 631.5 | 2.15 | TAG(56:4/FA18:2)+NH4 | 80 | 10 | 38 | 15 | 36.4 | 1 |
| 448 | 928.8 | 601.5 | 2.16 | TAG(56:4/FA20:1)+NH4 | 80 | 10 | 38 | 15 | 42.7 | 1 |
| 449 | 928.8 | 603.5 | 2.18 | TAG(56:4/FA20:2)+NH4 | 80 | 10 | 38 | 15 | 35.4 | 1 |
| 450 | 928.8 | 605.5 | 2.18 | TAG(56:4/FA20:3)+NH4 | 80 | 10 | 38 | 15 | 48.2 | 1 |
| 451 | 928.8 | 607.5 | 2.19 | TAG(56:4/FA20:4)+NH4 | 80 | 10 | 38 | 15 | 64.5 | 1 |
| 452 | 928.8 | 579.5 | 2.17 | TAG(56:4/FA22:4)+NH4 | 80 | 10 | 38 | 15 | 77 | 1 |
| 453 | 926.8 | 653.5 | 2.15 | TAG(56:5/FA16:0)+NH4 | 80 | 10 | 38 | 15 | 76.2 | 1 |
| 454 | 926.8 | 625.5 | 2.17 | TAG(56:5/FA18:0)+NH4 | 80 | 10 | 38 | 15 | 50.3 | 1 |
| 455 | 926.8 | 627.5 | 2.15 | TAG(56:5/FA18:1)+NH4 | 80 | 10 | 38 | 15 | 63.7 | 1 |
| 456 | 926.8 | 629.5 | 2.15 | TAG(56:5/FA18:2)+NH4 | 80 | 10 | 38 | 15 | 50.4 | 1 |
| 457 | 926.8 | 599.5 | 2.15 | TAG(56:5/FA20:1)+NH4 | 80 | 10 | 38 | 15 | 36.9 | 1 |

| | | | | | | | | | | |
|---|---|---|---|---|---|---|---|---|---|---|
| 458 | 926.8 | 601.5 | 2.15 | TAG(56:5/FA20:2)+NH4 | 80 | 10 | 38 | 15 | 35.7 | 1 |
| 459 | 926.8 | 603.5 | 2.15 | TAG(56:5/FA20:3)+NH4 | 80 | 10 | 38 | 15 | 38.6 | 1 |
| 460 | 926.8 | 605.5 | 2.16 | TAG(56:5/FA20:4)+NH4 | 80 | 10 | 38 | 15 | 50.3 | 1 |
| 461 | 926.8 | 577.5 | 2.16 | TAG(56:5/FA22:4)+NH4 | 80 | 10 | 38 | 15 | 58.5 | 1 |
| 462 | 926.8 | 579.5 | 2.15 | TAG(56:5/FA22:5)+NH4 | 80 | 10 | 38 | 15 | 52 | 1 |
| 463 | 924.8 | 651.5 | 2.14 | TAG(56:6/FA16:0)+NH4 | 80 | 10 | 38 | 15 | 58.6 | 1 |
| 464 | 924.8 | 623.5 | 2.14 | TAG(56:6/FA18:0)+NH4 | 80 | 10 | 38 | 15 | 44.4 | 1 |
| 465 | 924.8 | 625.5 | 2.14 | TAG(56:6/FA18:1)+NH4 | 80 | 10 | 38 | 15 | 48.9 | 1 |
| 466 | 924.8 | 627.5 | 2.14 | TAG(56:6/FA18:2)+NH4 | 80 | 10 | 38 | 15 | 41 | 1 |
| 467 | 924.8 | 629.5 | 2.14 | TAG(56:6/FA18:3)+NH4 | 80 | 10 | 38 | 15 | 39.2 | 1 |
| 468 | 924.8 | 599.5 | 2.15 | TAG(56:6/FA20:2)+NH4 | 80 | 10 | 38 | 15 | 34.1 | 1 |
| 469 | 924.8 | 601.5 | 2.14 | TAG(56:6/FA20:3)+NH4 | 80 | 10 | 38 | 15 | 36.4 | 1 |
| 470 | 924.8 | 603.5 | 2.14 | TAG(56:6/FA20:4)+NH4 | 80 | 10 | 38 | 15 | 48.3 | 1 |
| 471 | 924.8 | 605.5 | 2.15 | TAG(56:6/FA20:5)+NH4 | 80 | 10 | 38 | 15 | 45.9 | 1 |
| 472 | 924.8 | 575.5 | 2.14 | TAG(56:6/FA22:4)+NH4 | 80 | 10 | 38 | 15 | 40 | 1 |
| 473 | 924.8 | 577.5 | 2.14 | TAG(56:6/FA22:5)+NH4 | 80 | 10 | 38 | 15 | 44 | 1 |
| 474 | 924.8 | 579.5 | 2.15 | TAG(56:6/FA22:6)+NH4 | 80 | 10 | 38 | 15 | 42.7 | 1 |
| 475 | 922.8 | 649.5 | 2.11 | TAG(56:7/FA16:0)+NH4 | 80 | 10 | 38 | 15 | 49.6 | 1 |
| 476 | 922.8 | 651.5 | 2.14 | TAG(56:7/FA16:1)+NH4 | 80 | 10 | 38 | 15 | 47.9 | 1 |
| 477 | 922.8 | 621.5 | 2.18 | TAG(56:7/FA18:0)+NH4 | 80 | 10 | 38 | 15 | 54.7 | 1.01 |
| 478 | 922.8 | 623.5 | 2.14 | TAG(56:7/FA18:1)+NH4 | 80 | 10 | 38 | 15 | 74.7 | 1 |
| 479 | 922.8 | 625.5 | 2.13 | TAG(56:7/FA18:2)+NH4 | 80 | 10 | 38 | 15 | 39.4 | 1 |
| 480 | 922.8 | 627.5 | 2.13 | TAG(56:7/FA18:3)+NH4 | 80 | 10 | 38 | 15 | 40.5 | 1 |
| 481 | 922.8 | 599.5 | 2.14 | TAG(56:7/FA20:3)+NH4 | 80 | 10 | 38 | 15 | 40.1 | 1 |
| 482 | 922.8 | 601.5 | 2.13 | TAG(56:7/FA20:4)+NH4 | 80 | 10 | 38 | 15 | 40.9 | 1 |
| 483 | 922.8 | 603.5 | 2.11 | TAG(56:7/FA20:5)+NH4 | 80 | 10 | 38 | 15 | 42.2 | 1 |
| 484 | 922.8 | 573.5 | 2.14 | TAG(56:7/FA22:4)+NH4 | 80 | 10 | 38 | 15 | 44.6 | 1 |
| 485 | 922.8 | 575.5 | 2.13 | TAG(56:7/FA22:5)+NH4 | 80 | 10 | 38 | 15 | 57 | 1.01 |
| 486 | 922.8 | 577.5 | 2.14 | TAG(56:7/FA22:6)+NH4 | 80 | 10 | 38 | 15 | 41.8 | 1 |
| 487 | 920.8 | 647.5 | 2.14 | TAG(56:8/FA16:0)+NH4 | 80 | 10 | 38 | 15 | 45.6 | 1 |
| 488 | 920.8 | 649.5 | 2.15 | TAG(56:8/FA16:1)+NH4 | 80 | 10 | 38 | 15 | 53.2 | 1 |
| 489 | 920.8 | 621.5 | 2.15 | TAG(56:8/FA18:1)+NH4 | 80 | 10 | 38 | 15 | 69.7 | 1.01 |
| 490 | 920.8 | 623.5 | 2.13 | TAG(56:8/FA18:2)+NH4 | 80 | 10 | 38 | 15 | 55.5 | 1 |
| 491 | 920.8 | 625.5 | 2.13 | TAG(56:8/FA18:3)+NH4 | 80 | 10 | 38 | 15 | 44.9 | 1 |
| 492 | 920.8 | 599.5 | 2.13 | TAG(56:8/FA20:4)+NH4 | 80 | 10 | 38 | 15 | 41.1 | 1 |
| 493 | 920.8 | 601.5 | 2.14 | TAG(56:8/FA20:5)+NH4 | 80 | 10 | 38 | 15 | 41.1 | 1 |
| 494 | 920.8 | 573.5 | 2.13 | TAG(56:8/FA22:5)+NH4 | 80 | 10 | 38 | 15 | 43.4 | 1 |
| 495 | 920.8 | 575.5 | 2.14 | TAG(56:8/FA22:6)+NH4 | 80 | 10 | 38 | 15 | 39 | 1 |
| 496 | 918.8 | 623.5 | 2.13 | TAG(56:9/FA18:3)+NH4 | 80 | 10 | 38 | 15 | 41.5 | 1 |
| 497 | 918.8 | 597.5 | 2.13 | TAG(56:9/FA20:4)+NH4 | 80 | 10 | 38 | 15 | 62.2 | 1.01 |
| 498 | 918.8 | 599.5 | 2.14 | TAG(56:9/FA20:5)+NH4 | 80 | 10 | 38 | 15 | 45.7 | 1 |
| 499 | 918.8 | 573.5 | 2.14 | TAG(56:9/FA22:6)+NH4 | 80 | 10 | 38 | 15 | 37.7 | 1 |
| 500 | 928.7 | 583.4 | 2.26 | TAG(57:10/FA22:6)+NH4 | 80 | 10 | 38 | 15 | 49.1 | 1 |
| 501 | 946.9 | 647.6 | 2.16 | TAG(57:2/FA18:1)+NH4 | 80 | 10 | 38 | 15 | 80.3 | 1.16 |
| 502 | 944.9 | 647.6 | 2.15 | TAG(57:3/FA18:2)+NH4 | 80 | 10 | 38 | 15 | 66.6 | 1 |
| 503 | 944.8 | 647.5 | 2.14 | TAG(58:10/FA18:2)+NH4 | 80 | 10 | 38 | 15 | 51 | 1 |

| | | | | | | | | | | |
|---|---|---|---|---|---|---|---|---|---|---|
| 504 | 944.8 | 623.5 | 2.13 | TAG(58:10/FA20:4)+NH4 | 80 | 10 | 38 | 15 | 52.8 | 1 |
| 505 | 944.8 | 625.5 | 2.13 | TAG(58:10/FA20:5)+NH4 | 80 | 10 | 38 | 15 | 35.6 | 1 |
| 506 | 944.8 | 597.5 | 2.13 | TAG(58:10/FA22:5)+NH4 | 80 | 10 | 38 | 15 | 54.6 | 1.04 |
| 507 | 944.8 | 599.5 | 2.14 | TAG(58:10/FA22:6)+NH4 | 80 | 10 | 38 | 15 | 42.1 | 1.04 |
| 508 | 960.9 | 661.6 | 2.29 | TAG(58:2/FA18:1)+NH4 | 80 | 10 | 38 | 15 | 42.6 | 1 |
| 509 | 958.9 | 659.6 | 2.23 | TAG(58:3/FA18:1)+NH4 | 80 | 10 | 38 | 15 | 78.8 | 1 |
| 510 | 954.9 | 655.6 | 2.15 | TAG(58:5/FA18:1)+NH4 | 80 | 10 | 38 | 15 | 76.1 | 1 |
| 511 | 952.8 | 679.5 | 2.14 | TAG(58:6/FA16:0)+NH4 | 80 | 10 | 38 | 15 | 50.3 | 1 |
| 512 | 952.8 | 651.5 | 2.11 | TAG(58:6/FA18:0)+NH4 | 80 | 10 | 38 | 15 | 51.5 | 1 |
| 513 | 952.8 | 653.5 | 2.14 | TAG(58:6/FA18:1)+NH4 | 80 | 10 | 38 | 15 | 58.9 | 1 |
| 514 | 952.8 | 631.5 | 2.13 | TAG(58:6/FA20:4)+NH4 | 80 | 10 | 38 | 15 | 41.8 | 1 |
| 515 | 952.8 | 603.5 | 2.14 | TAG(58:6/FA22:4)+NH4 | 80 | 10 | 38 | 15 | 37.8 | 1 |
| 516 | 952.8 | 605.5 | 2.14 | TAG(58:6/FA22:5)+NH4 | 80 | 10 | 38 | 15 | 53.4 | 1 |
| 517 | 950.8 | 677.5 | 2.15 | TAG(58:7/FA16:0)+NH4 | 80 | 10 | 38 | 15 | 54 | 1 |
| 518 | 950.8 | 649.5 | 2.12 | TAG(58:7/FA18:0)+NH4 | 80 | 10 | 38 | 15 | 70.2 | 1 |
| 519 | 950.8 | 651.5 | 2.13 | TAG(58:7/FA18:1)+NH4 | 80 | 10 | 38 | 15 | 59.7 | 1 |
| 520 | 950.8 | 653.5 | 2.14 | TAG(58:7/FA18:2)+NH4 | 80 | 10 | 38 | 15 | 43.9 | 1 |
| 521 | 950.8 | 629.5 | 2.14 | TAG(58:7/FA20:4)+NH4 | 80 | 10 | 38 | 15 | 42.7 | 1 |
| 522 | 950.8 | 601.5 | 2.13 | TAG(58:7/FA22:4)+NH4 | 80 | 10 | 38 | 15 | 45.6 | 1 |
| 523 | 950.8 | 603.5 | 2.13 | TAG(58:7/FA22:5)+NH4 | 80 | 10 | 38 | 15 | 52.1 | 1 |
| 524 | 950.8 | 605.5 | 2.14 | TAG(58:7/FA22:6)+NH4 | 80 | 10 | 38 | 15 | 50.1 | 1 |
| 525 | 948.8 | 649.5 | 2.14 | TAG(58:8/FA18:1)+NH4 | 80 | 10 | 38 | 15 | 38.3 | 1 |
| 526 | 948.8 | 651.5 | 2.13 | TAG(58:8/FA18:2)+NH4 | 80 | 10 | 38 | 15 | 57.9 | 1 |
| 527 | 948.8 | 625.5 | 2.13 | TAG(58:8/FA20:3)+NH4 | 80 | 10 | 38 | 15 | 42.3 | 1 |
| 528 | 948.8 | 627.5 | 2.13 | TAG(58:8/FA20:4)+NH4 | 80 | 10 | 38 | 15 | 51.2 | 1.03 |
| 529 | 948.8 | 601.5 | 2.13 | TAG(58:8/FA22:5)+NH4 | 80 | 10 | 38 | 15 | 54.1 | 1 |
| 530 | 948.8 | 603.5 | 2.14 | TAG(58:8/FA22:6)+NH4 | 80 | 10 | 38 | 15 | 39.2 | 1 |
| 531 | 946.8 | 647.5 | 2.15 | TAG(58:9/FA18:1)+NH4 | 80 | 10 | 38 | 15 | 37.8 | 1 |
| 532 | 946.8 | 649.5 | 2.14 | TAG(58:9/FA18:2)+NH4 | 80 | 10 | 38 | 15 | 50.6 | 1 |
| 533 | 946.8 | 625.5 | 2.13 | TAG(58:9/FA20:4)+NH4 | 80 | 10 | 38 | 15 | 46.3 | 1 |
| 534 | 946.8 | 599.5 | 2.13 | TAG(58:9/FA22:5)+NH4 | 80 | 10 | 38 | 15 | 44.4 | 1 |
| 535 | 946.8 | 601.5 | 2.14 | TAG(58:9/FA22:6)+NH4 | 80 | 10 | 38 | 15 | 62.7 | 1 |
| 536 | 972.8 | 625.5 | 2.13 | TAG(60:10/FA22:5)+NH4 | 80 | 10 | 38 | 15 | 35.7 | 1 |
| 537 | 972.8 | 627.5 | 2.23 | TAG(60:10/FA22:6)+NH4 | 80 | 10 | 38 | 15 | 58.2 | 1.06 |
| 538 | 970.8 | 623.5 | 2.13 | TAG(60:11/FA22:5)+NH4 | 80 | 10 | 38 | 15 | 54.2 | 1.02 |
| 539 | 970.8 | 625.5 | 2.13 | TAG(60:11/FA22:6)+NH4 | 80 | 10 | 38 | 15 | 101.8 | 1.2 |
| 540 | 968.8 | 623.5 | 2.23 | TAG(60:12/FA22:6)+NH4 | 80 | 10 | 26 | 15 | 50.1 | 1.03 |
| 541 | 530.4 | 285.2 | 2.72 | DAG(14:0/14:0)+NH4 | 80 | 10 | 26 | 15 | 149.5 | 1.3 |
| 542 | 556.5 | 285.2 | 3.07 | DAG(14:0/16:1)+NH4 | 80 | 10 | 26 | 15 | 145.3 | 1.26 |
| 543 | 586.5 | 313.3 | 2.65 | DAG(16:0/16:0)+NH4 | 80 | 10 | 26 | 15 | 145.3 | 1.14 |
| 544 | 584.4 | 313.2 | 2.22 | DAG(16:0/16:1)+NH4 | 80 | 10 | 26 | 15 | 166.5 | 1.14 |
| 545 | 584.4 | 285.2 | 2.65 | DAG(14:0/18:1)+NH4 | 80 | 10 | 26 | 15 | 145.3 | 1.34 |
| 546 | 582.4 | 311.2 | 2.23 | DAG(16:1/16:1)+NH4 | 80 | 10 | 26 | 15 | 233.1 | 1.05 |
| 547 | 582.4 | 285.2 | 2.37 | DAG(14:0/18:2)+NH4 | 80 | 10 | 26 | 15 | 145.3 | 1.14 |
| 548 | 580.4 | 285.2 | 2.27 | DAG(14:0/18:3)+NH4 | 80 | 10 | 26 | 15 | 154.2 | 1.21 |
| 549 | 614.6 | 285.2 | 1.99 | DAG(14:0/20:0)+NH4 | 80 | 10 | 26 | 15 | 88.4 | 1.08 |

| | | | | | | | | | | |
|---|---|---|---|---|---|---|---|---|---|---|
| 550 | 614.4 | 313.2 | 2.18 | DAG(16:0/18:0)+NH4 | 80 | 10 | 26 | 15 | 109 | 1.12 |
| 551 | 612.6 | 311.3 | 2.72 | DAG(16:1/18:0)+NH4 | 80 | 10 | 26 | 15 | 81.7 | 1.04 |
| 552 | 612.6 | 313.2 | 2.39 | DAG(16:0/18:1)+NH4 | 80 | 10 | 26 | 15 | 145.3 | 1 |
| 553 | 610.4 | 311.2 | 2.2 | DAG(16:1/18:1)+NH4 | 80 | 10 | 26 | 15 | 107.1 | 1.08 |
| 554 | 610.4 | 313.2 | 2.27 | DAG(16:0/18:2)+NH4 | 80 | 10 | 26 | 15 | 70.5 | 1.04 |
| 555 | 608.5 | 311.2 | 2.5 | DAG(16:1/18:2)+NH4 | 80 | 10 | 26 | 15 | 159.8 | 1.36 |
| 556 | 608.5 | 313.2 | 2.21 | DAG(16:0/18:3)+NH4 | 80 | 10 | 26 | 15 | 145.3 | 1.31 |
| 557 | 606.4 | 311.2 | 2.72 | DAG(16:1/18:3)+NH4 | 80 | 10 | 26 | 15 | 89.2 | 1.13 |
| 558 | 606.4 | 285.2 | 3.02 | DAG(14:0/20:4)+NH4 | 80 | 10 | 26 | 15 | 156.4 | 1.18 |
| 559 | 640.6 | 283.2 | 2.82 | DAG(16:1/20:0)+NH4 | 80 | 10 | 26 | 15 | 131.5 | 1.31 |
| 560 | 640.4 | 341.3 | 2.5 | DAG(18:0/18:1)+NH4 | 80 | 10 | 26 | 15 | 134.3 | 1 |
| 561 | 638.4 | 339.3 | 2.23 | DAG(18:1/18:1)+NH4 | 80 | 10 | 26 | 15 | 183.8 | 1.28 |
| 562 | 638.4 | 341.3 | 2.36 | DAG(18:0/18:2)+NH4 | 80 | 10 | 26 | 15 | 92.5 | 1.1 |
| 563 | 636.5 | 339.3 | 2.23 | DAG(18:1/18:2)+NH4 | 80 | 10 | 26 | 15 | 145.3 | 1.67 |
| 564 | 636.5 | 341.3 | 2.37 | DAG(18:0/18:3)+NH4 | 80 | 10 | 26 | 15 | 78.6 | 1.13 |
| 565 | 636.6 | 311.3 | 2.28 | DAG(16:1/20:2)+NH4 | 80 | 10 | 26 | 15 | 145.3 | 1.93 |
| 566 | 636.5 | 313.3 | 2.27 | DAG(16:0/20:3)+NH4 | 80 | 10 | 26 | 15 | 145.3 | 1.5 |
| 567 | 634.5 | 313.3 | 2.3 | DAG(16:0/20:4)+NH4 | 80 | 10 | 26 | 15 | 93.7 | 1.41 |
| 568 | 632.4 | 337.3 | 2.29 | DAG(18:2/18:3)+NH4 | 80 | 10 | 26 | 15 | 145.3 | 1.3 |
| 569 | 632.4 | 311.3 | 2.19 | DAG(16:1/20:4)+NH4 | 80 | 10 | 26 | 15 | 92.9 | 1.15 |
| 570 | 632.4 | 313.3 | 2.32 | DAG(16:0/20:5)+NH4 | 80 | 10 | 26 | 15 | 70.3 | 1.17 |
| 571 | 630.5 | 285.3 | 2.47 | DAG(14:0/22:6)+NH4 | 80 | 10 | 26 | 15 | 163.7 | 1.18 |
| 572 | 666.6 | 339.3 | 2.22 | DAG(18:1/20:1)+NH4 | 80 | 10 | 26 | 15 | 71.7 | 1.4 |
| 573 | 664.6 | 339.3 | 2.24 | DAG(18:1/20:2)+NH4 | 80 | 10 | 26 | 15 | 94.5 | 1.1 |
| 574 | 662.6 | 339.3 | 2.45 | DAG(18:1/20:3)+NH4 | 80 | 10 | 26 | 15 | 90.7 | 1.29 |
| 575 | 660.5 | 337.3 | 2.27 | DAG(18:2/20:3)+NH4 | 80 | 10 | 26 | 15 | 82.2 | 1.51 |
| 576 | 660.5 | 339.3 | 2.23 | DAG(18:1/20:4)+NH4 | 80 | 10 | 26 | 15 | 145.3 | 1.09 |
| 577 | 660.5 | 313.3 | 2.02 | DAG(16:0/22:5)+NH4 | 80 | 10 | 26 | 15 | 76 | 1.22 |
| 578 | 658.5 | 337.3 | 2.24 | DAG(18:2/20:4)+NH4 | 80 | 10 | 26 | 15 | 145.3 | 1.32 |
| 579 | 658.5 | 339.3 | 2.28 | DAG(18:1/20:5)+NH4 | 80 | 10 | 26 | 15 | 107 | 1.33 |
| 580 | 658.5 | 313.3 | 2.26 | DAG(16:0/22:6)+NH4 | 80 | 10 | 26 | 15 | 104.6 | 1.25 |
| 581 | 656.5 | 337.3 | 2.27 | DAG(18:2/20:5)+NH4 | 80 | 10 | 26 | 15 | 145.3 | 1.31 |
| 582 | 656.5 | 311.3 | 2.65 | DAG(16:1/22:6)+NH4 | 80 | 10 | 26 | 15 | 89.3 | 1.24 |
| 583 | 698.6 | 369.3 | 2.23 | DAG(20:0/20:0)+NH4 | 80 | 10 | 26 | 15 | 206 | 1.15 |
| 584 | 688.6 | 339.3 | 2.32 | DAG(18:1/22:4)+NH4 | 80 | 10 | 26 | 15 | 43.8 | 1 |
| 585 | 686.6 | 337.3 | 1.52 | DAG(18:2/22:4)+NH4 | 80 | 10 | 26 | 15 | 74.5 | 1.09 |
| 586 | 686.6 | 339.3 | 2.26 | DAG(18:1/22:5)+NH4 | 80 | 10 | 26 | 15 | 145.3 | 9 |
| 587 | 686.6 | 341.3 | 2.22 | DAG(18:0/22:6)+NH4 | 80 | 10 | 26 | 15 | 101.8 | 1.27 |
| 588 | 684.6 | 337.3 | 2.22 | DAG(18:2/22:5)+NH4 | 80 | 10 | 26 | 15 | 72.1 | 1.02 |
| 589 | 684.6 | 339.3 | 2.17 | DAG(18:1/22:6)+NH4 | 80 | 10 | 26 | 15 | 93.3 | 1.32 |
| 590 | 682.5 | 337.3 | 2.39 | DAG(18:2/22:6)+NH4 | 80 | 10 | 26 | 15 | 141.5 | 1.21 |
| 591 | 313.2 | 239.2 | 2.35 | MAG(16:0)+NH4 | 80 | 10 | 25 | 15 | 178.4 | 1.02 |
| 592 | 311.2 | 237.2 | 2.72 | MAG(16:1)+NH4 | 80 | 10 | 25 | 15 | 157.5 | 1.02 |
| 593 | 341.2 | 267.2 | 2.39 | MAG(18:0)+NH4 | 80 | 10 | 25 | 15 | 184.4 | 1 |
| 594 | 339.2 | 265.2 | 2.65 | MAG(18:1)+NH4 | 80 | 10 | 25 | 15 | 310.6 | 1.03 |
| 595 | 337.2 | 263.2 | 2.32 | MAG(18:2)+NH4 | 80 | 10 | 25 | 15 | 63.1 | 1.01 |

| | | | | | | | | | | |
|---|---|---|---|---|---|---|---|---|---|---|
| 596 | 369.2 | 295.2 | 2.32 | MAG(20:0)+NH4 | 80 | 10 | 25 | 15 | 108.5 | 1.09 |
| 597 | 367.2 | 293.2 | 2.15 | MAG(20:1)+NH4 | 80 | 10 | 25 | 15 | 223.1 | 1.11 |
| 598 | 365.2 | 291.2 | 2.25 | MAG(20:2)+NH4 | 80 | 10 | 25 | 15 | 99.3 | 1.01 |
| 599 | 363.2 | 289.2 | 2.34 | MAG(20:3)+NH4 | 80 | 10 | 25 | 15 | 107.4 | 1.1 |
| 600 | 361.2 | 287.2 | 2.37 | MAG(20:4)+NH4 | 80 | 10 | 25 | 15 | 262.2 | 1.07 |
| 601 | 397.2 | 323.2 | 2.38 | MAG(22:0)+NH4 | 80 | 10 | 25 | 15 | 199 | 1.06 |
| 602 | 395.2 | 321.2 | 2.38 | MAG(22:1)+NH4 | 80 | 10 | 25 | 15 | 148.7 | 1.19 |
| 603 | 393.2 | 319.2 | 2.22 | MAG(22:2)+NH4 | 80 | 10 | 25 | 15 | 155.6 | 1.09 |
| 604 | 391.2 | 317.2 | 2.36 | MAG(22:3)+NH4 | 80 | 10 | 25 | 15 | 130.9 | 1.07 |
| 605 | 389.2 | 315.2 | 2.43 | MAG(22:4)+NH4 | 80 | 10 | 25 | 15 | 184.4 | 1.09 |
| 606 | 387.2 | 313.2 | 2.44 | MAG(22:5)+NH4 | 80 | 10 | 25 | 15 | 182.5 | 1.08 |
| 607 | 385.2 | 311.2 | 2.28 | MAG(22:6)+NH4 | 80 | 10 | 25 | 15 | 196 | 1.06 |
| 608 | 526.317 | 227.202 | 12.81 | LPC(14:0)+AcO | -80 | -10 | -50 | -15 | 38.2 | 1 |
| 609 | 554.346 | 255.233 | 12.65 | LPC(16:0)+AcO | -80 | -10 | -50 | -15 | 39.6 | 1.02 |
| 610 | 552.331 | 253.217 | 12.7 | LPC(16:1)+AcO | -80 | -10 | -50 | -15 | 38.1 | 1 |
| 611 | 582.378 | 283.264 | 12.49 | LPC(18:0)+AcO | -80 | -10 | -50 | -15 | 36.2 | 1 |
| 612 | 580.362 | 281.249 | 12.54 | LPC(18:1)+AcO | -80 | -10 | -50 | -15 | 41 | 1 |
| 613 | 578.346 | 279.233 | 12.6 | LPC(18:2)+AcO | -80 | -10 | -50 | -15 | 38.1 | 1 |
| 614 | 576.331 | 277.217 | 12.65 | LPC(18:3)+AcO | -80 | -10 | -50 | -15 | 39.7 | 1 |
| 615 | 610.409 | 311.3 | 12.37 | LPC(20:0)+AcO | -80 | -10 | -50 | -15 | 44.8 | 1.03 |
| 616 | 608.393 | 309.28 | 12.42 | LPC(20:1)+AcO | -80 | -10 | -50 | -15 | 676.4 | 1.03 |
| 617 | 606.378 | 307.264 | 12.48 | LPC(20:2)+AcO | -80 | -10 | -50 | -15 | 39.2 | 1.02 |
| 618 | 604.362 | 305.249 | 12.49 | LPC(20:3)+AcO | -80 | -10 | -50 | -15 | 37.8 | 1.02 |
| 619 | 602.346 | 303.233 | 12.48 | LPC(20:4)+AcO | -80 | -10 | -50 | -15 | 40.2 | 1 |
| 620 | 600.331 | 301.217 | 12.42 | LPC(20:5)+AcO | -80 | -10 | -50 | -15 | 37.8 | 1 |
| 621 | 630.364 | 331.264 | 12.41 | LPC(22:4)+AcO | -80 | -10 | -50 | -15 | 40.1 | 1.51 |
| 622 | 628.362 | 329.249 | 12.45 | LPC(22:5)+AcO | -80 | -10 | -50 | -15 | 37.6 | 1.11 |
| 623 | 626.346 | 327.233 | 12.42 | LPC(22:6)+AcO | -80 | -10 | -50 | -15 | 41.8 | 1.21 |
| 624 | 736.513 | 227.202 | 10.11 | PC(14:0/14:0)+AcO | -80 | -10 | -50 | -15 | 61.5 | 1 |
| 625 | 790.56 | 281.249 | 9.71 | PC(14:0/18:1)+AcO | -80 | -10 | -50 | -15 | 52.9 | 1.04 |
| 626 | 788.545 | 279.233 | 9.76 | PC(14:0/18:2)+AcO | -80 | -10 | -50 | -15 | 63.1 | 1 |
| 627 | 786.529 | 277.217 | 10.35 | PC(14:0/18:3)+AcO | -80 | -10 | -50 | -15 | 37.6 | 1 |
| 628 | 818.592 | 309.28 | 9.12 | PC(14:0/20:1)+AcO | -80 | -10 | -50 | -15 | 65.4 | 1 |
| 629 | 816.576 | 307.264 | 9.69 | PC(14:0/20:2)+AcO | -80 | -10 | -50 | -15 | 247 | 1.25 |
| 630 | 814.56 | 305.249 | 9.54 | PC(14:0/20:3)+AcO | -80 | -10 | -50 | -15 | 75.6 | 1.12 |
| 631 | 812.545 | 303.233 | 9.36 | PC(14:0/20:4)+AcO | -80 | -10 | -50 | -15 | 53.3 | 1.03 |
| 632 | 810.529 | 301.217 | 10.09 | PC(14:0/20:5)+AcO | -80 | -10 | -50 | -15 | 251.3 | 1 |
| 633 | 840.576 | 331.264 | 9.29 | PC(14:0/22:4)+AcO | -80 | -10 | -50 | -15 | 103.9 | 1.02 |
| 634 | 838.56 | 329.249 | 9.38 | PC(14:0/22:5)+AcO | -80 | -10 | -50 | -15 | 244.2 | 1.2 |
| 635 | 836.545 | 327.233 | 9.41 | PC(14:0/22:6)+AcO | -80 | -10 | -50 | -15 | 299.9 | 1.13 |
| 636 | 732.5 | 225.2 | 9.32 | PC(14:1/14:1)+AcO | -80 | -10 | -50 | -15 | 169.8 | 1.14 |
| 637 | 764.545 | 227.202 | 9.93 | PC(16:0/14:0)+AcO | -80 | -10 | -50 | -15 | 191 | 1.3 |
| 638 | 792.576 | 255.233 | 9.77 | PC(16:0/16:0)+AcO | -80 | -10 | -50 | -15 | 43.3 | 1 |
| 639 | 790.56 | 253.217 | 9.71 | PC(16:0/16:1)+AcO | -80 | -10 | -50 | -15 | 65.3 | 1 |
| 640 | 820.607 | 283.264 | 9.41 | PC(16:0/18:0)+AcO | -80 | -10 | -50 | -15 | 60.3 | 1 |
| 641 | 818.592 | 281.249 | 9.5 | PC(16:0/18:1)+AcO | -80 | -10 | -50 | -15 | 60.5 | 1 |

| | | | | | | | | | | |
|---|---|---|---|---|---|---|---|---|---|---|
| 642 | 816.576 | 279.233 | 9.54 | PC(16:0/18:2)+AcO | -80 | -10 | -50 | -15 | 54.8 | 1 |
| 643 | 814.56 | 277.217 | 9.56 | PC(16:0/18:3)+AcO | -80 | -10 | -50 | -15 | 47 | 1 |
| 644 | 846.623 | 309.28 | 8.18 | PC(16:0/20:1)+AcO | -80 | -10 | -50 | -15 | 70.6 | 1 |
| 645 | 844.607 | 307.264 | 9.44 | PC(16:0/20:2)+AcO | -80 | -10 | -50 | -15 | 86.3 | 1 |
| 646 | 842.592 | 305.249 | 9.37 | PC(16:0/20:3)+AcO | -80 | -10 | -50 | -15 | 46.9 | 1 |
| 647 | 840.576 | 303.233 | 9.18 | PC(16:0/20:4)+AcO | -80 | -10 | -50 | -15 | 41.8 | 1 |
| 648 | 838.56 | 301.217 | 9.25 | PC(16:0/20:5)+AcO | -80 | -10 | -50 | -15 | 37.8 | 1 |
| 649 | 868.607 | 331.264 | 9.16 | PC(16:0/22:4)+AcO | -80 | -10 | -50 | -15 | 51.7 | 1 |
| 650 | 866.592 | 329.249 | 9.23 | PC(16:0/22:5)+AcO | -80 | -10 | -50 | -15 | 57 | 1 |
| 651 | 816.576 | 281.249 | 9.47 | PC(16:1/18:1)+AcO | -80 | -10 | -50 | -15 | 47.1 | 1 |
| 652 | 814.56 | 253.217 | 9.54 | PC(16:1/18:2)+AcO | -80 | -10 | -50 | -15 | 87.9 | 1 |
| 653 | 864.576 | 327.233 | 9.1 | PC(16:0/22:6)+AcO | -80 | -10 | -50 | -15 | 58.1 | 1 |
| 654 | 792.576 | 227.202 | 9.75 | PC(18:0/14:0)+AcO | -80 | -10 | -50 | -15 | 36.2 | 1 |
| 655 | 818.592 | 253.217 | 9.5 | PC(18:0/16:1)+AcO | -80 | -10 | -50 | -15 | 60.8 | 1.04 |
| 656 | 848.639 | 283.264 | 9.34 | PC(18:0/18:0)+AcO | -80 | -10 | -50 | -15 | 71.8 | 1 |
| 657 | 846.623 | 281.249 | 9.35 | PC(18:0/18:1)+AcO | -80 | -10 | -50 | -15 | 68.4 | 1 |
| 658 | 844.607 | 279.233 | 9.37 | PC(18:0/18:2)+AcO | -80 | -10 | -50 | -15 | 51 | 1 |
| 659 | 842.592 | 277.217 | 9.34 | PC(18:0/18:3)+AcO | -80 | -10 | -50 | -15 | 56.8 | 1 |
| 660 | 876.67 | 283.264 | 10.67 | PC(18:0/20:0)+AcO | -80 | -10 | -50 | -15 | 69.3 | 1 |
| 661 | 874.654 | 309.28 | 9.31 | PC(18:0/20:1)+AcO | -80 | -10 | -50 | -15 | 54.8 | 1 |
| 662 | 872.639 | 307.264 | 9.28 | PC(18:0/20:2)+AcO | -80 | -10 | -50 | -15 | 62.1 | 1.02 |
| 663 | 870.623 | 305.249 | 9.21 | PC(18:0/20:3)+AcO | -80 | -10 | -50 | -15 | 63.6 | 1 |
| 664 | 868.607 | 303.233 | 9.01 | PC(18:0/20:4)+AcO | -80 | -10 | -50 | -15 | 59.1 | 1 |
| 665 | 866.592 | 301.217 | 9.06 | PC(18:0/20:5)+AcO | -80 | -10 | -50 | -15 | 44.7 | 1 |
| 666 | 896.639 | 331.264 | 8.99 | PC(18:0/22:4)+AcO | -80 | -10 | -50 | -15 | 45.1 | 1 |
| 667 | 894.623 | 329.249 | 9 | PC(18:0/22:5)+AcO | -80 | -10 | -50 | -15 | 48.8 | 1 |
| 668 | 892.607 | 327.233 | 8.91 | PC(18:0/22:6)+AcO | -80 | -10 | -50 | -15 | 75.7 | 1 |
| 669 | 816.576 | 281.249 | 5.48 | PC(18:1/16:1)+AcO | -80 | -10 | -50 | -15 | 39.5 | 1 |
| 670 | 844.607 | 281.249 | 9.32 | PC(18:1/18:1)+AcO | -80 | -10 | -50 | -15 | 80.2 | 1 |
| 671 | 842.592 | 279.233 | 9.36 | PC(18:1/18:2)+AcO | -80 | -10 | -50 | -15 | 56.7 | 1 |
| 672 | 840.576 | 277.217 | 9.36 | PC(18:1/18:3)+AcO | -80 | -10 | -50 | -15 | 59.3 | 1 |
| 673 | 872.639 | 309.28 | 9.23 | PC(18:1/20:1)+AcO | -80 | -10 | -50 | -15 | 66.2 | 1.01 |
| 674 | 870.623 | 307.264 | 9.26 | PC(18:1/20:2)+AcO | -80 | -10 | -50 | -15 | 63.3 | 1.03 |
| 675 | 868.607 | 305.249 | 9.21 | PC(18:1/20:3)+AcO | -80 | -10 | -50 | -15 | 60.4 | 1 |
| 676 | 866.592 | 303.233 | 9.01 | PC(18:1/20:4)+AcO | -80 | -10 | -50 | -15 | 46.9 | 1 |
| 677 | 864.576 | 301.217 | 9.07 | PC(18:1/20:5)+AcO | -80 | -10 | -50 | -15 | 56.9 | 1 |
| 678 | 894.623 | 331.264 | 7.72 | PC(18:1/22:4)+AcO | -80 | -10 | -50 | -15 | 53.4 | 1.04 |
| 679 | 892.607 | 329.249 | 8.98 | PC(18:1/22:5)+AcO | -80 | -10 | -50 | -15 | 50.9 | 1.03 |
| 680 | 890.592 | 327.233 | 8.91 | PC(18:1/22:6)+AcO | -80 | -10 | -50 | -15 | 79.7 | 1.01 |
| 681 | 814.56 | 279.233 | 9.53 | PC(18:2/16:1)+AcO | -80 | -10 | -50 | -15 | 44.5 | 1.01 |
| 682 | 840.576 | 279.233 | 9.41 | PC(18:2/18:2)+AcO | -80 | -10 | -50 | -15 | 68.8 | 1 |
| 683 | 838.56 | 277.217 | 6.81 | PC(18:2/18:3)+AcO | -80 | -10 | -50 | -15 | 44.7 | 1 |
| 684 | 870.623 | 309.28 | 9.24 | PC(18:2/20:1)+AcO | -80 | -10 | -50 | -15 | 57.5 | 1.01 |
| 685 | 868.607 | 307.264 | 9.29 | PC(18:2/20:2)+AcO | -80 | -10 | -50 | -15 | 56 | 1 |
| 686 | 866.592 | 305.249 | 9.25 | PC(18:2/20:3)+AcO | -80 | -10 | -50 | -15 | 52 | 1 |
| 687 | 864.576 | 303.233 | 9.08 | PC(18:2/20:4)+AcO | -80 | -10 | -50 | -15 | 57.1 | 1 |

| | | | | | | | | | | |
|---|---|---|---|---|---|---|---|---|---|---|
| 688 | 862.56 | 301.217 | 9.12 | PC(18:2/20:5)+AcO | -80 | -10 | -50 | -15 | 54.2 | 1 |
| 689 | 892.607 | 331.264 | 9.02 | PC(18:2/22:4)+AcO | -80 | -10 | -50 | -15 | 247 | 1.1 |
| 690 | 890.592 | 329.249 | 9.02 | PC(18:2/22:5)+AcO | -80 | -10 | -50 | -15 | 117.5 | 1.14 |
| 691 | 888.576 | 327.233 | 8.96 | PC(18:2/22:6)+AcO | -80 | -10 | -50 | -15 | 105.5 | 1.11 |
| 692 | 846.623 | 253.217 | 9.32 | PC(20:0/16:1)+AcO | -80 | -10 | -50 | -15 | 93.6 | 1.15 |
| 693 | 874.654 | 281.249 | 10.68 | PC(20:0/18:1)+AcO | -80 | -10 | -50 | -15 | 160.2 | 1.27 |
| 694 | 870.623 | 277.217 | 9.02 | PC(20:0/18:3)+AcO | -80 | -10 | -50 | -15 | 237.2 | 1 |
| 695 | 902.685 | 309.28 | 4.83 | PC(20:0/20:1)+AcO | -80 | -10 | -50 | -15 | 68.7 | 1.82 |
| 696 | 900.67 | 307.264 | 9.16 | PC(20:0/20:2)+AcO | -80 | -10 | -50 | -15 | 247 | 1.7 |
| 697 | 898.654 | 305.249 | 9 | PC(20:0/20:3)+AcO | -80 | -10 | -50 | -15 | 247 | 2.12 |
| 698 | 896.639 | 303.233 | 8.83 | PC(20:0/20:4)+AcO | -80 | -10 | -50 | -15 | 141.8 | 1.04 |
| 699 | 894.623 | 301.217 | 7.64 | PC(20:0/20:5)+AcO | -80 | -10 | -50 | -15 | 247 | 1 |
| 700 | 924.67 | 331.264 | 8.88 | PC(20:0/22:4)+AcO | -80 | -10 | -50 | -15 | 156.2 | 1.98 |
| 701 | 922.654 | 329.249 | 8.82 | PC(20:0/22:5)+AcO | -80 | -10 | -50 | -15 | 164.4 | 1.68 |
| 702 | 920.639 | 327.233 | 8.79 | PC(20:0/22:6)+AcO | -80 | -10 | -50 | -15 | 247 | 1.48 |
| 703 | 424.247 | 227.202 | 12.76 | LPE(14:0)-H | -80 | -10 | -50 | -15 | 37.6 | 1 |
| 704 | 452.278 | 255.233 | 13.11 | LPE(16:0)-H | -80 | -10 | -40 | -15 | 296.2 | 2.37 |
| 705 | 450.263 | 253.217 | 13.15 | LPE(16:1)-H | -80 | -10 | -40 | -15 | 74.8 | 1 |
| 706 | 480.31 | 283.264 | 12.98 | LPE(18:0)-H | -80 | -10 | -40 | -15 | 106.8 | 1.17 |
| 707 | 478.293 | 281.249 | 13 | LPE(18:1)-H | -80 | -10 | -40 | -15 | 26.9 | 1 |
| 708 | 476.278 | 279.233 | 13.07 | LPE(18:2)-H | -80 | -10 | -40 | -15 | 37.9 | 1 |
| 709 | 474.263 | 277.217 | 13.11 | LPE(18:3)-H | -80 | -10 | -40 | -15 | 43.6 | 1 |
| 710 | 508.341 | 311.3 | 12.87 | LPE(20:0)-H | -80 | -10 | -40 | -15 | 77.8 | 1.18 |
| 711 | 506.325 | 309.28 | 12.89 | LPE(20:1)-H | -80 | -10 | -40 | -15 | 41.3 | 1.22 |
| 712 | 504.31 | 307.264 | 12.95 | LPE(20:2)-H | -80 | -10 | -40 | -15 | 25.9 | 1.1 |
| 713 | 502.294 | 305.249 | 12.97 | LPE(20:3)-H | -80 | -10 | -40 | -15 | 43.1 | 1.17 |
| 714 | 500.278 | 303.233 | 12.93 | LPE(20:4)-H | -80 | -10 | -40 | -15 | 37 | 1.02 |
| 715 | 498.263 | 301.217 | 13.01 | LPE(20:5)-H | -80 | -10 | -40 | -15 | 48.2 | 1 |
| 716 | 528.31 | 331.264 | 12.86 | LPE(22:4)-H | -80 | -10 | -40 | -15 | 79.8 | 2.22 |
| 717 | 526.294 | 329.249 | 12.92 | LPE(22:5)-H | -80 | -10 | -40 | -15 | 40.2 | 1.15 |
| 718 | 524.278 | 327.233 | 9.32 | LPE(22:6)-H | -80 | -10 | -40 | -15 | 64.5 | 1.14 |
| 719 | 634.445 | 227.202 | 10.23 | PE(14:0/14:0)-H | -80 | -10 | -43 | -15 | 53.4 | 1 |
| 720 | 660.461 | 253.217 | 7.47 | PE(14:0/16:1)-H | -80 | -10 | -43 | -15 | 125.9 | 1.26 |
| 721 | 688.492 | 281.249 | 10.41 | PE(14:0/18:1)-H | -80 | -10 | -43 | -15 | 266.2 | 1.24 |
| 722 | 686.477 | 279.233 | 10.47 | PE(14:0/18:2)-H | -80 | -10 | -50 | -15 | 217.1 | 1.02 |
| 723 | 684.461 | 277.217 | 10.09 | PE(14:0/18:3)-H | -80 | -10 | -50 | -15 | 151.2 | 1.02 |
| 724 | 716.524 | 309.28 | 9.9 | PE(14:0/20:1)-H | -80 | -10 | -50 | -15 | 114.5 | 2.14 |
| 725 | 714.508 | 307.264 | 10.73 | PE(14:0/20:2)-H | -80 | -10 | -50 | -15 | 266.2 | 2.88 |
| 726 | 712.492 | 305.249 | 9.39 | PE(14:0/20:3)-H | -80 | -10 | -50 | -15 | 266.2 | 3.18 |
| 727 | 710.477 | 303.233 | 10.16 | PE(14:0/20:4)-H | -80 | -10 | -50 | -15 | 266.2 | 1.47 |
| 728 | 708.461 | 301.217 | 10.16 | PE(14:0/20:5)-H | -80 | -10 | -50 | -15 | 201.4 | 1.21 |
| 729 | 738.508 | 331.264 | 10.16 | PE(14:0/22:4)-H | -80 | -10 | -50 | -15 | 266.2 | 5.08 |
| 730 | 736.492 | 329.249 | 10.14 | PE(14:0/22:5)-H | -80 | -10 | -50 | -15 | 111.9 | 1.88 |
| 731 | 734.477 | 327.233 | 10.19 | PE(14:0/22:6)-H | -80 | -10 | -50 | -15 | 162 | 1.88 |
| 732 | 630.4 | 225.2 | 10.19 | PE(14:1/14:1)-H | -80 | -10 | -50 | -15 | 168.8 | 1.63 |
| 733 | 662.477 | 255.233 | 10.29 | PE(16:0/14:0)-H | -80 | -10 | -50 | -15 | 266.2 | 3.43 |

| | | | | | | | | | | |
|---|---|---|---|---|---|---|---|---|---|---|
| 734 | 690.508 | 255.233 | 10.29 | PE(16:0/16:0)-H | -80 | -10 | -50 | -15 | 228.7 | 1.04 |
| 735 | 688.492 | 253.217 | 10.55 | PE(16:0/16:1)-H | -80 | -10 | -50 | -15 | 56.7 | 1 |
| 736 | 716.524 | 281.249 | 10.53 | PE(16:0/18:1)-H | -80 | -10 | -50 | -15 | 63.3 | 1 |
| 737 | 714.508 | 279.233 | 10.51 | PE(16:0/18:2)-H | -80 | -10 | -50 | -15 | 72 | 1 |
| 738 | 712.492 | 277.217 | 10.53 | PE(16:0/18:3)-H | -80 | -10 | -50 | -15 | 46.3 | 1 |
| 739 | 744.555 | 309.28 | 10.63 | PE(16:0/20:1)-H | -80 | -10 | -50 | -15 | 49.2 | 1 |
| 740 | 742.539 | 307.264 | 10.46 | PE(16:0/20:2)-H | -80 | -10 | -50 | -15 | 109.7 | 1.04 |
| 741 | 740.524 | 305.249 | 10.33 | PE(16:0/20:3)-H | -80 | -10 | -50 | -15 | 68.1 | 1.01 |
| 742 | 738.508 | 303.233 | 10.33 | PE(16:0/20:4)-H | -80 | -10 | -50 | -15 | 266.2 | 1 |
| 743 | 736.492 | 301.217 | 10.19 | PE(16:0/20:5)-H | -80 | -10 | -50 | -15 | 45.2 | 1 |
| 744 | 766.539 | 331.264 | 10.19 | PE(16:0/22:4)-H | -80 | -10 | -50 | -15 | 51.3 | 1.08 |
| 745 | 764.524 | 329.249 | 10.19 | PE(16:0/22:5)-H | -80 | -10 | -50 | -15 | 59.2 | 1 |
| 746 | 762.508 | 327.233 | 10.19 | PE(16:0/22:6)-H | -80 | -10 | -50 | -15 | 60.4 | 1 |
| 747 | 690.508 | 283.264 | 10.77 | PE(18:0/14:0)-H | -80 | -10 | -50 | -15 | 40.6 | 1 |
| 748 | 718.539 | 283.264 | 10.58 | PE(18:0/16:0)-H | -80 | -10 | -50 | -15 | 249.6 | 1.13 |
| 749 | 716.524 | 283.264 | 10.48 | PE(18:0/16:1)-H | -80 | -10 | -50 | -15 | 185.1 | 1 |
| 750 | 746.57 | 283.264 | 10.48 | PE(18:0/18:0)-H | -80 | -10 | -50 | -15 | 77.1 | 1.02 |
| 751 | 744.555 | 281.249 | 10.48 | PE(18:0/18:1)-H | -80 | -10 | -50 | -15 | 50.1 | 1 |
| 752 | 742.539 | 279.233 | 10.48 | PE(18:0/18:2)-H | -80 | -10 | -50 | -15 | 53 | 1 |
| 753 | 740.524 | 277.217 | 10.48 | PE(18:0/18:3)-H | -80 | -10 | -50 | -15 | 121.6 | 1 |
| 754 | 772.586 | 309.28 | 10.48 | PE(18:0/20:1)-H | -80 | -10 | -50 | -15 | 156.7 | 1 |
| 755 | 770.57 | 307.264 | 10.14 | PE(18:0/20:2)-H | -80 | -10 | -50 | -15 | 328.9 | 1 |
| 756 | 768.555 | 305.249 | 10.2 | PE(18:0/20:3)-H | -80 | -10 | -50 | -15 | 67.6 | 1 |
| 757 | 766.539 | 303.233 | 9.51 | PE(18:0/20:4)-H | -80 | -10 | -50 | -15 | 266.2 | 1 |
| 758 | 764.524 | 301.217 | 9.98 | PE(18:0/20:5)-H | -80 | -10 | -50 | -15 | 235.4 | 1 |
| 759 | 794.57 | 331.264 | 9.98 | PE(18:0/22:4)-H | -80 | -10 | -50 | -15 | 228.8 | 1 |
| 760 | 792.555 | 329.249 | 9.33 | PE(18:0/22:5)-H | -80 | -10 | -50 | -15 | 163.6 | 1 |
| 761 | 790.539 | 327.233 | 9.23 | PE(18:0/22:6)-H | -80 | -10 | -50 | -15 | 164.1 | 1 |
| 762 | 714.508 | 281.249 | 10.36 | PE(18:1/16:1)-H | -80 | -10 | -50 | -15 | 212 | 1 |
| 763 | 742.539 | 281.249 | 10.31 | PE(18:1/18:1)-H | -80 | -10 | -50 | -15 | 266.2 | 1 |
| 764 | 740.524 | 279.233 | 10.3 | PE(18:1/18:2)-H | -80 | -10 | -50 | -15 | 63 | 1 |
| 765 | 738.508 | 277.217 | 10.34 | PE(18:1/18:3)-H | -80 | -10 | -50 | -15 | 59.5 | 1 |
| 766 | 770.57 | 309.28 | 10.32 | PE(18:1/20:1)-H | -80 | -10 | -50 | -15 | 64.8 | 1 |
| 767 | 768.555 | 307.264 | 10.32 | PE(18:1/20:2)-H | -80 | -10 | -50 | -15 | 89.8 | 1.03 |
| 768 | 766.539 | 305.249 | 10.2 | PE(18:1/20:3)-H | -80 | -10 | -50 | -15 | 65.1 | 1.03 |
| 769 | 764.524 | 303.233 | 9.13 | PE(18:1/20:4)-H | -80 | -10 | -50 | -15 | 63.5 | 1 |
| 770 | 762.508 | 301.217 | 10.21 | PE(18:1/20:5)-H | -80 | -10 | -50 | -15 | 62.3 | 1 |
| 771 | 792.555 | 331.264 | 9.99 | PE(18:1/22:4)-H | -80 | -10 | -50 | -15 | 147.1 | 1.11 |
| 772 | 790.539 | 329.249 | 9.95 | PE(18:1/22:5)-H | -80 | -10 | -50 | -15 | 113.1 | 1.03 |
| 773 | 788.524 | 327.233 | 9.05 | PE(18:1/22:6)-H | -80 | -10 | -50 | -15 | 85.1 | 1.03 |
| 774 | 712.492 | 279.233 | 10.16 | PE(18:2/16:1)-H | -80 | -10 | -50 | -15 | 59.2 | 1.04 |
| 775 | 738.508 | 279.233 | 10.16 | PE(18:2/18:2)-H | -80 | -10 | -50 | -15 | 81 | 1 |
| 776 | 736.492 | 277.217 | 10.37 | PE(18:2/18:3)-H | -80 | -10 | -50 | -15 | 266.2 | 1 |
| 777 | 768.555 | 309.28 | 10.37 | PE(18:2/20:1)-H | -80 | -10 | -50 | -15 | 43.8 | 1.01 |
| 778 | 766.539 | 307.264 | 10.37 | PE(18:2/20:2)-H | -80 | -10 | -50 | -15 | 59.4 | 1.03 |
| 779 | 764.524 | 305.249 | 8.82 | PE(18:2/20:3)-H | -80 | -10 | -50 | -15 | 51.3 | 1.05 |

| | | | | | | | | | | |
|---|---|---|---|---|---|---|---|---|---|---|
| 780 | 762.508 | 303.233 | 8.82 | PE(18:2/20:4)-H | -80 | -10 | -50 | -15 | 64.9 | 1.03 |
| 781 | 760.492 | 301.217 | 11.53 | PE(18:2/20:5)-H | -80 | -10 | -50 | -15 | 60.8 | 1 |
| 782 | 790.539 | 331.264 | 9.68 | PE(18:2/22:4)-H | -80 | -10 | -50 | -15 | 266.2 | 1.78 |
| 783 | 788.524 | 329.249 | 9.99 | PE(18:2/22:5)-H | -80 | -10 | -50 | -15 | 222.5 | 1.19 |
| 784 | 786.508 | 327.233 | 9.96 | PE(18:2/22:6)-H | -80 | -10 | -50 | -15 | 198.4 | 1.23 |
| 785 | 676.529 | 255.233 | 9.37 | PE(O-16:0/16:0)-H | -80 | -10 | -50 | -15 | 139.6 | 1.48 |
| 786 | 674.513 | 253.217 | 10.67 | PE(O-16:0/16:1)-H | -80 | -10 | -50 | -15 | 266.2 | 1.02 |
| 787 | 704.56 | 283.264 | 9.23 | PE(O-16:0/18:0)-H | -80 | -10 | -50 | -15 | 184.4 | 1.24 |
| 788 | 702.544 | 281.249 | 10.47 | PE(O-16:0/18:1)-H | -80 | -10 | -50 | -15 | 253.1 | 1.12 |
| 789 | 700.529 | 279.233 | 10.36 | PE(O-16:0/18:2)-H | -80 | -10 | -50 | -15 | 112.4 | 1 |
| 790 | 698.513 | 277.217 | 8.97 | PE(O-16:0/18:3)-H | -80 | -10 | -50 | -15 | 121.8 | 1 |
| 791 | 730.576 | 309.28 | 9.18 | PE(O-16:0/20:1)-H | -80 | -10 | -50 | -15 | 71.1 | 1.2 |
| 792 | 728.56 | 307.264 | 10.28 | PE(O-16:0/20:2)-H | -80 | -10 | -50 | -15 | 176.5 | 1.1 |
| 793 | 726.544 | 305.249 | 10.14 | PE(O-16:0/20:3)-H | -80 | -10 | -50 | -15 | 186.6 | 1.19 |
| 794 | 724.529 | 303.233 | 10 | PE(O-16:0/20:4)-H | -80 | -10 | -50 | -15 | 133.6 | 1.01 |
| 795 | 722.513 | 301.217 | 9.58 | PE(O-16:0/20:5)-H | -80 | -10 | -50 | -15 | 114.4 | 1 |
| 796 | 752.56 | 331.264 | 9.88 | PE(O-16:0/22:4)-H | -80 | -10 | -50 | -15 | 68 | 1.05 |
| 797 | 750.544 | 329.249 | 9.88 | PE(O-16:0/22:5)-H | -80 | -10 | -50 | -15 | 110.1 | 1 |
| 798 | 748.529 | 327.233 | 9.78 | PE(O-16:0/22:6)-H | -80 | -10 | -50 | -15 | 88.2 | 1 |
| 799 | 704.56 | 255.233 | 9.78 | PE(O-18:0/16:0)-H | -80 | -10 | -50 | -15 | 82.2 | 1.01 |
| 800 | 702.544 | 253.217 | 9.78 | PE(O-18:0/16:1)-H | -80 | -10 | -50 | -15 | 79.6 | 1 |
| 801 | 732.591 | 283.264 | 9.78 | PE(O-18:0/18:0)-H | -80 | -10 | -50 | -15 | 91.2 | 1 |
| 802 | 730.576 | 281.249 | 9.82 | PE(O-18:0/18:1)-H | -80 | -10 | -50 | -15 | 363.6 | 1.02 |
| 803 | 728.56 | 279.233 | 9.97 | PE(O-18:0/18:2)-H | -80 | -10 | -50 | -15 | 277 | 1 |
| 804 | 726.544 | 277.217 | 10.31 | PE(O-18:0/18:3)-H | -80 | -10 | -50 | -15 | 219.1 | 1 |
| 805 | 758.607 | 309.28 | 8.89 | PE(O-18:0/20:1)-H | -80 | -10 | -50 | -15 | 215.9 | 1.05 |
| 806 | 756.591 | 307.264 | 9.62 | PE(O-18:0/20:2)-H | -80 | -10 | -50 | -15 | 384 | 1.09 |
| 807 | 754.576 | 305.249 | 9.46 | PE(O-18:0/20:3)-H | -80 | -10 | -50 | -15 | 266.2 | 1.06 |
| 808 | 752.56 | 303.233 | 9.48 | PE(O-18:0/20:4)-H | -80 | -10 | -50 | -15 | 150.1 | 1 |
| 809 | 750.544 | 301.217 | 9.8 | PE(O-18:0/20:5)-H | -80 | -10 | -50 | -15 | 207 | 1 |
| 810 | 780.591 | 331.264 | 9.33 | PE(O-18:0/22:4)-H | -80 | -10 | -50 | -15 | 78.9 | 1.02 |
| 811 | 778.576 | 329.249 | 10.08 | PE(O-18:0/22:5)-H | -80 | -10 | -50 | -15 | 226.8 | 1 |
| 812 | 776.56 | 327.233 | 9.99 | PE(O-18:0/22:6)-H | -80 | -10 | -50 | -15 | 116.6 | 1 |
| 813 | 674.5 | 283.264 | 11.46 | PE(P-14:0/18:0)-H | -80 | -10 | -50 | -15 | 68.8 | 1.01 |
| 814 | 672.5 | 281.249 | 10.29 | PE(P-14:0/18:1)-H | -80 | -10 | -50 | -15 | 217.7 | 1.07 |
| 815 | 674.5 | 255.233 | 10.44 | PE(P-16:0/16:0)-H | -80 | -10 | -50 | -15 | 266.2 | 1.06 |
| 816 | 672.5 | 253.217 | 10.36 | PE(P-16:0/16:1)-H | -80 | -10 | -50 | -15 | 195.1 | 1 |
| 817 | 702.5 | 283.264 | 10.3 | PE(P-16:0/18:0)-H | -80 | -10 | -50 | -15 | 76.3 | 1.04 |
| 818 | 700.5 | 281.249 | 10.19 | PE(P-16:0/18:1)-H | -80 | -10 | -50 | -15 | 103.2 | 1.01 |
| 819 | 698.5 | 279.233 | 10.13 | PE(P-16:0/18:2)-H | -80 | -10 | -50 | -15 | 92.4 | 1 |
| 820 | 696.5 | 277.217 | 10.13 | PE(P-16:0/18:3)-H | -80 | -10 | -50 | -15 | 60.7 | 1 |
| 821 | 728.6 | 309.28 | 10.26 | PE(P-16:0/20:1)-H | -80 | -10 | -50 | -15 | 266.2 | 1.02 |
| 822 | 726.5 | 307.264 | 10.09 | PE(P-16:0/20:2)-H | -80 | -10 | -50 | -15 | 137.7 | 1.01 |
| 823 | 724.5 | 305.249 | 9.93 | PE(P-16:0/20:3)-H | -80 | -10 | -50 | -15 | 113 | 1.02 |
| 824 | 722.5 | 303.233 | 9.48 | PE(P-16:0/20:4)-H | -80 | -10 | -50 | -15 | 97.3 | 1 |
| 825 | 720.5 | 301.217 | 10.09 | PE(P-16:0/20:5)-H | -80 | -10 | -50 | -15 | 59.2 | 1 |

| | | | | | | | | | | |
|---|---|---|---|---|---|---|---|---|---|---|
| 826 | 750.5 | 331.264 | 10.09 | PE(P-16:0/22:4)-H | -80 | -10 | -50 | -15 | 59.1 | 1.01 |
| 827 | 748.5 | 329.249 | 10.09 | PE(P-16:0/22:5)-H | -80 | -10 | -50 | -15 | 56.3 | 1 |
| 828 | 746.5 | 327.233 | 10.09 | PE(P-16:0/22:6)-H | -80 | -10 | -50 | -15 | 55.7 | 1 |
| 829 | 698.5 | 281.249 | 9.61 | PE(P-16:1/18:1)-H | -80 | -10 | -50 | -15 | 45.4 | 1 |
| 830 | 702.5 | 255.233 | 9.98 | PE(P-18:0/16:0)-H | -80 | -10 | -50 | -15 | 305.3 | 1.05 |
| 831 | 700.5 | 253.217 | 9.83 | PE(P-18:0/16:1)-H | -80 | -10 | -50 | -15 | 266.2 | 1 |
| 832 | 730.6 | 283.264 | 10.22 | PE(P-18:0/18:0)-H | -80 | -10 | -50 | -15 | 266.2 | 1 |
| 833 | 728.6 | 281.249 | 10.01 | PE(P-18:0/18:1)-H | -80 | -10 | -50 | -15 | 218.9 | 1.01 |
| 834 | 726.5 | 279.233 | 10.07 | PE(P-18:0/18:2)-H | -80 | -10 | -50 | -15 | 232.1 | 1 |
| 835 | 724.5 | 277.217 | 10.07 | PE(P-18:0/18:3)-H | -80 | -10 | -50 | -15 | 83.6 | 1 |
| 836 | 756.6 | 309.28 | 10.07 | PE(P-18:0/20:1)-H | -80 | -10 | -50 | -15 | 84.9 | 1 |
| 837 | 754.6 | 307.264 | 9.67 | PE(P-18:0/20:2)-H | -80 | -10 | -50 | -15 | 284.2 | 1.03 |
| 838 | 752.6 | 305.249 | 9.67 | PE(P-18:0/20:3)-H | -80 | -10 | -50 | -15 | 224.9 | 1.01 |
| 839 | 750.5 | 303.233 | 9.67 | PE(P-18:0/20:4)-H | -80 | -10 | -50 | -15 | 98.6 | 1 |
| 840 | 748.5 | 301.217 | 9.67 | PE(P-18:0/20:5)-H | -80 | -10 | -50 | -15 | 68.6 | 1 |
| 841 | 778.6 | 331.264 | 9.67 | PE(P-18:0/22:4)-H | -80 | -10 | -50 | -15 | 53.2 | 1 |
| 842 | 776.6 | 329.249 | 9.67 | PE(P-18:0/22:5)-H | -80 | -10 | -50 | -15 | 77.9 | 1 |
| 843 | 774.5 | 327.233 | 9.67 | PE(P-18:0/22:6)-H | -80 | -10 | -50 | -15 | 64.3 | 1 |
| 844 | 700.5 | 255.233 | 10.21 | PE(P-18:1/16:0)-H | -80 | -10 | -50 | -15 | 65.4 | 1 |
| 845 | 698.5 | 253.217 | 10.21 | PE(P-18:1/16:1)-H | -80 | -10 | -50 | -15 | 86.2 | 1 |
| 846 | 728.6 | 283.2 | 10.21 | PE(P-18:1/18:0)-H | -80 | -10 | -50 | -15 | 77.5 | 1.04 |
| 847 | 726.5 | 281.249 | 10.21 | PE(P-18:1/18:1)-H | -80 | -10 | -50 | -15 | 160.3 | 1.02 |
| 848 | 724.5 | 279.233 | 10.21 | PE(P-18:1/18:2)-H | -80 | -10 | -50 | -15 | 85.2 | 1 |
| 849 | 722.5 | 277.217 | 10.21 | PE(P-18:1/18:3)-H | -80 | -10 | -50 | -15 | 61.1 | 1 |
| 850 | 754.6 | 309.28 | 10.21 | PE(P-18:1/20:1)-H | -80 | -10 | -50 | -15 | 266.2 | 1.06 |
| 851 | 752.6 | 307.264 | 10.21 | PE(P-18:1/20:2)-H | -80 | -10 | -50 | -15 | 126.8 | 1.06 |
| 852 | 750.5 | 305.249 | 10.21 | PE(P-18:1/20:3)-H | -80 | -10 | -50 | -15 | 266.2 | 1.05 |
| 853 | 748.5 | 303.233 | 10.21 | PE(P-18:1/20:4)-H | -80 | -10 | -50 | -15 | 80.1 | 1 |
| 854 | 746.5 | 301.217 | 10.21 | PE(P-18:1/20:5)-H | -80 | -10 | -50 | -15 | 55 | 1 |
| 855 | 776.6 | 331.264 | 10.21 | PE(P-18:1/22:4)-H | -80 | -10 | -50 | -15 | 266.2 | 1.02 |
| 856 | 774.5 | 329.249 | 10.21 | PE(P-18:1/22:5)-H | -80 | -10 | -50 | -15 | 70.5 | 1 |
| 857 | 772.5 | 327.233 | 10.21 | PE(P-18:1/22:6)-H | -80 | -10 | -50 | -15 | 74.6 | 1 |
| 858 | 722.5 | 279.233 | 10.21 | PE(P-18:2/18:2)-H | -80 | -10 | -50 | -15 | 266.2 | 1 |
| 859 | 746.5 | 303.233 | 10.21 | PE(P-18:2/20:4)-H | -80 | -10 | -50 | -15 | 50.5 | 1 |
| 860 | 770.5 | 327.233 | 10.21 | PE(P-18:2/22:6)-H | -80 | -10 | -50 | -15 | 48.6 | 1 |
| 861 | 455.241 | 227.202 | 11.26 | LPG(14:0)-H | -80 | -10 | -50 | -15 | 55.5 | 1.14 |
| 862 | 483.273 | 255.233 | 11.95 | LPG(16:0)-H | -80 | -10 | -50 | -15 | 413.3 | 8.72 |
| 863 | 481.257 | 253.217 | 11.69 | LPG(16:1)-H | -80 | -10 | -50 | -15 | 413.3 | 1.09 |
| 864 | 511.304 | 283.264 | 12.35 | LPG(18:0)-H | -80 | -10 | -50 | -15 | 158 | 2.1 |
| 865 | 509.289 | 281.249 | 11.66 | LPG(18:1)-H | -80 | -10 | -50 | -15 | 238.1 | 1.12 |
| 866 | 507.273 | 279.233 | 11.4 | LPG(18:2)-H | -80 | -10 | -50 | -15 | 38.5 | 1.05 |
| 867 | 505.257 | 277.217 | 11.79 | LPG(18:3)-H | -80 | -10 | -50 | -15 | 59.8 | 1.09 |
| 868 | 539.335 | 311.3 | 12.37 | LPG(20:0)-H | -80 | -10 | -50 | -15 | 413.3 | 2.05 |
| 869 | 537.32 | 309.28 | 12.41 | LPG(20:1)-H | -80 | -10 | -50 | -15 | 413.3 | 1.8 |
| 870 | 535.304 | 307.264 | 10.65 | LPG(20:2)-H | -80 | -10 | -50 | -15 | 413.3 | 3.34 |
| 871 | 533.289 | 305.249 | 11.59 | LPG(20:3)-H | -80 | -10 | -50 | -15 | 163 | 2.46 |

| | | | | | | | | | | |
|---|---|---|---|---|---|---|---|---|---|---|
| 872 | 531.273 | 303.233 | 11.6 | LPG(20:4)-H | -80 | -10 | -50 | -15 | 413.3 | 3.89 |
| 873 | 529.257 | 301.217 | 11.6 | LPG(20:5)-H | -80 | -10 | -50 | -15 | 413.3 | 4.04 |
| 874 | 559.304 | 331.264 | 11.4 | LPG(22:4)-H | -80 | -10 | -50 | -15 | 413.3 | 7.11 |
| 875 | 557.289 | 329.249 | 10.48 | LPG(22:5)-H | -80 | -10 | -50 | -15 | 413.3 | 3.68 |
| 876 | 555.273 | 327.233 | 11.2 | LPG(22:6)-H | -80 | -10 | -50 | -15 | 413.3 | 5.27 |
| 877 | 665.44 | 227.202 | 6.98 | PG(14:0/14:0)-H | -80 | -10 | -50 | -15 | 68.6 | 1 |
| 878 | 661.4 | 225.2 | 6.53 | PG(14:1/14:1)-H | -80 | -10 | -50 | -15 | 63.3 | 1.19 |
| 879 | 719.487 | 281.249 | 10.36 | PG(14:0/18:1)-H | -80 | -10 | -50 | -15 | 620.5 | 9.98 |
| 880 | 717.471 | 279.233 | 10.35 | PG(14:0/18:2)-H | -80 | -10 | -50 | -15 | 127.6 | 1.11 |
| 881 | 715.456 | 277.217 | 10.26 | PG(14:0/18:3)-H | -80 | -10 | -50 | -15 | 63.5 | 1.04 |
| 882 | 747.518 | 309.28 | 9.44 | PG(14:0/20:1)-H | -80 | -10 | -50 | -15 | 153.1 | 1.27 |
| 883 | 745.503 | 307.264 | 6.52 | PG(14:0/20:2)-H | -80 | -10 | -50 | -15 | 105.6 | 2.17 |
| 884 | 743.487 | 305.249 | 10.02 | PG(14:0/20:3)-H | -80 | -10 | -50 | -15 | 620.5 | 9.82 |
| 885 | 741.471 | 303.233 | 2.94 | PG(14:0/20:4)-H | -80 | -10 | -50 | -15 | 96.8 | 1.66 |
| 886 | 739.456 | 301.217 | 10.4 | PG(14:0/20:5)-H | -80 | -10 | -50 | -15 | 101.6 | 1.13 |
| 887 | 769.503 | 331.264 | 10.7 | PG(14:0/22:4)-H | -80 | -10 | -50 | -15 | 95.3 | 1.2 |
| 888 | 767.487 | 329.249 | 10.04 | PG(14:0/22:5)-H | -80 | -10 | -50 | -15 | 119 | 3.61 |
| 889 | 765.471 | 327.233 | 10.33 | PG(14:0/22:6)-H | -80 | -10 | -50 | -15 | 150 | 1.31 |
| 890 | 693.471 | 227.202 | 6.81 | PG(16:0/14:0)-H | -80 | -10 | -50 | -15 | 152.4 | 1.25 |
| 891 | 721.503 | 255.233 | 9.71 | PG(16:0/16:0)-H | -80 | -10 | -50 | -15 | 94.1 | 1.25 |
| 892 | 719.487 | 253.217 | 6.62 | PG(16:0/16:1)-H | -80 | -10 | -50 | -15 | 39.3 | 1 |
| 893 | 749.534 | 283.264 | 8.8 | PG(16:0/18:0)-H | -80 | -10 | -50 | -15 | 318.3 | 1.03 |
| 894 | 747.518 | 281.249 | 6.52 | PG(16:0/18:1)-H | -80 | -10 | -50 | -15 | 136.2 | 1 |
| 895 | 745.503 | 279.233 | 8.3 | PG(16:0/18:2)-H | -80 | -10 | -50 | -15 | 77.9 | 1 |
| 896 | 743.487 | 277.217 | 6.53 | PG(16:0/18:3)-H | -80 | -10 | -50 | -15 | 127.5 | 1 |
| 897 | 775.549 | 309.28 | 8.88 | PG(16:0/20:1)-H | -80 | -10 | -50 | -15 | 98.2 | 1.43 |
| 898 | 773.534 | 307.264 | 6.42 | PG(16:0/20:2)-H | -80 | -10 | -50 | -15 | 211.9 | 1.57 |
| 899 | 771.518 | 305.249 | 6.32 | PG(16:0/20:3)-H | -80 | -10 | -50 | -15 | 63.1 | 1.03 |
| 900 | 769.503 | 303.233 | 9.6 | PG(16:0/20:4)-H | -80 | -10 | -50 | -15 | 78.7 | 1.06 |
| 901 | 767.487 | 301.217 | 9.59 | PG(16:0/20:5)-H | -80 | -10 | -50 | -15 | 620.5 | 1.01 |
| 902 | 797.534 | 331.264 | 9.82 | PG(16:0/22:4)-H | -80 | -10 | -50 | -15 | 231.3 | 1.08 |
| 903 | 795.518 | 329.249 | 10.16 | PG(16:0/22:5)-H | -80 | -10 | -50 | -15 | 620.5 | 1.41 |
| 904 | 793.503 | 327.233 | 10 | PG(16:0/22:6)-H | -80 | -10 | -50 | -15 | 153.2 | 1.39 |
| 905 | 721.503 | 227.202 | 6.81 | PG(18:0/14:0)-H | -80 | -10 | -50 | -15 | 306.2 | 1.43 |
| 906 | 747.518 | 253.217 | 6.46 | PG(18:0/16:1)-H | -80 | -10 | -50 | -15 | 175.1 | 1.53 |
| 907 | 777.565 | 283.264 | 6.44 | PG(18:0/18:0)-H | -80 | -10 | -50 | -15 | 117 | 1.26 |
| 908 | 775.549 | 281.249 | 6.4 | PG(18:0/18:1)-H | -80 | -10 | -50 | -15 | 239.9 | 1.05 |
| 909 | 773.534 | 279.233 | 6.34 | PG(18:0/18:2)-H | -80 | -10 | -50 | -15 | 70.8 | 1 |
| 910 | 771.518 | 277.217 | 6.32 | PG(18:0/18:3)-H | -80 | -10 | -50 | -15 | 61.1 | 1 |
| 911 | 805.596 | 283.264 | 8.77 | PG(18:0/20:0)-H | -80 | -10 | -50 | -15 | 67.8 | 1.53 |
| 912 | 803.581 | 309.28 | 6.34 | PG(18:0/20:1)-H | -80 | -10 | -50 | -15 | 295.9 | 1.03 |
| 913 | 801.565 | 307.264 | 6.31 | PG(18:0/20:2)-H | -80 | -10 | -50 | -15 | 101.3 | 1.47 |
| 914 | 799.549 | 305.249 | 6.32 | PG(18:0/20:3)-H | -80 | -10 | -50 | -15 | 75.4 | 1.05 |
| 915 | 797.534 | 303.233 | 6.32 | PG(18:0/20:4)-H | -80 | -10 | -50 | -15 | 91.5 | 1.17 |
| 916 | 795.518 | 301.217 | 9.11 | PG(18:0/20:5)-H | -80 | -10 | -50 | -15 | 620.5 | 1.06 |
| 917 | 825.565 | 331.264 | 6.32 | PG(18:0/22:4)-H | -80 | -10 | -50 | -15 | 620.5 | 1.39 |

| | | | | | | | | | | |
|---|---|---|---|---|---|---|---|---|---|---|
| 918 | 823.549 | 329.249 | 6.32 | PG(18:0/22:5)-H | -80 | -10 | -50 | -15 | 620.5 | 1.78 |
| 919 | 821.534 | 327.233 | 6.32 | PG(18:0/22:6)-H | -80 | -10 | -50 | -15 | 202.6 | 1.61 |
| 920 | 745.503 | 281.249 | 6.32 | PG(18:1/16:1)-H | -80 | -10 | -50 | -15 | 620.5 | 3.76 |
| 921 | 773.534 | 281.249 | 6.32 | PG(18:1/18:1)-H | -80 | -10 | -50 | -15 | 56.1 | 1 |
| 922 | 771.518 | 279.233 | 6.32 | PG(18:1/18:2)-H | -80 | -10 | -50 | -15 | 45.9 | 1 |
| 923 | 769.503 | 277.217 | 6.32 | PG(18:1/18:3)-H | -80 | -10 | -50 | -15 | 55.6 | 1 |
| 924 | 801.565 | 309.28 | 9.26 | PG(18:1/20:1)-H | -80 | -10 | -50 | -15 | 74.6 | 1.03 |
| 925 | 799.549 | 307.264 | 1.45 | PG(18:1/20:2)-H | -80 | -10 | -50 | -15 | 242.5 | 1.12 |
| 926 | 797.534 | 305.249 | 6.32 | PG(18:1/20:3)-H | -80 | -10 | -50 | -15 | 96 | 1.01 |
| 927 | 795.518 | 303.233 | 8.8 | PG(18:1/20:4)-H | -80 | -10 | -50 | -15 | 62.9 | 1 |
| 928 | 793.503 | 301.217 | 8.8 | PG(18:1/20:5)-H | -80 | -10 | -50 | -15 | 60.2 | 1 |
| 929 | 823.549 | 331.264 | 9.11 | PG(18:1/22:4)-H | -80 | -10 | -50 | -15 | 54.7 | 1.07 |
| 930 | 821.534 | 329.249 | 9.11 | PG(18:1/22:5)-H | -80 | -10 | -50 | -15 | 59.7 | 1.02 |
| 931 | 819.518 | 327.233 | 9.11 | PG(18:1/22:6)-H | -80 | -10 | -50 | -15 | 67.6 | 1.01 |
| 932 | 743.487 | 279.233 | 9.11 | PG(18:2/16:1)-H | -80 | -10 | -50 | -15 | 53.9 | 1.03 |
| 933 | 769.503 | 279.233 | 9.11 | PG(18:2/18:2)-H | -80 | -10 | -50 | -15 | 64.8 | 1 |
| 934 | 767.487 | 277.217 | 9.27 | PG(18:2/18:3)-H | -80 | -10 | -50 | -15 | 61.6 | 1 |
| 935 | 799.549 | 309.28 | 9.19 | PG(18:2/20:1)-H | -80 | -10 | -50 | -15 | 620.5 | 1.14 |
| 936 | 797.534 | 307.264 | 9.19 | PG(18:2/20:2)-H | -80 | -10 | -50 | -15 | 145.2 | 1.3 |
| 937 | 795.518 | 305.249 | 9.19 | PG(18:2/20:3)-H | -80 | -10 | -50 | -15 | 64 | 1.1 |
| 938 | 793.503 | 303.233 | 9.19 | PG(18:2/20:4)-H | -80 | -10 | -50 | -15 | 65.5 | 1 |
| 939 | 791.487 | 301.217 | 7.72 | PG(18:2/20:5)-H | -80 | -10 | -50 | -15 | 59 | 1 |
| 940 | 821.534 | 331.264 | 8.9 | PG(18:2/22:4)-H | -80 | -10 | -50 | -15 | 620.5 | 1.56 |
| 941 | 819.518 | 329.249 | 8.9 | PG(18:2/22:5)-H | -80 | -10 | -50 | -15 | 57.6 | 1.21 |
| 942 | 817.503 | 327.233 | 8.9 | PG(18:2/22:6)-H | -80 | -10 | -50 | -15 | 79.4 | 1.15 |
| 943 | 775.549 | 253.217 | 9.33 | PG(20:0/16:1)-H | -80 | -10 | -50 | -15 | 67 | 1.29 |
| 944 | 803.581 | 281.249 | 9.33 | PG(20:0/18:1)-H | -80 | -10 | -50 | -15 | 58.3 | 1.14 |
| 945 | 801.565 | 279.233 | 9.33 | PG(20:0/18:2)-H | -80 | -10 | -50 | -15 | 61 | 1 |
| 946 | 799.549 | 277.217 | 9.44 | PG(20:0/18:3)-H | -80 | -10 | -50 | -15 | 54.7 | 1 |
| 947 | 831.612 | 309.28 | 9.11 | PG(20:0/20:1)-H | -80 | -10 | -50 | -15 | 620.5 | 1.29 |
| 948 | 829.596 | 307.264 | 9.03 | PG(20:0/20:2)-H | -80 | -10 | -50 | -15 | 620.5 | 1.55 |
| 949 | 827.581 | 305.249 | 8.95 | PG(20:0/20:3)-H | -80 | -10 | -50 | -15 | 620.5 | 1.24 |
| 950 | 825.565 | 303.233 | 8.95 | PG(20:0/20:4)-H | -80 | -10 | -50 | -15 | 55.1 | 1.01 |
| 951 | 823.549 | 301.217 | 8.95 | PG(20:0/20:5)-H | -80 | -10 | -50 | -15 | 37.2 | 1 |
| 952 | 853.596 | 331.264 | 8.77 | PG(20:0/22:4)-H | -80 | -10 | -50 | -15 | 156.4 | 1.31 |
| 953 | 851.581 | 329.249 | 8.77 | PG(20:0/22:5)-H | -80 | -10 | -50 | -15 | 56.3 | 1.02 |
| 954 | 849.565 | 327.233 | 8.77 | PG(20:0/22:6)-H | -80 | -10 | -50 | -15 | 67 | 1.03 |
| 955 | 543.258 | 227.202 | 11.51 | LPI(14:0)-H | -80 | -10 | -50 | -15 | 58.9 | 1.13 |
| 956 | 571.289 | 255.233 | 13.36 | LPI(16:0)-H | -80 | -10 | -50 | -15 | 948.2 | 2.57 |
| 957 | 569.273 | 253.217 | 13.5 | LPI(16:1)-H | -80 | -10 | -50 | -15 | 948.2 | 1.66 |
| 958 | 599.32 | 283.264 | 13.04 | LPI(18:0)-H | -80 | -10 | -50 | -15 | 948.2 | 5.13 |
| 959 | 597.305 | 281.249 | 13.04 | LPI(18:1)-H | -80 | -10 | -50 | -15 | 948.2 | 1.58 |
| 960 | 595.289 | 279.233 | 13.53 | LPI(18:2)-H | -80 | -10 | -50 | -15 | 948.2 | 3.5 |
| 961 | 593.273 | 277.217 | 13.53 | LPI(18:3)-H | -80 | -10 | -50 | -15 | 126.4 | 1.76 |
| 962 | 627.352 | 311.3 | 13.64 | LPI(20:0)-H | -80 | -10 | -50 | -15 | 948.2 | 6.19 |
| 963 | 625.336 | 309.28 | 13.68 | LPI(20:1)-H | -80 | -10 | -50 | -15 | 54.5 | 1.33 |

| | | | | | | | | | | |
|---|---|---|---|---|---|---|---|---|---|---|
| 964 | 623.32 | 307.264 | 13.35 | LPI(20:2)-H | -80 | -10 | -50 | -15 | 948.2 | 2.35 |
| 965 | 621.305 | 305.249 | 13.35 | LPI(20:3)-H | -80 | -10 | -50 | -15 | 948.2 | 4.42 |
| 966 | 619.289 | 303.233 | 13.59 | LPI(20:4)-H | -80 | -10 | -50 | -15 | 948.2 | 9 |
| 967 | 617.273 | 301.217 | 13.56 | LPI(20:5)-H | -80 | -10 | -50 | -15 | 948.2 | 3.42 |
| 968 | 647.32 | 331.264 | 13.61 | LPI(22:4)-H | -80 | -10 | -50 | -15 | 948.2 | 2.22 |
| 969 | 645.305 | 329.249 | 13.61 | LPI(22:5)-H | -80 | -10 | -50 | -15 | 948.2 | 2.99 |
| 970 | 643.289 | 327.233 | 13.61 | LPI(22:6)-H | -80 | -10 | -50 | -15 | 56.2 | 2.41 |
| 971 | 749.4 | 225.2 | 9.28 | PI(14:1/14:1)-H | -80 | -10 | -61 | -15 | 36.3 | 1.03 |
| 972 | 753.456 | 227.202 | 9.91 | PI(14:0/14:0)-H | -80 | -10 | -60 | -15 | 916.5 | 4.06 |
| 973 | 807.503 | 281.249 | 10.76 | PI(14:0/18:1)-H | -80 | -10 | -60 | -15 | 916.5 | 3.03 |
| 974 | 805.487 | 279.233 | 10.72 | PI(14:0/18:2)-H | -80 | -10 | -60 | -15 | 916.5 | 1.17 |
| 975 | 803.472 | 277.217 | 10.72 | PI(14:0/18:3)-H | -80 | -10 | -60 | -15 | 192.2 | 1.01 |
| 976 | 835.534 | 309.28 | 11.06 | PI(14:0/20:1)-H | -80 | -10 | -60 | -15 | 916.5 | 2.24 |
| 977 | 833.518 | 307.264 | 9.59 | PI(14:0/20:2)-H | -80 | -10 | -60 | -15 | 916.5 | 2.02 |
| 978 | 831.503 | 305.249 | 9.94 | PI(14:0/20:3)-H | -80 | -10 | -60 | -15 | 916.5 | 1.72 |
| 979 | 829.487 | 303.233 | 10.01 | PI(14:0/20:4)-H | -80 | -10 | -60 | -15 | 256 | 1.15 |
| 980 | 827.472 | 301.217 | 10.01 | PI(14:0/20:5)-H | -80 | -10 | -60 | -15 | 93.1 | 1.03 |
| 981 | 857.518 | 331.264 | 10.3 | PI(14:0/22:4)-H | -80 | -10 | -60 | -15 | 916.5 | 2.12 |
| 982 | 855.503 | 329.249 | 9.26 | PI(14:0/22:5)-H | -80 | -10 | -60 | -15 | 916.5 | 1.32 |
| 983 | 853.487 | 327.233 | 9.26 | PI(14:0/22:6)-H | -80 | -10 | -60 | -15 | 144.3 | 1.88 |
| 984 | 781.487 | 227.202 | 10.36 | PI(16:0/14:0)-H | -80 | -10 | -60 | -15 | 916.5 | 2.77 |
| 985 | 809.518 | 255.233 | 10.92 | PI(16:0/16:0)-H | -80 | -10 | -60 | -15 | 197.2 | 2.03 |
| 986 | 807.503 | 253.217 | 13.01 | PI(16:0/16:1)-H | -80 | -10 | -60 | -15 | 60.6 | 1 |
| 987 | 837.55 | 283.264 | 11.23 | PI(16:0/18:0)-H | -80 | -10 | -60 | -15 | 29.5 | 1.32 |
| 988 | 835.534 | 281.249 | 12.96 | PI(16:0/18:1)-H | -80 | -10 | -60 | -15 | 85.5 | 1 |
| 989 | 833.518 | 279.233 | 12.94 | PI(16:0/18:2)-H | -80 | -10 | -60 | -15 | 41.3 | 1.01 |
| 990 | 814.561 | 277.218 | 10.92 | PI(16:0/18:3)-H | -80 | -10 | -60 | -15 | 34.4 | 1 |
| 991 | 863.565 | 309.28 | 10.22 | PI(16:0/20:1)-H | -80 | -10 | -60 | -15 | 35.4 | 1 |
| 992 | 861.55 | 307.264 | 12.98 | PI(16:0/20:2)-H | -80 | -10 | -60 | -15 | 246.5 | 2.25 |
| 993 | 859.534 | 305.249 | 12.87 | PI(16:0/20:3)-H | -80 | -10 | -60 | -15 | 44.9 | 1.53 |
| 994 | 857.518 | 303.233 | 12.8 | PI(16:0/20:4)-H | -80 | -10 | -60 | -15 | 916.5 | 1.09 |
| 995 | 855.503 | 301.217 | 10.55 | PI(16:0/20:5)-H | -80 | -10 | -60 | -15 | 42.1 | 1.01 |
| 996 | 885.55 | 331.264 | 12.86 | PI(16:0/22:4)-H | -80 | -10 | -60 | -15 | 916.5 | 1.78 |
| 997 | 883.534 | 329.249 | 12.81 | PI(16:0/22:5)-H | -80 | -10 | -60 | -15 | 58 | 1.94 |
| 998 | 881.518 | 327.233 | 12.89 | PI(16:0/22:6)-H | -80 | -10 | -60 | -15 | 30.9 | 1.37 |
| 999 | 809.518 | 227.202 | 9.74 | PI(18:0/14:0)-H | -80 | -10 | -60 | -15 | 916.5 | 4.31 |
| 1000 | 835.534 | 253.217 | 12.99 | PI(18:0/16:1)-H | -80 | -10 | -60 | -15 | 916.5 | 2.52 |
| 1001 | 865.581 | 283.264 | 10.2 | PI(18:0/18:0)-H | -80 | -10 | -60 | -15 | 44.9 | 1.28 |
| 1002 | 863.565 | 281.249 | 12.96 | PI(18:0/18:1)-H | -80 | -10 | -60 | -15 | 61.2 | 1 |
| 1003 | 861.55 | 279.233 | 12.9 | PI(18:0/18:2)-H | -80 | -10 | -60 | -15 | 57.9 | 1.02 |
| 1004 | 859.534 | 277.217 | 12.9 | PI(18:0/18:3)-H | -80 | -10 | -60 | -15 | 43.8 | 1 |
| 1005 | 893.612 | 283.264 | 12.9 | PI(18:0/20:0)-H | -80 | -10 | -60 | -15 | 65.3 | 1.1 |
| 1006 | 891.597 | 309.28 | 12.74 | PI(18:0/20:1)-H | -80 | -10 | -60 | -15 | 43.9 | 1 |
| 1007 | 889.581 | 307.264 | 12.92 | PI(18:0/20:2)-H | -80 | -10 | -60 | -15 | 62.9 | 4.96 |
| 1008 | 887.565 | 305.249 | 12.82 | PI(18:0/20:3)-H | -80 | -10 | -60 | -15 | 64.5 | 1.36 |
| 1009 | 885.55 | 283.3 | 12.74 | PI(18:0/20:4)-H | -80 | -10 | -60 | -15 | 50.3 | 1 |

| | | | | | | | | | | |
|---|---|---|---|---|---|---|---|---|---|---|
| 1010 | 883.534 | 301.217 | 10.53 | PI(18:0/20:5)-H | -80 | -10 | -60 | -15 | 39.4 | 1 |
| 1011 | 913.581 | 331.264 | 12.74 | PI(18:0/22:4)-H | -80 | -10 | -60 | -15 | 170.7 | 1.41 |
| 1012 | 911.565 | 329.249 | 12.74 | PI(18:0/22:5)-H | -80 | -10 | -60 | -15 | 92.1 | 1.57 |
| 1013 | 909.55 | 327.233 | 10.95 | PI(18:0/22:6)-H | -80 | -10 | -60 | -15 | 49.8 | 1.25 |
| 1014 | 833.518 | 281.249 | 10.95 | PI(18:1/16:1)-H | -80 | -10 | -60 | -15 | 48.7 | 1.62 |
| 1015 | 861.55 | 281.249 | 12.87 | PI(18:1/18:1)-H | -80 | -10 | -60 | -15 | 916.5 | 1.39 |
| 1016 | 859.534 | 279.233 | 10.91 | PI(18:1/18:2)-H | -80 | -10 | -60 | -15 | 42 | 1.01 |
| 1017 | 857.518 | 277.217 | 10.91 | PI(18:1/18:3)-H | -80 | -10 | -60 | -15 | 34.1 | 1.02 |
| 1018 | 889.581 | 309.28 | 13.35 | PI(18:1/20:1)-H | -80 | -10 | -60 | -15 | 86.1 | 1.15 |
| 1019 | 887.565 | 307.264 | 10.82 | PI(18:1/20:2)-H | -80 | -10 | -60 | -15 | 916.5 | 2.04 |
| 1020 | 885.55 | 305.249 | 10.86 | PI(18:1/20:3)-H | -80 | -10 | -60 | -15 | 316.4 | 1.23 |
| 1021 | 883.534 | 303.233 | 9.28 | PI(18:1/20:4)-H | -80 | -10 | -60 | -15 | 195.6 | 1.01 |
| 1022 | 881.518 | 301.217 | 9.25 | PI(18:1/20:5)-H | -80 | -10 | -60 | -15 | 916.5 | 1 |
| 1023 | 911.565 | 331.264 | 9.28 | PI(18:1/22:4)-H | -80 | -10 | -60 | -15 | 916.5 | 1.86 |
| 1024 | 909.55 | 329.249 | 9.28 | PI(18:1/22:5)-H | -80 | -10 | -60 | -15 | 916.5 | 1.37 |
| 1025 | 907.534 | 327.233 | 10.93 | PI(18:1/22:6)-H | -80 | -10 | -60 | -15 | 916.5 | 1.2 |
| 1026 | 831.503 | 279.233 | 10.77 | PI(18:2/16:1)-H | -80 | -10 | -60 | -15 | 916.5 | 1.98 |
| 1027 | 857.518 | 279.233 | 10.54 | PI(18:2/18:2)-H | -80 | -10 | -60 | -15 | 916.5 | 9 |
| 1028 | 855.503 | 277.217 | 10.54 | PI(18:2/18:3)-H | -80 | -10 | -60 | -15 | 67.8 | 1 |
| 1029 | 887.565 | 309.28 | 10.54 | PI(18:2/20:1)-H | -80 | -10 | -60 | -15 | 916.5 | 2.37 |
| 1030 | 885.55 | 307.264 | 10.54 | PI(18:2/20:2)-H | -80 | -10 | -60 | -15 | 222.7 | 1.61 |
| 1031 | 883.534 | 305.249 | 9.21 | PI(18:2/20:3)-H | -80 | -10 | -60 | -15 | 240.4 | 1.58 |
| 1032 | 881.518 | 303.233 | 10.59 | PI(18:2/20:4)-H | -80 | -10 | -60 | -15 | 916.5 | 1.11 |
| 1033 | 879.503 | 301.217 | 9.12 | PI(18:2/20:5)-H | -80 | -10 | -60 | -15 | 110.5 | 1.01 |
| 1034 | 909.55 | 331.264 | 8.46 | PI(18:2/22:4)-H | -80 | -10 | -60 | -15 | 916.5 | 3.68 |
| 1035 | 907.534 | 329.249 | 8.97 | PI(18:2/22:5)-H | -80 | -10 | -60 | -15 | 125.1 | 2.04 |
| 1036 | 905.518 | 327.233 | 9.82 | PI(18:2/22:6)-H | -80 | -10 | -60 | -15 | 916.5 | 1.87 |
| 1037 | 863.565 | 253.217 | 9.82 | PI(20:0/16:1)-H | -80 | -10 | -60 | -15 | 916.5 | 3.04 |
| 1038 | 891.597 | 281.249 | 9.85 | PI(20:0/18:1)-H | -80 | -10 | -60 | -15 | 916.5 | 1.36 |
| 1039 | 889.581 | 279.233 | 10.68 | PI(20:0/18:2)-H | -80 | -10 | -60 | -15 | 916.5 | 1.04 |
| 1040 | 887.565 | 277.217 | 9.06 | PI(20:0/18:3)-H | -80 | -10 | -60 | -15 | 63 | 1.05 |
| 1041 | 919.628 | 309.28 | 9.64 | PI(20:0/20:1)-H | -80 | -10 | -60 | -15 | 916.5 | 1.58 |
| 1042 | 917.612 | 307.264 | 9.05 | PI(20:0/20:2)-H | -80 | -10 | -60 | -15 | 63 | 1.84 |
| 1043 | 915.597 | 305.249 | 9.05 | PI(20:0/20:3)-H | -80 | -10 | -60 | -15 | 916.5 | 1.97 |
| 1044 | 913.581 | 303.233 | 8.91 | PI(20:0/20:4)-H | -80 | -10 | -60 | -15 | 916.5 | 1.26 |
| 1045 | 911.565 | 301.217 | 8.91 | PI(20:0/20:5)-H | -80 | -10 | -60 | -15 | 207.2 | 1.04 |
| 1046 | 939.597 | 329.249 | 8.79 | PI(20:0/22:5)-H | -80 | -10 | -60 | -15 | 916.5 | 3.03 |
| 1047 | 937.581 | 327.233 | 8.79 | PI(20:0/22:6)-H | -80 | -10 | -60 | -15 | 916.5 | 3.37 |
| 1048 | 468.233 | 227.202 | 8.58 | LPS(14:0)-H | -80 | -10 | -50 | -15 | 988 | 7.11 |
| 1049 | 496.268 | 255.233 | 2.35 | LPS(16:0)-H | -80 | -10 | -50 | -15 | 988 | 5.09 |
| 1050 | 494.252 | 253.217 | 2.59 | LPS(16:1)-H | -80 | -10 | -50 | -15 | 988 | 4.04 |
| 1051 | 524.299 | 283.264 | 2.71 | LPS(18:0)-H | -80 | -10 | -50 | -15 | 988 | 3.04 |
| 1052 | 522.284 | 281.249 | 2.81 | LPS(18:1)-H | -80 | -10 | -50 | -15 | 988 | 2.05 |
| 1053 | 520.268 | 279.233 | 2.61 | LPS(18:2)-H | -80 | -10 | -50 | -15 | 988 | 1.43 |
| 1054 | 518.252 | 277.217 | 2.19 | LPS(18:3)-H | -80 | -10 | -50 | -15 | 988 | 2.1 |
| 1055 | 552.331 | 311.3 | 2.55 | LPS(20:0)-H | -80 | -10 | -50 | -15 | 988 | 1.85 |

| | | | | | | | | | | |
|---|---|---|---|---|---|---|---|---|---|---|
| 1056 | 550.315 | 309.28 | 2.55 | LPS(20:1)-H | -80 | -10 | -50 | -15 | 48.5 | 1.11 |
| 1057 | 548.299 | 307.264 | 2.55 | LPS(20:2)-H | -80 | -10 | -50 | -15 | 30.5 | 1.05 |
| 1058 | 546.284 | 305.249 | 2.68 | LPS(20:3)-H | -80 | -10 | -50 | -15 | 988 | 2.88 |
| 1059 | 544.268 | 303.233 | 2.71 | LPS(20:4)-H | -80 | -10 | -50 | -15 | 988 | 2.32 |
| 1060 | 542.252 | 301.217 | 2.64 | LPS(20:5)-H | -80 | -10 | -50 | -15 | 988 | 1.47 |
| 1061 | 572.299 | 331.264 | 2.61 | LPS(22:4)-H | -80 | -10 | -50 | -15 | 165.6 | 2.57 |
| 1062 | 570.284 | 329.249 | 2.55 | LPS(22:5)-H | -80 | -10 | -50 | -15 | 988 | 2.97 |
| 1063 | 568.268 | 327.233 | 2.11 | LPS(22:6)-H | -80 | -10 | -50 | -15 | 988 | 2.66 |
| 1064 | 674.4 | 225.2 | 10.5 | PS(14:1/14:1) | -80 | -10 | -51 | -15 | 106.7 | 1.17 |
| 1065 | 678.435 | 227.202 | 10.71 | PS(14:0/14:0)-H | -80 | -10 | -50 | -15 | 359.3 | 7.77 |
| 1066 | 732.482 | 281.249 | 9.48 | PS(14:0/18:1)-H | -80 | -10 | -50 | -15 | 84.8 | 1.9 |
| 1067 | 730.466 | 279.233 | 9.75 | PS(14:0/18:2)-H | -80 | -10 | -50 | -15 | 268.4 | 1.03 |
| 1068 | 728.451 | 277.217 | 8.64 | PS(14:0/18:3)-H | -80 | -10 | -50 | -15 | 227.8 | 1.01 |
| 1069 | 760.513 | 309.28 | 9.47 | PS(14:0/20:1)-H | -80 | -10 | -50 | -15 | 359.3 | 1.98 |
| 1070 | 758.498 | 307.264 | 10.87 | PS(14:0/20:2)-H | -80 | -10 | -50 | -15 | 113.8 | 2.27 |
| 1071 | 756.482 | 305.249 | 9.8 | PS(14:0/20:3)-H | -80 | -10 | -50 | -15 | 359.3 | 2.73 |
| 1072 | 754.466 | 303.233 | 9.23 | PS(14:0/20:4)-H | -80 | -10 | -50 | -15 | 139.5 | 1.23 |
| 1073 | 752.451 | 301.217 | 9.49 | PS(14:0/20:5)-H | -80 | -10 | -50 | -15 | 237.2 | 1.04 |
| 1074 | 782.498 | 331.264 | 9.84 | PS(14:0/22:4)-H | -80 | -10 | -50 | -15 | 359.3 | 1.21 |
| 1075 | 780.482 | 329.249 | 9.7 | PS(14:0/22:5)-H | -80 | -10 | -50 | -15 | 359.3 | 1.34 |
| 1076 | 778.466 | 327.233 | 9.7 | PS(14:0/22:6)-H | -80 | -10 | -50 | -15 | 228.8 | 1.28 |
| 1077 | 706.466 | 227.202 | 9.7 | PS(16:0/14:0)-H | -80 | -10 | -50 | -15 | 286.2 | 1.48 |
| 1078 | 734.498 | 255.233 | 10.91 | PS(16:0/16:0)-H | -80 | -10 | -50 | -15 | 359.3 | 1.93 |
| 1079 | 732.482 | 253.217 | 10.42 | PS(16:0/16:1)-H | -80 | -10 | -50 | -15 | 79.8 | 1.08 |
| 1080 | 760.513 | 283.264 | 9.4 | PS(16:0/18:0)-H | -80 | -10 | -50 | -15 | 359.3 | 9 |
| 1081 | 760.513 | 281.249 | 10.55 | PS(16:0/18:1)-H | -80 | -10 | -50 | -15 | 70.9 | 1.01 |
| 1082 | 758.498 | 279.233 | 10.73 | PS(16:0/18:2)-H | -80 | -10 | -50 | -15 | 41.2 | 1 |
| 1083 | 756.482 | 277.217 | 10.7 | PS(16:0/18:3)-H | -80 | -10 | -50 | -15 | 35.7 | 1 |
| 1084 | 788.545 | 309.28 | 10.7 | PS(16:0/20:1)-H | -80 | -10 | -50 | -15 | 87 | 1.07 |
| 1085 | 786.529 | 307.264 | 10.7 | PS(16:0/20:2)-H | -80 | -10 | -50 | -15 | 359.3 | 2.47 |
| 1086 | 784.513 | 305.249 | 9.19 | PS(16:0/20:3)-H | -80 | -10 | -50 | -15 | 150.6 | 1.89 |
| 1087 | 782.498 | 303.233 | 10.09 | PS(16:0/20:4)-H | -80 | -10 | -50 | -15 | 359.3 | 1.09 |
| 1088 | 780.482 | 301.217 | 10.77 | PS(16:0/20:5)-H | -80 | -10 | -50 | -15 | 255 | 1.02 |
| 1089 | 810.529 | 331.264 | 9.12 | PS(16:0/22:4)-H | -80 | -10 | -50 | -15 | 73.8 | 1.11 |
| 1090 | 808.513 | 329.249 | 9.25 | PS(16:0/22:5)-H | -80 | -10 | -50 | -15 | 359.3 | 1.42 |
| 1091 | 806.498 | 327.233 | 8.73 | PS(16:0/22:6)-H | -80 | -10 | -50 | -15 | 359.3 | 1.38 |
| 1092 | 734.498 | 227.202 | 10.04 | PS(18:0/14:0)-H | -80 | -10 | -50 | -15 | 359.3 | 1.33 |
| 1093 | 760.513 | 253.217 | 9.79 | PS(18:0/16:1)-H | -80 | -10 | -50 | -15 | 75.8 | 1.48 |
| 1094 | 790.56 | 283.264 | 9.7 | PS(18:0/18:0)-H | -80 | -10 | -50 | -15 | 268.2 | 1.08 |
| 1095 | 788.545 | 281.249 | 9.68 | PS(18:0/18:1)-H | -80 | -10 | -50 | -15 | 62 | 1 |
| 1096 | 786.529 | 279.233 | 9.33 | PS(18:0/18:2)-H | -80 | -10 | -50 | -15 | 59.9 | 1 |
| 1097 | 784.513 | 277.217 | 10.33 | PS(18:0/18:3)-H | -80 | -10 | -50 | -15 | 310.5 | 1 |
| 1098 | 818.592 | 283.264 | 8.89 | PS(18:0/20:0)-H | -80 | -10 | -50 | -15 | 55.5 | 1.03 |
| 1099 | 816.576 | 309.28 | 9.64 | PS(18:0/20:1)-H | -80 | -10 | -50 | -15 | 52.7 | 1 |
| 1100 | 814.56 | 307.264 | 9.84 | PS(18:0/20:2)-H | -80 | -10 | -50 | -15 | 96 | 1.21 |
| 1101 | 812.545 | 305.249 | 9.14 | PS(18:0/20:3)-H | -80 | -10 | -50 | -15 | 134.9 | 1.29 |

| | | | | | | | | | | |
|---|---|---|---|---|---|---|---|---|---|---|
| 1102 | 810.529 | 303.233 | 8.86 | PS(18:0/20:4)-H | -80 | -10 | -50 | -15 | 359.3 | 1.35 |
| 1103 | 808.513 | 301.217 | 10.15 | PS(18:0/20:5)-H | -80 | -10 | -50 | -15 | 99.4 | 1.01 |
| 1104 | 838.56 | 331.264 | 8.84 | PS(18:0/22:4)-H | -80 | -10 | -50 | -15 | 134.1 | 1.12 |
| 1105 | 836.545 | 329.249 | 8.84 | PS(18:0/22:5)-H | -80 | -10 | -50 | -15 | 66.3 | 1.46 |
| 1106 | 834.529 | 327.233 | 8.84 | PS(18:0/22:6)-H | -80 | -10 | -50 | -15 | 359.3 | 1.29 |
| 1107 | 758.498 | 281.249 | 9.4 | PS(18:1/16:1)-H | -80 | -10 | -50 | -15 | 359.3 | 3.03 |
| 1108 | 786.529 | 281.249 | 9.87 | PS(18:1/18:1)-H | -80 | -10 | -50 | -15 | 50.1 | 1 |
| 1109 | 784.513 | 279.233 | 9.27 | PS(18:1/18:2)-H | -80 | -10 | -50 | -15 | 325.9 | 1.01 |
| 1110 | 782.498 | 277.217 | 10.37 | PS(18:1/18:3)-H | -80 | -10 | -50 | -15 | 53.7 | 1 |
| 1111 | 814.56 | 309.28 | 9.88 | PS(18:1/20:1)-H | -80 | -10 | -50 | -15 | 146.7 | 1.1 |
| 1112 | 812.545 | 307.264 | 8.63 | PS(18:1/20:2)-H | -80 | -10 | -50 | -15 | 359.3 | 1.71 |
| 1113 | 810.529 | 305.249 | 9.08 | PS(18:1/20:3)-H | -80 | -10 | -50 | -15 | 151.6 | 1.44 |
| 1114 | 808.513 | 303.233 | 8.81 | PS(18:1/20:4)-H | -80 | -10 | -50 | -15 | 88.3 | 1.03 |
| 1115 | 806.498 | 301.217 | 9.96 | PS(18:1/20:5)-H | -80 | -10 | -50 | -15 | 86.6 | 1 |
| 1116 | 836.545 | 331.264 | 8.83 | PS(18:1/22:4)-H | -80 | -10 | -50 | -15 | 359.3 | 1.23 |
| 1117 | 834.529 | 329.249 | 8.82 | PS(18:1/22:5)-H | -80 | -10 | -50 | -15 | 359.3 | 1.2 |
| 1118 | 832.513 | 327.233 | 8.93 | PS(18:1/22:6)-H | -80 | -10 | -50 | -15 | 110.8 | 1.15 |
| 1119 | 756.482 | 279.233 | 9.43 | PS(18:2/16:1)-H | -80 | -10 | -50 | -15 | 359.3 | 1.28 |
| 1120 | 782.498 | 279.233 | 10.32 | PS(18:2/18:2)-H | -80 | -10 | -50 | -15 | 56.5 | 1 |
| 1121 | 780.482 | 277.217 | 10.56 | PS(18:2/18:3)-H | -80 | -10 | -50 | -15 | 75.3 | 1 |
| 1122 | 812.545 | 309.28 | 10.21 | PS(18:2/20:1)-H | -80 | -10 | -50 | -15 | 204.8 | 1.22 |
| 1123 | 810.529 | 307.264 | 8.79 | PS(18:2/20:2)-H | -80 | -10 | -50 | -15 | 172.6 | 1.25 |
| 1124 | 808.513 | 305.249 | 8.64 | PS(18:2/20:3)-H | -80 | -10 | -50 | -15 | 264.3 | 1.17 |
| 1125 | 806.498 | 303.233 | 8.84 | PS(18:2/20:4)-H | -80 | -10 | -50 | -15 | 431.7 | 1.01 |
| 1126 | 804.482 | 301.217 | 9.33 | PS(18:2/20:5)-H | -80 | -10 | -50 | -15 | 359.3 | 1 |
| 1127 | 834.529 | 331.264 | 7.58 | PS(18:2/22:4)-H | -80 | -10 | -50 | -15 | 168.6 | 1.22 |
| 1128 | 832.513 | 329.249 | 8.93 | PS(18:2/22:5)-H | -80 | -10 | -50 | -15 | 137.8 | 1.14 |
| 1129 | 830.498 | 327.233 | 9.16 | PS(18:2/22:6)-H | -80 | -10 | -50 | -15 | 287.7 | 1.13 |
| 1130 | 788.545 | 253.217 | 9.68 | PS(20:0/16:1)-H | -80 | -10 | -50 | -15 | 185.4 | 1.12 |
| 1131 | 814.56 | 281.249 | 10.9 | PS(20:0/18:1)-H | -80 | -10 | -50 | -15 | 59.7 | 1.02 |
| 1132 | 814.561 | 279.234 | 9.5 | PS(20:0/18:2)-H | -80 | -10 | -50 | -15 | 106.7 | 1.01 |
| 1133 | 812.545 | 277.217 | 9.55 | PS(20:0/18:3)-H | -80 | -10 | -50 | -15 | 54.9 | 1 |
| 1134 | 844.607 | 309.28 | 8.94 | PS(20:0/20:1)-H | -80 | -10 | -50 | -15 | 110.6 | 1.09 |
| 1135 | 842.592 | 307.264 | 9.31 | PS(20:0/20:2)-H | -80 | -10 | -50 | -15 | 58.7 | 1.2 |
| 1136 | 840.576 | 305.249 | 9.34 | PS(20:0/20:3)-H | -80 | -10 | -50 | -15 | 176.9 | 1.09 |
| 1137 | 838.56 | 303.233 | 8.83 | PS(20:0/20:4)-H | -80 | -10 | -50 | -15 | 74.4 | 1 |
| 1138 | 836.545 | 301.217 | 8.64 | PS(20:0/20:5)-H | -80 | -10 | -50 | -15 | 131.1 | 1 |
| 1139 | 866.592 | 331.264 | 8.81 | PS(20:0/22:4)-H | -80 | -10 | -50 | -15 | 359.3 | 1.28 |
| 1140 | 864.576 | 329.249 | 8.96 | PS(20:0/22:5)-H | -80 | -10 | -50 | -15 | 139.2 | 1.08 |
| 1141 | 862.56 | 327.233 | 9.04 | PS(20:0/22:6)-H | -80 | -10 | -50 | -15 | 120 | 2 |
| 1142 | 225.1 | 225.1 | 13.75 | L PA(14:0)-H | -80 | -10 | -50 | -15 | 120 | 2 |
| 1143 | 227.1 | 227.1 | 13.77 | L PA(16:0)-H | -80 | -10 | -50 | -15 | 120 | 2 |
| 1144 | 251.1 | 251.1 | 12.56 | L PA(16:1)-H | -80 | -10 | -50 | -15 | 120 | 2 |
| 1145 | 253.1 | 253.1 | 2.08 | L PA(18:0)-H | -80 | -10 | -50 | -15 | 120 | 2 |
| 1146 | 255.1 | 255.1 | 3.33 | L PA(18:1)-H | -80 | -10 | -50 | -15 | 120 | 2 |
| 1147 | 277.2 | 277.2 | 14.42 | L PA(18:2)-H | -80 | -10 | -50 | -15 | 120 | 2 |

| | | | | | | | | | | |
|---|---|---|---|---|---|---|---|---|---|---|
| 1148 | 591.403 | 227.202 | 0.34 | PA(14:0/14:0)-H | -80 | -10 | -50 | -15 | 120 | 2 |
| 1149 | 645.45 | 281.249 | 12.09 | PA(14:0/18:1)-H | -80 | -10 | -50 | -15 | 120 | 2 |
| 1150 | 643.434 | 279.233 | 11.16 | PA(14:0/18:2)-H | -80 | -10 | -50 | -15 | 120 | 2 |
| 1151 | 641.419 | 277.217 | 8.9 | PA(14:0/18:3)-H | -80 | -10 | -50 | -15 | 120 | 2 |
| 1152 | 673.481 | 309.28 | 8.9 | PA(14:0/20:1)-H | -80 | -10 | -50 | -15 | 120 | 2 |
| 1153 | 671.466 | 307.264 | 9.77 | PA(14:0/20:2)-H | -80 | -10 | -50 | -15 | 120 | 2 |
| 1154 | 669.45 | 305.249 | 9.77 | PA(14:0/20:3)-H | -80 | -10 | -50 | -15 | 120 | 2 |
| 1155 | 667.434 | 303.233 | 9.77 | PA(14:0/20:4)-H | -80 | -10 | -50 | -15 | 120 | 2 |
| 1156 | 665.419 | 301.217 | 9.69 | PA(14:0/20:5)-H | -80 | -10 | -50 | -15 | 120 | 2 |
| 1157 | 695.466 | 331.264 | 9.69 | PA(14:0/22:4)-H | -80 | -10 | -50 | -15 | 120 | 2 |
| 1158 | 693.45 | 329.249 | 9.69 | PA(14:0/22:5)-H | -80 | -10 | -50 | -15 | 120 | 2 |
| 1159 | 691.434 | 327.233 | 9.69 | PA(14:0/22:6)-H | -80 | -10 | -50 | -15 | 120 | 2 |
| 1160 | 619.434 | 227.202 | 10.16 | PA(16:0/14:0)-H | -80 | -10 | -50 | -15 | 120 | 2 |
| 1161 | 647.466 | 255.233 | 9.94 | PA(16:0/16:0)-H | -80 | -10 | -50 | -15 | 120 | 2 |
| 1162 | 645.45 | 253.217 | 9.88 | PA(16:0/16:1)-H | -80 | -10 | -50 | -15 | 120 | 2 |
| 1163 | 675.497 | 283.264 | 11.26 | PA(16:0/18:0)-H | -80 | -10 | -50 | -15 | 120 | 2 |
| 1164 | 673.481 | 281.249 | 10.58 | PA(16:0/18:1)-H | -80 | -10 | -50 | -15 | 120 | 2 |
| 1165 | 671.466 | 279.233 | 9.7 | PA(16:0/18:2)-H | -80 | -10 | -50 | -15 | 120 | 2 |
| 1166 | 669.45 | 277.217 | 10.3 | PA(16:0/18:3)-H | -80 | -10 | -50 | -15 | 120 | 2 |
| 1167 | 701.513 | 309.28 | 10.3 | PA(16:0/20:1)-H | -80 | -10 | -50 | -15 | 120 | 2 |
| 1168 | 699.497 | 307.264 | 9.96 | PA(16:0/20:2)-H | -80 | -10 | -50 | -15 | 120 | 2 |
| 1169 | 697.481 | 305.249 | 10.06 | PA(16:0/20:3)-H | -80 | -10 | -50 | -15 | 120 | 2 |
| 1170 | 695.466 | 303.233 | 9.35 | PA(16:0/20:4)-H | -80 | -10 | -50 | -15 | 120 | 2 |
| 1171 | 693.45 | 301.217 | 11.53 | PA(16:0/20:5)-H | -80 | -10 | -50 | -15 | 120 | 2 |
| 1172 | 723.497 | 331.264 | 11.53 | PA(16:0/22:4)-H | -80 | -10 | -50 | -15 | 120 | 2 |
| 1173 | 721.481 | 329.249 | 9.6 | PA(16:0/22:5)-H | -80 | -10 | -50 | -15 | 120 | 2 |
| 1174 | 719.466 | 327.233 | 9.97 | PA(16:0/22:6)-H | -80 | -10 | -50 | -15 | 120 | 2 |
| 1175 | 647.466 | 227.202 | 10.72 | PA(18:0/14:0)-H | -80 | -10 | -50 | -15 | 120 | 2 |
| 1176 | 673.481 | 253.217 | 10.34 | PA(18:0/16:1)-H | -80 | -10 | -50 | -15 | 120 | 2 |
| 1177 | 703.528 | 283.264 | 10.36 | PA(18:0/18:0)-H | -80 | -10 | -50 | -15 | 120 | 2 |
| 1178 | 701.513 | 281.249 | 10.23 | PA(18:0/18:1)-H | -80 | -10 | -50 | -15 | 120 | 2 |
| 1179 | 699.497 | 279.233 | 10.17 | PA(18:0/18:2)-H | -80 | -10 | -50 | -15 | 120 | 2 |
| 1180 | 697.481 | 277.217 | 10.23 | PA(18:0/18:3)-H | -80 | -10 | -50 | -15 | 120 | 2 |
| 1181 | 731.56 | 283.264 | 10.84 | PA(18:0/20:0)-H | -80 | -10 | -50 | -15 | 120 | 2 |
| 1182 | 729.544 | 309.28 | 10.31 | PA(18:0/20:1)-H | -80 | -10 | -50 | -15 | 120 | 2 |
| 1183 | 727.528 | 307.264 | 10 | PA(18:0/20:2)-H | -80 | -10 | -50 | -15 | 120 | 2 |
| 1184 | 725.513 | 305.249 | 9.93 | PA(18:0/20:3)-H | -80 | -10 | -50 | -15 | 120 | 2 |
| 1185 | 723.497 | 303.233 | 9.72 | PA(18:0/20:4)-H | -80 | -10 | -50 | -15 | 120 | 2 |
| 1186 | 721.481 | 301.217 | 9.76 | PA(18:0/20:5)-H | -80 | -10 | -50 | -15 | 120 | 2 |
| 1187 | 751.528 | 331.264 | 9.73 | PA(18:0/22:4)-H | -80 | -10 | -50 | -15 | 120 | 2 |
| 1188 | 749.513 | 329.249 | 9.77 | PA(18:0/22:5)-H | -80 | -10 | -50 | -15 | 120 | 2 |
| 1189 | 747.497 | 327.233 | 9.64 | PA(18:0/22:6)-H | -80 | -10 | -50 | -15 | 120 | 2 |
| 1190 | 671.466 | 281.249 | 11.41 | PA(18:1/16:1)-H | -80 | -10 | -50 | -15 | 120 | 2 |
| 1191 | 699.497 | 281.249 | 9.69 | PA(18:1/18:1)-H | -80 | -10 | -50 | -15 | 120 | 2 |
| 1192 | 697.481 | 279.233 | 9.68 | PA(18:1/18:2)-H | -80 | -10 | -50 | -15 | 120 | 2 |
| 1193 | 695.466 | 277.217 | 9.68 | PA(18:1/18:3)-H | -80 | -10 | -50 | -15 | 120 | 2 |

| | | | | | | | | | | |
|---|---|---|---|---|---|---|---|---|---|---|
| 1194 | 727.528 | 309.28 | 10.25 | PA(18:1/20:1)-H | -80 | -10 | -50 | -15 | 120 | 2 |
| 1195 | 725.513 | 307.264 | 9.56 | PA(18:1/20:2)-H | -80 | -10 | -50 | -15 | 120 | 2 |
| 1196 | 723.497 | 305.249 | 9.49 | PA(18:1/20:3)-H | -80 | -10 | -50 | -15 | 120 | 2 |
| 1197 | 721.481 | 303.233 | 9.34 | PA(18:1/20:4)-H | -80 | -10 | -50 | -15 | 120 | 2 |
| 1198 | 719.466 | 301.217 | 9.43 | PA(18:1/20:5)-H | -80 | -10 | -50 | -15 | 120 | 2 |
| 1199 | 749.513 | 331.264 | 9.26 | PA(18:1/22:4)-H | -80 | -10 | -50 | -15 | 120 | 2 |
| 1200 | 747.497 | 329.249 | 9.43 | PA(18:1/22:5)-H | -80 | -10 | -50 | -15 | 120 | 2 |
| 1201 | 745.481 | 327.233 | 9.28 | PA(18:1/22:6)-H | -80 | -10 | -50 | -15 | 120 | 2 |
| 1202 | 669.45 | 279.233 | 9.76 | PA(18:2/16:1)-H | -80 | -10 | -50 | -15 | 120 | 2 |
| 1203 | 695.466 | 279.233 | 9.7 | PA(18:2/18:2)-H | -80 | -10 | -50 | -15 | 120 | 2 |
| 1204 | 693.45 | 277.217 | 9.7 | PA(18:2/18:3)-H | -80 | -10 | -50 | -15 | 120 | 2 |
| 1205 | 725.513 | 309.28 | 10.8 | PA(18:2/20:1)-H | -80 | -10 | -50 | -15 | 120 | 2 |
| 1206 | 723.497 | 307.264 | 9.43 | PA(18:2/20:2)-H | -80 | -10 | -50 | -15 | 120 | 2 |
| 1207 | 721.481 | 305.249 | 10.06 | PA(18:2/20:3)-H | -80 | -10 | -50 | -15 | 120 | 2 |
| 1208 | 719.466 | 303.233 | 9.16 | PA(18:2/20:4)-H | -80 | -10 | -50 | -15 | 120 | 2 |
| 1209 | 717.45 | 301.217 | 9.16 | PA(18:2/20:5)-H | -80 | -10 | -50 | -15 | 120 | 2 |
| 1210 | 747.497 | 331.264 | 9.16 | PA(18:2/22:4)-H | -80 | -10 | -50 | -15 | 120 | 2 |
| 1211 | 745.481 | 329.249 | 10.72 | PA(18:2/22:5)-H | -80 | -10 | -50 | -15 | 120 | 2 |
| 1212 | 743.466 | 327.233 | 10.29 | PA(18:2/22:6)-H | -80 | -10 | -50 | -15 | 120 | 2 |
| 1213 | 701.513 | 253.217 | 10.29 | PA(20:0/16:1)-H | -80 | -10 | -50 | -15 | 120 | 2 |
| 1214 | 729.544 | 281.249 | 10.24 | PA(20:0/18:1)-H | -80 | -10 | -50 | -15 | 120 | 2 |
| 1215 | 727.528 | 279.233 | 10.05 | PA(20:0/18:2)-H | -80 | -10 | -50 | -15 | 120 | 2 |
| 1216 | 725.513 | 277.217 | 10.06 | PA(20:0/18:3)-H | -80 | -10 | -50 | -15 | 120 | 2 |
| 1217 | 757.575 | 309.28 | 10.67 | PA(20:0/20:1)-H | -80 | -10 | -50 | -15 | 120 | 2 |
| 1218 | 755.56 | 307.264 | 9.83 | PA(20:0/20:2)-H | -80 | -10 | -50 | -15 | 120 | 2 |
| 1219 | 753.544 | 305.249 | 9.76 | PA(20:0/20:3)-H | -80 | -10 | -50 | -15 | 120 | 2 |
| 1220 | 751.528 | 303.233 | 9.64 | PA(20:0/20:4)-H | -80 | -10 | -50 | -15 | 120 | 2 |
| 1221 | 749.513 | 301.217 | 9.61 | PA(20:0/20:5)-H | -80 | -10 | -50 | -15 | 120 | 2 |
| 1222 | 779.56 | 331.264 | 9.64 | PA(20:0/22:4)-H | -80 | -10 | -50 | -15 | 120 | 2 |
| 1223 | 777.544 | 329.249 | 9.64 | PA(20:0/22:5)-H | -80 | -10 | -50 | -15 | 120 | 2 |
| 1224 | 775.528 | 327.233 | 9.49 | PA(20:0/22:6)-H | -80 | -10 | -50 | -15 | 120 | 2 |
| S.No. | Q1 | Q3 | Retention time | ID (Internal standards) | DP | EP | CE | CXP | WINDOW (SEC) | DWELL WEGHT |
| 1 | 738.663 | 184.2 | 11.84 | SM (d18:1-18:1(d9)) | 80 | 10 | 43 | 15 | 120 | 2 |
| 2 | 552.5 | 264.3 | 2.5 | Ceramide (17:0) | 80 | 10 | 43 | 15 | 127.4 | 2 |
| 3 | 829.37 | 570.5 | 2.2 | TAG (15:0-18:1(d7)-15:0) | 80 | 10 | 38 | 15 | 180 | 2 |
| 4 | 605.5 | 346.3 | 2.28 | DAG (15:0-18:1(d7)) | 80 | 10 | 25 | 15 | 200 | 2 |
| 5 | 587.409 | 288.296 | 12.54 | LPC (18:1(d7)) | -80 | -10 | -50 | -15 | 120 | 2 |
| 6 | 811.625 | 288.298 | 9.61 | PC (15:0-18:1(d7)) | -80 | -10 | -50 | -15 | 180 | 2 |
| 7 | 485.343 | 288.298 | 13 | LPE (18:1(d7)) | -80 | -10 | -50 | -15 | 150 | 2 |
| 8 | 709.557 | 288.298 | 10.57 | PE (15:0-18:1(d7)) | -80 | -10 | -50 | -15 | 180 | 2 |
| 9 | 740.552 | 288.298 | 6.53 | PG (15:0-18:1(d7)) | -80 | -10 | -50 | -15 | 620.5 | 4.2 |
| 10 | 828.56 | 288.298 | 12.97 | PI (15:0-18:1(d7)) | -80 | -10 | -50 | -15 | 120 | 2 |
| 11 | 753.547 | 288.298 | 9.58 | PS (15:0-18:1(d7)) | -80 | -10 | -50 | -15 | 400 | 2.17 |
| 12 | 666.515 | 288.298 | 11.66 | PA (15:0-18:1(d7)) | -80 | -10 | -50 | -15 | 120 | 2 |

Supplementary- table 2:

| S.No. | Lipid class | Chain length | Un-saturation | Abundance (c.p.s.) |
|---|---|---|---|---|
| 1 | Triglyceride | 42 | 1 | 811367.76 |
| 2 | Triglyceride | 42 | 2 | 820309.50 |
| 3 | Triglyceride | 44 | 0 | 1889691.19 |
| 4 | Triglyceride | 44 | 1 | 1607071.49 |
| 5 | Triglyceride | 44 | 2 | 3545918.46 |
| 6 | Triglyceride | 44 | 3 | 205829.78 |
| 7 | Triglyceride | 46 | 0 | 4098134.77 |
| 8 | Triglyceride | 46 | 1 | 9479326.70 |
| 9 | Triglyceride | 46 | 2 | 11754738.12 |
| 10 | Triglyceride | 46 | 3 | 3280398.83 |
| 11 | Triglyceride | 46 | 4 | 552746.31 |
| 12 | Triglyceride | 47 | 0 | 716157.48 |
| 13 | Triglyceride | 47 | 1 | 382891.78 |
| 14 | Triglyceride | 47 | 2 | 835052.67 |
| 15 | Triglyceride | 48 | 0 | 16502695.61 |
| 16 | Triglyceride | 48 | 1 | 41214794.05 |
| 17 | Triglyceride | 48 | 2 | 40746576.58 |
| 18 | Triglyceride | 48 | 3 | 21115895.16 |
| 19 | Triglyceride | 48 | 4 | 11102440.52 |
| 20 | Triglyceride | 48 | 5 | 2253911.36 |
| 21 | Triglyceride | 49 | 0 | 3083820.03 |
| 22 | Triglyceride | 49 | 1 | 10864412.35 |
| 23 | Triglyceride | 49 | 2 | 8608646.22 |
| 24 | Triglyceride | 49 | 3 | 1924646.16 |
| 25 | Triglyceride | 50 | 0 | 34780705.65 |
| 26 | Triglyceride | 50 | 1 | 204298901.49 |
| 27 | Triglyceride | 50 | 2 | 283851672.53 |
| 28 | Triglyceride | 50 | 3 | 162174064.22 |
| 29 | Triglyceride | 50 | 4 | 25828409.94 |
| 30 | Triglyceride | 50 | 5 | 4744665.99 |
| 31 | Triglyceride | 50 | 6 | 253435.34 |
| 32 | Triglyceride | 51 | 0 | 17138202.27 |
| 33 | Triglyceride | 51 | 1 | 19427195.71 |
| 34 | Triglyceride | 51 | 2 | 30104469.12 |
| 35 | Triglyceride | 51 | 3 | 11865280.03 |
| 36 | Triglyceride | 51 | 4 | 6986149.45 |
| 37 | Triglyceride | 51 | 5 | 1379938.29 |
| 38 | Triglyceride | 52 | 0 | 19914688.18 |
| 39 | Triglyceride | 52 | 1 | 177898566.00 |
| 40 | Triglyceride | 52 | 2 | 822185377.32 |
| 41 | Triglyceride | 52 | 3 | 1138049923.52 |

| | | | | |
|---|---|---|---|---|
| 42 | Triglyceride | 52 | 4 | 641631175.95 |
| 43 | Triglyceride | 52 | 5 | 130213513.11 |
| 44 | Triglyceride | 52 | 6 | 14790578.81 |
| 45 | Triglyceride | 52 | 7 | 1019005.52 |
| 46 | Triglyceride | 52 | 8 | 473614.14 |
| 47 | Triglyceride | 53 | 0 | 1601398.34 |
| 48 | Triglyceride | 53 | 1 | 11053244.27 |
| 49 | Triglyceride | 53 | 2 | 38690484.95 |
| 50 | Triglyceride | 53 | 3 | 29970008.42 |
| 51 | Triglyceride | 53 | 4 | 21446406.53 |
| 52 | Triglyceride | 53 | 5 | 615823.28 |
| 53 | Triglyceride | 53 | 6 | 301631.37 |
| 54 | Triglyceride | 54 | 0 | 5245645.35 |
| 55 | Triglyceride | 54 | 1 | 48208560.39 |
| 56 | Triglyceride | 54 | 2 | 202606946.01 |
| 57 | Triglyceride | 54 | 3 | 439147734.51 |
| 58 | Triglyceride | 54 | 4 | 531644374.43 |
| 59 | Triglyceride | 54 | 5 | 416224602.55 |
| 60 | Triglyceride | 54 | 6 | 206243666.48 |
| 61 | Triglyceride | 54 | 7 | 52378554.06 |
| 62 | Triglyceride | 54 | 8 | 7011866.25 |
| 63 | Triglyceride | 55 | 1 | 3709166.50 |
| 64 | Triglyceride | 55 | 2 | 5703325.59 |
| 65 | Triglyceride | 55 | 3 | 11077926.49 |
| 66 | Triglyceride | 55 | 4 | 13387239.89 |
| 67 | Triglyceride | 55 | 5 | 10748796.70 |
| 68 | Triglyceride | 55 | 7 | 317837.09 |
| 69 | Triglyceride | 56 | 10 | 2225567.09 |
| 70 | Triglyceride | 56 | 1 | 7957579.40 |
| 71 | Triglyceride | 56 | 2 | 27478512.54 |
| 72 | Triglyceride | 56 | 3 | 73732927.41 |
| 73 | Triglyceride | 56 | 4 | 73870722.65 |
| 74 | Triglyceride | 56 | 5 | 67984387.23 |
| 75 | Triglyceride | 56 | 6 | 64120050.95 |
| 76 | Triglyceride | 56 | 7 | 48751487.00 |
| 77 | Triglyceride | 56 | 8 | 22465226.12 |
| 78 | Triglyceride | 56 | 9 | 3635250.13 |
| 79 | Triglyceride | 57 | 10 | 74015.83 |
| 80 | Triglyceride | 57 | 2 | 1371787.89 |
| 81 | Triglyceride | 57 | 3 | 813856.56 |
| 82 | Triglyceride | 58 | 10 | 3763983.55 |
| 83 | Triglyceride | 58 | 2 | 5380361.00 |
| 84 | Triglyceride | 58 | 6 | 5271076.57 |
| 85 | Triglyceride | 58 | 7 | 8277494.56 |
| 86 | Triglyceride | 58 | 8 | 10175817.99 |
| 87 | Triglyceride | 58 | 9 | 8526499.15 |

| | | | | |
|---|---|---|---|---|
| 88 | Triglyceride | 60 | 10 | 436566.91 |
| 89 | Triglyceride | 60 | 11 | 252546.15 |
| 90 | Triglyceride | 60 | 12 | 97536.40 |
| 91 | Phospholipid | 28 | 0 | 288788.26 |
| 92 | Phospholipid | 28 | 2 | 37290.01 |
| 93 | Phospholipid | 30 | 0 | 2843154.68 |
| 94 | Phospholipid | 32 | 0 | 46696177.33 |
| 95 | Phospholipid | 32 | 1 | 12028870.95 |
| 96 | Phospholipid | 32 | 2 | 5493335.46 |
| 97 | Phospholipid | 32 | 3 | 633554.24 |
| 98 | Phospholipid | 34 | 0 | 14921002.91 |
| 99 | Phospholipid | 34 | 1 | 110866574.20 |
| 100 | Phospholipid | 34 | 2 | 292710433.02 |
| 101 | Phospholipid | 34 | 3 | 80774166.70 |
| 102 | Phospholipid | 34 | 4 | 1161214.17 |
| 103 | Phospholipid | 34 | 5 | 314113.89 |
| 104 | Phospholipid | 36 | 0 | 12648874.12 |
| 105 | Phospholipid | 36 | 1 | 110434087.17 |
| 106 | Phospholipid | 36 | 2 | 388545030.09 |
| 107 | Phospholipid | 36 | 3 | 120956037.48 |
| 108 | Phospholipid | 36 | 4 | 95460746.10 |
| 109 | Phospholipid | 36 | 5 | 2805766.69 |
| 110 | Phospholipid | 36 | 6 | 175056.99 |
| 111 | Phospholipid | 38 | 0 | 6505349.38 |
| 112 | Phospholipid | 38 | 1 | 5501593.47 |
| 113 | Phospholipid | 38 | 2 | 21721448.95 |
| 114 | Phospholipid | 38 | 3 | 32184037.44 |
| 115 | Phospholipid | 38 | 4 | 6243053.92 |
| 116 | Phospholipid | 38 | 5 | 29007233.62 |
| 117 | Phospholipid | 38 | 6 | 11955408.01 |
| 118 | Phospholipid | 38 | 7 | 183959.38 |
| 119 | Phospholipid | 40 | 1 | 115393.73 |
| 120 | Phospholipid | 40 | 2 | 251188.51 |
| 121 | Phospholipid | 40 | 3 | 2670761.29 |
| 122 | Phospholipid | 40 | 4 | 21425072.83 |
| 123 | Phospholipid | 40 | 5 | 9084793.88 |
| 124 | Phospholipid | 40 | 6 | 5704598.55 |
| 125 | Phospholipid | 40 | 7 | 1093411.56 |
| 126 | Phospholipid | 40 | 8 | 217216.16 |
| 127 | Phospholipid | 42 | 4 | 1509600.44 |
| 128 | Phospholipid | 42 | 5 | 1473665.06 |
| 129 | Phospholipid | 42 | 6 | 759968.54 |

Supplementary- table 3:

| S.No. | Lipid species | Different types of lipids within class | No. of Isomer | Avg |
|---|---|---|---|---|
| 1 | SM(14:0)+H | _ | _ | 63890834.58 |
| 2 | SM(16:0)+H | _ | _ | 584909027.42 |
| 3 | SM(18:0)+H | _ | _ | 104342785.94 |
| 4 | SM(18:1)+H | _ | _ | 51032868.17 |
| 5 | SM(20:0)+H | _ | _ | 1781226922.79 |
| 6 | SM(20:1)+H | _ | _ | 131394903.33 |
| 7 | SM(22:0)+H | _ | _ | 1468315557.26 |
| 8 | SM(22:1)+H | _ | _ | 569935148.70 |
| 9 | SM(24:0)+H | _ | _ | 309766142.22 |
| 10 | SM(24:1)+H | _ | _ | 609032297.24 |
| 11 | SM(26:0)+H | _ | _ | 31708675.54 |
| 12 | SM(26:1)+H | _ | _ | 3584916.76 |
| 13 | CE(24:0)+H | _ | _ | 1179657.15 |
| 14 | CE(22:6)+H | _ | _ | 3319912.83 |
| 15 | CE(20:0)+H | _ | _ | 11840992.18 |
| 16 | CE(20:1)+H | _ | _ | 4259836.92 |
| 17 | CE(22:5)+H | _ | _ | 260612.73 |
| 18 | CE(16:0)+H | _ | _ | 95546.20 |
| 19 | CE(16:1)+H | _ | _ | 75411.83 |
| 20 | CE(18:0)+H | _ | _ | 1010605.01 |
| 21 | CE(18:1)+H | _ | _ | 370006.07 |
| 22 | CE(18:2)+H | _ | _ | 945879.47 |
| 23 | CE(18:3)+H | _ | _ | 1617531.78 |
| 24 | CE(20:2)+H | _ | _ | 403153.98 |
| 25 | CE(20:3)+H | _ | _ | 135502.96 |
| 26 | CE(20:4)+H | _ | _ | 291818.22 |
| 27 | CE(20:5)+H | _ | _ | 514272.65 |
| 28 | CE(22:0)+H | _ | _ | 169154.80 |
| 29 | CE(22:1)+H | _ | _ | 454230.44 |
| 30 | CE(22:2)+H | _ | _ | 2771122.81 |
| 31 | CE(22:4)+H | _ | _ | 479027.25 |
| 32 | CER(14:0)+H | _ | _ | 70229.43 |
| 33 | CER(16:0)+H | _ | _ | 1630860.46 |
| 34 | CER(18:0)+H | _ | _ | 466982.19 |
| 35 | CER(20:0)+H | _ | _ | 383056.74 |
| 36 | CER(22:0)+H | _ | _ | 2516961.87 |
| 37 | CER(22:1)+H | _ | _ | 2961059.17 |
| 38 | CER(24:0)+H | _ | _ | 8039109.95 |
| 39 | CER(24:1)+H | _ | _ | 7307158.13 |
| 40 | DCER(16:0)+H | _ | _ | 521865.44 |

| | | | | |
|---|---|---|---|---|
| 41 | DCER(18:1)+H | _ | _ | 41203.84 |
| 42 | DCER(20:0)+H | _ | _ | 159437.31 |
| 43 | DCER(22:0)+H | _ | _ | 831693.00 |
| 44 | DCER(22:1)+H | _ | _ | 559917.07 |
| 45 | DCER(24:0)+H | _ | _ | 1532185.17 |
| 46 | DCER(24:1)+H | _ | _ | 901089.36 |
| 47 | DCER(26:0)+H | _ | _ | 67402.93 |
| 48 | HCER(14:0)+H | _ | _ | 56351.98 |
| 49 | HCER(16:0)+H | _ | _ | 5547047.68 |
| 50 | HCER(18:0)+H | _ | _ | 444276.79 |
| 51 | HCER(18:1)+H | _ | _ | 832875.72 |
| 52 | HCER(20:0)+H | _ | _ | 583873.16 |
| 53 | HCER(20:1)+H | _ | _ | 662283.03 |
| 54 | HCER(22:0)+H | _ | _ | 4859249.83 |
| 55 | HCER(22:1)+H | _ | _ | 5465968.81 |
| 56 | HCER(24:0)+H | _ | _ | 8761649.76 |
| 57 | HCER(24:1)+H | _ | _ | 8279999.47 |
| 58 | HCER(26:0)+H | _ | _ | 92801.71 |
| 59 | HCER(26:1)+H | _ | _ | 141229.42 |
| 60 | HCER(d18:0/18:0)+H | _ | _ | 83194.90 |
| 61 | HCER(d18:0/20:0)+H | _ | _ | 12767.05 |
| 62 | HCER(d18:0/22:0)+H | _ | _ | 101364.07 |
| 63 | HCER(d18:0/24:0)+H | _ | _ | 159561.01 |
| 64 | HCER(d18:0/24:1)+H | _ | _ | 230506.85 |
| 65 | HCER(d18:0/26:0)+H | _ | _ | 65382.26 |
| 66 | HCER(d18:0/26:1)+H | _ | _ | 137131.52 |
| 67 | LCER(14:0)+H | _ | _ | 108100.38 |
| 68 | LCER(16:0)+H | _ | _ | 96444.44 |
| 69 | LCER(18:0)+H | _ | _ | 109958.33 |
| 70 | LCER(18:1)+H | _ | _ | 185111.33 |
| 71 | LCER(20:0)+H | _ | _ | 183419.76 |
| 72 | LCER(20:1)+H | _ | _ | 144543.49 |
| 73 | LCER(22:0)+H | _ | _ | 72851.25 |
| 74 | LCER(24:0)+H | _ | _ | 33112.57 |
| 75 | LCER(24:1)+H | _ | _ | 41519.54 |
| 76 | LCER(26:1)+H | _ | _ | 11122.94 |
| 77 | LCER(d18:0/18:0)+H | _ | _ | 62607.70 |
| 78 | LCER(d18:0/20:0)+H | _ | _ | 59326.25 |
| 79 | LCER(d18:0/24:0)+H | _ | _ | 8070.76 |
| 80 | LCER(d18:0/24:1)+H | _ | _ | 28439.07 |
| 81 | TAG(42:1/FA14:0)+NH4 | Type1 | Isomer1 | 157557.22 |
| 82 | TAG(42:1/FA16:1)+NH4 | Type1 | Isomer2 | 95609.58 |
| 83 | TAG(42:1/FA18:1)+NH4 | Type1 | Isomer3 | 558200.96 |
| 84 | TAG(42:2/FA18:2)+NH4 | Type2 | Isomer1 | 820309.50 |
| 85 | TAG(44:0/FA16:0)+NH4 | Type3 | Isomer1 | 1377190.11 |
| 86 | TAG(44:0/FA18:0)+NH4 | Type3 | Isomer2 | 512501.09 |

| | | | | |
|---|---|---|---|---|
| 87 | TAG(44:1/FA14:0)+NH4 | Type4 | Isomer1 | 732510.07 |
| 88 | TAG(44:1/FA16:0)+NH4 | Type4 | Isomer2 | 874561.42 |
| 89 | TAG(44:2/FA14:0)+NH4 | Type5 | Isomer1 | 1089103.60 |
| 90 | TAG(44:2/FA16:0)+NH4 | Type5 | Isomer2 | 582489.33 |
| 91 | TAG(44:2/FA18:1)+NH4 | Type5 | Isomer3 | 302078.34 |
| 92 | TAG(44:2/FA18:2)+NH4 | Type5 | Isomer4 | 1572247.18 |
| 93 | TAG(44:3/FA18:2)+NH4 | Type6 | Isomer1 | 205829.78 |
| 94 | TAG(46:0/FA16:0)+NH4 | Type7 | Isomer1 | 3463252.59 |
| 95 | TAG(46:0/FA18:0)+NH4 | Type7 | Isomer2 | 634882.18 |
| 96 | TAG(46:1/FA14:0)+NH4 | Type8 | Isomer1 | 1738453.90 |
| 97 | TAG(46:1/FA16:0)+NH4 | Type8 | Isomer2 | 3199773.52 |
| 98 | TAG(46:1/FA16:1)+NH4 | Type8 | Isomer3 | 886803.52 |
| 99 | TAG(46:1/FA18:0)+NH4 | Type8 | Isomer4 | 356722.51 |
| 100 | TAG(46:1/FA18:1)+NH4 | Type8 | Isomer5 | 3297573.24 |
| 101 | TAG(46:2/FA14:0)+NH4 | Type9 | Isomer1 | 1403156.97 |
| 102 | TAG(46:2/FA16:0)+NH4 | Type9 | Isomer2 | 3815252.11 |
| 103 | TAG(46:2/FA16:1)+NH4 | Type9 | Isomer3 | 461970.94 |
| 104 | TAG(46:2/FA18:1)+NH4 | Type9 | Isomer4 | 1228216.13 |
| 105 | TAG(46:2/FA18:2)+NH4 | Type9 | Isomer5 | 4846141.98 |
| 106 | TAG(46:3/FA14:0)+NH4 | Type10 | Isomer1 | 191191.00 |
| 107 | TAG(46:3/FA16:0)+NH4 | Type10 | Isomer2 | 601545.46 |
| 108 | TAG(46:3/FA16:1)+NH4 | Type10 | Isomer3 | 288309.69 |
| 109 | TAG(46:3/FA18:1)+NH4 | Type10 | Isomer4 | 689998.16 |
| 110 | TAG(46:3/FA18:2)+NH4 | Type10 | Isomer5 | 858729.17 |
| 111 | TAG(46:3/FA18:3)+NH4 | Type10 | Isomer6 | 650625.36 |
| 112 | TAG(46:4/FA18:2)+NH4 | Type11 | Isomer1 | 552746.31 |
| 113 | TAG(47:0/FA14:0)+NH4 | Type12 | Isomer1 | 298757.57 |
| 114 | TAG(47:0/FA17:0)+NH4 | Type12 | Isomer2 | 417399.91 |
| 115 | TAG(47:1/FA14:0)+NH4 | Type13 | Isomer3 | 382891.78 |
| 116 | TAG(47:2/FA14:0)+NH4 | Type14 | Isomer1 | 329938.05 |
| 117 | TAG(47:2/FA18:2)+NH4 | Type14 | Isomer2 | 505114.62 |
| 118 | TAG(48:0/FA14:0)+NH4 | Type15 | Isomer1 | 1915686.67 |
| 119 | TAG(48:0/FA16:0)+NH4 | Type15 | Isomer2 | 12581375.37 |
| 120 | TAG(48:0/FA18:0)+NH4 | Type15 | Isomer5 | 2005633.58 |
| 121 | TAG(48:1/FA14:0)+NH4 | Type16 | Isomer1 | 9953050.41 |
| 122 | TAG(48:1/FA16:0)+NH4 | Type16 | Isomer2 | 14231599.29 |
| 123 | TAG(48:1/FA16:1)+NH4 | Type16 | Isomer3 | 2063518.29 |
| 124 | TAG(48:1/FA18:0)+NH4 | Type16 | Isomer4 | 1170773.04 |
| 125 | TAG(48:1/FA18:1)+NH4 | Type16 | Isomer5 | 13795853.02 |
| 126 | TAG(48:2/FA14:0)+NH4 | Type17 | Isomer1 | 8498210.42 |
| 127 | TAG(48:2/FA16:0)+NH4 | Type17 | Isomer2 | 9166218.98 |
| 128 | TAG(48:2/FA16:1)+NH4 | Type17 | Isomer3 | 2597917.60 |
| 129 | TAG(48:2/FA18:0)+NH4 | Type17 | Isomer4 | 1500641.87 |
| 130 | TAG(48:2/FA18:1)+NH4 | Type17 | Isomer5 | 7761460.51 |
| 131 | TAG(48:2/FA18:2)+NH4 | Type17 | Isomer6 | 11222127.20 |
| 132 | TAG(48:3/FA14:0)+NH4 | Type18 | Isomer1 | 1979621.87 |

| | | | | |
|---|---|---|---|---|
| 133 | TAG(48:3/FA16:0)+NH4 | Type18 | Isomer2 | 1561641.30 |
| 134 | TAG(48:3/FA16:1)+NH4 | Type18 | Isomer3 | 1358739.38 |
| 135 | TAG(48:3/FA18:1)+NH4 | Type18 | Isomer4 | 6400075.38 |
| 136 | TAG(48:3/FA18:2)+NH4 | Type18 | Isomer5 | 8394103.70 |
| 137 | TAG(48:3/FA18:3)+NH4 | Type18 | Isomer6 | 1421713.54 |
| 138 | TAG(48:4/FA14:0)+NH4 | Type19 | Isomer1 | 153226.68 |
| 139 | TAG(48:4/FA16:0)+NH4 | Type19 | Isomer2 | 197192.39 |
| 140 | TAG(48:4/FA16:1)+NH4 | Type19 | Isomer3 | 205685.70 |
| 141 | TAG(48:4/FA18:1)+NH4 | Type19 | Isomer4 | 856391.67 |
| 142 | TAG(48:4/FA18:2)+NH4 | Type19 | Isomer5 | 8313773.38 |
| 143 | TAG(48:4/FA18:3)+NH4 | Type19 | Isomer6 | 1240673.95 |
| 144 | TAG(48:4/FA20:4)+NH4 | Type19 | Isomer7 | 135496.74 |
| 145 | TAG(48:5/FA18:2)+NH4 | Type20 | Isomer1 | 1011148.80 |
| 146 | TAG(48:5/FA18:3)+NH4 | Type20 | Isomer2 | 1242762.55 |
| 147 | TAG(49:0/FA16:0)+NH4 | Type21 | Isomer1 | 2347030.79 |
| 148 | TAG(49:0/FA18:0)+NH4 | Type21 | Isomer2 | 736789.24 |
| 149 | TAG(49:1/FA14:0)+NH4 | Type22 | Isomer1 | 650293.97 |
| 150 | TAG(49:1/FA16:0)+NH4 | Type22 | Isomer2 | 4358995.70 |
| 151 | TAG(49:1/FA16:1)+NH4 | Type22 | Isomer3 | 459407.30 |
| 152 | TAG(49:1/FA17:0)+NH4 | Type22 | Isomer4 | 858200.32 |
| 153 | TAG(49:1/FA18:1)+NH4 | Type22 | Isomer5 | 4537515.07 |
| 154 | TAG(49:2/FA14:0)+NH4 | Type23 | Isomer1 | 592648.44 |
| 155 | TAG(49:2/FA16:0)+NH4 | Type23 | Isomer2 | 2780632.56 |
| 156 | TAG(49:2/FA17:0)+NH4 | Type23 | Isomer3 | 357899.47 |
| 157 | TAG(49:2/FA18:1)+NH4 | Type23 | Isomer4 | 1503130.14 |
| 158 | TAG(49:2/FA18:2)+NH4 | Type23 | Isomer5 | 3374335.61 |
| 159 | TAG(49:3/FA16:0)+NH4 | Type24 | Isomer1 | 463337.45 |
| 160 | TAG(49:3/FA18:2)+NH4 | Type24 | Isomer2 | 1113331.21 |
| 161 | TAG(49:3/FA18:3)+NH4 | Type24 | Isomer3 | 347977.50 |
| 162 | TAG(50:0/FA14:0)+NH4 | Type25 | Isomer1 | 746360.76 |
| 163 | TAG(50:0/FA16:0)+NH4 | Type25 | Isomer2 | 23364380.78 |
| 164 | TAG(50:0/FA18:0)+NH4 | Type25 | Isomer3 | 10669964.11 |
| 165 | TAG(50:1/FA14:0)+NH4 | Type26 | Isomer1 | 4933652.61 |
| 166 | TAG(50:1/FA16:0)+NH4 | Type26 | Isomer2 | 112434722.14 |
| 167 | TAG(50:1/FA16:1)+NH4 | Type26 | Isomer3 | 3243258.67 |
| 168 | TAG(50:1/FA18:0)+NH4 | Type26 | Isomer4 | 6443483.99 |
| 169 | TAG(50:1/FA18:1)+NH4 | Type26 | Isomer5 | 76782161.69 |
| 170 | TAG(50:1/FA20:1)+NH4 | Type26 | Isomer6 | 461622.39 |
| 171 | TAG(50:2/FA14:0)+NH4 | Type27 | Isomer1 | 22061171.55 |
| 172 | TAG(50:2/FA16:0)+NH4 | Type27 | Isomer2 | 110500381.62 |
| 173 | TAG(50:2/FA16:1)+NH4 | Type27 | Isomer3 | 26354408.84 |
| 174 | TAG(50:2/FA18:0)+NH4 | Type27 | Isomer4 | 3656802.79 |
| 175 | TAG(50:2/FA18:1)+NH4 | Type27 | Isomer5 | 62231593.51 |
| 176 | TAG(50:2/FA18:2)+NH4 | Type27 | Isomer6 | 58782250.85 |
| 177 | TAG(50:2/FA20:2)+NH4 | Type27 | Isomer7 | 265063.38 |
| 178 | TAG(50:3/FA14:0)+NH4 | Type28 | Isomer1 | 27274238.65 |

| | | | | |
|---|---|---|---|---|
| 179 | TAG(50:3/FA16:0)+NH4 | Type28 | Isomer2 | 29274318.15 |
| 180 | TAG(50:3/FA16:1)+NH4 | Type28 | Isomer3 | 20736479.56 |
| 181 | TAG(50:3/FA18:0)+NH4 | Type28 | Isomer4 | 630992.51 |
| 182 | TAG(50:3/FA18:1)+NH4 | Type28 | Isomer5 | 28910184.87 |
| 183 | TAG(50:3/FA18:2)+NH4 | Type28 | Isomer6 | 48156539.76 |
| 184 | TAG(50:3/FA18:3)+NH4 | Type28 | Isomer7 | 6742100.55 |
| 185 | TAG(50:3/FA20:3)+NH4 | Type28 | Isomer8 | 449210.17 |
| 186 | TAG(50:4/FA14:0)+NH4 | Type29 | Isomer1 | 7033829.84 |
| 187 | TAG(50:4/FA16:0)+NH4 | Type29 | Isomer2 | 1175362.44 |
| 188 | TAG(50:4/FA16:1)+NH4 | Type29 | Isomer3 | 1600693.05 |
| 189 | TAG(50:4/FA18:1)+NH4 | Type29 | Isomer4 | 1619102.53 |
| 190 | TAG(50:4/FA18:2)+NH4 | Type29 | Isomer5 | 11501011.56 |
| 191 | TAG(50:4/FA18:3)+NH4 | Type29 | Isomer6 | 2395295.76 |
| 192 | TAG(50:4/FA20:3)+NH4 | Type29 | Isomer7 | 127482.49 |
| 193 | TAG(50:4/FA20:4)+NH4 | Type29 | Isomer8 | 375632.26 |
| 194 | TAG(50:5/FA14:0)+NH4 | Type30 | Isomer1 | 1131301.28 |
| 195 | TAG(50:5/FA16:0)+NH4 | Type30 | Isomer2 | 161173.90 |
| 196 | TAG(50:5/FA16:1)+NH4 | Type30 | Isomer3 | 190585.68 |
| 197 | TAG(50:5/FA18:1)+NH4 | Type30 | Isomer4 | 184904.32 |
| 198 | TAG(50:5/FA18:2)+NH4 | Type30 | Isomer5 | 1267873.54 |
| 199 | TAG(50:5/FA18:3)+NH4 | Type30 | Isomer6 | 1504474.35 |
| 200 | TAG(50:5/FA20:4)+NH4 | Type30 | Isomer7 | 196181.01 |
| 201 | TAG(50:5/FA20:5)+NH4 | Type30 | Isomer8 | 108171.91 |
| 202 | TAG(50:6/FA20:4)+NH4 | Type31 | Isomer1 | 253435.34 |
| 203 | TAG(51:0/FA16:0)+NH4 | Type31 | Isomer2 | 1903564.64 |
| 204 | TAG(51:0/FA17:0)+NH4 | Type31 | Isomer3 | 14064793.16 |
| 205 | TAG(51:0/FA18:0)+NH4 | Type32 | Isomer1 | 1169844.47 |
| 206 | TAG(51:1/FA16:0)+NH4 | Type33 | Isomer1 | 6442401.11 |
| 207 | TAG(51:1/FA17:0)+NH4 | Type33 | Isomer2 | 4695093.90 |
| 208 | TAG(51:1/FA18:0)+NH4 | Type33 | Isomer3 | 1466866.20 |
| 209 | TAG(51:1/FA18:1)+NH4 | Type33 | Isomer4 | 6822834.50 |
| 210 | TAG(51:2/FA16:0)+NH4 | Type34 | Isomer1 | 7193005.66 |
| 211 | TAG(51:2/FA17:0)+NH4 | Type34 | Isomer2 | 3901675.36 |
| 212 | TAG(51:2/FA18:1)+NH4 | Type34 | Isomer3 | 14412628.21 |
| 213 | TAG(51:2/FA18:2)+NH4 | Type34 | Isomer4 | 4597159.89 |
| 214 | TAG(51:3/FA16:1)+NH4 | Type35 | Isomer1 | 1035212.38 |
| 215 | TAG(51:3/FA17:0)+NH4 | Type35 | Isomer2 | 751949.20 |
| 216 | TAG(51:3/FA18:2)+NH4 | Type35 | Isomer3 | 9513400.44 |
| 217 | TAG(51:3/FA18:3)+NH4 | Type35 | Isomer4 | 564718.01 |
| 218 | TAG(51:4/FA16:1)+NH4 | Type36 | Isomer1 | 319184.09 |
| 219 | TAG(51:4/FA18:2)+NH4 | Type36 | Isomer2 | 5259749.61 |
| 220 | TAG(51:4/FA18:3)+NH4 | Type36 | Isomer3 | 1168947.74 |
| 221 | TAG(51:4/FA20:4)+NH4 | Type36 | Isomer4 | 238268.01 |
| 222 | TAG(51:5/FA18:2)+NH4 | Type37 | Isomer1 | 771541.99 |
| 223 | TAG(51:5/FA18:3)+NH4 | Type37 | Isomer2 | 608396.31 |
| 224 | TAG(52:0/FA16:0)+NH4 | Type38 | Isomer1 | 7680744.25 |

| | | | | |
|---|---|---|---|---|
| 225 | TAG(52:0/FA18:0)+NH4 | Type38 | Isomer2 | 10665622.89 |
| 226 | TAG(52:0/FA20:0)+NH4 | Type38 | Isomer3 | 1568321.04 |
| 227 | TAG(52:1/FA16:0)+NH4 | Type39 | Isomer1 | 55009828.59 |
| 228 | TAG(52:1/FA16:1)+NH4 | Type39 | Isomer2 | 633099.20 |
| 229 | TAG(52:1/FA18:0)+NH4 | Type39 | Isomer3 | 44678116.23 |
| 230 | TAG(52:1/FA18:1)+NH4 | Type39 | Isomer4 | 73095797.11 |
| 231 | TAG(52:1/FA20:0)+NH4 | Type39 | Isomer5 | 2501078.32 |
| 232 | TAG(52:1/FA20:1)+NH4 | Type39 | Isomer6 | 1980646.55 |
| 233 | TAG(52:2/FA14:0)+NH4 | Type40 | Isomer1 | 1891593.57 |
| 234 | TAG(52:2/FA16:0)+NH4 | Type40 | Isomer2 | 282384932.38 |
| 235 | TAG(52:2/FA16:1)+NH4 | Type40 | Isomer3 | 6347653.16 |
| 236 | TAG(52:2/FA18:0)+NH4 | Type40 | Isomer4 | 37657707.66 |
| 237 | TAG(52:2/FA18:1)+NH4 | Type40 | Isomer5 | 431088516.46 |
| 238 | TAG(52:2/FA18:2)+NH4 | Type40 | Isomer6 | 56452984.62 |
| 239 | TAG(52:2/FA20:0)+NH4 | Type40 | Isomer7 | 1654078.54 |
| 240 | TAG(52:2/FA20:1)+NH4 | Type40 | Isomer8 | 3552055.26 |
| 241 | TAG(52:2/FA20:2)+NH4 | Type40 | Isomer9 | 1155855.68 |
| 242 | TAG(52:3/FA14:0)+NH4 | Type41 | Isomer1 | 1844225.36 |
| 243 | TAG(52:3/FA16:0)+NH4 | Type41 | Isomer2 | 369600055.44 |
| 244 | TAG(52:3/FA16:1)+NH4 | Type41 | Isomer3 | 32812389.39 |
| 245 | TAG(52:3/FA18:0)+NH4 | Type41 | Isomer4 | 5715274.36 |
| 246 | TAG(52:3/FA18:1)+NH4 | Type41 | Isomer5 | 360528558.86 |
| 247 | TAG(52:3/FA18:2)+NH4 | Type41 | Isomer6 | 354094037.97 |
| 248 | TAG(52:3/FA18:3)+NH4 | Type41 | Isomer7 | 6451634.63 |
| 249 | TAG(52:3/FA20:0)+NH4 | Type41 | Isomer8 | 1059705.79 |
| 250 | TAG(52:3/FA20:1)+NH4 | Type41 | Isomer9 | 2037511.89 |
| 251 | TAG(52:3/FA20:2)+NH4 | Type41 | Isomer10 | 1630749.94 |
| 252 | TAG(52:3/FA20:3)+NH4 | Type41 | Isomer11 | 2275779.88 |
| 253 | TAG(52:4/FA14:0)+NH4 | Type42 | Isomer1 | 1072544.91 |
| 254 | TAG(52:4/FA16:0)+NH4 | Type42 | Isomer2 | 189023507.27 |
| 255 | TAG(52:4/FA16:1)+NH4 | Type42 | Isomer3 | 36981338.82 |
| 256 | TAG(52:4/FA18:0)+NH4 | Type42 | Isomer4 | 653333.32 |
| 257 | TAG(52:4/FA18:1)+NH4 | Type42 | Isomer5 | 58058124.99 |
| 258 | TAG(52:4/FA18:2)+NH4 | Type42 | Isomer6 | 299558642.17 |
| 259 | TAG(52:4/FA18:3)+NH4 | Type42 | Isomer7 | 46350465.35 |
| 260 | TAG(52:4/FA20:0)+NH4 | Type42 | Isomer8 | 1542697.82 |
| 261 | TAG(52:4/FA20:2)+NH4 | Type42 | Isomer9 | 759076.51 |
| 262 | TAG(52:4/FA20:3)+NH4 | Type42 | Isomer10 | 2527134.72 |
| 263 | TAG(52:4/FA20:4)+NH4 | Type42 | Isomer11 | 4756816.46 |
| 264 | TAG(52:4/FA22:4)+NH4 | Type42 | Isomer12 | 347493.63 |
| 265 | TAG(52:5/FA14:0)+NH4 | Type43 | Isomer1 | 748373.79 |
| 266 | TAG(52:5/FA16:0)+NH4 | Type43 | Isomer2 | 24697272.04 |
| 267 | TAG(52:5/FA16:1)+NH4 | Type43 | Isomer3 | 15568198.66 |
| 268 | TAG(52:5/FA18:1)+NH4 | Type43 | Isomer4 | 5235667.25 |
| 269 | TAG(52:5/FA18:2)+NH4 | Type43 | Isomer5 | 41773153.43 |
| 270 | TAG(52:5/FA18:3)+NH4 | Type43 | Isomer6 | 36550474.76 |

| | | | | |
|---|---|---|---|---|
| 271 | TAG(52:5/FA20:3)+NH4 | Type43 | Isomer7 | 837471.44 |
| 272 | TAG(52:5/FA20:4)+NH4 | Type43 | Isomer8 | 3232365.48 |
| 273 | TAG(52:5/FA20:5)+NH4 | Type43 | Isomer9 | 714580.44 |
| 274 | TAG(52:5/FA22:5)+NH4 | Type43 | Isomer10 | 855955.84 |
| 275 | TAG(52:6/FA14:0)+NH4 | Type44 | Isomer1 | 331226.75 |
| 276 | TAG(52:6/FA16:0)+NH4 | Type44 | Isomer2 | 1714043.40 |
| 277 | TAG(52:6/FA16:1)+NH4 | Type44 | Isomer3 | 1850691.59 |
| 278 | TAG(52:6/FA18:1)+NH4 | Type44 | Isomer4 | 490382.85 |
| 279 | TAG(52:6/FA18:2)+NH4 | Type44 | Isomer5 | 3498086.44 |
| 280 | TAG(52:6/FA18:3)+NH4 | Type44 | Isomer6 | 4726658.30 |
| 281 | TAG(52:6/FA20:4)+NH4 | Type44 | Isomer7 | 1232899.56 |
| 282 | TAG(52:6/FA20:5)+NH4 | Type44 | Isomer8 | 554639.61 |
| 283 | TAG(52:6/FA22:6)+NH4 | Type44 | Isomer9 | 391950.30 |
| 284 | TAG(52:7/FA18:1)+NH4 | Type45 | Isomer1 | 529862.34 |
| 285 | TAG(52:7/FA20:5)+NH4 | Type45 | Isomer2 | 194143.40 |
| 286 | TAG(52:7/FA22:6)+NH4 | Type45 | Isomer3 | 294999.78 |
| 287 | TAG(52:8/FA18:2)+NH4 | Type46 | Isomer1 | 473614.14 |
| 288 | TAG(53:0/FA16:0)+NH4 | Type47 | Isomer1 | 1601398.34 |
| 289 | TAG(53:1/FA16:0)+NH4 | Type48 | Isomer1 | 2676999.89 |
| 290 | TAG(53:1/FA17:0)+NH4 | Type48 | Isomer2 | 1753238.82 |
| 291 | TAG(53:1/FA18:0)+NH4 | Type48 | Isomer3 | 2239859.55 |
| 292 | TAG(53:1/FA18:1)+NH4 | Type48 | Isomer4 | 4383146.00 |
| 293 | TAG(53:2/FA16:0)+NH4 | Type49 | Isomer1 | 7167911.21 |
| 294 | TAG(53:2/FA17:0)+NH4 | Type49 | Isomer2 | 9150515.98 |
| 295 | TAG(53:2/FA18:1)+NH4 | Type49 | Isomer3 | 19139899.54 |
| 296 | TAG(53:2/FA18:2)+NH4 | Type49 | Isomer4 | 3232158.23 |
| 297 | TAG(53:3/FA16:0)+NH4 | Type50 | Isomer1 | 5954138.11 |
| 298 | TAG(53:3/FA17:0)+NH4 | Type50 | Isomer2 | 10816080.97 |
| 299 | TAG(53:3/FA18:2)+NH4 | Type50 | Isomer3 | 13199789.34 |
| 300 | TAG(53:4/FA16:0)+NH4 | Type51 | Isomer1 | 2823853.23 |
| 301 | TAG(53:4/FA17:0)+NH4 | Type51 | Isomer2 | 4963796.97 |
| 302 | TAG(53:4/FA18:2)+NH4 | Type51 | Isomer3 | 11821345.00 |
| 303 | TAG(53:4/FA18:3)+NH4 | Type51 | Isomer4 | 1475021.53 |
| 304 | TAG(53:4/FA20:4)+NH4 | Type51 | Isomer5 | 362389.79 |
| 305 | TAG(53:5/FA20:4)+NH4 | Type52 | Isomer1 | 615823.28 |
| 306 | TAG(53:6/FA20:4)+NH4 | Type53 | Isomer1 | 301631.37 |
| 307 | TAG(54:0/FA16:0)+NH4 | Type54 | Isomer1 | 2243867.52 |
| 308 | TAG(54:0/FA18:0)+NH4 | Type54 | Isomer2 | 3001777.83 |
| 309 | TAG(54:1/FA16:0)+NH4 | Type55 | Isomer1 | 5075234.22 |
| 310 | TAG(54:1/FA18:0)+NH4 | Type55 | Isomer2 | 16058925.82 |
| 311 | TAG(54:1/FA18:1)+NH4 | Type55 | Isomer3 | 13862352.04 |
| 312 | TAG(54:1/FA20:0)+NH4 | Type55 | Isomer4 | 10727283.93 |
| 313 | TAG(54:1/FA20:1)+NH4 | Type55 | Isomer5 | 2484764.37 |
| 314 | TAG(54:2/FA16:0)+NH4 | Type56 | Isomer1 | 19792483.28 |
| 315 | TAG(54:2/FA18:0)+NH4 | Type56 | Isomer2 | 52194521.52 |
| 316 | TAG(54:2/FA18:1)+NH4 | Type56 | Isomer3 | 94050379.69 |

| | | | | |
|---|---|---|---|---|
| 317 | TAG(54:2/FA18:2)+NH4 | Type56 | Isomer4 | 10967433.41 |
| 318 | TAG(54:2/FA20:0)+NH4 | Type56 | Isomer5 | 8374539.73 |
| 319 | TAG(54:2/FA20:1)+NH4 | Type56 | Isomer6 | 16264944.39 |
| 320 | TAG(54:2/FA20:2)+NH4 | Type56 | Isomer7 | 962643.98 |
| 321 | TAG(54:3/FA16:0)+NH4 | Type57 | Isomer1 | 20563570.32 |
| 322 | TAG(54:3/FA16:1)+NH4 | Type57 | Isomer2 | 3964045.85 |
| 323 | TAG(54:3/FA18:0)+NH4 | Type57 | Isomer3 | 63728737.17 |
| 324 | TAG(54:3/FA18:1)+NH4 | Type57 | Isomer4 | 253205250.74 |
| 325 | TAG(54:3/FA18:2)+NH4 | Type57 | Isomer5 | 70392790.58 |
| 326 | TAG(54:3/FA18:3)+NH4 | Type57 | Isomer6 | 1136792.41 |
| 327 | TAG(54:3/FA20:1)+NH4 | Type57 | Isomer7 | 16138748.86 |
| 328 | TAG(54:3/FA20:2)+NH4 | Type57 | Isomer8 | 8438372.99 |
| 329 | TAG(54:3/FA20:3)+NH4 | Type57 | Isomer9 | 1579425.59 |
| 330 | TAG(54:4/FA16:0)+NH4 | Type58 | Isomer1 | 15319341.93 |
| 331 | TAG(54:4/FA16:1)+NH4 | Type58 | Isomer2 | 3089381.81 |
| 332 | TAG(54:4/FA18:0)+NH4 | Type58 | Isomer3 | 37847940.89 |
| 333 | TAG(54:4/FA18:1)+NH4 | Type58 | Isomer4 | 254541012.34 |
| 334 | TAG(54:4/FA18:2)+NH4 | Type58 | Isomer5 | 182769437.49 |
| 335 | TAG(54:4/FA18:3)+NH4 | Type58 | Isomer6 | 9964669.23 |
| 336 | TAG(54:4/FA20:1)+NH4 | Type58 | Isomer7 | 3196018.44 |
| 337 | TAG(54:4/FA20:2)+NH4 | Type58 | Isomer8 | 7391450.40 |
| 338 | TAG(54:4/FA20:3)+NH4 | Type58 | Isomer9 | 12875708.29 |
| 339 | TAG(54:4/FA20:4)+NH4 | Type58 | Isomer10 | 3801278.17 |
| 340 | TAG(54:4/FA22:4)+NH4 | Type58 | Isomer11 | 848135.42 |
| 341 | TAG(54:5/FA16:0)+NH4 | Type59 | Isomer1 | 12377957.17 |
| 342 | TAG(54:5/FA16:1)+NH4 | Type59 | Isomer2 | 1909972.32 |
| 343 | TAG(54:5/FA18:0)+NH4 | Type59 | Isomer3 | 6028134.86 |
| 344 | TAG(54:5/FA18:1)+NH4 | Type59 | Isomer4 | 126624241.98 |
| 345 | TAG(54:5/FA18:2)+NH4 | Type59 | Isomer5 | 200328877.22 |
| 346 | TAG(54:5/FA18:3)+NH4 | Type59 | Isomer6 | 32716182.31 |
| 347 | TAG(54:5/FA20:2)+NH4 | Type59 | Isomer7 | 1387520.23 |
| 348 | TAG(54:5/FA20:3)+NH4 | Type59 | Isomer8 | 9842251.64 |
| 349 | TAG(54:5/FA20:4)+NH4 | Type59 | Isomer9 | 22294920.41 |
| 350 | TAG(54:5/FA20:5)+NH4 | Type59 | Isomer10 | 492256.59 |
| 351 | TAG(54:5/FA22:4)+NH4 | Type59 | Isomer11 | 770545.53 |
| 352 | TAG(54:5/FA22:5)+NH4 | Type59 | Isomer12 | 1451742.29 |
| 353 | TAG(54:6/FA16:0)+NH4 | Type60 | Isomer1 | 5254307.63 |
| 354 | TAG(54:6/FA16:1)+NH4 | Type60 | Isomer2 | 1420670.75 |
| 355 | TAG(54:6/FA18:1)+NH4 | Type60 | Isomer3 | 25139210.98 |
| 356 | TAG(54:6/FA18:2)+NH4 | Type60 | Isomer4 | 112559087.12 |
| 357 | TAG(54:6/FA18:3)+NH4 | Type60 | Isomer5 | 34755784.09 |
| 358 | TAG(54:6/FA20:3)+NH4 | Type60 | Isomer6 | 1464885.81 |
| 359 | TAG(54:6/FA20:4)+NH4 | Type60 | Isomer7 | 19324093.86 |
| 360 | TAG(54:6/FA20:5)+NH4 | Type60 | Isomer8 | 3449250.35 |
| 361 | TAG(54:6/FA22:5)+NH4 | Type60 | Isomer9 | 1319584.26 |
| 362 | TAG(54:6/FA22:6)+NH4 | Type60 | Isomer10 | 1556791.63 |

| | | | | |
|---|---|---|---|---|
| 363 | TAG(54:7/FA16:1)+NH4 | Type61 | Isomer1 | 502849.36 |
| 364 | TAG(54:7/FA18:1)+NH4 | Type61 | Isomer2 | 1974305.33 |
| 365 | TAG(54:7/FA18:2)+NH4 | Type61 | Isomer3 | 24267734.89 |
| 366 | TAG(54:7/FA18:3)+NH4 | Type61 | Isomer4 | 18386601.33 |
| 367 | TAG(54:7/FA20:4)+NH4 | Type61 | Isomer5 | 2349780.35 |
| 368 | TAG(54:7/FA20:5)+NH4 | Type61 | Isomer6 | 2771002.83 |
| 369 | TAG(54:7/FA22:5)+NH4 | Type61 | Isomer7 | 504029.40 |
| 370 | TAG(54:7/FA22:6)+NH4 | Type61 | Isomer8 | 1622250.58 |
| 371 | TAG(54:8/FA18:2)+NH4 | Type62 | Isomer1 | 1810753.86 |
| 372 | TAG(54:8/FA18:3)+NH4 | Type62 | Isomer2 | 3818229.45 |
| 373 | TAG(54:8/FA20:4)+NH4 | Type62 | Isomer3 | 226508.08 |
| 374 | TAG(54:8/FA20:5)+NH4 | Type62 | Isomer4 | 438886.60 |
| 375 | TAG(54:8/FA22:6)+NH4 | Type62 | Isomer5 | 717488.27 |
| 376 | TAG(55:1/FA16:0)+NH4 | Type63 | Isomer1 | 1851035.05 |
| 377 | TAG(55:1/FA18:1)+NH4 | Type63 | Isomer2 | 1858131.44 |
| 378 | TAG(55:2/FA18:1)+NH4 | Type64 | Isomer1 | 4389799.75 |
| 379 | TAG(55:2/FA18:2)+NH4 | Type64 | Isomer2 | 1313525.84 |
| 380 | TAG(55:3/FA18:1)+NH4 | Type65 | Isomer1 | 8034995.54 |
| 381 | TAG(55:3/FA18:2)+NH4 | Type65 | Isomer2 | 3042930.95 |
| 382 | TAG(55:4/FA18:1)+NH4 | Type66 | Isomer1 | 7919436.59 |
| 383 | TAG(55:4/FA18:2)+NH4 | Type66 | Isomer2 | 5467803.30 |
| 384 | TAG(55:5/FA18:1)+NH4 | Type67 | Isomer1 | 4135061.05 |
| 385 | TAG(55:5/FA18:2)+NH4 | Type67 | Isomer2 | 5919875.47 |
| 386 | TAG(55:5/FA20:4)+NH4 | Type67 | Isomer3 | 693860.17 |
| 387 | TAG(55:7/FA22:6)+NH4 | Type68 | Isomer1 | 317837.09 |
| 388 | TAG(56:10/FA18:2)+NH4 | Type69 | Isomer1 | 2225567.09 |
| 389 | TAG(56:1/FA16:0)+NH4 | Type70 | Isomer1 | 3294493.58 |
| 390 | TAG(56:1/FA18:1)+NH4 | Type70 | Isomer2 | 4663085.83 |
| 391 | TAG(56:2/FA16:0)+NH4 | Type71 | Isomer1 | 3676291.18 |
| 392 | TAG(56:2/FA18:0)+NH4 | Type71 | Isomer2 | 5399636.77 |
| 393 | TAG(56:2/FA20:0)+NH4 | Type71 | Isomer3 | 12796023.83 |
| 394 | TAG(56:2/FA20:1)+NH4 | Type71 | Isomer4 | 5606560.75 |
| 395 | TAG(56:3/FA18:0)+NH4 | Type72 | Isomer1 | 4465332.26 |
| 396 | TAG(56:3/FA18:1)+NH4 | Type72 | Isomer2 | 29393605.63 |
| 397 | TAG(56:3/FA18:2)+NH4 | Type72 | Isomer3 | 6853134.08 |
| 398 | TAG(56:3/FA20:0)+NH4 | Type72 | Isomer4 | 11587289.52 |
| 399 | TAG(56:3/FA20:1)+NH4 | Type72 | Isomer5 | 20051065.15 |
| 400 | TAG(56:3/FA20:2)+NH4 | Type72 | Isomer6 | 1382500.77 |
| 401 | TAG(56:4/FA16:0)+NH4 | Type73 | Isomer1 | 1742408.09 |
| 402 | TAG(56:4/FA18:0)+NH4 | Type73 | Isomer2 | 2732435.38 |
| 403 | TAG(56:4/FA18:1)+NH4 | Type73 | Isomer3 | 22744446.94 |
| 404 | TAG(56:4/FA18:2)+NH4 | Type73 | Isomer4 | 19288270.51 |
| 405 | TAG(56:4/FA20:1)+NH4 | Type73 | Isomer5 | 19170757.95 |
| 406 | TAG(56:4/FA20:2)+NH4 | Type73 | Isomer6 | 4790008.82 |
| 407 | TAG(56:4/FA20:3)+NH4 | Type73 | Isomer7 | 1930127.56 |
| 408 | TAG(56:4/FA20:4)+NH4 | Type73 | Isomer8 | 1010104.20 |

| | | | | |
|---|---|---|---|---|
| 409 | TAG(56:4/FA22:4)+NH4 | Type73 | Isomer9 | 462163.22 |
| 410 | TAG(56:5/FA16:0)+NH4 | Type74 | Isomer1 | 4007651.71 |
| 411 | TAG(56:5/FA18:0)+NH4 | Type74 | Isomer2 | 3122692.39 |
| 412 | TAG(56:5/FA18:1)+NH4 | Type74 | Isomer3 | 12451740.31 |
| 413 | TAG(56:5/FA18:2)+NH4 | Type74 | Isomer4 | 16055447.38 |
| 414 | TAG(56:5/FA20:1)+NH4 | Type74 | Isomer5 | 9267159.66 |
| 415 | TAG(56:5/FA20:2)+NH4 | Type74 | Isomer6 | 4791389.98 |
| 416 | TAG(56:5/FA20:3)+NH4 | Type74 | Isomer7 | 5748530.56 |
| 417 | TAG(56:5/FA20:4)+NH4 | Type74 | Isomer8 | 7730240.69 |
| 418 | TAG(56:5/FA22:4)+NH4 | Type74 | Isomer9 | 3878227.96 |
| 419 | TAG(56:5/FA22:5)+NH4 | Type74 | Isomer10 | 931306.58 |
| 420 | TAG(56:6/FA16:0)+NH4 | Type75 | Isomer1 | 5508424.01 |
| 421 | TAG(56:6/FA18:0)+NH4 | Type75 | Isomer2 | 1813386.26 |
| 422 | TAG(56:6/FA18:1)+NH4 | Type75 | Isomer3 | 9591028.00 |
| 423 | TAG(56:6/FA18:2)+NH4 | Type75 | Isomer4 | 9347796.01 |
| 424 | TAG(56:6/FA18:3)+NH4 | Type75 | Isomer5 | 2261436.34 |
| 425 | TAG(56:6/FA20:2)+NH4 | Type75 | Isomer6 | 2428012.07 |
| 426 | TAG(56:6/FA20:3)+NH4 | Type75 | Isomer7 | 4994507.79 |
| 427 | TAG(56:6/FA20:4)+NH4 | Type75 | Isomer8 | 15482814.59 |
| 428 | TAG(56:6/FA20:5)+NH4 | Type75 | Isomer9 | 1017463.46 |
| 429 | TAG(56:6/FA22:4)+NH4 | Type75 | Isomer10 | 3344609.35 |
| 430 | TAG(56:6/FA22:5)+NH4 | Type75 | Isomer11 | 7279887.11 |
| 431 | TAG(56:6/FA22:6)+NH4 | Type75 | Isomer12 | 1050685.96 |
| 432 | TAG(56:7/FA16:0)+NH4 | Type76 | Isomer1 | 4685195.57 |
| 433 | TAG(56:7/FA16:1)+NH4 | Type76 | Isomer2 | 366025.73 |
| 434 | TAG(56:7/FA18:0)+NH4 | Type76 | Isomer3 | 614004.58 |
| 435 | TAG(56:7/FA18:1)+NH4 | Type76 | Isomer4 | 3890952.61 |
| 436 | TAG(56:7/FA18:2)+NH4 | Type76 | Isomer5 | 6719104.32 |
| 437 | TAG(56:7/FA18:3)+NH4 | Type76 | Isomer6 | 1477183.49 |
| 438 | TAG(56:7/FA20:3)+NH4 | Type76 | Isomer7 | 2202123.38 |
| 439 | TAG(56:7/FA20:4)+NH4 | Type76 | Isomer8 | 12034256.00 |
| 440 | TAG(56:7/FA20:5)+NH4 | Type76 | Isomer9 | 2375794.00 |
| 441 | TAG(56:7/FA22:4)+NH4 | Type76 | Isomer10 | 376986.75 |
| 442 | TAG(56:7/FA22:5)+NH4 | Type76 | Isomer11 | 5893153.66 |
| 443 | TAG(56:7/FA22:6)+NH4 | Type76 | Isomer12 | 8116706.92 |
| 444 | TAG(56:8/FA16:0)+NH4 | Type77 | Isomer1 | 1508171.80 |
| 445 | TAG(56:8/FA16:1)+NH4 | Type77 | Isomer2 | 337328.64 |
| 446 | TAG(56:8/FA18:1)+NH4 | Type77 | Isomer3 | 1584703.62 |
| 447 | TAG(56:8/FA18:2)+NH4 | Type77 | Isomer4 | 2721111.60 |
| 448 | TAG(56:8/FA18:3)+NH4 | Type77 | Isomer5 | 992928.13 |
| 449 | TAG(56:8/FA20:4)+NH4 | Type77 | Isomer6 | 5640430.62 |
| 450 | TAG(56:8/FA20:5)+NH4 | Type77 | Isomer7 | 2162394.92 |
| 451 | TAG(56:8/FA22:5)+NH4 | Type77 | Isomer8 | 729956.40 |
| 452 | TAG(56:8/FA22:6)+NH4 | Type77 | Isomer9 | 6788200.39 |
| 453 | TAG(56:9/FA18:3)+NH4 | Type78 | Isomer1 | 400725.54 |
| 454 | TAG(56:9/FA20:4)+NH4 | Type78 | Isomer2 | 927180.39 |

| | | | | |
|---|---|---|---|---|
| 455 | TAG(56:9/FA20:5)+NH4 | Type78 | Isomer3 | 1043155.82 |
| 456 | TAG(56:9/FA22:6)+NH4 | Type78 | Isomer4 | 1264188.38 |
| 457 | TAG(57:10/FA22:6)+NH4 | Type79 | Isomer1 | 74015.83 |
| 458 | TAG(57:2/FA18:1)+NH4 | Type80 | Isomer1 | 1371787.89 |
| 459 | TAG(57:3/FA18:2)+NH4 | Type81 | Isomer1 | 813856.56 |
| 460 | TAG(58:10/FA18:2)+NH4 | Type82 | Isomer1 | 815087.76 |
| 461 | TAG(58:10/FA20:4)+NH4 | Type82 | Isomer2 | 476554.56 |
| 462 | TAG(58:10/FA20:5)+NH4 | Type82 | Isomer3 | 197314.78 |
| 463 | TAG(58:10/FA22:5)+NH4 | Type82 | Isomer4 | 171818.00 |
| 464 | TAG(58:10/FA22:6)+NH4 | Type82 | Isomer5 | 2103208.45 |
| 465 | TAG(58:2/FA18:1)+NH4 | Type83 | Isomer1 | 5380361.00 |
| 466 | TAG(58:6/FA16:0)+NH4 | Type84 | Isomer1 | 771164.55 |
| 467 | TAG(58:6/FA18:0)+NH4 | Type84 | Isomer2 | 494262.43 |
| 468 | TAG(58:6/FA18:1)+NH4 | Type84 | Isomer3 | 1638151.44 |
| 469 | TAG(58:6/FA20:4)+NH4 | Type84 | Isomer4 | 869189.28 |
| 470 | TAG(58:6/FA22:4)+NH4 | Type84 | Isomer5 | 870731.40 |
| 471 | TAG(58:6/FA22:5)+NH4 | Type84 | Isomer6 | 627577.47 |
| 472 | TAG(58:7/FA16:0)+NH4 | Type85 | Isomer1 | 694370.87 |
| 473 | TAG(58:7/FA18:0)+NH4 | Type85 | Isomer2 | 541515.66 |
| 474 | TAG(58:7/FA18:1)+NH4 | Type85 | Isomer3 | 1181260.08 |
| 475 | TAG(58:7/FA18:2)+NH4 | Type85 | Isomer4 | 1063210.32 |
| 476 | TAG(58:7/FA20:4)+NH4 | Type85 | Isomer5 | 712580.09 |
| 477 | TAG(58:7/FA22:4)+NH4 | Type85 | Isomer6 | 801390.12 |
| 478 | TAG(58:7/FA22:5)+NH4 | Type85 | Isomer7 | 1690522.51 |
| 479 | TAG(58:7/FA22:6)+NH4 | Type85 | Isomer8 | 1592644.90 |
| 480 | TAG(58:8/FA18:1)+NH4 | Type86 | Isomer1 | 1513330.84 |
| 481 | TAG(58:8/FA18:2)+NH4 | Type86 | Isomer2 | 900850.25 |
| 482 | TAG(58:8/FA20:3)+NH4 | Type86 | Isomer3 | 206796.18 |
| 483 | TAG(58:8/FA20:4)+NH4 | Type86 | Isomer4 | 598284.62 |
| 484 | TAG(58:8/FA22:5)+NH4 | Type86 | Isomer5 | 1628548.31 |
| 485 | TAG(58:8/FA22:6)+NH4 | Type86 | Isomer6 | 5328007.79 |
| 486 | TAG(58:9/FA18:1)+NH4 | Type87 | Isomer1 | 1124951.22 |
| 487 | TAG(58:9/FA18:2)+NH4 | Type87 | Isomer2 | 1020003.54 |
| 488 | TAG(58:9/FA20:4)+NH4 | Type87 | Isomer3 | 625264.16 |
| 489 | TAG(58:9/FA22:5)+NH4 | Type87 | Isomer4 | 882293.50 |
| 490 | TAG(58:9/FA22:6)+NH4 | Type87 | Isomer5 | 4873986.72 |
| 491 | TAG(60:10/FA22:5)+NH4 | Type88 | Isomer1 | 157266.85 |
| 492 | TAG(60:10/FA22:6)+NH4 | Type88 | Isomer2 | 279300.06 |
| 493 | TAG(60:11/FA22:5)+NH4 | Type89 | Isomer1 | 48869.42 |
| 494 | TAG(60:11/FA22:6)+NH4 | Type89 | Isomer2 | 203676.72 |
| 495 | TAG(60:12/FA22:6)+NH4 | Type90 | Isomer1 | 97536.40 |
| 496 | DAG(14:0/18:1)+NH4 | Type1 | Isomer1 | 152836.50 |
| 497 | DAG(14:0/18:3)+NH4 | Type2 | Isomer1 | 131022.64 |
| 498 | DAG(14:0/20:0)+NH4 | Type3 | Isomer1 | 111652.57 |
| 499 | DAG(16:0/18:0)+NH4 | Type3 | Isomer2 | 169333.75 |
| 500 | DAG(16:1/18:2)+NH4 | Type4 | Isomer1 | 57113.37 |

| | | | | |
|---|---|---|---|---|
| 501 | DAG(16:0/18:3)+NH4 | Type4 | Isomer2 | 57400.69 |
| 502 | DAG(16:1/18:3)+NH4 | Type5 | Isomer1 | 66690.97 |
| 503 | DAG(14:0/20:4)+NH4 | Type5 | Isomer2 | 40195.56 |
| 504 | DAG(18:0/18:1)+NH4 | Type6 | Isomer1 | 37068.98 |
| 505 | DAG(18:1/18:1)+NH4 | Type7 | Isomer1 | 121753.46 |
| 506 | DAG(18:1/18:2)+NH4 | Type8 | Isomer1 | 61187.83 |
| 507 | DAG(16:0/20:3)+NH4 | Type8 | Isomer2 | 34565.13 |
| 508 | DAG(18:2/18:3)+NH4 | Type9 | Isomer1 | 57379.87 |
| 509 | DAG(16:1/20:4)+NH4 | Type9 | Isomer2 | 78064.14 |
| 510 | DAG(18:1/20:1)+NH4 | Type10 | Isomer1 | 107799.19 |
| 511 | DAG(18:1/20:2)+NH4 | Type11 | Isomer1 | 42821.77 |
| 512 | DAG(18:1/20:3)+NH4 | Type12 | Isomer1 | 21625.28 |
| 513 | DAG(18:2/20:4)+NH4 | Type13 | Isomer1 | 36651.61 |
| 514 | DAG(18:1/20:5)+NH4 | Type13 | Isomer2 | 42707.79 |
| 515 | DAG(16:0/22:6)+NH4 | Type13 | Isomer3 | 34619.74 |
| 516 | DAG(18:2/20:5)+NH4 | Type14 | Isomer1 | 50633.62 |
| 517 | DAG(20:0/20:0)+NH4 | Type15 | Isomer1 | 6143154.40 |
| 518 | DAG(18:2/22:5)+NH4 | Type16 | Isomer1 | 42919.56 |
| 519 | DAG(18:1/22:6)+NH4 | Type16 | Isomer2 | 42743.67 |
| 520 | LPC(14:0)+AcO | _ | _ | 304284.46 |
| 521 | LPC(16:0)+AcO | _ | _ | 43716182.55 |
| 522 | LPC(16:1)+AcO | _ | _ | 1052940.44 |
| 523 | LPC(18:0)+AcO | _ | _ | 25513124.86 |
| 524 | LPC(18:1)+AcO | _ | _ | 11411518.71 |
| 525 | LPC(18:2)+AcO | _ | _ | 17085200.42 |
| 526 | LPC(18:3)+AcO | _ | _ | 226265.56 |
| 527 | LPC(20:0)+AcO | _ | _ | 220829.04 |
| 528 | LPC(20:1)+AcO | _ | _ | 360259.09 |
| 529 | LPC(20:2)+AcO | _ | _ | 314708.54 |
| 530 | LPC(20:3)+AcO | _ | _ | 1003535.41 |
| 531 | LPC(20:4)+AcO | _ | _ | 1005492.12 |
| 532 | LPC(20:5)+AcO | _ | _ | 20756.00 |
| 533 | LPC(22:4)+AcO | _ | _ | 87462.67 |
| 534 | LPC(22:5)+AcO | _ | _ | 53393.02 |
| 535 | LPC(22:6)+AcO | _ | _ | 18248.94 |
| 536 | PC(14:0/14:0)+AcO | Type1 | Isomer1 | 182675.10 |
| 537 | PC(14:0/18:1)+AcO | Type2 | Isomer1 | 2822208.23 |
| 538 | PC(14:0/18:2)+AcO | Type3 | Isomer1 | 4222808.34 |
| 539 | PC(14:0/18:3)+AcO | Type5 | Isomer1 | 567869.80 |
| 540 | PC(14:0/20:2)+AcO | Type5 | Isomer2 | 76859.54 |
| 541 | PC(14:0/20:3)+AcO | Type6 | Isomer1 | 246606.47 |
| 542 | PC(14:0/20:4)+AcO | Type6 | Isomer2 | 565061.25 |
| 543 | PC(14:0/20:5)+AcO | Type7 | Isomer1 | 266356.45 |
| 544 | PC(14:0/22:4)+AcO | Type8 | Isomer1 | 58947.91 |
| 545 | PC(14:0/22:5)+AcO | Type9 | Isomer1 | 67874.04 |
| 546 | PC(14:0/22:6)+AcO | Type10 | Isomer1 | 85896.04 |

| | | | | |
|---|---|---|---|---|
| 547 | PC(14:1/14:1)+AcO | Type10 | Isomer2 | 37290.01 |
| 548 | PC(16:0/14:0)+AcO | Type11 | Isomer1 | 2573343.54 |
| 549 | PC(16:0/16:0)+AcO | Type11 | Isomer2 | 41899759.81 |
| 550 | PC(16:0/16:1)+AcO | Type11 | Isomer3 | 7438608.07 |
| 551 | PC(16:0/18:0)+AcO | Type11 | Isomer4 | 12495345.40 |
| 552 | PC(16:0/18:1)+AcO | Type12 | Isomer1 | 93448598.93 |
| 553 | PC(16:0/18:2)+AcO | Type12 | Isomer2 | 233362047.59 |
| 554 | PC(16:0/18:3)+AcO | Type12 | Isomer3 | 3623848.07 |
| 555 | PC(16:0/20:1)+AcO | Type12 | Isomer4 | 877983.77 |
| 556 | PC(16:0/20:2)+AcO | Type13 | Isomer1 | 3103088.02 |
| 557 | PC(16:0/20:3)+AcO | Type14 | Isomer1 | 28939429.82 |
| 558 | PC(16:0/20:4)+AcO | Type15 | Isomer1 | 52570511.51 |
| 559 | PC(16:0/20:5)+AcO | Type16 | Isomer1 | 1439397.30 |
| 560 | PC(16:0/22:4)+AcO | Type16 | Isomer2 | 2482320.86 |
| 561 | PC(16:0/22:5)+AcO | Type16 | Isomer3 | 5220513.64 |
| 562 | PC(16:1/18:1)+AcO | Type17 | Isomer1 | 3120454.12 |
| 563 | PC(16:1/18:2)+AcO | Type17 | Isomer2 | 1132701.11 |
| 564 | PC(16:0/22:6)+AcO | Type17 | Isomer3 | 3421125.39 |
| 565 | PC(18:0/14:0)+AcO | Type18 | Isomer1 | 190876.70 |
| 566 | PC(18:0/16:1)+AcO | Type18 | Isomer2 | 451612.43 |
| 567 | PC(18:0/18:0)+AcO | Type18 | Isomer3 | 1928097.08 |
| 568 | PC(18:0/18:1)+AcO | Type19 | Isomer1 | 18794377.79 |
| 569 | PC(18:0/18:2)+AcO | Type19 | Isomer2 | 120378433.34 |
| 570 | PC(18:0/18:3)+AcO | Type19 | Isomer3 | 1385735.17 |
| 571 | PC(18:0/20:0)+AcO | Type19 | Isomer4 | 839919.18 |
| 572 | PC(18:0/20:1)+AcO | Type20 | Isomer1 | 278183.81 |
| 573 | PC(18:0/20:2)+AcO | Type20 | Isomer2 | 1517951.99 |
| 574 | PC(18:0/20:3)+AcO | Type20 | Isomer3 | 17441848.85 |
| 575 | PC(18:0/20:4)+AcO | Type21 | Isomer1 | 37918895.95 |
| 576 | PC(18:0/20:5)+AcO | Type22 | Isomer1 | 1032311.50 |
| 577 | PC(18:0/22:4)+AcO | Type23 | Isomer1 | 1449082.16 |
| 578 | PC(18:0/22:5)+AcO | Type23 | Isomer2 | 2825534.89 |
| 579 | PC(18:0/22:6)+AcO | Type24 | Isomer1 | 2091211.98 |
| 580 | PC(18:1/16:1)+AcO | Type24 | Isomer2 | 3369816.49 |
| 581 | PC(18:1/18:1)+AcO | Type25 | Isomer1 | 12147445.61 |
| 582 | PC(18:1/18:2)+AcO | Type25 | Isomer2 | 30147163.14 |
| 583 | PC(18:1/18:3)+AcO | Type25 | Isomer3 | 436091.59 |
| 584 | PC(18:1/20:1)+AcO | Type25 | Isomer4 | 212629.68 |
| 585 | PC(18:1/20:2)+AcO | Type26 | Isomer1 | 506553.03 |
| 586 | PC(18:1/20:3)+AcO | Type26 | Isomer2 | 3565651.90 |
| 587 | PC(18:1/20:4)+AcO | Type26 | Isomer3 | 6888023.76 |
| 588 | PC(18:1/20:5)+AcO | Type26 | Isomer4 | 202030.30 |
| 589 | PC(18:1/22:4)+AcO | Type27 | Isomer1 | 244705.41 |
| 590 | PC(18:1/22:5)+AcO | Type27 | Isomer2 | 393608.79 |
| 591 | PC(18:1/22:6)+AcO | Type27 | Isomer3 | 355783.54 |
| 592 | PC(18:2/16:1)+AcO | Type27 | Isomer4 | 4410273.72 |

| | | | | |
|---|---|---|---|---|
| 593 | PC(18:2/18:2)+AcO | Type28 | Isomer1 | 15493832.51 |
| 594 | PC(18:2/18:3)+AcO | Type28 | Isomer2 | 356885.61 |
| 595 | PC(18:2/20:1)+AcO | Type28 | Isomer3 | 428567.28 |
| 596 | PC(18:2/20:2)+AcO | Type29 | Isomer1 | 457415.35 |
| 597 | PC(18:2/20:3)+AcO | Type30 | Isomer1 | 1198325.13 |
| 598 | PC(18:2/20:4)+AcO | Type31 | Isomer1 | 2703068.84 |
| 599 | PC(18:2/20:5)+AcO | Type32 | Isomer1 | 92399.08 |
| 600 | PC(18:2/22:4)+AcO | Type33 | Isomer1 | 63426.59 |
| 601 | PC(18:2/22:5)+AcO | Type33 | Isomer2 | 86541.92 |
| 602 | PC(18:2/22:6)+AcO | Type34 | Isomer1 | 75422.52 |
| 603 | PC(20:0/16:1)+AcO | Type34 | Isomer2 | 40938.60 |
| 604 | PC(20:0/18:1)+AcO | Type34 | Isomer3 | 524774.16 |
| 605 | PC(20:0/18:3)+AcO | Type35 | Isomer1 | 12301.01 |
| 606 | PC(20:0/20:1)+AcO | Type35 | Isomer2 | 18523.43 |
| 607 | PC(20:0/20:2)+AcO | Type35 | Isomer3 | 10350.61 |
| 608 | PC(20:0/20:3)+AcO | Type36 | Isomer1 | 185608.47 |
| 609 | PC(20:0/20:4)+AcO | Type36 | Isomer2 | 621002.82 |
| 610 | PC(20:0/20:5)+AcO | Type37 | Isomer1 | 11538.27 |
| 611 | PC(20:0/22:4)+AcO | Type38 | Isomer1 | 17042.49 |
| 612 | PC(20:0/22:6)+AcO | Type40 | Isomer1 | 30664.85 |
| 613 | LPE(16:0)-H | _ | _ | 809803.56 |
| 614 | LPE(18:0)-H | _ | _ | 3956433.47 |
| 615 | LPE(18:1)-H | _ | _ | 3603409.52 |
| 616 | LPE(18:2)-H | _ | _ | 3844946.49 |
| 617 | LPE(18:3)-H | _ | _ | 47621.28 |
| 618 | LPE(20:0)-H | _ | _ | 49040.48 |
| 619 | LPE(20:1)-H | _ | _ | 102928.78 |
| 620 | LPE(20:2)-H | _ | _ | 56756.61 |
| 621 | LPE(20:3)-H | _ | _ | 290330.93 |
| 622 | LPE(20:4)-H | _ | _ | 786467.13 |
| 623 | LPE(20:5)-H | _ | _ | 7794.87 |
| 624 | LPE(22:4)-H | _ | _ | 64986.65 |
| 625 | LPE(22:5)-H | _ | _ | 76903.44 |
| 626 | LPE(22:6)-H | _ | _ | 26654.06 |
| 627 | PE(14:0/14:0)-H | Type1 | Isomer1 | 37128.19 |
| 628 | PE(14:0/18:1)-H | Type3 | Isomer1 | 283719.82 |
| 629 | PE(14:0/18:2)-H | Type5 | Isomer1 | 345447.30 |
| 630 | PE(14:0/18:3)-H | Type5 | Isomer2 | 10733.59 |
| 631 | PE(14:0/20:1)-H | Type6 | Isomer1 | 7044.00 |
| 632 | PE(14:0/20:2)-H | Type6 | Isomer2 | 7675.58 |
| 633 | PE(14:0/20:3)-H | Type7 | Isomer1 | 25271.67 |
| 634 | PE(14:0/20:4)-H | Type8 | Isomer1 | 45934.42 |
| 635 | PE(14:0/22:4)-H | Type9 | Isomer1 | 16777.78 |
| 636 | PE(14:0/22:5)-H | Type10 | Isomer1 | 14142.44 |
| 637 | PE(14:0/22:6)-H | Type10 | Isomer2 | 16843.28 |
| 638 | PE(16:0/14:0)-H | Type10 | Isomer3 | 203890.07 |

| | | | | |
|---|---|---|---|---|
| 639 | PE(16:0/16:0)-H | Type11 | Isomer1 | 2980617.99 |
| 640 | PE(16:0/16:1)-H | Type11 | Isomer2 | 975389.03 |
| 641 | PE(16:0/18:1)-H | Type11 | Isomer3 | 12732650.06 |
| 642 | PE(16:0/18:2)-H | Type12 | Isomer1 | 30014323.06 |
| 643 | PE(16:0/18:3)-H | Type12 | Isomer2 | 732315.43 |
| 644 | PE(16:0/20:1)-H | Type12 | Isomer3 | 194814.92 |
| 645 | PE(16:0/20:2)-H | Type13 | Isomer1 | 405964.23 |
| 646 | PE(16:0/20:3)-H | Type15 | Isomer1 | 1891977.74 |
| 647 | PE(16:0/20:4)-H | Type16 | Isomer1 | 8657711.50 |
| 648 | PE(16:0/20:5)-H | Type16 | Isomer2 | 105797.35 |
| 649 | PE(16:0/22:4)-H | Type17 | Isomer1 | 909508.97 |
| 650 | PE(16:0/22:5)-H | Type17 | Isomer2 | 1601175.67 |
| 651 | PE(16:0/22:6)-H | Type17 | Isomer3 | 1099779.66 |
| 652 | PE(18:0/14:0)-H | Type18 | Isomer1 | 70911.70 |
| 653 | PE(18:0/16:0)-H | Type18 | Isomer2 | 651908.95 |
| 654 | PE(18:0/16:1)-H | Type18 | Isomer3 | 257200.54 |
| 655 | PE(18:0/18:0)-H | Type19 | Isomer1 | 5207680.54 |
| 656 | PE(18:0/18:1)-H | Type19 | Isomer2 | 84912093.94 |
| 657 | PE(18:0/18:2)-H | Type19 | Isomer3 | 224331555.72 |
| 658 | PE(18:0/18:3)-H | Type19 | Isomer4 | 3297257.74 |
| 659 | PE(18:0/20:1)-H | Type20 | Isomer1 | 925561.94 |
| 660 | PE(18:0/20:2)-H | Type20 | Isomer2 | 2685810.87 |
| 661 | PE(18:0/20:3)-H | Type20 | Isomer3 | 10976935.96 |
| 662 | PE(18:0/20:4)-H | Type21 | Isomer1 | 49175395.85 |
| 663 | PE(18:0/20:5)-H | Type23 | Isomer1 | 645383.84 |
| 664 | PE(18:0/22:4)-H | Type24 | Isomer1 | 2330173.63 |
| 665 | PE(18:0/22:5)-H | Type24 | Isomer2 | 3367089.78 |
| 666 | PE(18:0/22:6)-H | Type25 | Isomer1 | 1692011.40 |
| 667 | PE(18:1/16:1)-H | Type25 | Isomer2 | 1151761.50 |
| 668 | PE(18:1/18:1)-H | Type25 | Isomer3 | 13290537.01 |
| 669 | PE(18:1/18:2)-H | Type26 | Isomer1 | 28574727.24 |
| 670 | PE(18:1/18:3)-H | Type26 | Isomer2 | 604413.02 |
| 671 | PE(18:1/20:1)-H | Type26 | Isomer3 | 222992.70 |
| 672 | PE(18:1/20:2)-H | Type26 | Isomer4 | 234501.92 |
| 673 | PE(18:1/20:3)-H | Type27 | Isomer1 | 1107487.49 |
| 674 | PE(18:1/20:4)-H | Type27 | Isomer2 | 5480070.64 |
| 675 | PE(18:1/20:5)-H | Type27 | Isomer3 | 82969.27 |
| 676 | PE(18:1/22:4)-H | Type27 | Isomer4 | 261234.56 |
| 677 | PE(18:1/22:5)-H | Type28 | Isomer1 | 259556.21 |
| 678 | PE(18:1/22:6)-H | Type28 | Isomer2 | 205990.02 |
| 679 | PE(18:2/16:1)-H | Type28 | Isomer3 | 2265975.00 |
| 680 | PE(18:2/18:2)-H | Type29 | Isomer1 | 8647443.27 |
| 681 | PE(18:2/18:3)-H | Type33 | Isomer2 | 393187.39 |
| 682 | PE(18:2/20:1)-H | Type34 | Isomer1 | 229380.12 |
| 683 | PE(18:2/20:2)-H | Type34 | Isomer2 | 175114.14 |
| 684 | PE(18:2/20:3)-H | Type35 | Isomer1 | 262406.49 |

| | | | | |
|---|---|---|---|---|
| 685 | PE(18:2/20:4)-H | Type35 | Isomer2 | 926228.05 |
| 686 | PE(18:2/20:5)-H | Type35 | Isomer3 | 13938.36 |
| 687 | PE(18:2/22:4)-H | Type36 | Isomer1 | 60071.40 |
| 688 | PE(18:2/22:5)-H | Type36 | Isomer2 | 49004.38 |
| 689 | PE(18:2/22:6)-H | Type37 | Isomer1 | 27981.59 |
| 690 | PE(O-16:0/18:0)-H | _ | _ | 88351.15 |
| 691 | PE(O-16:0/18:1)-H | _ | _ | 1108405.78 |
| 692 | PE(O-16:0/18:2)-H | _ | _ | 2201707.41 |
| 693 | PE(O-16:0/18:3)-H | _ | _ | 58470.04 |
| 694 | PE(O-16:0/20:1)-H | _ | _ | 91036.60 |
| 695 | PE(O-16:0/20:2)-H | _ | _ | 55971.45 |
| 696 | PE(O-16:0/20:3)-H | _ | _ | 376695.73 |
| 697 | PE(O-16:0/20:4)-H | _ | _ | 2375081.58 |
| 698 | PE(O-16:0/20:5)-H | _ | _ | 156859.09 |
| 699 | PE(O-16:0/22:4)-H | _ | _ | 877771.66 |
| 700 | PE(O-16:0/22:5)-H | _ | _ | 1242782.19 |
| 701 | PE(O-16:0/22:6)-H | _ | _ | 365072.31 |
| 702 | PE(O-18:0/16:0)-H | _ | _ | 3639554.90 |
| 703 | PE(O-18:0/16:1)-H | _ | _ | 457959.95 |
| 704 | PE(O-18:0/18:0)-H | _ | _ | 279546.89 |
| 705 | PE(O-18:0/18:1)-H | _ | _ | 3251504.55 |
| 706 | PE(O-18:0/18:2)-H | _ | _ | 8299159.77 |
| 707 | PE(O-18:0/18:3)-H | _ | _ | 156800.33 |
| 708 | PE(O-18:0/20:1)-H | _ | _ | 106025.89 |
| 709 | PE(O-18:0/20:2)-H | _ | _ | 150671.36 |
| 710 | PE(O-18:0/20:3)-H | _ | _ | 1110812.21 |
| 711 | PE(O-18:0/20:4)-H | _ | _ | 5777322.83 |
| 712 | PE(O-18:0/20:5)-H | _ | _ | 289176.37 |
| 713 | PE(O-18:0/22:4)-H | _ | _ | 872717.63 |
| 714 | PE(O-18:0/22:5)-H | _ | _ | 1012273.16 |
| 715 | PE(O-18:0/22:6)-H | _ | _ | 424188.13 |
| 716 | PE(P-14:0/18:0)-H | _ | _ | 126153.39 |
| 717 | PE(P-14:0/18:1)-H | _ | _ | 151277.61 |
| 718 | PE(P-16:0/16:0)-H | _ | _ | 669660.73 |
| 719 | PE(P-16:0/16:1)-H | _ | _ | 200408.83 |
| 720 | PE(P-16:0/18:0)-H | _ | _ | 386783.76 |
| 721 | PE(P-16:0/18:1)-H | _ | _ | 7815718.21 |
| 722 | PE(P-16:0/18:2)-H | _ | _ | 22186322.58 |
| 723 | PE(P-16:0/18:3)-H | _ | _ | 271853.17 |
| 724 | PE(P-16:0/20:1)-H | _ | _ | 345267.42 |
| 725 | PE(P-16:0/20:2)-H | _ | _ | 279185.65 |
| 726 | PE(P-16:0/20:3)-H | _ | _ | 4009258.91 |
| 727 | PE(P-16:0/20:4)-H | _ | _ | 25218675.01 |
| 728 | PE(P-16:0/20:5)-H | _ | _ | 337482.99 |
| 729 | PE(P-16:0/22:4)-H | _ | _ | 6666907.18 |
| 730 | PE(P-16:0/22:5)-H | _ | _ | 6882465.80 |

| | | | | |
|---|---|---|---|---|
| 731 | PE(P-16:0/22:6)-H | _ | _ | 2072535.64 |
| 732 | PE(P-16:1/18:1)-H | _ | _ | 172438.77 |
| 733 | PE(P-18:0/16:0)-H | _ | _ | 2468623.63 |
| 734 | PE(P-18:0/16:1)-H | _ | _ | 570292.44 |
| 735 | PE(P-18:0/18:0)-H | _ | _ | 428571.09 |
| 736 | PE(P-18:0/18:1)-H | _ | _ | 11055888.54 |
| 737 | PE(P-18:0/18:2)-H | _ | _ | 33621185.36 |
| 738 | PE(P-18:0/18:3)-H | _ | _ | 470885.02 |
| 739 | PE(P-18:0/20:1)-H | _ | _ | 252072.40 |
| 740 | PE(P-18:0/20:2)-H | _ | _ | 449879.75 |
| 741 | PE(P-18:0/20:3)-H | _ | _ | 7462302.89 |
| 742 | PE(P-18:0/20:4)-H | _ | _ | 44748816.88 |
| 743 | PE(P-18:0/20:5)-H | _ | _ | 986890.05 |
| 744 | PE(P-18:0/22:4)-H | _ | _ | 4997315.45 |
| 745 | PE(P-18:0/22:5)-H | _ | _ | 5229045.24 |
| 746 | PE(P-18:0/22:6)-H | _ | _ | 2433519.54 |
| 747 | PE(P-18:1/16:0)-H | _ | _ | 1642706.32 |
| 748 | PE(P-18:1/16:1)-H | _ | _ | 190971.99 |
| 749 | PE(P-18:1/18:0)-H | _ | _ | 257347.49 |
| 750 | PE(P-18:1/18:1)-H | _ | _ | 4927860.83 |
| 751 | PE(P-18:1/18:2)-H | _ | _ | 13737179.15 |
| 752 | PE(P-18:1/20:1)-H | _ | _ | 144368.47 |
| 753 | PE(P-18:1/20:2)-H | _ | _ | 174700.25 |
| 754 | PE(P-18:1/20:3)-H | _ | _ | 2420492.16 |
| 755 | PE(P-18:1/20:4)-H | _ | _ | 18018429.99 |
| 756 | PE(P-18:1/20:5)-H | _ | _ | 289220.90 |
| 757 | PE(P-18:1/22:4)-H | _ | _ | 1419696.15 |
| 758 | PE(P-18:1/22:5)-H | _ | _ | 1515476.68 |
| 759 | PE(P-18:1/22:6)-H | _ | _ | 863287.00 |
| 760 | PE(P-18:2/18:2)-H | _ | _ | 1549621.83 |
| 761 | PE(P-18:2/20:4)-H | _ | _ | 1369409.29 |
| 762 | PE(P-18:2/22:6)-H | _ | _ | 76476.00 |
| 763 | LPG(16:0)-H | _ | _ | 100346.08 |
| 764 | LPG(18:0)-H | _ | _ | 80071.89 |
| 765 | LPG(18:1)-H | _ | _ | 163319.69 |
| 766 | LPG(18:2)-H | _ | _ | 111220.49 |
| 767 | LPG(20:0)-H | _ | _ | 16411.94 |
| 768 | LPG(20:2)-H | _ | _ | 7766.91 |
| 769 | LPG(22:4)-H | _ | _ | 4372.18 |
| 770 | LPG(22:5)-H | _ | _ | 2754.42 |
| 771 | PG(14:0/14:0)-H | Type1 | Isomer1 | 49045.60 |
| 772 | PG(14:0/18:2)-H | Type3 | Isomer1 | 186229.94 |
| 773 | PG(14:0/18:3)-H | Type5 | Isomer1 | 42543.75 |
| 774 | PG(14:0/20:1)-H | Type5 | Isomer2 | 9628.32 |
| 775 | PG(14:0/20:3)-H | Type6 | Isomer1 | 14668.91 |
| 776 | PG(14:0/20:4)-H | Type7 | Isomer1 | 79654.00 |

| | | | | |
|---|---|---|---|---|
| 777 | PG(14:0/20:5)-H | Type8 | Isomer1 | 47757.44 |
| 778 | PG(14:0/22:4)-H | Type9 | Isomer1 | 5009.17 |
| 779 | PG(14:0/22:5)-H | Type10 | Isomer1 | 32234.52 |
| 780 | PG(14:0/22:6)-H | Type10 | Isomer2 | 43100.00 |
| 781 | PG(16:0/14:0)-H | Type10 | Isomer3 | 41147.50 |
| 782 | PG(16:0/16:0)-H | Type11 | Isomer1 | 491432.40 |
| 783 | PG(16:0/16:1)-H | Type11 | Isomer2 | 240538.63 |
| 784 | PG(16:0/18:0)-H | Type12 | Isomer1 | 689232.39 |
| 785 | PG(16:0/18:1)-H | Type12 | Isomer2 | 2583771.06 |
| 786 | PG(16:0/18:2)-H | Type12 | Isomer3 | 1245846.23 |
| 787 | PG(16:0/18:3)-H | Type13 | Isomer1 | 28507.55 |
| 788 | PG(16:0/20:1)-H | Type14 | Isomer1 | 21125.94 |
| 789 | PG(16:0/20:2)-H | Type15 | Isomer1 | 251412.37 |
| 790 | PG(16:0/20:3)-H | Type16 | Isomer1 | 109722.53 |
| 791 | PG(16:0/20:4)-H | Type16 | Isomer2 | 334107.45 |
| 792 | PG(16:0/20:5)-H | Type16 | Isomer3 | 112776.56 |
| 793 | PG(16:0/22:5)-H | Type17 | Isomer1 | 39341.26 |
| 794 | PG(16:0/22:6)-H | Type17 | Isomer2 | 27512.17 |
| 795 | PG(18:0/14:0)-H | Type17 | Isomer3 | 23046.60 |
| 796 | PG(18:0/16:1)-H | Type18 | Isomer1 | 44104.55 |
| 797 | PG(18:0/18:0)-H | Type18 | Isomer2 | 143947.82 |
| 798 | PG(18:0/18:1)-H | Type18 | Isomer3 | 2052322.41 |
| 799 | PG(18:0/18:2)-H | Type19 | Isomer1 | 1930054.50 |
| 800 | PG(18:0/18:3)-H | Type19 | Isomer2 | 20490.89 |
| 801 | PG(18:0/20:0)-H | Type19 | Isomer3 | 213104.23 |
| 802 | PG(18:0/20:1)-H | Type19 | Isomer4 | 24882.59 |
| 803 | PG(18:0/20:2)-H | Type20 | Isomer1 | 172604.83 |
| 804 | PG(18:0/20:3)-H | Type20 | Isomer2 | 55416.82 |
| 805 | PG(18:0/20:4)-H | Type20 | Isomer3 | 150904.44 |
| 806 | PG(18:0/22:5)-H | Type21 | Isomer1 | 15785.24 |
| 807 | PG(18:0/22:6)-H | Type22 | Isomer1 | 5260.38 |
| 808 | PG(18:1/16:1)-H | Type23 | Isomer1 | 17238548.96 |
| 809 | PG(18:1/18:1)-H | Type23 | Isomer2 | 3366340.16 |
| 810 | PG(18:1/18:2)-H | Type24 | Isomer1 | 23272232.49 |
| 811 | PG(18:1/18:3)-H | Type24 | Isomer2 | 222766.59 |
| 812 | PG(18:1/20:1)-H | Type24 | Isomer3 | 79338.47 |
| 813 | PG(18:1/20:2)-H | Type25 | Isomer1 | 383501.77 |
| 814 | PG(18:1/20:3)-H | Type25 | Isomer2 | 3195968.99 |
| 815 | PG(18:1/20:4)-H | Type25 | Isomer3 | 4410238.19 |
| 816 | PG(18:1/20:5)-H | Type26 | Isomer1 | 130492.47 |
| 817 | PG(18:1/22:4)-H | Type26 | Isomer2 | 252691.34 |
| 818 | PG(18:1/22:5)-H | Type26 | Isomer3 | 400879.49 |
| 819 | PG(18:1/22:6)-H | Type27 | Isomer1 | 197544.37 |
| 820 | PG(18:2/16:1)-H | Type27 | Isomer2 | 62872828.01 |
| 821 | PG(18:2/18:2)-H | Type27 | Isomer3 | 5687424.96 |
| 822 | PG(18:2/18:3)-H | Type28 | Isomer1 | 78068.80 |

| | | | | |
|---|---|---|---|---|
| 823 | PG(18:2/20:1)-H | Type28 | Isomer2 | 41393.05 |
| 824 | PG(18:2/20:2)-H | Type28 | Isomer3 | 112588.64 |
| 825 | PG(18:2/20:3)-H | Type29 | Isomer1 | 485301.16 |
| 826 | PG(18:2/20:4)-H | Type31 | Isomer1 | 1204059.40 |
| 827 | PG(18:2/20:5)-H | Type32 | Isomer1 | 24526.40 |
| 828 | PG(18:2/22:4)-H | Type33 | Isomer1 | 44194.54 |
| 829 | PG(18:2/22:5)-H | Type34 | Isomer1 | 61878.24 |
| 830 | PG(18:2/22:6)-H | Type34 | Isomer2 | 34414.92 |
| 831 | PG(20:0/16:1)-H | Type34 | Isomer3 | 72980.67 |
| 832 | PG(20:0/18:1)-H | Type35 | Isomer1 | 1000622.49 |
| 833 | PG(20:0/18:2)-H | Type35 | Isomer2 | 2806271.09 |
| 834 | PG(20:0/20:2)-H | Type35 | Isomer3 | 45982.03 |
| 835 | PG(20:0/20:3)-H | Type36 | Isomer1 | 362948.53 |
| 836 | PG(20:0/20:4)-H | Type36 | Isomer2 | 2186928.87 |
| 837 | PG(20:0/20:5)-H | Type37 | Isomer1 | 37387.85 |
| 838 | PG(20:0/22:4)-H | Type38 | Isomer1 | 313902.24 |
| 839 | PG(20:0/22:5)-H | Type39 | Isomer1 | 217473.44 |
| 840 | PG(20:0/22:6)-H | Type40 | Isomer1 | 77131.81 |
| 841 | PI(14:0/14:0)-H | Type1 | Isomer1 | 5774.39 |
| 842 | PI(14:0/18:2)-H | Type3 | Isomer1 | 357829.49 |
| 843 | PI(14:0/20:1)-H | Type5 | Isomer1 | 9672.58 |
| 844 | PI(14:0/20:2)-H | Type6 | Isomer1 | 11773.63 |
| 845 | PI(14:0/20:3)-H | Type7 | Isomer1 | 67824.00 |
| 846 | PI(14:0/20:4)-H | Type9 | Isomer1 | 272254.64 |
| 847 | PI(14:0/22:4)-H | Type10 | Isomer1 | 35562.57 |
| 848 | PI(14:0/22:5)-H | Type10 | Isomer2 | 14492.07 |
| 849 | PI(14:0/22:6)-H | Type10 | Isomer3 | 6594.91 |
| 850 | PI(16:0/14:0)-H | Type11 | Isomer1 | 11507.49 |
| 851 | PI(16:0/16:0)-H | Type11 | Isomer2 | 887701.54 |
| 852 | PI(16:0/16:1)-H | Type12 | Isomer1 | 35956.60 |
| 853 | PI(16:0/18:0)-H | Type12 | Isomer2 | 701405.30 |
| 854 | PI(16:0/18:1)-H | Type13 | Isomer1 | 331620.17 |
| 855 | PI(16:0/18:2)-H | Type15 | Isomer1 | 740912.19 |
| 856 | PI(16:0/18:3)-H | Type16 | Isomer1 | 619949.46 |
| 857 | PI(16:0/20:1)-H | Type16 | Isomer2 | 8585.85 |
| 858 | PI(16:0/20:2)-H | Type16 | Isomer3 | 22435.50 |
| 859 | PI(16:0/20:4)-H | Type17 | Isomer1 | 354877.51 |
| 860 | PI(16:0/20:5)-H | Type17 | Isomer2 | 14739.74 |
| 861 | PI(16:0/22:4)-H | Type17 | Isomer3 | 10982.40 |
| 862 | PI(16:0/22:5)-H | Type18 | Isomer1 | 25514.83 |
| 863 | PI(18:0/16:1)-H | Type18 | Isomer2 | 36731.37 |
| 864 | PI(18:0/18:0)-H | Type19 | Isomer1 | 654641.27 |
| 865 | PI(18:0/18:1)-H | Type19 | Isomer2 | 264080.62 |
| 866 | PI(18:0/18:2)-H | Type19 | Isomer3 | 1422559.30 |
| 867 | PI(18:0/18:3)-H | Type19 | Isomer4 | 93648.29 |
| 868 | PI(18:0/20:0)-H | Type20 | Isomer1 | 1327411.51 |

| | | | | |
|---|---|---|---|---|
| 869 | PI(18:0/20:2)-H | Type20 | Isomer2 | 31221.79 |
| 870 | PI(18:0/20:3)-H | Type21 | Isomer1 | 566673.47 |
| 871 | PI(18:0/20:4)-H | Type22 | Isomer1 | 4248223.79 |
| 872 | PI(18:0/20:5)-H | Type23 | Isomer1 | 32085.62 |
| 873 | PI(18:0/22:4)-H | Type24 | Isomer1 | 17828.46 |
| 874 | PI(18:0/22:5)-H | Type24 | Isomer2 | 42819.20 |
| 875 | PI(18:0/22:6)-H | Type25 | Isomer1 | 23318.78 |
| 876 | PI(18:1/18:1)-H | Type25 | Isomer2 | 413938.30 |
| 877 | PI(18:1/18:2)-H | Type25 | Isomer3 | 298809.31 |
| 878 | PI(18:1/18:3)-H | Type26 | Isomer1 | 73080.48 |
| 879 | PI(18:1/20:2)-H | Type26 | Isomer2 | 43514.97 |
| 880 | PI(18:1/20:3)-H | Type26 | Isomer3 | 401608.60 |
| 881 | PI(18:1/20:5)-H | Type26 | Isomer4 | 13884.40 |
| 882 | PI(18:1/22:4)-H | Type27 | Isomer1 | 29745.88 |
| 883 | PI(18:1/22:6)-H | Type27 | Isomer2 | 11867.10 |
| 884 | PI(18:2/18:2)-H | Type27 | Isomer3 | 604666.31 |
| 885 | PI(18:2/20:1)-H | Type28 | Isomer1 | 18356.81 |
| 886 | PI(18:2/20:2)-H | Type28 | Isomer2 | 15351.09 |
| 887 | PI(18:2/20:3)-H | Type30 | Isomer1 | 100641.37 |
| 888 | PI(18:2/20:4)-H | Type31 | Isomer1 | 429304.11 |
| 889 | PI(18:2/22:4)-H | Type32 | Isomer1 | 8847.85 |
| 890 | PI(20:0/16:1)-H | Type33 | Isomer1 | 32230.35 |
| 891 | PI(20:0/18:1)-H | Type33 | Isomer2 | 168396.61 |
| 892 | PI(20:0/18:2)-H | Type34 | Isomer1 | 172176.02 |
| 893 | PI(20:0/20:1)-H | Type34 | Isomer2 | 15763.18 |
| 894 | PI(20:0/20:2)-H | Type34 | Isomer3 | 12969.84 |
| 895 | PI(20:0/20:3)-H | Type35 | Isomer1 | 44049.77 |
| 896 | PI(20:0/20:4)-H | Type35 | Isomer2 | 202779.95 |
| 897 | PI(20:0/20:5)-H | Type36 | Isomer1 | 6235.51 |
| 898 | PI(20:0/22:5)-H | Type39 | Isomer1 | 6340.55 |
| 899 | LPS(16:0)-H | _ | _ | 4601.95 |
| 900 | LPS(20:0)-H | _ | _ | 86160.93 |
| 901 | LPS(20:1)-H | _ | _ | 139152.57 |
| 902 | PS(14:0/14:0)-H | Type1 | Isomer1 | 14164.98 |
| 903 | PS(14:0/18:1)-H | Type3 | Isomer1 | 218377.65 |
| 904 | PS(14:0/18:2)-H | Type5 | Isomer1 | 381020.38 |
| 905 | PS(14:0/18:3)-H | Type5 | Isomer2 | 12407.11 |
| 906 | PS(14:0/20:1)-H | Type6 | Isomer1 | 7608.32 |
| 907 | PS(14:0/20:2)-H | Type7 | Isomer1 | 5764.35 |
| 908 | PS(14:0/20:3)-H | Type8 | Isomer1 | 49452.38 |
| 909 | PS(14:0/20:4)-H | Type9 | Isomer1 | 198309.86 |
| 910 | PS(14:0/22:4)-H | Type10 | Isomer1 | 32699.00 |
| 911 | PS(14:0/22:5)-H | Type10 | Isomer2 | 37823.04 |
| 912 | PS(14:0/22:6)-H | Type10 | Isomer3 | 22622.76 |
| 913 | PS(16:0/14:0)-H | Type11 | Isomer1 | 13266.07 |
| 914 | PS(16:0/16:0)-H | Type11 | Isomer2 | 128691.03 |

| | | | | |
|---|---|---|---|---|
| 915 | PS(16:0/18:0)-H | Type11 | Isomer3 | 348341.00 |
| 916 | PS(16:0/18:1)-H | Type12 | Isomer1 | 779833.97 |
| 917 | PS(16:0/18:2)-H | Type12 | Isomer2 | 700398.49 |
| 918 | PS(16:0/18:3)-H | Type12 | Isomer3 | 144313.77 |
| 919 | PS(16:0/20:1)-H | Type13 | Isomer1 | 3448.41 |
| 920 | PS(16:0/20:2)-H | Type15 | Isomer1 | 14653.97 |
| 921 | PS(16:0/20:3)-H | Type16 | Isomer1 | 115323.45 |
| 922 | PS(16:0/20:4)-H | Type16 | Isomer2 | 256328.61 |
| 923 | PS(16:0/20:5)-H | Type16 | Isomer3 | 89009.23 |
| 924 | PS(16:0/22:4)-H | Type17 | Isomer1 | 28799.63 |
| 925 | PS(18:0/14:0)-H | Type17 | Isomer2 | 23139.56 |
| 926 | PS(18:0/16:1)-H | Type17 | Isomer3 | 113652.23 |
| 927 | PS(18:0/18:0)-H | Type18 | Isomer1 | 4714507.40 |
| 928 | PS(18:0/18:1)-H | Type18 | Isomer2 | 1008409.16 |
| 929 | PS(18:0/18:2)-H | Type18 | Isomer3 | 974342.35 |
| 930 | PS(18:0/18:3)-H | Type19 | Isomer1 | 237868.39 |
| 931 | PS(18:0/20:0)-H | Type19 | Isomer2 | 3969253.06 |
| 932 | PS(18:0/20:1)-H | Type19 | Isomer3 | 52426.33 |
| 933 | PS(18:0/20:2)-H | Type19 | Isomer4 | 42074.79 |
| 934 | PS(18:0/20:3)-H | Type20 | Isomer1 | 42107.55 |
| 935 | PS(18:0/20:4)-H | Type20 | Isomer2 | 380497.56 |
| 936 | PS(18:0/20:5)-H | Type20 | Isomer3 | 81991.12 |
| 937 | PS(18:0/22:4)-H | Type21 | Isomer1 | 23716.32 |
| 938 | PS(18:0/22:5)-H | Type22 | Isomer1 | 33330.48 |
| 939 | PS(18:0/22:6)-H | Type23 | Isomer1 | 6418.01 |
| 940 | PS(18:1/16:1)-H | Type23 | Isomer2 | 1418732.13 |
| 941 | PS(18:1/18:1)-H | Type24 | Isomer1 | 332362.43 |
| 942 | PS(18:1/18:2)-H | Type24 | Isomer2 | 1304852.21 |
| 943 | PS(18:1/18:3)-H | Type25 | Isomer1 | 94334.06 |
| 944 | PS(18:1/20:2)-H | Type25 | Isomer2 | 26433.00 |
| 945 | PS(18:1/20:3)-H | Type25 | Isomer3 | 214580.76 |
| 946 | PS(18:1/20:4)-H | Type25 | Isomer4 | 771374.84 |
| 947 | PS(18:1/20:5)-H | Type26 | Isomer1 | 42863.74 |
| 948 | PS(18:1/22:4)-H | Type26 | Isomer2 | 50262.36 |
| 949 | PS(18:1/22:5)-H | Type26 | Isomer3 | 66414.94 |
| 950 | PS(18:1/22:6)-H | Type26 | Isomer4 | 41607.74 |
| 951 | PS(18:2/16:1)-H | Type27 | Isomer1 | 4513549.80 |
| 952 | PS(18:2/18:2)-H | Type27 | Isomer2 | 1133388.05 |
| 953 | PS(18:2/18:3)-H | Type27 | Isomer3 | 46775.78 |
| 954 | PS(18:2/20:1)-H | Type28 | Isomer1 | 46087.45 |
| 955 | PS(18:2/20:2)-H | Type28 | Isomer2 | 58607.39 |
| 956 | PS(18:2/20:3)-H | Type29 | Isomer1 | 382800.84 |
| 957 | PS(18:2/20:4)-H | Type30 | Isomer1 | 1646203.74 |
| 958 | PS(18:2/20:5)-H | Type31 | Isomer1 | 53095.54 |
| 959 | PS(18:2/22:4)-H | Type32 | Isomer1 | 72311.04 |
| 960 | PS(18:2/22:5)-H | Type33 | Isomer1 | 83194.26 |

| | | | | |
|---|---|---|---|---|
| 961 | PS(18:2/22:6)-H | Type33 | Isomer2 | 79397.12 |
| 962 | PS(20:0/16:1)-H | Type34 | Isomer1 | 277715.06 |
| 963 | PS(20:0/18:1)-H | Type34 | Isomer2 | 368895.25 |
| 964 | PS(20:0/18:2)-H | Type35 | Isomer1 | 4372426.77 |
| 965 | PS(20:0/18:3)-H | Type35 | Isomer2 | 104021.02 |
| 966 | PS(20:0/20:1)-H | Type35 | Isomer3 | 53766.89 |
| 967 | PS(20:0/20:2)-H | Type36 | Isomer1 | 109357.48 |
| 968 | PS(20:0/20:3)-H | Type36 | Isomer2 | 531609.49 |
| 969 | PS(20:0/20:4)-H | Type37 | Isomer1 | 1773541.70 |
| 970 | PS(20:0/22:4)-H | Type38 | Isomer1 | 117925.39 |
| 971 | PS(20:0/22:5)-H | Type39 | Isomer1 | 129249.98 |
| 972 | PS(20:0/22:6)-H | Type40 | Isomer1 | 71217.92 |
| 973 | PA(14:0/22:5)-H | Type6 | Isomer1 | 2562.81 |
| 974 | PA(16:0/16:1)-H | Type9 | Isomer1 | 14072.92 |
| 975 | PA(16:0/18:0)-H | Type10 | Isomer1 | 34769.87 |
| 976 | PA(16:0/18:2)-H | Type11 | Isomer1 | 245519.16 |
| 977 | PA(16:0/20:4)-H | Type12 | Isomer1 | 36103.39 |
| 978 | PA(16:0/22:5)-H | Type16 | Isomer1 | 7629.18 |
| 979 | PA(18:0/16:1)-H | Type16 | Isomer2 | 52845.66 |
| 980 | PA(18:0/18:1)-H | Type17 | Isomer1 | 1818239.89 |
| 981 | PA(18:0/18:2)-H | Type17 | Isomer2 | 5578222.67 |
| 982 | PA(18:0/18:3)-H | Type18 | Isomer1 | 72121.14 |
| 983 | PA(18:0/20:0)-H | Type18 | Isomer2 | 155661.39 |
| 984 | PA(18:0/20:1)-H | Type19 | Isomer1 | 72888.19 |
| 985 | PA(18:0/20:2)-H | Type19 | Isomer2 | 62853.32 |
| 986 | PA(18:0/20:3)-H | Type19 | Isomer3 | 894528.34 |
| 987 | PA(18:0/20:4)-H | Type20 | Isomer1 | 6080316.31 |
| 988 | PA(18:0/20:5)-H | Type22 | Isomer1 | 81414.08 |
| 989 | PA(18:0/22:4)-H | Type23 | Isomer1 | 1563117.17 |
| 990 | PA(18:0/22:5)-H | Type23 | Isomer2 | 1620437.99 |
| 991 | PA(18:0/22:6)-H | Type24 | Isomer1 | 485499.49 |
| 992 | PA(18:1/18:1)-H | Type24 | Isomer2 | 581684.61 |
| 993 | PA(18:1/18:2)-H | Type24 | Isomer3 | 1194677.94 |
| 994 | PA(18:1/18:3)-H | Type25 | Isomer1 | 17256.25 |
| 995 | PA(18:1/20:1)-H | Type25 | Isomer2 | 3835.03 |
| 996 | PA(18:1/20:2)-H | Type25 | Isomer3 | 28595.03 |
| 997 | PA(18:1/20:3)-H | Type26 | Isomer1 | 156720.26 |
| 998 | PA(18:1/20:4)-H | Type26 | Isomer2 | 260690.47 |
| 999 | PA(18:1/20:5)-H | Type26 | Isomer3 | 6234.39 |
| 1000 | PA(18:1/22:4)-H | Type27 | Isomer1 | 27890.59 |
| 1001 | PA(18:1/22:5)-H | Type27 | Isomer2 | 28130.28 |
| 1002 | PA(18:2/16:1)-H | Type27 | Isomer3 | 26081.35 |
| 1003 | PA(18:2/18:2)-H | Type28 | Isomer1 | 87412.62 |
| 1004 | PA(18:2/20:2)-H | Type28 | Isomer2 | 6017.36 |
| 1005 | PA(18:2/20:4)-H | Type30 | Isomer1 | 19652.07 |
| 1006 | PA(18:2/22:4)-H | Type31 | Isomer1 | 3437.35 |

| | | | | |
|---|---|---|---|---|
| 1007 | PA(20:0/16:1)-H | Type32 | Isomer1 | 54739.79 |
| 1008 | PA(20:0/18:1)-H | Type33 | Isomer1 | 2084962.10 |
| 1009 | PA(20:0/18:2)-H | Type33 | Isomer2 | 9339261.61 |
| 1010 | PA(20:0/18:3)-H | Type34 | Isomer1 | 103320.01 |
| 1011 | PA(20:0/20:1)-H | Type34 | Isomer2 | 27340.24 |
| 1012 | PA(20:0/20:2)-H | Type34 | Isomer3 | 72528.55 |
| 1013 | PA(20:0/20:3)-H | Type35 | Isomer1 | 1546545.03 |
| 1014 | PA(20:0/20:4)-H | Type35 | Isomer2 | 11256901.74 |
| 1015 | PA(20:0/20:5)-H | Type35 | Isomer3 | 258104.54 |
| 1016 | PA(20:0/22:4)-H | Type38 | Isomer1 | 1060730.33 |
| 1017 | PA(20:0/22:5)-H | Type39 | Isomer1 | 1120601.09 |
| 1018 | PA(20:0/22:6)-H | Type40 | Isomer1 | 580953.96 |

Supplementary- table 4:

| S.No. | Lipid class | Chain length | Un-saturation | Sum |
|---|---|---|---|---|
| 1 | PC | 28 | 0 | 182675.10 |
| 2 | PC | 32 | 2 | 4222808.34 |
| 3 | PC | 32 | 3 | 567869.80 |
| 4 | PC | 34 | 4 | 565061.25 |
| 5 | PC | 34 | 5 | 266356.45 |
| 6 | PC | 36 | 6 | 85896.04 |
| 7 | PC | 28 | 2 | 37290.01 |
| 8 | PC | 30 | 0 | 2573343.54 |
| 9 | PC | 32 | 1 | 10260816.30 |
| 10 | PC | 34 | 0 | 12495345.40 |
| 11 | PC | 32 | 0 | 42090636.51 |
| 12 | PC | 34 | 1 | 93900211.36 |
| 13 | PC | 36 | 0 | 1928097.08 |
| 14 | PC | 38 | 0 | 839919.18 |
| 15 | PC | 34 | 2 | 239929177.74 |
| 16 | PC | 36 | 2 | 135628966.97 |
| 17 | PC | 36 | 3 | 60472328.13 |
| 18 | PC | 38 | 2 | 1730581.66 |
| 19 | PC | 34 | 3 | 9413429.38 |
| 20 | PC | 36 | 4 | 68559383.52 |
| 21 | PC | 36 | 5 | 1864156.95 |
| 22 | PC | 38 | 4 | 44424284.06 |
| 23 | PC | 38 | 5 | 14339174.03 |
| 24 | PC | 38 | 6 | 6326224.53 |
| 25 | PC | 38 | 7 | 92399.08 |

| | | | | |
|---|---|---|---|---|
| 26 | PC | 40 | 6 | 2548247.37 |
| 27 | PC | 40 | 7 | 442325.46 |
| 28 | PC | 40 | 8 | 75422.52 |
| 29 | PC | 36 | 1 | 19713300.16 |
| 30 | PC | 38 | 1 | 802957.97 |
| 31 | PC | 38 | 3 | 18389270.17 |
| 32 | PC | 40 | 1 | 18523.43 |
| 33 | PC | 40 | 2 | 10350.61 |
| 34 | PC | 40 | 3 | 185608.47 |
| 35 | PC | 40 | 4 | 2070084.98 |
| 36 | PC | 40 | 5 | 3081778.56 |
| 37 | PC | 42 | 4 | 17042.49 |
| 38 | PC | 42 | 6 | 30664.85 |
| 39 | PE | 28 | 0 | 37128.19 |
| 40 | PE | 32 | 2 | 345447.30 |
| 41 | PE | 32 | 3 | 10733.59 |
| 42 | PE | 34 | 4 | 45934.42 |
| 43 | PE | 36 | 6 | 16843.28 |
| 44 | PE | 30 | 0 | 203890.07 |
| 45 | PE | 32 | 1 | 1259108.85 |
| 46 | PE | 32 | 0 | 3051529.69 |
| 47 | PE | 34 | 0 | 651908.95 |
| 48 | PE | 34 | 1 | 12996894.60 |
| 49 | PE | 36 | 0 | 5207680.54 |
| 50 | PE | 36 | 1 | 85106908.86 |
| 51 | PE | 38 | 1 | 925561.94 |
| 52 | PE | 40 | 4 | 2330173.63 |
| 53 | PE | 34 | 2 | 31173760.14 |
| 54 | PE | 36 | 2 | 238028056.95 |
| 55 | PE | 36 | 3 | 33763962.72 |
| 56 | PE | 38 | 2 | 2908803.57 |
| 57 | PE | 40 | 5 | 3628324.34 |
| 58 | PE | 34 | 3 | 3023562.10 |
| 59 | PE | 36 | 4 | 17926345.57 |
| 60 | PE | 36 | 5 | 513127.18 |
| 61 | PE | 38 | 3 | 11440818.00 |
| 62 | PE | 38 | 4 | 51367506.45 |
| 63 | PE | 38 | 5 | 7989036.64 |
| 64 | PE | 38 | 6 | 2108976.99 |
| 65 | PE | 38 | 7 | 13938.36 |
| 66 | PE | 40 | 6 | 2011639.01 |
| 67 | PE | 40 | 7 | 254994.40 |
| 68 | PE | 40 | 8 | 27981.59 |
| 69 | PG | 28 | 0 | 49045.60 |
| 70 | PG | 32 | 2 | 186229.94 |
| 71 | PG | 32 | 3 | 42543.75 |

| | | | | |
|---|---|---|---|---|
| 72 | PG | 34 | 4 | 79654.00 |
| 73 | PG | 34 | 5 | 47757.44 |
| 74 | PG | 36 | 6 | 43100.00 |
| 75 | PG | 30 | 0 | 41147.50 |
| 76 | PG | 32 | 1 | 240538.63 |
| 77 | PG | 34 | 0 | 689232.39 |
| 78 | PG | 32 | 0 | 514479.01 |
| 79 | PG | 34 | 1 | 2637503.93 |
| 80 | PG | 36 | 0 | 143947.82 |
| 81 | PG | 38 | 0 | 213104.23 |
| 82 | PG | 34 | 2 | 18484395.19 |
| 83 | PG | 36 | 2 | 5547807.03 |
| 84 | PG | 36 | 3 | 23402445.90 |
| 85 | PG | 34 | 3 | 62916004.47 |
| 86 | PG | 36 | 4 | 6249308.17 |
| 87 | PG | 36 | 5 | 223079.87 |
| 88 | PG | 38 | 3 | 480311.63 |
| 89 | PG | 38 | 4 | 3459462.07 |
| 90 | PG | 38 | 5 | 4934880.61 |
| 91 | PG | 38 | 6 | 1362064.04 |
| 92 | PG | 38 | 7 | 24526.40 |
| 93 | PG | 40 | 6 | 450334.42 |
| 94 | PG | 40 | 7 | 259422.60 |
| 95 | PG | 40 | 8 | 34414.92 |
| 96 | PG | 36 | 1 | 2146429.02 |
| 97 | PG | 38 | 1 | 1025505.08 |
| 98 | PG | 38 | 2 | 3058214.39 |
| 99 | PG | 40 | 2 | 45982.03 |
| 100 | PG | 40 | 3 | 362948.53 |
| 101 | PG | 40 | 4 | 2186928.87 |
| 102 | PG | 40 | 5 | 305864.43 |
| 103 | PG | 42 | 4 | 313902.24 |
| 104 | PG | 42 | 5 | 217473.44 |
| 105 | PG | 42 | 6 | 77131.81 |
| 106 | PI | 28 | 0 | 5774.39 |
| 107 | PI | 32 | 2 | 357829.49 |
| 108 | PI | 34 | 4 | 272254.64 |
| 109 | PI | 36 | 6 | 6594.91 |
| 110 | PI | 30 | 0 | 11507.49 |
| 111 | PI | 32 | 0 | 887701.54 |
| 112 | PI | 32 | 1 | 35956.60 |
| 113 | PI | 34 | 0 | 701405.30 |
| 114 | PI | 34 | 2 | 752685.82 |
| 115 | PI | 34 | 3 | 687773.45 |
| 116 | PI | 36 | 5 | 29231.81 |
| 117 | PI | 34 | 1 | 378024.12 |

| | | | | |
|---|---|---|---|---|
| 118 | PI | 36 | 0 | 654641.27 |
| 119 | PI | 38 | 0 | 1327411.51 |
| 120 | PI | 36 | 2 | 1858933.10 |
| 121 | PI | 36 | 3 | 392457.60 |
| 122 | PI | 40 | 7 | 11867.10 |
| 123 | PI | 36 | 4 | 1068186.86 |
| 124 | PI | 38 | 3 | 628545.25 |
| 125 | PI | 38 | 4 | 4676165.88 |
| 126 | PI | 38 | 5 | 158241.81 |
| 127 | PI | 38 | 6 | 443188.51 |
| 128 | PI | 40 | 6 | 32166.64 |
| 129 | PI | 36 | 1 | 304896.82 |
| 130 | PI | 38 | 1 | 168396.61 |
| 131 | PI | 38 | 2 | 203397.81 |
| 132 | PI | 40 | 1 | 15763.18 |
| 133 | PI | 40 | 2 | 12969.84 |
| 134 | PI | 40 | 3 | 44049.77 |
| 135 | PI | 40 | 4 | 220608.41 |
| 136 | PI | 40 | 5 | 78800.58 |
| 137 | PI | 42 | 5 | 6340.55 |
| 138 | PS | 28 | 0 | 14164.98 |
| 139 | PS | 32 | 1 | 218377.65 |
| 140 | PS | 32 | 2 | 381020.38 |
| 141 | PS | 32 | 3 | 12407.11 |
| 142 | PS | 34 | 4 | 198309.86 |
| 143 | PS | 36 | 6 | 22622.76 |
| 144 | PS | 30 | 0 | 13266.07 |
| 145 | PS | 34 | 0 | 348341.00 |
| 146 | PS | 32 | 0 | 151830.59 |
| 147 | PS | 34 | 1 | 901094.53 |
| 148 | PS | 36 | 0 | 4714507.40 |
| 149 | PS | 38 | 0 | 3969253.06 |
| 150 | PS | 34 | 2 | 2124894.97 |
| 151 | PS | 36 | 2 | 1321358.75 |
| 152 | PS | 36 | 3 | 1658044.05 |
| 153 | PS | 40 | 5 | 83592.84 |
| 154 | PS | 34 | 3 | 4707315.95 |
| 155 | PS | 36 | 4 | 1516749.72 |
| 156 | PS | 36 | 5 | 173608.06 |
| 157 | PS | 38 | 4 | 682485.34 |
| 158 | PS | 38 | 5 | 1236166.79 |
| 159 | PS | 38 | 6 | 1689067.48 |
| 160 | PS | 38 | 7 | 53095.54 |
| 161 | PS | 40 | 6 | 145143.99 |
| 162 | PS | 40 | 7 | 124802.00 |
| 163 | PS | 40 | 8 | 79397.12 |

| | | | | |
|---|---|---|---|---|
| 164 | PS | 36 | 1 | 1289572.63 |
| 165 | PS | 38 | 1 | 421321.58 |
| 166 | PS | 38 | 2 | 4414501.56 |
| 167 | PS | 38 | 3 | 218649.02 |
| 168 | PS | 40 | 1 | 53766.89 |
| 169 | PS | 40 | 2 | 109357.48 |
| 170 | PS | 40 | 3 | 531609.49 |
| 171 | PS | 40 | 4 | 1797258.02 |
| 172 | PS | 42 | 4 | 117925.39 |
| 173 | PS | 42 | 5 | 129249.98 |
| 174 | PS | 42 | 6 | 71217.92 |
| 175 | PA | 36 | 5 | 2562.81 |
| 176 | PA | 32 | 1 | 14072.92 |
| 177 | PA | 34 | 0 | 34769.87 |
| 178 | PA | 34 | 2 | 245519.16 |
| 179 | PA | 34 | 1 | 52845.66 |
| 180 | PA | 38 | 0 | 155661.39 |
| 181 | PA | 36 | 2 | 6159907.29 |
| 182 | PA | 36 | 3 | 1266799.09 |
| 183 | PA | 38 | 5 | 349733.73 |
| 184 | PA | 34 | 3 | 26081.35 |
| 185 | PA | 36 | 4 | 140772.26 |
| 186 | PA | 38 | 4 | 6243053.92 |
| 187 | PA | 38 | 6 | 25886.46 |
| 188 | PA | 40 | 6 | 517067.12 |
| 189 | PA | 36 | 1 | 1872979.67 |
| 190 | PA | 38 | 1 | 2157850.29 |
| 191 | PA | 38 | 2 | 9405949.96 |
| 192 | PA | 38 | 3 | 1026443.37 |
| 193 | PA | 40 | 1 | 27340.24 |
| 194 | PA | 40 | 2 | 72528.55 |
| 195 | PA | 40 | 3 | 1546545.03 |
| 196 | PA | 40 | 4 | 12820018.91 |
| 197 | PA | 40 | 5 | 1906433.13 |
| 198 | PA | 42 | 4 | 1060730.33 |
| 199 | PA | 42 | 5 | 1120601.09 |
| 200 | PA | 42 | 6 | 580953.96 |

Supplementary- table 5:

| Lipid standards | LoB (AUC) |
|---|---|
| SM | 175092.84 |
| Cer | 364623.75 |
| TG | 140710.81 |
| DG | 88877.39 |

| | |
|---|---|
| LPC | 45326.22 |
| PC | 25164.35 |
| LPE | 48182.94 |
| PE | 96532.49 |
| PG | 168948.15 |
| PI | 46159.94 |
| PS | 68699.00 |
| PA | 441488.86 |

Supplementary-table 6:

| Sample Name _(n=3) | % RECOVERY | RSD | REFERENCE | RSD |
|---|---|---|---|---|
| SM | 105.26 | 5.90 | 100.00 | 3.82 |
| CER | 100.32 | 25.68 | 100.00 | 23.61 |
| TAG | 113.19 | 4.15 | 100.00 | 27.42 |
| DAG | 137.25 | 5.73 | 100.00 | 17.81 |
| LPC | 97.03 | 8.77 | 100.00 | 1.98 |
| PC | 75.04 | 28.03 | 100.00 | 20.85 |
| LPE | 76.28 | 34.75 | 100.00 | 31.83 |
| PE | 80.90 | 5.90 | 100.00 | 8.80 |
| PG | 105.52 | 7.58 | 100.00 | 2.99 |
| PI | 69.75 | 19.67 | 100.00 | 15.36 |
| PS | 74.38 | 24.59 | 100.00 | 36.81 |
| PA | 74.96 | 31.19 | 100.00 | 25.19 |

Supplementary- table 7:

| S.No. | Lipid species | CV1 | CV2 | CV3 |
|---|---|---|---|---|
| 1 | SM(14:0)+H | 19.21 | 8.81 | 2.87 |
| 2 | SM(16:0)+H | 10.07 | 1.84 | 7.18 |
| 3 | SM(18:0)+H | 14.09 | 4.1 | 10.91 |
| 4 | SM(18:1)+H | 17.45 | 4.27 | 8.69 |
| 5 | SM(20:0)+H | 3.07 | 1.14 | 1.14 |
| 6 | SM(20:1)+H | 12.49 | 3.64 | 15.05 |
| 7 | SM(22:0)+H | 2.67 | 2.59 | 2.48 |
| 8 | SM(22:1)+H | 8.11 | 1.88 | 8.13 |
| 9 | SM(24:0)+H | 4.97 | 3.43 | 4.5 |
| 10 | SM(24:1)+H | 3.4 | 3.17 | 4.26 |
| 11 | SM(26:0)+H | 4.07 | 7.87 | 18.96 |
| 12 | SM(26:1)+H | 13.53 | 6.43 | 3.87 |
| 13 | CE(24:0)+H | 42.99 | 5.25 | 37.59 |

| | | | | |
|---|---|---|---|---|
| 14 | CE(22:6)+H | 24.01 | 37.24 | 21.34 |
| 15 | CE(20:0)+H | 26.72 | 51.61 | 17.36 |
| 16 | CE(20:1)+H | 34.5 | 54.11 | 30.57 |
| 17 | CE(22:5)+H | 33.2 | NA | 20.12 |
| 18 | CE(14:0)+H | NA | 48.05 | 21.85 |
| 19 | CE(16:0)+H | 35.96 | NA | NA |
| 20 | CE(16:1)+H | 59.06 | NA | 36.37 |
| 21 | CE(18:0)+H | 17.98 | NA | 18.48 |
| 22 | CE(18:1)+H | 23.17 | NA | 25.29 |
| 23 | CE(18:2)+H | 37.12 | 7.29 | 28.23 |
| 24 | CE(18:3)+H | 31.52 | 24.13 | 26.56 |
| 25 | CE(20:2)+H | 35.07 | NA | 19.82 |
| 26 | CE(20:3)+H | 49.9 | 71.68 | 17.34 |
| 27 | CE(20:4)+H | 41.4 | 11.98 | 41.74 |
| 28 | CE(20:5)+H | 19.65 | 16.64 | 28.35 |
| 29 | CE(22:0)+H | 34.32 | NA | 21.35 |
| 30 | CE(22:1)+H | 46.71 | NA | 18.73 |
| 31 | CE(22:2)+H | 25.83 | 11.61 | 16.67 |
| 32 | CE(22:4)+H | 31.96 | 60.19 | 19.6 |
| 33 | CE(24:1)+H | NA | NA | NA |
| 34 | CER(14:0)+H | 23.76 | NA | NA |
| 35 | CER(16:0)+H | 36.59 | NA | 30.17 |
| 36 | CER(18:0)+H | 22.83 | 38.57 | 25.33 |
| 37 | CER(18:1)+H | NA | NA | NA |
| 38 | CER(20:0)+H | 57.61 | 29.04 | 23.34 |
| 39 | CER(20:1)+H | NA | NA | NA |
| 40 | CER(22:0)+H | 36.35 | 17.66 | 25.2 |
| 41 | CER(22:1)+H | 13.13 | 20.97 | 7.97 |
| 42 | CER(24:0)+H | 10.98 | 33.95 | 8.62 |
| 43 | CER(24:1)+H | 7.4 | 13.97 | 3.02 |
| 44 | CER(26:0)+H | NA | NA | NA |
| 45 | CER(26:1)+H | NA | NA | NA |
| 46 | DCER(14:0)+H | NA | 106.11 | NA |
| 47 | DCER(16:0)+H | 12.96 | NA | NA |
| 48 | DCER(18:0)+H | NA | 41.27 | NA |
| 49 | DCER(18:1)+H | 38.99 | 48 | NA |
| 50 | DCER(20:0)+H | 38.66 | 28.02 | 13.09 |
| 51 | DCER(20:1)+H | NA | 68.28 | NA |
| 52 | DCER(22:0)+H | 10.86 | 40.62 | 8.84 |
| 53 | DCER(22:1)+H | 16.25 | 19.03 | 8.35 |
| 54 | DCER(24:0)+H | 1.37 | 43.08 | 13.16 |
| 55 | DCER(24:1)+H | 6.52 | 41.98 | 9.46 |
| 56 | DCER(26:0)+H | 80.12 | NA | NA |

| | | | | |
|---|---|---|---|---|
| 57 | DCER(26:1)+H | NA | NA | NA |
| 58 | HCER(14:0)+H | 31.26 | 34.98 | NA |
| 59 | HCER(16:0)+H | 14.07 | 20.91 | 13.76 |
| 60 | HCER(18:0)+H | 16.39 | 14.78 | 27.01 |
| 61 | HCER(18:1)+H | 11.69 | 22.16 | 9.15 |
| 62 | HCER(20:0)+H | 17.24 | 30.21 | 15.4 |
| 63 | HCER(20:1)+H | 8.9 | 20.8 | 6.59 |
| 64 | HCER(22:0)+H | 11.11 | 27 | 15.68 |
| 65 | HCER(22:1)+H | 9.34 | 16.07 | 5.2 |
| 66 | HCER(24:0)+H | 14.56 | 21.01 | 10.7 |
| 67 | HCER(24:1)+H | 14.41 | 43.55 | 8.9 |
| 68 | HCER(26:0)+H | 55.61 | NA | 41.96 |
| 69 | HCER(26:1)+H | 37.27 | 24.91 | 47.42 |
| 70 | HCER(d18:0/18:0)+H | 25.85 | 35.7 | 38.78 |
| 71 | HCER(d18:0/20:0)+H | 59.07 | 21.47 | NA |
| 72 | HCER(d18:0/22:0)+H | 9.13 | 23.6 | 18.99 |
| 73 | HCER(d18:0/24:0)+H | 14.28 | 21.16 | 14.17 |
| 74 | HCER(d18:0/24:1)+H | 5.03 | 109.66 | 17.72 |
| 75 | HCER(d18:0/26:0)+H | 36.31 | 37.08 | 13.02 |
| 76 | HCER(d18:0/26:1)+H | 15.51 | 48.54 | 25.26 |
| 77 | LCER(14:0)+H | 28.03 | 49.7 | 16.49 |
| 78 | LCER(16:0)+H | 17.51 | 40.75 | 35.09 |
| 79 | LCER(18:0)+H | 16.11 | 22.69 | 37.59 |
| 80 | LCER(18:1)+H | 11.85 | 25.98 | NA |
| 81 | LCER(20:0)+H | 32.12 | 18.04 | 16.91 |
| 82 | LCER(20:1)+H | 34.19 | 18.25 | NA |
| 83 | LCER(22:0)+H | 26.82 | 55.8 | NA |
| 84 | LCER(22:1)+H | NA | 11.04 | 54.28 |
| 85 | LCER(24:0)+H | 127.65 | 51.72 | NA |
| 86 | LCER(24:1)+H | 43.24 | 41.8 | 36.73 |
| 87 | LCER(26:0)+H | NA | 41.67 | 104.31 |
| 88 | LCER(26:1)+H | 106.23 | 74.34 | NA |
| 89 | LCER(d18:0/18:0)+H | 31.38 | 22.24 | 55.62 |
| 90 | LCER(d18:0/20:0)+H | 58.87 | 67.48 | 44.95 |
| 91 | LCER(d18:0/22:0)+H | NA | 82.42 | 53.53 |
| 92 | LCER(d18:0/24:0)+H | 64.4 | NA | 35.54 |
| 93 | LCER(d18:0/24:1)+H | 53.68 | NA | NA |
| 94 | LCER(d18:0/26:0)+H | NA | 19.97 | 28.97 |
| 95 | LCER(d18:0/26:1)+H | NA | NA | NA |
| 96 | TAG(40:0/FA14:0)+NH4 | NA | NA | NA |
| 97 | TAG(40:0/FA16:0)+NH4 | NA | NA | NA |
| 98 | TAG(42:0/FA14:0)+NH4 | NA | NA | NA |
| 99 | TAG(42:0/FA16:0)+NH4 | NA | NA | NA |

| | | | | |
|---|---|---|---|---|
| 100 | TAG(42:1/FA14:0)+NH4 | 31.3 | 39.28 | NA |
| 101 | TAG(42:1/FA16:0)+NH4 | NA | 42.44 | 8.86 |
| 102 | TAG(42:1/FA16:1)+NH4 | 17.3 | NA | NA |
| 103 | TAG(42:1/FA18:1)+NH4 | 14.82 | NA | NA |
| 104 | TAG(42:2/FA18:2)+NH4 | 10.88 | 1.21 | 19.01 |
| 105 | TAG(44:0/FA14:0)+NH4 | NA | NA | NA |
| 106 | TAG(44:0/FA16:0)+NH4 | 0.96 | NA | NA |
| 107 | TAG(44:0/FA18:0)+NH4 | 4.94 | NA | NA |
| 108 | TAG(44:1/FA14:0)+NH4 | 11.03 | 25.5 | NA |
| 109 | TAG(44:1/FA16:0)+NH4 | 19.82 | NA | NA |
| 110 | TAG(44:1/FA16:1)+NH4 | NA | NA | NA |
| 111 | TAG(44:1/FA18:1)+NH4 | NA | NA | 17.06 |
| 112 | TAG(44:2/FA14:0)+NH4 | 15.05 | NA | 10.94 |
| 113 | TAG(44:2/FA16:0)+NH4 | 19.72 | 36.24 | 11.03 |
| 114 | TAG(44:2/FA16:1)+NH4 | NA | NA | NA |
| 115 | TAG(44:2/FA18:1)+NH4 | 17.49 | NA | NA |
| 116 | TAG(44:2/FA18:2)+NH4 | 15.12 | 12.66 | 10.01 |
| 117 | TAG(44:3/FA18:2)+NH4 | 30.78 | 24.87 | 15.68 |
| 118 | TAG(45:0/FA14:0)+NH4 | NA | NA | NA |
| 119 | TAG(45:0/FA16:0)+NH4 | NA | NA | NA |
| 120 | TAG(45:1/FA16:0)+NH4 | NA | NA | NA |
| 121 | TAG(45:1/FA18:1)+NH4 | NA | NA | NA |
| 122 | TAG(46:0/FA14:0)+NH4 | NA | NA | NA |
| 123 | TAG(46:0/FA16:0)+NH4 | 9.08 | NA | NA |
| 124 | TAG(46:0/FA18:0)+NH4 | 16.49 | 37.95 | NA |
| 125 | TAG(46:1/FA14:0)+NH4 | 15.38 | 7.77 | NA |
| 126 | TAG(46:1/FA16:0)+NH4 | 6.02 | 1.16 | NA |
| 127 | TAG(46:1/FA16:1)+NH4 | 12.34 | NA | NA |
| 128 | TAG(46:1/FA18:0)+NH4 | 40.75 | NA | NA |
| 129 | TAG(46:1/FA18:1)+NH4 | 15.26 | 4.5 | 18.13 |
| 130 | TAG(46:2/FA14:0)+NH4 | 11.05 | 11.94 | NA |
| 131 | TAG(46:2/FA16:0)+NH4 | 10.35 | 11.41 | 11.34 |
| 132 | TAG(46:2/FA16:1)+NH4 | 33.78 | NA | NA |
| 133 | TAG(46:2/FA18:1)+NH4 | 12.6 | NA | 7.08 |
| 134 | TAG(46:2/FA18:2)+NH4 | 8.13 | 12.14 | 8.76 |
| 135 | TAG(46:3/FA14:0)+NH4 | 28.81 | 12 | 19.23 |
| 136 | TAG(46:3/FA16:0)+NH4 | 18.94 | 25.54 | 8.69 |
| 137 | TAG(46:3/FA16:1)+NH4 | 52.76 | 15.17 | NA |
| 138 | TAG(46:3/FA18:1)+NH4 | 12.18 | 27.62 | 13.45 |
| 139 | TAG(46:3/FA18:2)+NH4 | 16.65 | 22.95 | 14.53 |
| 140 | TAG(46:3/FA18:3)+NH4 | 20.34 | 26.49 | 8.35 |
| 141 | TAG(46:4/FA18:2)+NH4 | 13.02 | 11.85 | 6.52 |
| 142 | TAG(47:0/FA14:0)+NH4 | 16.14 | NA | NA |

| | | | | |
|---|---|---|---|---|
| 143 | TAG(47:0/FA16:0)+NH4 | NA | NA | NA |
| 144 | TAG(47:0/FA17:0)+NH4 | 35.43 | NA | NA |
| 145 | TAG(47:1/FA14:0)+NH4 | 20.67 | NA | NA |
| 146 | TAG(47:1/FA16:0)+NH4 | NA | NA | NA |
| 147 | TAG(47:1/FA16:1)+NH4 | NA | NA | NA |
| 148 | TAG(47:1/FA17:0)+NH4 | NA | NA | NA |
| 149 | TAG(47:1/FA18:1)+NH4 | NA | NA | NA |
| 150 | TAG(47:2/FA14:0)+NH4 | 25.75 | 5.36 | NA |
| 151 | TAG(47:2/FA16:1)+NH4 | NA | NA | NA |
| 152 | TAG(47:2/FA18:1)+NH4 | NA | NA | NA |
| 153 | TAG(47:2/FA18:2)+NH4 | 35.14 | 31.24 | NA |
| 154 | TAG(48:0/FA14:0)+NH4 | 14.97 | 6.52 | 7.29 |
| 155 | TAG(48:0/FA16:0)+NH4 | 11.78 | 0.37 | NA |
| 156 | TAG(48:0/FA18:0)+NH4 | 17.96 | 4.78 | 6.14 |
| 157 | TAG(48:1/FA14:0)+NH4 | 17.1 | 3.81 | 6.2 |
| 158 | TAG(48:1/FA16:0)+NH4 | 7.97 | 8.11 | 10.43 |
| 159 | TAG(48:1/FA16:1)+NH4 | 8.75 | 4.75 | NA |
| 160 | TAG(48:1/FA18:0)+NH4 | 23.43 | 11.3 | 11.76 |
| 161 | TAG(48:1/FA18:1)+NH4 | 8.81 | 5.33 | 11.3 |
| 162 | TAG(48:2/FA14:0)+NH4 | 11.66 | 47.02 | 11.96 |
| 163 | TAG(48:2/FA16:0)+NH4 | 11.37 | 57.05 | 7.94 |
| 164 | TAG(48:2/FA16:1)+NH4 | 20.6 | 16.39 | NA |
| 165 | TAG(48:2/FA18:0)+NH4 | 19.67 | 40.87 | 13.09 |
| 166 | TAG(48:2/FA18:1)+NH4 | 16.72 | 45.42 | 3.34 |
| 167 | TAG(48:2/FA18:2)+NH4 | 7.66 | 11 | 10 |
| 168 | TAG(48:3/FA14:0)+NH4 | 11.8 | 18.24 | 11.38 |
| 169 | TAG(48:3/FA16:0)+NH4 | 14.85 | 17.91 | 13.65 |
| 170 | TAG(48:3/FA16:1)+NH4 | 14.13 | 1.79 | NA |
| 171 | TAG(48:3/FA18:1)+NH4 | 12.62 | 10.03 | 7.59 |
| 172 | TAG(48:3/FA18:2)+NH4 | 7.78 | 78.75 | 4.91 |
| 173 | TAG(48:3/FA18:3)+NH4 | 16.36 | 109.51 | 10.13 |
| 174 | TAG(48:4/FA14:0)+NH4 | 28.7 | 23.03 | 25.14 |
| 175 | TAG(48:4/FA16:0)+NH4 | 10.78 | 41.37 | 22.06 |
| 176 | TAG(48:4/FA16:1)+NH4 | 33.37 | 12.24 | NA |
| 177 | TAG(48:4/FA18:1)+NH4 | 7.69 | 22.6 | 19.99 |
| 178 | TAG(48:4/FA18:2)+NH4 | 12.33 | 16.97 | 4.21 |
| 179 | TAG(48:4/FA18:3)+NH4 | 25.87 | 35.39 | 6.51 |
| 180 | TAG(48:4/FA20:4)+NH4 | 28.47 | 32.25 | 16.65 |
| 181 | TAG(48:5/FA18:2)+NH4 | 18.49 | 54.08 | 9.57 |
| 182 | TAG(48:5/FA18:3)+NH4 | 14.15 | 25.74 | 5.88 |
| 183 | TAG(49:0/FA16:0)+NH4 | 2.49 | NA | NA |
| 184 | TAG(49:0/FA17:0)+NH4 | NA | NA | NA |
| 185 | TAG(49:0/FA18:0)+NH4 | 19.69 | NA | NA |

| | | | | |
|---|---|---|---|---|
| 186 | TAG(49:1/FA14:0)+NH4 | 13.33 | NA | NA |
| 187 | TAG(49:1/FA16:0)+NH4 | 6.11 | NA | NA |
| 188 | TAG(49:1/FA16:1)+NH4 | 17.4 | NA | NA |
| 189 | TAG(49:1/FA17:0)+NH4 | 8.99 | NA | NA |
| 190 | TAG(49:1/FA18:1)+NH4 | 4.74 | 11.77 | NA |
| 191 | TAG(49:2/FA14:0)+NH4 | 19.3 | 15.55 | NA |
| 192 | TAG(49:2/FA16:0)+NH4 | 11.14 | 10.75 | NA |
| 193 | TAG(49:2/FA16:1)+NH4 | NA | NA | NA |
| 194 | TAG(49:2/FA17:0)+NH4 | 22.19 | NA | NA |
| 195 | TAG(49:2/FA18:1)+NH4 | 9.46 | NA | NA |
| 196 | TAG(49:2/FA18:2)+NH4 | 12.98 | 8.3 | NA |
| 197 | TAG(49:3/FA16:0)+NH4 | 25.62 | 24.06 | 14.7 |
| 198 | TAG(49:3/FA16:1)+NH4 | NA | NA | NA |
| 199 | TAG(49:3/FA18:2)+NH4 | 7.77 | 21.51 | NA |
| 200 | TAG(49:3/FA18:3)+NH4 | 7.35 | 16.62 | 21.42 |
| 201 | TAG(50:0/FA14:0)+NH4 | 5.27 | NA | 18.63 |
| 202 | TAG(50:0/FA16:0)+NH4 | 10.48 | 9.65 | 8.91 |
| 203 | TAG(50:0/FA18:0)+NH4 | 11.94 | 11.13 | 10.35 |
| 204 | TAG(50:1/FA14:0)+NH4 | 18.08 | 1.56 | 2.89 |
| 205 | TAG(50:1/FA16:0)+NH4 | 3.66 | 7.67 | 4.42 |
| 206 | TAG(50:1/FA16:1)+NH4 | 14.54 | 5.88 | NA |
| 207 | TAG(50:1/FA18:0)+NH4 | 12.86 | 3.21 | 12.55 |
| 208 | TAG(50:1/FA18:1)+NH4 | 14.27 | 6.25 | 6.86 |
| 209 | TAG(50:1/FA20:1)+NH4 | 13.69 | NA | NA |
| 210 | TAG(50:2/FA14:0)+NH4 | 9.61 | 5.06 | 7.78 |
| 211 | TAG(50:2/FA16:0)+NH4 | 10.51 | 4.84 | 6.11 |
| 212 | TAG(50:2/FA16:1)+NH4 | 9.69 | 2.62 | 3.42 |
| 213 | TAG(50:2/FA18:0)+NH4 | 16.19 | 4.81 | 6.87 |
| 214 | TAG(50:2/FA18:1)+NH4 | 8.43 | 5.21 | 4.17 |
| 215 | TAG(50:2/FA18:2)+NH4 | 8.95 | 2.83 | 5.4 |
| 216 | TAG(50:2/FA20:2)+NH4 | 17.45 | 21.72 | NA |
| 217 | TAG(50:3/FA14:0)+NH4 | 13.51 | 6.74 | 6.56 |
| 218 | TAG(50:3/FA16:0)+NH4 | 12.05 | 7.65 | 6.74 |
| 219 | TAG(50:3/FA16:1)+NH4 | 10.77 | 3.26 | 7.4 |
| 220 | TAG(50:3/FA18:0)+NH4 | 30 | 13.46 | 8.04 |
| 221 | TAG(50:3/FA18:1)+NH4 | 6.88 | 5.74 | 8.87 |
| 222 | TAG(50:3/FA18:2)+NH4 | 5.5 | 5.9 | 5.3 |
| 223 | TAG(50:3/FA18:3)+NH4 | 7.78 | 5.88 | 11.37 |
| 224 | TAG(50:3/FA20:3)+NH4 | 15.13 | 18.24 | 21.05 |
| 225 | TAG(50:4/FA14:0)+NH4 | 7.53 | 9.47 | 5.99 |
| 226 | TAG(50:4/FA16:0)+NH4 | 17.14 | 16.16 | 10.25 |
| 227 | TAG(50:4/FA16:1)+NH4 | 35.59 | 5.93 | 7.02 |
| 228 | TAG(50:4/FA18:1)+NH4 | 10.93 | 9.89 | 20.98 |

| | | | | |
|---|---|---|---|---|
| 229 | TAG(50:4/FA18:2)+NH4 | 8.85 | 11.14 | 6.28 |
| 230 | TAG(50:4/FA18:3)+NH4 | 18.06 | 8.09 | 8.5 |
| 231 | TAG(50:4/FA20:3)+NH4 | 45.06 | 15.66 | 22.09 |
| 232 | TAG(50:4/FA20:4)+NH4 | 23.71 | 14.8 | 5.51 |
| 233 | TAG(50:5/FA14:0)+NH4 | 26.29 | 18.99 | 7.6 |
| 234 | TAG(50:5/FA16:0)+NH4 | 57.79 | 26.09 | 21.47 |
| 235 | TAG(50:5/FA16:1)+NH4 | 22.49 | 11.59 | 24.05 |
| 236 | TAG(50:5/FA18:1)+NH4 | 32.83 | 20.16 | 21.39 |
| 237 | TAG(50:5/FA18:2)+NH4 | 25.6 | 17.29 | 6.24 |
| 238 | TAG(50:5/FA18:3)+NH4 | 19.94 | 11.94 | 6.82 |
| 239 | TAG(50:5/FA20:4)+NH4 | 10.81 | 7.62 | 31.58 |
| 240 | TAG(50:5/FA20:5)+NH4 | 65.76 | NA | 20.95 |
| 241 | TAG(50:6/FA20:4)+NH4 | 44.34 | 14.4 | 16.3 |
| 242 | TAG(51:0/FA16:0)+NH4 | 11.5 | NA | NA |
| 243 | TAG(51:0/FA17:0)+NH4 | 3.54 | NA | NA |
| 244 | TAG(51:0/FA18:0)+NH4 | 8.47 | 7.93 | NA |
| 245 | TAG(51:1/FA16:0)+NH4 | 14.98 | 8.67 | NA |
| 246 | TAG(51:1/FA17:0)+NH4 | 19.41 | 8.47 | NA |
| 247 | TAG(51:1/FA18:0)+NH4 | 14.1 | 17.59 | NA |
| 248 | TAG(51:1/FA18:1)+NH4 | 10.29 | 3.87 | NA |
| 249 | TAG(51:2/FA16:0)+NH4 | 10.32 | 6.36 | 7.48 |
| 250 | TAG(51:2/FA16:1)+NH4 | NA | NA | NA |
| 251 | TAG(51:2/FA17:0)+NH4 | 15.36 | 8.29 | 11.01 |
| 252 | TAG(51:2/FA18:1)+NH4 | 9.79 | 3.41 | 15.1 |
| 253 | TAG(51:2/FA18:2)+NH4 | 14.87 | 3.91 | 5.58 |
| 254 | TAG(51:3/FA16:1)+NH4 | 18.22 | 19.83 | NA |
| 255 | TAG(51:3/FA17:0)+NH4 | 29.61 | 13.65 | 11.63 |
| 256 | TAG(51:3/FA18:2)+NH4 | 12.52 | 4.93 | 13.02 |
| 257 | TAG(51:3/FA18:3)+NH4 | 8.1 | 11.98 | 29.1 |
| 258 | TAG(51:4/FA16:1)+NH4 | 23.82 | 14.16 | NA |
| 259 | TAG(51:4/FA18:2)+NH4 | 14.48 | 11.26 | 3.63 |
| 260 | TAG(51:4/FA18:3)+NH4 | 15.57 | 10.35 | 13.87 |
| 261 | TAG(51:4/FA20:4)+NH4 | 36.17 | 23.48 | 24.47 |
| 262 | TAG(51:5/FA18:2)+NH4 | 10.3 | 20.36 | 16.83 |
| 263 | TAG(51:5/FA18:3)+NH4 | 26.87 | 9.59 | 10.83 |
| 264 | TAG(52:0/FA16:0)+NH4 | 13.06 | 2.85 | 7.74 |
| 265 | TAG(52:0/FA18:0)+NH4 | 7.09 | 4.67 | 15.87 |
| 266 | TAG(52:0/FA20:0)+NH4 | 15.79 | NA | 18.24 |
| 267 | TAG(52:1/FA16:0)+NH4 | 3.13 | 6.23 | 5.82 |
| 268 | TAG(52:1/FA16:1)+NH4 | 22.26 | 8.19 | NA |
| 269 | TAG(52:1/FA18:0)+NH4 | 11.79 | 3.1 | 2.79 |
| 270 | TAG(52:1/FA18:1)+NH4 | 7.93 | 2.8 | 6.22 |
| 271 | TAG(52:1/FA20:0)+NH4 | 20.97 | 8.93 | 19.86 |

| | | | | |
|---|---|---|---|---|
| 272 | TAG(52:1/FA20:1)+NH4 | 12.08 | 12.28 | 11.4 |
| 273 | TAG(52:2/FA14:0)+NH4 | 26.24 | 22.49 | 16.83 |
| 274 | TAG(52:2/FA16:0)+NH4 | 6.91 | 4.9 | 2.82 |
| 275 | TAG(52:2/FA16:1)+NH4 | 10.65 | 2.78 | 1.98 |
| 276 | TAG(52:2/FA18:0)+NH4 | 7.64 | 3.97 | 5.03 |
| 277 | TAG(52:2/FA18:1)+NH4 | 11.75 | 1.64 | 7.55 |
| 278 | TAG(52:2/FA18:2)+NH4 | 8.3 | 3.88 | 7.05 |
| 279 | TAG(52:2/FA20:0)+NH4 | 6.39 | 8.11 | 17.31 |
| 280 | TAG(52:2/FA20:1)+NH4 | 26.69 | 3.85 | 19.56 |
| 281 | TAG(52:2/FA20:2)+NH4 | 14.67 | 14.44 | 12.76 |
| 282 | TAG(52:3/FA14:0)+NH4 | 20.27 | 2.6 | 14.21 |
| 283 | TAG(52:3/FA16:0)+NH4 | 6.91 | 2.06 | 2.42 |
| 284 | TAG(52:3/FA16:1)+NH4 | 4.31 | 3.83 | 9.61 |
| 285 | TAG(52:3/FA18:0)+NH4 | 9.56 | 2.39 | 5.72 |
| 286 | TAG(52:3/FA18:1)+NH4 | 5.47 | 4.66 | 2.43 |
| 287 | TAG(52:3/FA18:2)+NH4 | 3.65 | 4.1 | 7.99 |
| 288 | TAG(52:3/FA18:3)+NH4 | 8.54 | 4.11 | 7.67 |
| 289 | TAG(52:3/FA20:0)+NH4 | 17.5 | 5.75 | 17.21 |
| 290 | TAG(52:3/FA20:1)+NH4 | 13.9 | 9.4 | 14.35 |
| 291 | TAG(52:3/FA20:2)+NH4 | 16.47 | 7.84 | 5.32 |
| 292 | TAG(52:3/FA20:3)+NH4 | 9.98 | 11.57 | 8.11 |
| 293 | TAG(52:4/FA14:0)+NH4 | 25.13 | 13.59 | 16.64 |
| 294 | TAG(52:4/FA16:0)+NH4 | 7.97 | 5.44 | 7.48 |
| 295 | TAG(52:4/FA16:1)+NH4 | 5.03 | 4.87 | 12.07 |
| 296 | TAG(52:4/FA18:0)+NH4 | 9.39 | 6.8 | 10.94 |
| 297 | TAG(52:4/FA18:1)+NH4 | 3.93 | 5.55 | 6.44 |
| 298 | TAG(52:4/FA18:2)+NH4 | 5.92 | 6.01 | 6.11 |
| 299 | TAG(52:4/FA18:3)+NH4 | 8.43 | 6.13 | 9.25 |
| 300 | TAG(52:4/FA20:0)+NH4 | 14.92 | 7.33 | 13 |
| 301 | TAG(52:4/FA20:2)+NH4 | 24.56 | 8.33 | 19.67 |
| 302 | TAG(52:4/FA20:3)+NH4 | 7.48 | 8.77 | 6.57 |
| 303 | TAG(52:4/FA20:4)+NH4 | 12.14 | 4.37 | 12.35 |
| 304 | TAG(52:4/FA22:4)+NH4 | 23.95 | 13.7 | 19.07 |
| 305 | TAG(52:5/FA14:0)+NH4 | 15.71 | 8.53 | 16.78 |
| 306 | TAG(52:5/FA16:0)+NH4 | 4.5 | 8.46 | 2.65 |
| 307 | TAG(52:5/FA16:1)+NH4 | 8.08 | 8.41 | 5.19 |
| 308 | TAG(52:5/FA18:1)+NH4 | 8.35 | 9.7 | 9.16 |
| 309 | TAG(52:5/FA18:2)+NH4 | 4.97 | 7.81 | 7.61 |
| 310 | TAG(52:5/FA18:3)+NH4 | 5.78 | 6.11 | 9.22 |
| 311 | TAG(52:5/FA20:3)+NH4 | 15.03 | 10.03 | 20.54 |
| 312 | TAG(52:5/FA20:4)+NH4 | 4.59 | 11.31 | 6.04 |
| 313 | TAG(52:5/FA20:5)+NH4 | 11.7 | 26.62 | 14.77 |
| 314 | TAG(52:5/FA22:5)+NH4 | 17.76 | 28.45 | 6.06 |

| | | | | |
|---|---|---|---|---|
| 315 | TAG(52:6/FA14:0)+NH4 | 38.98 | 20.79 | 28.25 |
| 316 | TAG(52:6/FA16:0)+NH4 | 13.37 | 16.14 | 15.14 |
| 317 | TAG(52:6/FA16:1)+NH4 | 12.42 | 10.48 | 12.1 |
| 318 | TAG(52:6/FA18:1)+NH4 | 18.73 | 12.49 | 22.2 |
| 319 | TAG(52:6/FA18:2)+NH4 | 10.21 | 8.65 | 10.17 |
| 320 | TAG(52:6/FA18:3)+NH4 | 10.06 | 10.11 | 14.29 |
| 321 | TAG(52:6/FA20:4)+NH4 | 10.16 | 13.53 | 12.29 |
| 322 | TAG(52:6/FA20:5)+NH4 | 16.7 | 13.66 | 23.9 |
| 323 | TAG(52:6/FA22:6)+NH4 | 24.97 | 11.6 | 21.45 |
| 324 | TAG(52:7/FA16:0)+NH4 | NA | NA | NA |
| 325 | TAG(52:7/FA18:1)+NH4 | 19.68 | 13.27 | 7.45 |
| 326 | TAG(52:7/FA20:5)+NH4 | 34.89 | 26.52 | 22.45 |
| 327 | TAG(52:7/FA22:6)+NH4 | 24.86 | 17.64 | 18.84 |
| 328 | TAG(52:8/FA16:1)+NH4 | NA | NA | NA |
| 329 | TAG(52:8/FA18:2)+NH4 | 15.43 | 17.78 | 21.68 |
| 330 | TAG(53:0/FA16:0)+NH4 | 19.23 | 20.33 | 4.72 |
| 331 | TAG(53:1/FA16:0)+NH4 | 19.53 | 6.98 | 10.26 |
| 332 | TAG(53:1/FA17:0)+NH4 | 6.32 | 4.38 | 11.91 |
| 333 | TAG(53:1/FA18:0)+NH4 | 13.52 | 9.6 | 5.52 |
| 334 | TAG(53:1/FA18:1)+NH4 | 6.22 | 9.73 | 16.42 |
| 335 | TAG(53:2/FA16:0)+NH4 | 13.39 | 12.41 | 14.78 |
| 336 | TAG(53:2/FA17:0)+NH4 | 3.07 | 3.57 | 6.51 |
| 337 | TAG(53:2/FA18:1)+NH4 | 10.77 | 4.86 | 8.14 |
| 338 | TAG(53:2/FA18:2)+NH4 | 11.65 | 5 | 6.81 |
| 339 | TAG(53:3/FA16:0)+NH4 | 20.17 | 18.47 | 8.77 |
| 340 | TAG(53:3/FA17:0)+NH4 | 14.41 | 8.84 | 5.75 |
| 341 | TAG(53:3/FA18:2)+NH4 | 10.01 | 6.77 | 7.53 |
| 342 | TAG(53:4/FA16:0)+NH4 | 38.15 | 7.15 | 14.99 |
| 343 | TAG(53:4/FA17:0)+NH4 | 6.15 | 5.01 | 9.21 |
| 344 | TAG(53:4/FA18:2)+NH4 | 9.21 | 6.53 | 5.66 |
| 345 | TAG(53:4/FA18:3)+NH4 | 8.49 | 1.52 | 7.21 |
| 346 | TAG(53:4/FA20:4)+NH4 | 35.12 | 17.79 | 14.47 |
| 347 | TAG(53:5/FA20:4)+NH4 | 20.52 | 17.81 | 22.75 |
| 348 | TAG(53:6/FA20:4)+NH4 | 20.46 | 13.64 | 9.22 |
| 349 | TAG(54:0/FA16:0)+NH4 | 18.41 | 18.31 | 12.6 |
| 350 | TAG(54:0/FA18:0)+NH4 | 14.65 | 16.13 | 5.86 |
| 351 | TAG(54:1/FA16:0)+NH4 | 18.91 | 4.14 | 8.29 |
| 352 | TAG(54:1/FA18:0)+NH4 | 15.52 | 4.79 | 3.47 |
| 353 | TAG(54:1/FA18:1)+NH4 | 10.58 | 6.97 | 9.91 |
| 354 | TAG(54:1/FA20:0)+NH4 | 15.35 | 4.8 | 13.27 |
| 355 | TAG(54:1/FA20:1)+NH4 | 11.67 | 10.31 | 10.93 |
| 356 | TAG(54:2/FA16:0)+NH4 | 13.37 | 8.27 | 17.79 |
| 357 | TAG(54:2/FA18:0)+NH4 | 6.51 | 3.65 | 7.74 |

| | | | | |
|---|---|---|---|---|
| 358 | TAG(54:2/FA18:1)+NH4 | 7.04 | 6.6 | 5.29 |
| 359 | TAG(54:2/FA18:2)+NH4 | 4.87 | 10.22 | 5.21 |
| 360 | TAG(54:2/FA20:0)+NH4 | 13.65 | 5.81 | 15.48 |
| 361 | TAG(54:2/FA20:1)+NH4 | 20.2 | 7.76 | 4.94 |
| 362 | TAG(54:2/FA20:2)+NH4 | 11.07 | 15.52 | 6.62 |
| 363 | TAG(54:3/FA16:0)+NH4 | 11.25 | 3.12 | 13.5 |
| 364 | TAG(54:3/FA16:1)+NH4 | 36.55 | 10.25 | 19.08 |
| 365 | TAG(54:3/FA18:0)+NH4 | 8.16 | 5.51 | 6.98 |
| 366 | TAG(54:3/FA18:1)+NH4 | 6.23 | 4.22 | 5.3 |
| 367 | TAG(54:3/FA18:2)+NH4 | 6.48 | 3.17 | 6.21 |
| 368 | TAG(54:3/FA18:3)+NH4 | 18.21 | 12.5 | 7.61 |
| 369 | TAG(54:3/FA20:1)+NH4 | 15.22 | 6.67 | 17.39 |
| 370 | TAG(54:3/FA20:2)+NH4 | 10.44 | 4.99 | 6.97 |
| 371 | TAG(54:3/FA20:3)+NH4 | 8.42 | 10.73 | 23.64 |
| 372 | TAG(54:4/FA16:0)+NH4 | 10.4 | 5.15 | 8.17 |
| 373 | TAG(54:4/FA16:1)+NH4 | 13.35 | 4.63 | 16.59 |
| 374 | TAG(54:4/FA18:0)+NH4 | 4 | 5.45 | 9.58 |
| 375 | TAG(54:4/FA18:1)+NH4 | 6.34 | 3.18 | 7.34 |
| 376 | TAG(54:4/FA18:2)+NH4 | 13.45 | 5.65 | 5.89 |
| 377 | TAG(54:4/FA18:3)+NH4 | 9.44 | 2.09 | 6.44 |
| 378 | TAG(54:4/FA20:1)+NH4 | 17.88 | 5.75 | 9.01 |
| 379 | TAG(54:4/FA20:2)+NH4 | 7.33 | 6.6 | 11.9 |
| 380 | TAG(54:4/FA20:3)+NH4 | 8.58 | 6.75 | 8.3 |
| 381 | TAG(54:4/FA20:4)+NH4 | 8.47 | 3.84 | 7.42 |
| 382 | TAG(54:4/FA22:4)+NH4 | 9.33 | 7.47 | 12.86 |
| 383 | TAG(54:5/FA16:0)+NH4 | 9.52 | 7.32 | 4.52 |
| 384 | TAG(54:5/FA16:1)+NH4 | 10.17 | 4.65 | 14.45 |
| 385 | TAG(54:5/FA18:0)+NH4 | 11.77 | 6.8 | 8.2 |
| 386 | TAG(54:5/FA18:1)+NH4 | 9.01 | 2.11 | 3.73 |
| 387 | TAG(54:5/FA18:2)+NH4 | 6.25 | 5.41 | 6.02 |
| 388 | TAG(54:5/FA18:3)+NH4 | 2.57 | 8.47 | 4.34 |
| 389 | TAG(54:5/FA20:2)+NH4 | 7.13 | 9.5 | 15.58 |
| 390 | TAG(54:5/FA20:3)+NH4 | 10.26 | 2.11 | 3.75 |
| 391 | TAG(54:5/FA20:4)+NH4 | 11.33 | 3.82 | 7.04 |
| 392 | TAG(54:5/FA20:5)+NH4 | 6.03 | 8.26 | 8.66 |
| 393 | TAG(54:5/FA22:4)+NH4 | 20.09 | 8.23 | 13.05 |
| 394 | TAG(54:5/FA22:5)+NH4 | 12.67 | 20.7 | 7.73 |
| 395 | TAG(54:6/FA16:0)+NH4 | 5.84 | 3.2 | 9.13 |
| 396 | TAG(54:6/FA16:1)+NH4 | 10.5 | 2.85 | 10.66 |
| 397 | TAG(54:6/FA18:1)+NH4 | 5.54 | 7.01 | 6.71 |
| 398 | TAG(54:6/FA18:2)+NH4 | 4.25 | 5.46 | 6.51 |
| 399 | TAG(54:6/FA18:3)+NH4 | 8.17 | 5.71 | 10.64 |
| 400 | TAG(54:6/FA20:3)+NH4 | 7.01 | 4.91 | 8.56 |

| | | | | |
|---|---|---|---|---|
| 401 | TAG(54:6/FA20:4)+NH4 | 13.13 | 4.51 | 5.04 |
| 402 | TAG(54:6/FA20:5)+NH4 | 7.52 | 2.29 | 13.91 |
| 403 | TAG(54:6/FA22:5)+NH4 | 15.2 | 5.93 | 5.18 |
| 404 | TAG(54:6/FA22:6)+NH4 | 6.62 | 9.02 | 11.73 |
| 405 | TAG(54:7/FA16:1)+NH4 | 16.65 | 13.82 | 8.76 |
| 406 | TAG(54:7/FA18:1)+NH4 | 18.14 | 6.39 | 8.94 |
| 407 | TAG(54:7/FA18:2)+NH4 | 5.02 | 11.48 | 5.63 |
| 408 | TAG(54:7/FA18:3)+NH4 | 14.19 | 6.44 | 9.51 |
| 409 | TAG(54:7/FA20:4)+NH4 | 10.73 | 7.86 | 10.95 |
| 410 | TAG(54:7/FA20:5)+NH4 | 14.34 | 6.98 | 4.92 |
| 411 | TAG(54:7/FA22:5)+NH4 | 17.35 | 9.73 | 20.19 |
| 412 | TAG(54:7/FA22:6)+NH4 | 9.39 | 10.73 | 7.69 |
| 413 | TAG(54:8/FA18:2)+NH4 | 10.72 | 7.66 | 15.22 |
| 414 | TAG(54:8/FA18:3)+NH4 | 7.61 | 9.57 | 11.38 |
| 415 | TAG(54:8/FA20:4)+NH4 | 33.22 | 21.32 | 13.36 |
| 416 | TAG(54:8/FA20:5)+NH4 | 17.71 | 16.78 | 3.82 |
| 417 | TAG(54:8/FA22:6)+NH4 | 9.38 | 19.77 | 14.5 |
| 418 | TAG(55:1/FA16:0)+NH4 | 19.71 | 3.1 | 20.84 |
| 419 | TAG(55:1/FA18:1)+NH4 | 15.79 | 6.11 | 15.29 |
| 420 | TAG(55:2/FA18:1)+NH4 | 3.78 | 11.79 | 11.33 |
| 421 | TAG(55:2/FA18:2)+NH4 | 16.97 | 2.26 | 12.22 |
| 422 | TAG(55:3/FA18:1)+NH4 | 17.55 | 7.83 | 8.59 |
| 423 | TAG(55:3/FA18:2)+NH4 | 18.94 | 10.93 | 17.75 |
| 424 | TAG(55:4/FA18:1)+NH4 | 21.09 | 4 | 5.45 |
| 425 | TAG(55:4/FA18:2)+NH4 | 11.18 | 6.62 | 7.77 |
| 426 | TAG(55:5/FA18:1)+NH4 | 31.76 | 12.43 | 21.91 |
| 427 | TAG(55:5/FA18:2)+NH4 | 19.59 | 9.74 | 15.23 |
| 428 | TAG(55:5/FA20:4)+NH4 | 9.7 | 8.75 | 23.52 |
| 429 | TAG(55:7/FA22:6)+NH4 | 14.95 | 17.39 | 12.61 |
| 430 | TAG(56:10/FA18:2)+NH4 | 13.46 | 14.28 | 4.5 |
| 431 | TAG(56:1/FA16:0)+NH4 | 8.72 | 13.04 | NA |
| 432 | TAG(56:1/FA18:1)+NH4 | 12.3 | 7.91 | 20.7 |
| 433 | TAG(56:2/FA16:0)+NH4 | 10.52 | 14.49 | 12.5 |
| 434 | TAG(56:2/FA18:0)+NH4 | 10.41 | 3.19 | 4.13 |
| 435 | TAG(56:2/FA20:0)+NH4 | 15.94 | 15.74 | 15.01 |
| 436 | TAG(56:2/FA20:1)+NH4 | 11.55 | 20.37 | 3.54 |
| 437 | TAG(56:3/FA16:0)+NH4 | NA | 5.31 | 3.59 |
| 438 | TAG(56:3/FA18:0)+NH4 | 9.14 | 14.2 | 13.22 |
| 439 | TAG(56:3/FA18:1)+NH4 | 10.42 | 15.18 | 9.13 |
| 440 | TAG(56:3/FA18:2)+NH4 | 10.59 | 13.1 | 12.76 |
| 441 | TAG(56:3/FA20:0)+NH4 | 13.4 | 17.48 | 6.11 |
| 442 | TAG(56:3/FA20:1)+NH4 | 14.91 | 15.88 | 8.27 |
| 443 | TAG(56:3/FA20:2)+NH4 | 19.51 | 11.92 | 17.34 |

| | | | | |
|---|---|---|---|---|
| 444 | TAG(56:4/FA16:0)+NH4 | 21.45 | 9.02 | 8.08 |
| 445 | TAG(56:4/FA18:0)+NH4 | 15.25 | 11.28 | 8.12 |
| 446 | TAG(56:4/FA18:1)+NH4 | 12.83 | 11.9 | 15.68 |
| 447 | TAG(56:4/FA18:2)+NH4 | 14.49 | 7.31 | 11.39 |
| 448 | TAG(56:4/FA20:1)+NH4 | 12.12 | 10.37 | 10.78 |
| 449 | TAG(56:4/FA20:2)+NH4 | 5.72 | 10.35 | 8.95 |
| 450 | TAG(56:4/FA20:3)+NH4 | 9.1 | 12.64 | 13.09 |
| 451 | TAG(56:4/FA20:4)+NH4 | 2.08 | 11.82 | 8.87 |
| 452 | TAG(56:4/FA22:4)+NH4 | 23.46 | 10.29 | 15.34 |
| 453 | TAG(56:5/FA16:0)+NH4 | 15.22 | 10.06 | 10.19 |
| 454 | TAG(56:5/FA18:0)+NH4 | 13.17 | 11.59 | 9.31 |
| 455 | TAG(56:5/FA18:1)+NH4 | 17.7 | 10.41 | 3.84 |
| 456 | TAG(56:5/FA18:2)+NH4 | 14.45 | 6.61 | 5.64 |
| 457 | TAG(56:5/FA20:1)+NH4 | 10.89 | 9.61 | 10.76 |
| 458 | TAG(56:5/FA20:2)+NH4 | 5.4 | 5.71 | 13.54 |
| 459 | TAG(56:5/FA20:3)+NH4 | 6.37 | 14.24 | 18.25 |
| 460 | TAG(56:5/FA20:4)+NH4 | 13.85 | 15.6 | 6.21 |
| 461 | TAG(56:5/FA22:4)+NH4 | 12.95 | 10.41 | 6.61 |
| 462 | TAG(56:5/FA22:5)+NH4 | 16.48 | 21.88 | 19.49 |
| 463 | TAG(56:6/FA16:0)+NH4 | 8.74 | 15.07 | 4.28 |
| 464 | TAG(56:6/FA18:0)+NH4 | 21.59 | 9.3 | 15.47 |
| 465 | TAG(56:6/FA18:1)+NH4 | 6.95 | 16.58 | 9.39 |
| 466 | TAG(56:6/FA18:2)+NH4 | 8.91 | 10.12 | 6.74 |
| 467 | TAG(56:6/FA18:3)+NH4 | 12.96 | 24.04 | 9.56 |
| 468 | TAG(56:6/FA20:2)+NH4 | 14.61 | 5.51 | 10.96 |
| 469 | TAG(56:6/FA20:3)+NH4 | 10.82 | 10.23 | 5.5 |
| 470 | TAG(56:6/FA20:4)+NH4 | 13.11 | 10.98 | 9.24 |
| 471 | TAG(56:6/FA20:5)+NH4 | 9 | 10.36 | 16.53 |
| 472 | TAG(56:6/FA22:4)+NH4 | 16.28 | 9.27 | 8.09 |
| 473 | TAG(56:6/FA22:5)+NH4 | 12.14 | 14.15 | 7.31 |
| 474 | TAG(56:6/FA22:6)+NH4 | 10.17 | 13.39 | 11.07 |
| 475 | TAG(56:7/FA16:0)+NH4 | 14.12 | 16.19 | 8.01 |
| 476 | TAG(56:7/FA16:1)+NH4 | 25.23 | 20.17 | 27.37 |
| 477 | TAG(56:7/FA18:0)+NH4 | 15.55 | 8.63 | 14.45 |
| 478 | TAG(56:7/FA18:1)+NH4 | 13.31 | 9.7 | 12.43 |
| 479 | TAG(56:7/FA18:2)+NH4 | 11.15 | 11.62 | 3.61 |
| 480 | TAG(56:7/FA18:3)+NH4 | 7.53 | 20.63 | 17.26 |
| 481 | TAG(56:7/FA20:3)+NH4 | 7.23 | 13.12 | 9.34 |
| 482 | TAG(56:7/FA20:4)+NH4 | 7.52 | 11.24 | 9.1 |
| 483 | TAG(56:7/FA20:5)+NH4 | 17.83 | 10.38 | 20.15 |
| 484 | TAG(56:7/FA22:4)+NH4 | 18.48 | 12.67 | 11.22 |
| 485 | TAG(56:7/FA22:5)+NH4 | 12.99 | 12.94 | 8.62 |
| 486 | TAG(56:7/FA22:6)+NH4 | 8.79 | 28.47 | 10.52 |

| | | | | |
|---|---|---|---|---|
| 487 | TAG(56:8/FA16:0)+NH4 | 14.39 | 15.13 | 13.79 |
| 488 | TAG(56:8/FA16:1)+NH4 | 12.76 | 16.33 | NA |
| 489 | TAG(56:8/FA18:1)+NH4 | 9.41 | 17.76 | 12.12 |
| 490 | TAG(56:8/FA18:2)+NH4 | 12.85 | 12.54 | 6.76 |
| 491 | TAG(56:8/FA18:3)+NH4 | 10.63 | 19.77 | 15.51 |
| 492 | TAG(56:8/FA20:4)+NH4 | 15.81 | 17.68 | 11.18 |
| 493 | TAG(56:8/FA20:5)+NH4 | 15.09 | 8.53 | 10.93 |
| 494 | TAG(56:8/FA22:5)+NH4 | 12.46 | 24.89 | 15.91 |
| 495 | TAG(56:8/FA22:6)+NH4 | 9.94 | 25.08 | 6.55 |
| 496 | TAG(56:9/FA18:3)+NH4 | 20.32 | 7.41 | 17.91 |
| 497 | TAG(56:9/FA20:4)+NH4 | 10.86 | 24.55 | 20 |
| 498 | TAG(56:9/FA20:5)+NH4 | 7.26 | 5.36 | 16.22 |
| 499 | TAG(56:9/FA22:6)+NH4 | 14.57 | 25.67 | 10.11 |
| 500 | TAG(57:10/FA22:6)+NH4 | 76.58 | NA | 65.4 |
| 501 | TAG(57:2/FA18:1)+NH4 | 3.85 | 15.13 | 13.56 |
| 502 | TAG(57:3/FA18:2)+NH4 | 21.96 | 16.69 | 22.94 |
| 503 | TAG(58:10/FA18:2)+NH4 | 31.52 | 19.98 | 21.43 |
| 504 | TAG(58:10/FA20:4)+NH4 | 15.46 | 14.72 | 23.39 |
| 505 | TAG(58:10/FA20:5)+NH4 | 14.34 | 19.63 | 24.68 |
| 506 | TAG(58:10/FA22:5)+NH4 | 28.9 | 15.1 | 19.16 |
| 507 | TAG(58:10/FA22:6)+NH4 | 6.02 | 16.96 | 11.23 |
| 508 | TAG(58:2/FA18:1)+NH4 | 14.68 | 15.19 | 14.86 |
| 509 | TAG(58:3/FA18:1)+NH4 | NA | 11.66 | 10.86 |
| 510 | TAG(58:5/FA18:1)+NH4 | NA | 24.35 | 17.52 |
| 511 | TAG(58:6/FA16:0)+NH4 | 14.7 | 12.01 | 16.84 |
| 512 | TAG(58:6/FA18:0)+NH4 | 27.91 | 8.56 | 18.95 |
| 513 | TAG(58:6/FA18:1)+NH4 | 24.34 | 21.58 | 7.59 |
| 514 | TAG(58:6/FA20:4)+NH4 | 20.18 | 10.91 | 13.51 |
| 515 | TAG(58:6/FA22:4)+NH4 | 19.91 | 21.13 | 8.3 |
| 516 | TAG(58:6/FA22:5)+NH4 | 10.72 | 19.97 | 21.57 |
| 517 | TAG(58:7/FA16:0)+NH4 | 7.2 | 19.56 | 22.37 |
| 518 | TAG(58:7/FA18:0)+NH4 | 27.74 | 14.12 | 12.23 |
| 519 | TAG(58:7/FA18:1)+NH4 | 19.44 | 18.61 | 10.98 |
| 520 | TAG(58:7/FA18:2)+NH4 | 15.24 | 18.46 | 11.96 |
| 521 | TAG(58:7/FA20:4)+NH4 | 24.88 | 24.62 | 15.31 |
| 522 | TAG(58:7/FA22:4)+NH4 | 24.08 | 18.68 | 14.94 |
| 523 | TAG(58:7/FA22:5)+NH4 | 20.35 | 50 | 17.53 |
| 524 | TAG(58:7/FA22:6)+NH4 | 7.86 | 15.42 | 14.62 |
| 525 | TAG(58:8/FA18:1)+NH4 | 16.11 | 25.18 | 13.79 |
| 526 | TAG(58:8/FA18:2)+NH4 | 11.45 | 17.36 | 25.98 |
| 527 | TAG(58:8/FA20:3)+NH4 | 38.61 | 25.39 | 31.5 |
| 528 | TAG(58:8/FA20:4)+NH4 | 30.49 | 26.45 | 13.63 |
| 529 | TAG(58:8/FA22:5)+NH4 | 9.95 | 30.15 | 13.94 |

| | | | | |
|---|---|---|---|---|
| 530 | TAG(58:8/FA22:6)+NH4 | 7.31 | 28.57 | 8.31 |
| 531 | TAG(58:9/FA18:1)+NH4 | 9.56 | 22.64 | 17.14 |
| 532 | TAG(58:9/FA18:2)+NH4 | 19.98 | 36.25 | 15.59 |
| 533 | TAG(58:9/FA20:4)+NH4 | 19.97 | 21.59 | 4.95 |
| 534 | TAG(58:9/FA22:5)+NH4 | 5.69 | 21 | 24.5 |
| 535 | TAG(58:9/FA22:6)+NH4 | 6.89 | 24.95 | 7.07 |
| 536 | TAG(60:10/FA22:5)+NH4 | 32.52 | 40.04 | 18.41 |
| 537 | TAG(60:10/FA22:6)+NH4 | 25.32 | 38.12 | 43.9 |
| 538 | TAG(60:11/FA22:5)+NH4 | 59.32 | 36.51 | 34.76 |
| 539 | TAG(60:11/FA22:6)+NH4 | 21.85 | 26.82 | 14.54 |
| 540 | TAG(60:12/FA22:6)+NH4 | 60.12 | 4.8 | 52.38 |
| 541 | DAG(14:0/14:0)+NH4 | NA | NA | NA |
| 542 | DAG(14:0/16:1)+NH4 | NA | NA | NA |
| 543 | DAG(16:0/16:0)+NH4 | NA | NA | NA |
| 544 | DAG(16:0/16:1)+NH4 | NA | NA | NA |
| 545 | DAG(14:0/18:1)+NH4 | 36.5 | NA | NA |
| 546 | DAG(16:1/16:1)+NH4 | NA | 50.02 | NA |
| 547 | DAG(14:0/18:2)+NH4 | NA | NA | NA |
| 548 | DAG(14:0/18:3)+NH4 | 50.58 | 64.98 | NA |
| 549 | DAG(14:0/20:0)+NH4 | 5.87 | 55.77 | 48.74 |
| 550 | DAG(16:0/18:0)+NH4 | 44.12 | NA | 31.45 |
| 551 | DAG(16:1/18:0)+NH4 | NA | NA | NA |
| 552 | DAG(16:0/18:1)+NH4 | NA | 7.21 | NA |
| 553 | DAG(16:1/18:1)+NH4 | NA | NA | NA |
| 554 | DAG(16:0/18:2)+NH4 | NA | NA | NA |
| 555 | DAG(16:1/18:2)+NH4 | 43.96 | NA | NA |
| 556 | DAG(16:0/18:3)+NH4 | 27.33 | 85.02 | 49.67 |
| 557 | DAG(16:1/18:3)+NH4 | 40.96 | NA | NA |
| 558 | DAG(14:0/20:4)+NH4 | 73.88 | NA | 17.1 |
| 559 | DAG(16:1/20:0)+NH4 | NA | NA | NA |
| 560 | DAG(18:0/18:1)+NH4 | 74.49 | NA | NA |
| 561 | DAG(18:1/18:1)+NH4 | 32.11 | NA | 37.96 |
| 562 | DAG(18:0/18:2)+NH4 | NA | NA | NA |
| 563 | DAG(18:1/18:2)+NH4 | 77.08 | 15.25 | NA |
| 564 | DAG(18:0/18:3)+NH4 | NA | 63.56 | NA |
| 565 | DAG(16:1/20:2)+NH4 | NA | NA | NA |
| 566 | DAG(16:0/20:3)+NH4 | 51.77 | 86.99 | NA |
| 567 | DAG(16:0/20:4)+NH4 | NA | NA | 38.59 |
| 568 | DAG(18:2/18:3)+NH4 | 28.76 | 41.79 | NA |
| 569 | DAG(16:1/20:4)+NH4 | 48.15 | NA | 73.86 |
| 570 | DAG(16:0/20:5)+NH4 | NA | NA | NA |
| 571 | DAG(14:0/22:6)+NH4 | NA | NA | NA |
| 572 | DAG(18:1/20:1)+NH4 | 53.76 | NA | 12.64 |

| | | | | |
|---|---|---|---|---|
| 573 | DAG(18:1/20:2)+NH4 | 61.76 | 76.84 | 107.07 |
| 574 | DAG(18:1/20:3)+NH4 | 68.19 | NA | NA |
| 575 | DAG(18:2/20:3)+NH4 | NA | NA | 57.29 |
| 576 | DAG(18:1/20:4)+NH4 | NA | 66.92 | 40.38 |
| 577 | DAG(16:0/22:5)+NH4 | NA | NA | NA |
| 578 | DAG(18:2/20:4)+NH4 | 92.06 | 34.23 | 30.46 |
| 579 | DAG(18:1/20:5)+NH4 | 83.14 | 77.96 | 104.68 |
| 580 | DAG(16:0/22:6)+NH4 | 62.97 | NA | 55.04 |
| 581 | DAG(18:2/20:5)+NH4 | 50.96 | NA | 27.76 |
| 582 | DAG(16:1/22:6)+NH4 | NA | NA | NA |
| 583 | DAG(20:0/20:0)+NH4 | 11.41 | 68.74 | 5.38 |
| 584 | DAG(18:1/22:4)+NH4 | NA | NA | NA |
| 585 | DAG(18:2/22:4)+NH4 | NA | NA | NA |
| 586 | DAG(18:1/22:5)+NH4 | NA | NA | NA |
| 587 | DAG(18:0/22:6)+NH4 | NA | NA | NA |
| 588 | DAG(18:2/22:5)+NH4 | 45.51 | 63.57 | NA |
| 589 | DAG(18:1/22:6)+NH4 | 58.84 | 87.34 | 29.8 |
| 590 | DAG(18:2/22:6)+NH4 | NA | 85.21 | 45.24 |
| 591 | MAG(16:0)+NH4 | NA | NA | NA |
| 592 | MAG(16:1)+NH4 | NA | NA | NA |
| 593 | MAG(18:0)+NH4 | NA | NA | NA |
| 594 | MAG(18:1)+NH4 | NA | NA | NA |
| 595 | MAG(18:2)+NH4 | NA | NA | NA |
| 596 | MAG(20:0)+NH4 | NA | NA | NA |
| 597 | MAG(20:1)+NH4 | NA | NA | NA |
| 598 | MAG(20:2)+NH4 | NA | NA | NA |
| 599 | MAG(20:3)+NH4 | NA | NA | NA |
| 600 | MAG(20:4)+NH4 | NA | NA | NA |
| 601 | MAG(22:0)+NH4 | NA | NA | NA |
| 602 | MAG(22:1)+NH4 | NA | NA | NA |
| 603 | MAG(22:2)+NH4 | NA | NA | NA |
| 604 | MAG(22:3)+NH4 | NA | NA | NA |
| 605 | MAG(22:4)+NH4 | NA | NA | NA |
| 606 | MAG(22:5)+NH4 | NA | NA | NA |
| 607 | MAG(22:6)+NH4 | NA | 139.2 | NA |
| 608 | LPC(14:0)+AcO | 26.78 | 10.1 | 7.34 |
| 609 | LPC(16:0)+AcO | 15.99 | 6.62 | 4.9 |
| 610 | LPC(16:1)+AcO | 13.89 | 13.59 | 8.59 |
| 611 | LPC(18:0)+AcO | 13.23 | 11.26 | 13.41 |
| 612 | LPC(18:1)+AcO | 30.62 | 5.4 | 13.51 |
| 613 | LPC(18:2)+AcO | 38.72 | 6.5 | 20.21 |
| 614 | LPC(18:3)+AcO | 23.6 | 18.9 | 23.84 |
| 615 | LPC(20:0)+AcO | 57.71 | 21.83 | 24.59 |

| | | | | |
|---|---|---|---|---|
| 616 | LPC(20:1)+AcO | 32.83 | 5.39 | 10.82 |
| 617 | LPC(20:2)+AcO | 27.48 | 6.81 | 10.5 |
| 618 | LPC(20:3)+AcO | 8.27 | 8.16 | 15.24 |
| 619 | LPC(20:4)+AcO | 23.33 | 2.53 | 11.94 |
| 620 | LPC(20:5)+AcO | 20.04 | 33.38 | 59.96 |
| 621 | LPC(22:4)+AcO | 13.59 | 12.88 | 24.99 |
| 622 | LPC(22:5)+AcO | 73.78 | 16.51 | 23.8 |
| 623 | LPC(22:6)+AcO | 39.15 | 46.4 | 28.16 |
| 624 | PC(14:0/14:0)+AcO | 18.93 | 44.68 | 9.6 |
| 625 | PC(14:0/18:1)+AcO | 4.37 | 14.72 | 5.65 |
| 626 | PC(14:0/18:2)+AcO | 6.34 | 10.2 | 6.98 |
| 627 | PC(14:0/18:3)+AcO | 21.09 | 19.45 | 44.88 |
| 628 | PC(14:0/20:1)+AcO | NA | 49.24 | 35.98 |
| 629 | PC(14:0/20:2)+AcO | 21.11 | 23.96 | 31.17 |
| 630 | PC(14:0/20:3)+AcO | 12.46 | 14.1 | 31.96 |
| 631 | PC(14:0/20:4)+AcO | 9.99 | 18.24 | 20.23 |
| 632 | PC(14:0/20:5)+AcO | 13.2 | 77.72 | 11.64 |
| 633 | PC(14:0/22:4)+AcO | 21.85 | 65.1 | 27.93 |
| 634 | PC(14:0/22:5)+AcO | 35.43 | 59.16 | 35.39 |
| 635 | PC(14:0/22:6)+AcO | 17.88 | 31.36 | 30.77 |
| 636 | PC(14:1/14:1)+AcO | 42.86 | NA | 27.45 |
| 637 | PC(16:0/14:0)+AcO | 5.92 | 24.18 | 10.61 |
| 638 | PC(16:0/16:0)+AcO | 5.25 | 6.05 | 1.76 |
| 639 | PC(16:0/16:1)+AcO | 5.14 | 2.77 | 6.76 |
| 640 | PC(16:0/18:0)+AcO | 16.19 | 7.6 | 6.66 |
| 641 | PC(16:0/18:1)+AcO | 3.55 | 2.77 | 2.61 |
| 642 | PC(16:0/18:2)+AcO | 3.64 | 3.29 | 4.12 |
| 643 | PC(16:0/18:3)+AcO | 2.69 | 4.41 | 6.74 |
| 644 | PC(16:0/20:1)+AcO | 5.6 | 12.13 | 12.28 |
| 645 | PC(16:0/20:2)+AcO | 4.64 | 15.44 | 3.51 |
| 646 | PC(16:0/20:3)+AcO | 7.96 | 3.19 | 4.76 |
| 647 | PC(16:0/20:4)+AcO | 5.7 | 3.97 | 7.2 |
| 648 | PC(16:0/20:5)+AcO | 4.44 | 11.24 | 6.93 |
| 649 | PC(16:0/22:4)+AcO | 7.09 | 3.29 | 4.97 |
| 650 | PC(16:0/22:5)+AcO | 5.95 | 7.59 | 6.72 |
| 651 | PC(16:1/18:1)+AcO | 8.34 | 8.04 | 16.33 |
| 652 | PC(16:1/18:2)+AcO | 9.46 | 22.04 | 11.03 |
| 653 | PC(16:0/22:6)+AcO | 2.62 | 4.44 | 7.42 |
| 654 | PC(18:0/14:0)+AcO | 15.32 | 7.39 | 19.7 |
| 655 | PC(18:0/16:1)+AcO | 8.5 | 12.24 | 10.7 |
| 656 | PC(18:0/18:0)+AcO | 4.88 | 11.54 | 9.6 |
| 657 | PC(18:0/18:1)+AcO | 3.28 | 5.74 | 2.97 |
| 658 | PC(18:0/18:2)+AcO | 2.06 | 6.71 | 3.41 |

| | | | | |
|---|---|---|---|---|
| 659 | PC(18:0/18:3)+AcO | 2.65 | 5.05 | 11.02 |
| 660 | PC(18:0/20:0)+AcO | 15.47 | 46.76 | 30.88 |
| 661 | PC(18:0/20:1)+AcO | 10.16 | 10.06 | 25.62 |
| 662 | PC(18:0/20:2)+AcO | 4.96 | 8.04 | 10.68 |
| 663 | PC(18:0/20:3)+AcO | 5.41 | 12.67 | 6.06 |
| 664 | PC(18:0/20:4)+AcO | 7.07 | 6.39 | 2.57 |
| 665 | PC(18:0/20:5)+AcO | 12.02 | 5.52 | 7.93 |
| 666 | PC(18:0/22:4)+AcO | 8.5 | 5.01 | 2.12 |
| 667 | PC(18:0/22:5)+AcO | 11.26 | 6.72 | 5.87 |
| 668 | PC(18:0/22:6)+AcO | 9.39 | 11.46 | 12.68 |
| 669 | PC(18:1/16:1)+AcO | 4.82 | 5.75 | 12.4 |
| 670 | PC(18:1/18:1)+AcO | 7.26 | 7.71 | 3.9 |
| 671 | PC(18:1/18:2)+AcO | 3.25 | 7.64 | 7.75 |
| 672 | PC(18:1/18:3)+AcO | 8.95 | 16.16 | 7.87 |
| 673 | PC(18:1/20:1)+AcO | 23.01 | 17.2 | 20.84 |
| 674 | PC(18:1/20:2)+AcO | 7.81 | 9.77 | 7.85 |
| 675 | PC(18:1/20:3)+AcO | 5.84 | 15.19 | 9.31 |
| 676 | PC(18:1/20:4)+AcO | 7.99 | 13.26 | 6.86 |
| 677 | PC(18:1/20:5)+AcO | 14.97 | 15.63 | 10.43 |
| 678 | PC(18:1/22:4)+AcO | 13.03 | 22.36 | 25.8 |
| 679 | PC(18:1/22:5)+AcO | 12.71 | 14.11 | 7 |
| 680 | PC(18:1/22:6)+AcO | 14.46 | 14.79 | 20.28 |
| 681 | PC(18:2/16:1)+AcO | 3.4 | 14.02 | 19.98 |
| 682 | PC(18:2/18:2)+AcO | 7.5 | 6.05 | 4.69 |
| 683 | PC(18:2/18:3)+AcO | 12.2 | 61.42 | 21.24 |
| 684 | PC(18:2/20:1)+AcO | 12.9 | 12.71 | 5.2 |
| 685 | PC(18:2/20:2)+AcO | 9.44 | 8.1 | 12.11 |
| 686 | PC(18:2/20:3)+AcO | 10.04 | 23.03 | 8.71 |
| 687 | PC(18:2/20:4)+AcO | 15.49 | 12.95 | 10.65 |
| 688 | PC(18:2/20:5)+AcO | 9.01 | 29.22 | 34.51 |
| 689 | PC(18:2/22:4)+AcO | 22.78 | 13.83 | 22.4 |
| 690 | PC(18:2/22:5)+AcO | 25.81 | 15.18 | 24.55 |
| 691 | PC(18:2/22:6)+AcO | 20.57 | 11.05 | 18.36 |
| 692 | PC(20:0/16:1)+AcO | 49.99 | 2.58 | 11.72 |
| 693 | PC(20:0/18:1)+AcO | 5.94 | 14.31 | 30.11 |
| 694 | PC(20:0/18:3)+AcO | 53.44 | 41.33 | 60.86 |
| 695 | PC(20:0/20:1)+AcO | 7 | 66.45 | 108.07 |
| 696 | PC(20:0/20:2)+AcO | 31.17 | 22.71 | NA |
| 697 | PC(20:0/20:3)+AcO | 20.36 | 15.27 | 15.71 |
| 698 | PC(20:0/20:4)+AcO | 17.9 | 17.85 | 13.52 |
| 699 | PC(20:0/20:5)+AcO | 48.22 | 75 | 27.65 |
| 700 | PC(20:0/22:4)+AcO | 68.64 | 35.22 | 58.4 |
| 701 | PC(20:0/22:5)+AcO | NA | 31.96 | 46.4 |

| | | | | |
|---|---|---|---|---|
| 702 | PC(20:0/22:6)+AcO | 35.5 | 35.72 | 30.86 |
| 703 | LPE(14:0)-H | NA | 114.32 | NA |
| 704 | LPE(16:0)-H | 91.12 | 2.69 | 12.11 |
| 705 | LPE(16:1)-H | NA | 13.82 | 41.41 |
| 706 | LPE(18:0)-H | 23.25 | 2.82 | 15.89 |
| 707 | LPE(18:1)-H | 21.19 | 2.85 | 8 |
| 708 | LPE(18:2)-H | 36.16 | 1.68 | 9.75 |
| 709 | LPE(18:3)-H | 87.67 | 6.01 | 7.4 |
| 710 | LPE(20:0)-H | 44.51 | 19.74 | 40.63 |
| 711 | LPE(20:1)-H | 10.15 | 30.59 | 19.91 |
| 712 | LPE(20:2)-H | 34.55 | 9.42 | 21.24 |
| 713 | LPE(20:3)-H | 23.23 | 16.02 | 12.7 |
| 714 | LPE(20:4)-H | 22.55 | 3.3 | 14.06 |
| 715 | LPE(20:5)-H | 64.36 | 21.6 | 33.65 |
| 716 | LPE(22:4)-H | 15.37 | 15.52 | 14.59 |
| 717 | LPE(22:5)-H | 32.32 | 16.74 | 18.96 |
| 718 | LPE(22:6)-H | 43.63 | 23.05 | 10.9 |
| 719 | PE(14:0/14:0)-H | 37.33 | 11.72 | 53.33 |
| 720 | PE(14:0/16:1)-H | NA | NA | 39.35 |
| 721 | PE(14:0/18:1)-H | 21 | 36.8 | 33.34 |
| 722 | PE(14:0/18:2)-H | 9.99 | 30.23 | 20.48 |
| 723 | PE(14:0/18:3)-H | 44.91 | NA | 63.2 |
| 724 | PE(14:0/20:1)-H | 74.22 | NA | 73.72 |
| 725 | PE(14:0/20:2)-H | 44.68 | NA | 98.62 |
| 726 | PE(14:0/20:3)-H | 22.53 | 14.62 | 40.58 |
| 727 | PE(14:0/20:4)-H | 39.34 | 68.22 | 61.32 |
| 728 | PE(14:0/20:5)-H | NA | NA | 4.5 |
| 729 | PE(14:0/22:4)-H | 58.26 | 59.05 | 77.56 |
| 730 | PE(14:0/22:5)-H | 57.93 | 40.11 | 49.97 |
| 731 | PE(14:0/22:6)-H | 53.39 | 26.95 | 56.51 |
| 732 | PE(14:1/14:1)-H | NA | NA | NA |
| 733 | PE(16:0/14:0)-H | 15.63 | 23.43 | 16.35 |
| 734 | PE(16:0/16:0)-H | 32.05 | 12.81 | 17.85 |
| 735 | PE(16:0/16:1)-H | 10.51 | NA | 10.7 |
| 736 | PE(16:0/18:1)-H | 8.68 | 8.93 | 3.24 |
| 737 | PE(16:0/18:2)-H | 5.58 | 11.66 | 4.29 |
| 738 | PE(16:0/18:3)-H | 11.17 | 5.83 | 10.34 |
| 739 | PE(16:0/20:1)-H | 23.98 | 41.79 | 20.34 |
| 740 | PE(16:0/20:2)-H | 8.16 | 29.1 | 7.73 |
| 741 | PE(16:0/20:3)-H | 5.12 | 17.09 | 10.84 |
| 742 | PE(16:0/20:4)-H | 8.53 | 13.8 | 5.47 |
| 743 | PE(16:0/20:5)-H | 19.36 | 37.94 | 5.61 |
| 744 | PE(16:0/22:4)-H | 9.96 | 12.7 | 9.44 |

| | | | | |
|---|---|---|---|---|
| 745 | PE(16:0/22:5)-H | 8 | 9.27 | 5.58 |
| 746 | PE(16:0/22:6)-H | 4.13 | 39.18 | 13.69 |
| 747 | PE(18:0/14:0)-H | 17.39 | 10.38 | 39.37 |
| 748 | PE(18:0/16:0)-H | 15.15 | 25.71 | 23.53 |
| 749 | PE(18:0/16:1)-H | 26.77 | 26.18 | 13.87 |
| 750 | PE(18:0/18:0)-H | 9.32 | 9.53 | 6 |
| 751 | PE(18:0/18:1)-H | 9.49 | 5.79 | 8.86 |
| 752 | PE(18:0/18:2)-H | 6.39 | 27.35 | 9.57 |
| 753 | PE(18:0/18:3)-H | 9.13 | 42.05 | 12.55 |
| 754 | PE(18:0/20:1)-H | 11.12 | 12.59 | 28.75 |
| 755 | PE(18:0/20:2)-H | 10.53 | 11.15 | 37.09 |
| 756 | PE(18:0/20:3)-H | 26.1 | 10.74 | 24.81 |
| 757 | PE(18:0/20:4)-H | 13.09 | 11.23 | 31.69 |
| 758 | PE(18:0/20:5)-H | 21.14 | 35.23 | 13.4 |
| 759 | PE(18:0/22:4)-H | 14.05 | 23.54 | 41.21 |
| 760 | PE(18:0/22:5)-H | 14.17 | 14.16 | 24.14 |
| 761 | PE(18:0/22:6)-H | 16.97 | 49.18 | 14.03 |
| 762 | PE(18:1/16:1)-H | 4.52 | NA | 13.43 |
| 763 | PE(18:1/18:1)-H | 7.42 | 10.18 | 4.7 |
| 764 | PE(18:1/18:2)-H | 10.27 | 14.61 | 3.49 |
| 765 | PE(18:1/18:3)-H | 11.61 | 6.89 | 9.3 |
| 766 | PE(18:1/20:1)-H | 13.59 | 13.59 | 25.45 |
| 767 | PE(18:1/20:2)-H | 14.66 | 32.99 | 9.8 |
| 768 | PE(18:1/20:3)-H | 12.49 | 22.62 | 18.53 |
| 769 | PE(18:1/20:4)-H | 6.09 | 17.32 | 18.43 |
| 770 | PE(18:1/20:5)-H | 26.9 | 26 | 30.94 |
| 771 | PE(18:1/22:4)-H | 10.78 | 22.02 | 33.09 |
| 772 | PE(18:1/22:5)-H | 14.25 | 7.26 | 30.94 |
| 773 | PE(18:1/22:6)-H | 6.7 | 15.97 | 17.66 |
| 774 | PE(18:2/16:1)-H | 6.83 | 22.98 | 16.43 |
| 775 | PE(18:2/18:2)-H | 21.8 | 13.06 | 4.59 |
| 776 | PE(18:2/18:3)-H | 13.89 | 17.58 | 14.23 |
| 777 | PE(18:2/20:1)-H | 14.82 | 10.17 | 13.12 |
| 778 | PE(18:2/20:2)-H | 8.78 | 42.16 | 31.17 |
| 779 | PE(18:2/20:3)-H | 14.89 | 26.34 | 20.98 |
| 780 | PE(18:2/20:4)-H | 15.24 | 18.43 | 23.7 |
| 781 | PE(18:2/20:5)-H | 32.86 | 65.28 | 65.99 |
| 782 | PE(18:2/22:4)-H | 45.48 | 51.31 | 46.64 |
| 783 | PE(18:2/22:5)-H | 27.49 | 13.34 | 38.07 |
| 784 | PE(18:2/22:6)-H | 51.49 | 47.83 | 38.34 |
| 785 | PE(O-16:0/16:0)-H | NA | 21.03 | 26.43 |
| 786 | PE(O-16:0/16:1)-H | NA | 44.8 | 26.14 |
| 787 | PE(O-16:0/18:0)-H | 16.86 | 25.91 | 40.63 |

| | | | | |
|---|---|---|---|---|
| 788 | PE(O-16:0/18:1)-H | 8.72 | 19.04 | 11.98 |
| 789 | PE(O-16:0/18:2)-H | 6.02 | 6.63 | 9.94 |
| 790 | PE(O-16:0/18:3)-H | 11.36 | 31.48 | 31.44 |
| 791 | PE(O-16:0/20:1)-H | 22.11 | 58.23 | 36.96 |
| 792 | PE(O-16:0/20:2)-H | 22.37 | 31.61 | 28.86 |
| 793 | PE(O-16:0/20:3)-H | 13.66 | 6.64 | 14.61 |
| 794 | PE(O-16:0/20:4)-H | 6.66 | 6.66 | 12.9 |
| 795 | PE(O-16:0/20:5)-H | 13.19 | 18.23 | 10.26 |
| 796 | PE(O-16:0/22:4)-H | 7.31 | 19.37 | 5.62 |
| 797 | PE(O-16:0/22:5)-H | 10.77 | 17.93 | 7.93 |
| 798 | PE(O-16:0/22:6)-H | 12.61 | 12.48 | 14.8 |
| 799 | PE(O-18:0/16:0)-H | 16.26 | 6.91 | 12.79 |
| 800 | PE(O-18:0/16:1)-H | 14.17 | 9.32 | 21.41 |
| 801 | PE(O-18:0/18:0)-H | 17.14 | 37.02 | 37.39 |
| 802 | PE(O-18:0/18:1)-H | 2.54 | 10.85 | 7.75 |
| 803 | PE(O-18:0/18:2)-H | 10.69 | 23.01 | 25.29 |
| 804 | PE(O-18:0/18:3)-H | 25.83 | 30.37 | 49.87 |
| 805 | PE(O-18:0/20:1)-H | 26.69 | 16.63 | 34.96 |
| 806 | PE(O-18:0/20:2)-H | 12.74 | 12.86 | 18.81 |
| 807 | PE(O-18:0/20:3)-H | 13.15 | 12.93 | 6.59 |
| 808 | PE(O-18:0/20:4)-H | 5.72 | 18.83 | 13.11 |
| 809 | PE(O-18:0/20:5)-H | 12.27 | 15.04 | 24.56 |
| 810 | PE(O-18:0/22:4)-H | 12.78 | 19.56 | 8.46 |
| 811 | PE(O-18:0/22:5)-H | 3.94 | 24.03 | 12.92 |
| 812 | PE(O-18:0/22:6)-H | 9.28 | 32.5 | 13.97 |
| 813 | PE(P-14:0/18:0)-H | 18.62 | NA | 60.72 |
| 814 | PE(P-14:0/18:1)-H | 17.3 | 29.53 | 26.15 |
| 815 | PE(P-16:0/16:0)-H | 6.68 | 20.27 | 8.46 |
| 816 | PE(P-16:0/16:1)-H | 10.71 | 20.58 | 13 |
| 817 | PE(P-16:0/18:0)-H | 15.68 | 30.37 | 10.3 |
| 818 | PE(P-16:0/18:1)-H | 10.13 | 8.54 | 5.9 |
| 819 | PE(P-16:0/18:2)-H | 3.51 | 5.62 | 6.13 |
| 820 | PE(P-16:0/18:3)-H | 11.16 | 35.58 | 13.75 |
| 821 | PE(P-16:0/20:1)-H | 9.63 | 13.22 | 3.27 |
| 822 | PE(P-16:0/20:2)-H | 18.75 | 12.13 | 14.8 |
| 823 | PE(P-16:0/20:3)-H | 11.1 | 6.86 | 7.11 |
| 824 | PE(P-16:0/20:4)-H | 9.18 | 12.58 | 7.17 |
| 825 | PE(P-16:0/20:5)-H | 23.83 | 21.63 | 20.51 |
| 826 | PE(P-16:0/22:4)-H | 9.83 | 17.8 | 8.75 |
| 827 | PE(P-16:0/22:5)-H | 12.3 | 19.51 | 13.33 |
| 828 | PE(P-16:0/22:6)-H | 7.32 | 12.7 | 7.86 |
| 829 | PE(P-16:1/18:1)-H | 18.37 | 32.75 | 27.89 |
| 830 | PE(P-18:0/16:0)-H | 8.24 | 8.81 | 22.99 |

| | | | | |
|---|---|---|---|---|
| 831 | PE(P-18:0/16:1)-H | 16.21 | 22.15 | 30.31 |
| 832 | PE(P-18:0/18:0)-H | 11.9 | 18.51 | 15.06 |
| 833 | PE(P-18:0/18:1)-H | 6.77 | 8.27 | 7.31 |
| 834 | PE(P-18:0/18:2)-H | 4.46 | 11.15 | 10.72 |
| 835 | PE(P-18:0/18:3)-H | 5.7 | 18.85 | 14.16 |
| 836 | PE(P-18:0/20:1)-H | 11.54 | 29.81 | 28.07 |
| 837 | PE(P-18:0/20:2)-H | 18.61 | 20.74 | 19.77 |
| 838 | PE(P-18:0/20:3)-H | 6.06 | 14.11 | 9.73 |
| 839 | PE(P-18:0/20:4)-H | 11.05 | 12.94 | 5.56 |
| 840 | PE(P-18:0/20:5)-H | 10.31 | 22.82 | 7.69 |
| 841 | PE(P-18:0/22:4)-H | 8.34 | 16.27 | 7.41 |
| 842 | PE(P-18:0/22:5)-H | 9.06 | 23.6 | 6.64 |
| 843 | PE(P-18:0/22:6)-H | 12.99 | 9.93 | 7.93 |
| 844 | PE(P-18:1/16:0)-H | 6.36 | 7.57 | 11.1 |
| 845 | PE(P-18:1/16:1)-H | 20.19 | 27.2 | 39.88 |
| 846 | PE(P-18:1/18:0)-H | 7.18 | 23.37 | 19.88 |
| 847 | PE(P-18:1/18:1)-H | 9.97 | 5.86 | 4.58 |
| 848 | PE(P-18:1/18:2)-H | 7.6 | 11.87 | 8.95 |
| 849 | PE(P-18:1/18:3)-H | NA | 16.58 | 26.31 |
| 850 | PE(P-18:1/20:1)-H | 13.27 | 22.95 | 14.4 |
| 851 | PE(P-18:1/20:2)-H | 12.23 | 28.21 | 20.75 |
| 852 | PE(P-18:1/20:3)-H | 8.58 | 17.8 | 5.96 |
| 853 | PE(P-18:1/20:4)-H | 10.37 | 17.29 | 8.15 |
| 854 | PE(P-18:1/20:5)-H | 16.59 | 24.31 | 25.21 |
| 855 | PE(P-18:1/22:4)-H | 8.45 | 22.05 | 4.76 |
| 856 | PE(P-18:1/22:5)-H | 4.33 | 11.61 | 9.4 |
| 857 | PE(P-18:1/22:6)-H | 6.43 | 19.36 | 14.06 |
| 858 | PE(P-18:2/18:2)-H | 8.34 | 7.96 | 8.88 |
| 859 | PE(P-18:2/20:4)-H | 9.62 | 19.34 | 8.32 |
| 860 | PE(P-18:2/22:6)-H | 29.58 | 31.11 | 23.41 |
| 861 | LPG(14:0)-H | NA | NA | NA |
| 862 | LPG(16:0)-H | 26.33 | 13.75 | 40.08 |
| 863 | LPG(16:1)-H | NA | 28.9 | NA |
| 864 | LPG(18:0)-H | 35.91 | 9.36 | 50.48 |
| 865 | LPG(18:1)-H | 20.74 | 5.65 | 16.2 |
| 866 | LPG(18:2)-H | 46.3 | 13.07 | 26.59 |
| 867 | LPG(18:3)-H | NA | 67.84 | NA |
| 868 | LPG(20:0)-H | 12.25 | NA | NA |
| 869 | LPG(20:1)-H | NA | 29.95 | 48.32 |
| 870 | LPG(20:2)-H | 103.03 | 47.49 | 85.73 |
| 871 | LPG(20:3)-H | NA | NA | NA |
| 872 | LPG(20:4)-H | NA | NA | 47.82 |
| 873 | LPG(20:5)-H | NA | NA | 68.93 |

| | | | | |
|---|---|---|---|---|
| 874 | LPG(22:4)-H | 19.04 | NA | NA |
| 875 | LPG(22:5)-H | 69.02 | NA | NA |
| 876 | LPG(22:6)-H | NA | NA | NA |
| 877 | PG(14:0/14:0)-H | 24.84 | 33.48 | 20.17 |
| 878 | PG(14:1/14:1)-H | NA | NA | NA |
| 879 | PG(14:0/18:1)-H | NA | 38.69 | 28.96 |
| 880 | PG(14:0/18:2)-H | 21.48 | 17.29 | 31.4 |
| 881 | PG(14:0/18:3)-H | 39.31 | NA | NA |
| 882 | PG(14:0/20:1)-H | 37.22 | NA | 74.22 |
| 883 | PG(14:0/20:2)-H | NA | NA | NA |
| 884 | PG(14:0/20:3)-H | 42.18 | NA | 31.2 |
| 885 | PG(14:0/20:4)-H | 37.42 | 20.7 | 26.18 |
| 886 | PG(14:0/20:5)-H | 17.09 | NA | 41.16 |
| 887 | PG(14:0/22:4)-H | 39.7 | NA | 53.89 |
| 888 | PG(14:0/22:5)-H | 38.88 | 28.82 | 37.57 |
| 889 | PG(14:0/22:6)-H | 30.74 | NA | 14.65 |
| 890 | PG(16:0/14:0)-H | 12.13 | NA | 20.32 |
| 891 | PG(16:0/16:0)-H | 16.38 | 34.12 | 15.27 |
| 892 | PG(16:0/16:1)-H | 23.18 | NA | NA |
| 893 | PG(16:0/18:0)-H | 21.37 | 49.19 | 42.26 |
| 894 | PG(16:0/18:1)-H | 6.35 | 21.71 | 12.11 |
| 895 | PG(16:0/18:2)-H | 12.96 | 32.45 | 22.54 |
| 896 | PG(16:0/18:3)-H | 29.48 | 38.49 | 45.86 |
| 897 | PG(16:0/20:1)-H | 23.31 | 61.58 | NA |
| 898 | PG(16:0/20:2)-H | 11.49 | 37.04 | 9.4 |
| 899 | PG(16:0/20:3)-H | 34.58 | 55.08 | 34.72 |
| 900 | PG(16:0/20:4)-H | 12.7 | 34.48 | 9.26 |
| 901 | PG(16:0/20:5)-H | 34.1 | 36.72 | 53.25 |
| 902 | PG(16:0/22:4)-H | NA | NA | 46.61 |
| 903 | PG(16:0/22:5)-H | 28.24 | 55.19 | 42.06 |
| 904 | PG(16:0/22:6)-H | 18.5 | NA | 19.43 |
| 905 | PG(18:0/14:0)-H | 27.6 | 33.36 | NA |
| 906 | PG(18:0/16:1)-H | 11.98 | 40.73 | NA |
| 907 | PG(18:0/18:0)-H | 15.96 | 25.93 | 29.01 |
| 908 | PG(18:0/18:1)-H | 10.71 | 23.44 | 10.68 |
| 909 | PG(18:0/18:2)-H | 13 | 21.84 | 7.58 |
| 910 | PG(18:0/18:3)-H | 47.77 | 19.24 | 65.88 |
| 911 | PG(18:0/20:0)-H | 20.22 | 51.49 | 24.02 |
| 912 | PG(18:0/20:1)-H | 19.98 | 56.29 | 39.95 |
| 913 | PG(18:0/20:2)-H | 12.92 | 32.53 | 18.25 |
| 914 | PG(18:0/20:3)-H | 40.08 | 38.4 | 45.85 |
| 915 | PG(18:0/20:4)-H | 22.11 | 28.38 | 12.61 |
| 916 | PG(18:0/20:5)-H | NA | 33.21 | 18.91 |

| | | | | |
|---|---|---|---|---|
| 917 | PG(18:0/22:4)-H | NA | NA | NA |
| 918 | PG(18:0/22:5)-H | 87.52 | NA | NA |
| 919 | PG(18:0/22:6)-H | 116.07 | 36.43 | 63.31 |
| 920 | PG(18:1/16:1)-H | 16.98 | 21.04 | 10.3 |
| 921 | PG(18:1/18:1)-H | 10.41 | 9.94 | 23.64 |
| 922 | PG(18:1/18:2)-H | 16.27 | 32.15 | 19.39 |
| 923 | PG(18:1/18:3)-H | 22.82 | 27.26 | 25.16 |
| 924 | PG(18:1/20:1)-H | 19.43 | 39.94 | 46.48 |
| 925 | PG(18:1/20:2)-H | 26.16 | 14.8 | 16.1 |
| 926 | PG(18:1/20:3)-H | 6.04 | 15.24 | 18.86 |
| 927 | PG(18:1/20:4)-H | 26.69 | 24.18 | 26.32 |
| 928 | PG(18:1/20:5)-H | 13.53 | 54.56 | 33.66 |
| 929 | PG(18:1/22:4)-H | 26.77 | 35.03 | 18.61 |
| 930 | PG(18:1/22:5)-H | 9 | 20.86 | 16.64 |
| 931 | PG(18:1/22:6)-H | 52.24 | 39.17 | 31.25 |
| 932 | PG(18:2/16:1)-H | 3.96 | 20.07 | 8.23 |
| 933 | PG(18:2/18:2)-H | 21.68 | 19.97 | 19.23 |
| 934 | PG(18:2/18:3)-H | 28.48 | 32.49 | 30.53 |
| 935 | PG(18:2/20:1)-H | 38.32 | 65.34 | 23 |
| 936 | PG(18:2/20:2)-H | 22.67 | 19.79 | 36.63 |
| 937 | PG(18:2/20:3)-H | 14.52 | 22.3 | 26.76 |
| 938 | PG(18:2/20:4)-H | 15.69 | 14.27 | 5.27 |
| 939 | PG(18:2/20:5)-H | 61.77 | 58.53 | 32.66 |
| 940 | PG(18:2/22:4)-H | 32.07 | 56.14 | 63.99 |
| 941 | PG(18:2/22:5)-H | 37.13 | 41.89 | 33.2 |
| 942 | PG(18:2/22:6)-H | 40.18 | 15.89 | 33.4 |
| 943 | PG(20:0/16:1)-H | 14.59 | 50.88 | 31.88 |
| 944 | PG(20:0/18:1)-H | 1.09 | 20.81 | 11.32 |
| 945 | PG(20:0/18:2)-H | 4.34 | 36.77 | 20.39 |
| 946 | PG(20:0/18:3)-H | NA | NA | 93.41 |
| 947 | PG(20:0/20:1)-H | NA | NA | 40.61 |
| 948 | PG(20:0/20:2)-H | 49.96 | 22.17 | 48.49 |
| 949 | PG(20:0/20:3)-H | 20.89 | 19.05 | 10.69 |
| 950 | PG(20:0/20:4)-H | 14.37 | 18.41 | 9.41 |
| 951 | PG(20:0/20:5)-H | 42.92 | 41.71 | 26.03 |
| 952 | PG(20:0/22:4)-H | 10.92 | 10.42 | 7.14 |
| 953 | PG(20:0/22:5)-H | 19.34 | 41.94 | 11.57 |
| 954 | PG(20:0/22:6)-H | 11.01 | 39.98 | 22.99 |
| 955 | LPI(14:0)-H | NA | NA | NA |
| 956 | LPI(16:0)-H | NA | NA | NA |
| 957 | LPI(16:1)-H | NA | NA | 52.3 |
| 958 | LPI(18:0)-H | NA | 12.73 | 46.34 |
| 959 | LPI(18:1)-H | NA | NA | NA |

| | | | | |
|---|---|---|---|---|
| 960 | LPI(18:2)-H | NA | NA | 66.4 |
| 961 | LPI(18:3)-H | NA | NA | 47.16 |
| 962 | LPI(20:0)-H | NA | 66.9 | NA |
| 963 | LPI(20:1)-H | NA | NA | NA |
| 964 | LPI(20:2)-H | NA | NA | NA |
| 965 | LPI(20:3)-H | NA | NA | 98.82 |
| 966 | LPI(20:4)-H | NA | NA | NA |
| 967 | LPI(20:5)-H | NA | NA | NA |
| 968 | LPI(22:4)-H | NA | NA | 69.65 |
| 969 | LPI(22:5)-H | NA | NA | NA |
| 970 | LPI(22:6)-H | NA | NA | NA |
| 971 | PI(14:1/14:1)-H | NA | NA | NA |
| 972 | PI(14:0/14:0)-H | 61.18 | NA | 48.53 |
| 973 | PI(14:0/18:1)-H | NA | 25.01 | 13.86 |
| 974 | PI(14:0/18:2)-H | 10.55 | 9.31 | 8.38 |
| 975 | PI(14:0/18:3)-H | NA | NA | 24.26 |
| 976 | PI(14:0/20:1)-H | 62.01 | NA | 34.63 |
| 977 | PI(14:0/20:2)-H | 44.01 | NA | 92.93 |
| 978 | PI(14:0/20:3)-H | 15.17 | 27.24 | 28.7 |
| 979 | PI(14:0/20:4)-H | 12.98 | 9.56 | 6.96 |
| 980 | PI(14:0/20:5)-H | NA | NA | NA |
| 981 | PI(14:0/22:4)-H | 26.34 | 14.08 | 25.74 |
| 982 | PI(14:0/22:5)-H | 31.1 | NA | 46.43 |
| 983 | PI(14:0/22:6)-H | 14.2 | NA | 17.55 |
| 984 | PI(16:0/14:0)-H | 74.4 | NA | NA |
| 985 | PI(16:0/16:0)-H | 18.52 | 18.27 | 26.17 |
| 986 | PI(16:0/16:1)-H | 32.02 | 9.85 | 15.02 |
| 987 | PI(16:0/18:0)-H | 20.97 | 51.6 | 12.41 |
| 988 | PI(16:0/18:1)-H | 18.18 | 13.7 | 5.3 |
| 989 | PI(16:0/18:2)-H | 14.15 | 8.59 | 8.93 |
| 990 | PI(16:0/18:3)-H | 34.83 | 23.68 | 5.14 |
| 991 | PI(16:0/20:1)-H | 13.97 | NA | 55.32 |
| 992 | PI(16:0/20:2)-H | 38.84 | 29.83 | 40.87 |
| 993 | PI(16:0/20:3)-H | NA | 18.41 | 14.12 |
| 994 | PI(16:0/20:4)-H | 19.16 | 11.19 | 9.89 |
| 995 | PI(16:0/20:5)-H | 39.48 | NA | 40.27 |
| 996 | PI(16:0/22:4)-H | 33.94 | 26.28 | 64 |
| 997 | PI(16:0/22:5)-H | 16.4 | 19.33 | 18.06 |
| 998 | PI(16:0/22:6)-H | NA | NA | NA |
| 999 | PI(18:0/14:0)-H | NA | NA | 75.67 |
| 1000 | PI(18:0/16:1)-H | 37.73 | 19.57 | 15.42 |
| 1001 | PI(18:0/18:0)-H | 11.65 | 12.43 | 58.3 |
| 1002 | PI(18:0/18:1)-H | 14.31 | 17.44 | 13.01 |

| | | | | |
|---|---|---|---|---|
| 1003 | PI(18:0/18:2)-H | 7.94 | 3.74 | 10.02 |
| 1004 | PI(18:0/18:3)-H | 26.74 | NA | 27.11 |
| 1005 | PI(18:0/20:0)-H | 11.31 | 10.3 | 6.95 |
| 1006 | PI(18:0/20:1)-H | NA | 23.93 | 65.46 |
| 1007 | PI(18:0/20:2)-H | 39.94 | 10.73 | 18.31 |
| 1008 | PI(18:0/20:3)-H | 13.41 | 17.26 | 12.91 |
| 1009 | PI(18:0/20:4)-H | 10.23 | 5.93 | 5.68 |
| 1010 | PI(18:0/20:5)-H | 20.96 | NA | 27.08 |
| 1011 | PI(18:0/22:4)-H | 54.67 | 17.66 | 43.66 |
| 1012 | PI(18:0/22:5)-H | 30.72 | 18.78 | 17.58 |
| 1013 | PI(18:0/22:6)-H | 22.88 | 25.99 | 25.96 |
| 1014 | PI(18:1/16:1)-H | NA | 25.38 | 25.41 |
| 1015 | PI(18:1/18:1)-H | 11.5 | 16.6 | 15.68 |
| 1016 | PI(18:1/18:2)-H | 14.99 | 6.84 | 9.2 |
| 1017 | PI(18:1/18:3)-H | 32.56 | NA | 49.81 |
| 1018 | PI(18:1/20:1)-H | NA | 30.42 | 41.91 |
| 1019 | PI(18:1/20:2)-H | 22.89 | 37.86 | 22.8 |
| 1020 | PI(18:1/20:3)-H | 37.01 | 55.99 | 25.76 |
| 1021 | PI(18:1/20:4)-H | NA | 50.28 | 25.41 |
| 1022 | PI(18:1/20:5)-H | 82.93 | NA | NA |
| 1023 | PI(18:1/22:4)-H | 65.4 | NA | 44.74 |
| 1024 | PI(18:1/22:5)-H | NA | NA | 49.16 |
| 1025 | PI(18:1/22:6)-H | 148.77 | NA | 52.51 |
| 1026 | PI(18:2/16:1)-H | NA | 2.63 | 7.12 |
| 1027 | PI(18:2/18:2)-H | 25.6 | 38.26 | 14.94 |
| 1028 | PI(18:2/18:3)-H | NA | NA | 15.57 |
| 1029 | PI(18:2/20:1)-H | 38.11 | NA | 42.24 |
| 1030 | PI(18:2/20:2)-H | 36.75 | NA | NA |
| 1031 | PI(18:2/20:3)-H | 44.49 | 50.6 | 28.11 |
| 1032 | PI(18:2/20:4)-H | 23.52 | 8.28 | 9.34 |
| 1033 | PI(18:2/20:5)-H | NA | 93.25 | NA |
| 1034 | PI(18:2/22:4)-H | 87.37 | 23.79 | 40.2 |
| 1035 | PI(18:2/22:5)-H | NA | NA | 65.36 |
| 1036 | PI(18:2/22:6)-H | NA | NA | 51.84 |
| 1037 | PI(20:0/16:1)-H | 48.14 | 39.14 | 30.94 |
| 1038 | PI(20:0/18:1)-H | 16.95 | 20.75 | 22.81 |
| 1039 | PI(20:0/18:2)-H | 12.85 | 27.19 | 18.48 |
| 1040 | PI(20:0/18:3)-H | NA | NA | 30.74 |
| 1041 | PI(20:0/20:1)-H | 57.23 | 63.06 | 68.01 |
| 1042 | PI(20:0/20:2)-H | 53.13 | NA | 120.5 |
| 1043 | PI(20:0/20:3)-H | 43.25 | NA | 15.79 |
| 1044 | PI(20:0/20:4)-H | 20.09 | 9.35 | 20.39 |
| 1045 | PI(20:0/20:5)-H | 73.16 | NA | NA |

| | | | | |
|---|---|---|---|---|
| 1046 | PI(20:0/22:5)-H | 53.42 | 25.5 | 16.65 |
| 1047 | PI(20:0/22:6)-H | NA | NA | NA |
| 1048 | LPS(14:0)-H | NA | NA | NA |
| 1049 | LPS(16:0)-H | 63.47 | NA | 45.37 |
| 1050 | LPS(16:1)-H | NA | NA | NA |
| 1051 | LPS(18:0)-H | NA | 27.9 | 37.82 |
| 1052 | LPS(18:1)-H | NA | 60.44 | 46.22 |
| 1053 | LPS(18:2)-H | NA | NA | 45.37 |
| 1054 | LPS(18:3)-H | NA | NA | NA |
| 1055 | LPS(20:0)-H | 35.22 | NA | NA |
| 1056 | LPS(20:1)-H | 12.1 | NA | NA |
| 1057 | LPS(20:2)-H | NA | NA | NA |
| 1058 | LPS(20:3)-H | NA | NA | NA |
| 1059 | LPS(20:4)-H | NA | NA | NA |
| 1060 | LPS(20:5)-H | NA | NA | NA |
| 1061 | LPS(22:4)-H | NA | NA | NA |
| 1062 | LPS(22:5)-H | NA | NA | NA |
| 1063 | LPS(22:6)-H | NA | NA | NA |
| 1064 | PS(14:1/14:1) | NA | NA | NA |
| 1065 | PS(14:0/14:0)-H | 54.09 | NA | 34.54 |
| 1066 | PS(14:0/18:1)-H | 15.31 | 45.36 | 41.95 |
| 1067 | PS(14:0/18:2)-H | 12.14 | 16.57 | 16.1 |
| 1068 | PS(14:0/18:3)-H | 49.65 | NA | NA |
| 1069 | PS(14:0/20:1)-H | 58.97 | 69.86 | 52.76 |
| 1070 | PS(14:0/20:2)-H | 35.35 | NA | 35.85 |
| 1071 | PS(14:0/20:3)-H | 49.77 | NA | 43.29 |
| 1072 | PS(14:0/20:4)-H | 13.79 | 25.58 | 14.06 |
| 1073 | PS(14:0/20:5)-H | NA | 70.71 | 58.24 |
| 1074 | PS(14:0/22:4)-H | 17.72 | 22.94 | 20.2 |
| 1075 | PS(14:0/22:5)-H | 13.95 | 40.45 | 35.54 |
| 1076 | PS(14:0/22:6)-H | 18.78 | 92.48 | 36.85 |
| 1077 | PS(16:0/14:0)-H | 42.71 | NA | 60.34 |
| 1078 | PS(16:0/16:0)-H | 19.73 | 15.86 | 8.85 |
| 1079 | PS(16:0/16:1)-H | NA | 36.79 | 13.96 |
| 1080 | PS(16:0/18:0)-H | 12.65 | 44.53 | 12.4 |
| 1081 | PS(16:0/18:1)-H | 19.09 | 21.81 | 12.51 |
| 1082 | PS(16:0/18:2)-H | 51.38 | 4.49 | 56.25 |
| 1083 | PS(16:0/18:3)-H | 18.87 | 32.84 | 36.22 |
| 1084 | PS(16:0/20:1)-H | 44.02 | 44.68 | 40.5 |
| 1085 | PS(16:0/20:2)-H | 51.7 | 50.7 | 29.36 |
| 1086 | PS(16:0/20:3)-H | 23.62 | 51.65 | 29.74 |
| 1087 | PS(16:0/20:4)-H | 8.59 | 40.77 | 34.03 |
| 1088 | PS(16:0/20:5)-H | 16.4 | 22.17 | 36.14 |

| | | | | |
|---|---|---|---|---|
| 1089 | PS(16:0/22:4)-H | 27.86 | NA | 35.69 |
| 1090 | PS(16:0/22:5)-H | NA | 18.17 | NA |
| 1091 | PS(16:0/22:6)-H | NA | 38.81 | 49.69 |
| 1092 | PS(18:0/14:0)-H | 42.64 | 29.66 | 50.21 |
| 1093 | PS(18:0/16:1)-H | 12.3 | 30.11 | 30.76 |
| 1094 | PS(18:0/18:0)-H | 14.39 | 12.43 | 6.74 |
| 1095 | PS(18:0/18:1)-H | 15.76 | 41.36 | 18.57 |
| 1096 | PS(18:0/18:2)-H | 18.57 | 89.47 | 35.02 |
| 1097 | PS(18:0/18:3)-H | 18.7 | 41.64 | 36.51 |
| 1098 | PS(18:0/20:0)-H | 22.75 | 11.31 | 18.57 |
| 1099 | PS(18:0/20:1)-H | 19.37 | 83.18 | 25.36 |
| 1100 | PS(18:0/20:2)-H | 21.18 | NA | 43.64 |
| 1101 | PS(18:0/20:3)-H | 142.77 | 43.02 | 54.16 |
| 1102 | PS(18:0/20:4)-H | 30.08 | 9.23 | 27.82 |
| 1103 | PS(18:0/20:5)-H | 26.49 | NA | 25.88 |
| 1104 | PS(18:0/22:4)-H | 47.76 | 53.87 | 17.04 |
| 1105 | PS(18:0/22:5)-H | 16.07 | 48.18 | 53.48 |
| 1106 | PS(18:0/22:6)-H | 88.86 | NA | 61.48 |
| 1107 | PS(18:1/16:1)-H | 24.25 | 18.54 | 8.28 |
| 1108 | PS(18:1/18:1)-H | 16.01 | 66.74 | 42.6 |
| 1109 | PS(18:1/18:2)-H | 25.33 | 13.13 | 39.85 |
| 1110 | PS(18:1/18:3)-H | 45.31 | 30.67 | 48.52 |
| 1111 | PS(18:1/20:1)-H | NA | 36.56 | 74.77 |
| 1112 | PS(18:1/20:2)-H | 36.11 | 41.03 | 42.61 |
| 1113 | PS(18:1/20:3)-H | 21.93 | 44.38 | 39.86 |
| 1114 | PS(18:1/20:4)-H | 14.04 | 16.39 | 21.02 |
| 1115 | PS(18:1/20:5)-H | 3.83 | NA | 27.58 |
| 1116 | PS(18:1/22:4)-H | 22.46 | 16.94 | 36.42 |
| 1117 | PS(18:1/22:5)-H | 23.06 | 20.65 | 25.87 |
| 1118 | PS(18:1/22:6)-H | 19.43 | 44.82 | 29.75 |
| 1119 | PS(18:2/16:1)-H | 10.63 | 27.62 | 21.99 |
| 1120 | PS(18:2/18:2)-H | 33.44 | 44.22 | 13.18 |
| 1121 | PS(18:2/18:3)-H | 19.28 | 17.27 | 71.43 |
| 1122 | PS(18:2/20:1)-H | 15.2 | 74.05 | 46.03 |
| 1123 | PS(18:2/20:2)-H | 35.04 | 42.49 | 55.15 |
| 1124 | PS(18:2/20:3)-H | 8.43 | 15.29 | 16.29 |
| 1125 | PS(18:2/20:4)-H | 11.89 | 18.65 | 7.09 |
| 1126 | PS(18:2/20:5)-H | 27.68 | 66.81 | 25.08 |
| 1127 | PS(18:2/22:4)-H | 7.74 | 42.78 | 26.9 |
| 1128 | PS(18:2/22:5)-H | 40.64 | 40.49 | 43.27 |
| 1129 | PS(18:2/22:6)-H | 20.84 | 68.61 | 19.68 |
| 1130 | PS(20:0/16:1)-H | 9.18 | 14 | 4.82 |
| 1131 | PS(20:0/18:1)-H | 15.67 | 31.68 | 7.08 |

| | | | | |
|---|---|---|---|---|
| 1132 | PS(20:0/18:2)-H | 4.42 | 13.09 | 10.63 |
| 1133 | PS(20:0/18:3)-H | 18.42 | 14.4 | 26.18 |
| 1134 | PS(20:0/20:1)-H | 33.87 | 20.37 | 50.92 |
| 1135 | PS(20:0/20:2)-H | 21.51 | 18.26 | 36.61 |
| 1136 | PS(20:0/20:3)-H | 11.46 | 23.19 | 9.22 |
| 1137 | PS(20:0/20:4)-H | 12.38 | 5.23 | 13.21 |
| 1138 | PS(20:0/20:5)-H | NA | 38.45 | 55.44 |
| 1139 | PS(20:0/22:4)-H | 27.48 | 42.3 | 13.29 |
| 1140 | PS(20:0/22:5)-H | 9.89 | 11.96 | 24.73 |
| 1141 | PS(20:0/22:6)-H | 13.58 | 28.79 | 8.57 |
| 1142 | L PA(14:0)-H | NA | NA | NA |
| 1143 | L PA(16:0)-H | NA | NA | NA |
| 1144 | L PA(16:1)-H | NA | NA | NA |
| 1145 | L PA(18:0)-H | NA | NA | NA |
| 1146 | L PA(18:1)-H | NA | NA | NA |
| 1147 | L PA(18:2)-H | NA | NA | NA |
| 1148 | PA(14:0/14:0)-H | NA | NA | NA |
| 1149 | PA(14:0/18:1)-H | NA | NA | 1.92 |
| 1150 | PA(14:0/18:2)-H | NA | NA | NA |
| 1151 | PA(14:0/18:3)-H | NA | NA | 30.91 |
| 1152 | PA(14:0/20:1)-H | NA | NA | NA |
| 1153 | PA(14:0/20:2)-H | NA | NA | NA |
| 1154 | PA(14:0/20:3)-H | NA | NA | NA |
| 1155 | PA(14:0/20:4)-H | NA | NA | 42.23 |
| 1156 | PA(14:0/20:5)-H | NA | NA | NA |
| 1157 | PA(14:0/22:4)-H | NA | NA | NA |
| 1158 | PA(14:0/22:5)-H | 39.59 | NA | NA |
| 1159 | PA(14:0/22:6)-H | NA | NA | NA |
| 1160 | PA(16:0/14:0)-H | NA | 71.49 | NA |
| 1161 | PA(16:0/16:0)-H | NA | 76.84 | 20.5 |
| 1162 | PA(16:0/16:1)-H | 83.93 | 37.03 | NA |
| 1163 | PA(16:0/18:0)-H | 15.74 | NA | NA |
| 1164 | PA(16:0/18:1)-H | NA | NA | NA |
| 1165 | PA(16:0/18:2)-H | 29.45 | 23.76 | 17.99 |
| 1166 | PA(16:0/18:3)-H | NA | NA | NA |
| 1167 | PA(16:0/20:1)-H | NA | NA | NA |
| 1168 | PA(16:0/20:2)-H | NA | NA | NA |
| 1169 | PA(16:0/20:3)-H | NA | 46.26 | 6.22 |
| 1170 | PA(16:0/20:4)-H | 31.29 | 57.37 | 25.51 |
| 1171 | PA(16:0/20:5)-H | NA | NA | NA |
| 1172 | PA(16:0/22:4)-H | NA | NA | NA |
| 1173 | PA(16:0/22:5)-H | 48.06 | NA | NA |
| 1174 | PA(16:0/22:6)-H | NA | NA | NA |

| | | | | |
|---|---|---|---|---|
| 1175 | PA(18:0/14:0)-H | NA | NA | 43.2 |
| 1176 | PA(18:0/16:1)-H | 19.26 | 58.86 | 40.14 |
| 1177 | PA(18:0/18:0)-H | NA | 63.68 | 33.63 |
| 1178 | PA(18:0/18:1)-H | 9.45 | 8.5 | 13.89 |
| 1179 | PA(18:0/18:2)-H | 6.33 | 9.27 | 4.48 |
| 1180 | PA(18:0/18:3)-H | 18.43 | 11.51 | 27.46 |
| 1181 | PA(18:0/20:0)-H | 40.75 | 13.69 | 35.66 |
| 1182 | PA(18:0/20:1)-H | 27.18 | 8.62 | 24.6 |
| 1183 | PA(18:0/20:2)-H | 33.92 | 32.37 | 31.12 |
| 1184 | PA(18:0/20:3)-H | 2.79 | 8.21 | 13.58 |
| 1185 | PA(18:0/20:4)-H | 7.23 | 9.76 | 13.78 |
| 1186 | PA(18:0/20:5)-H | 16.7 | 27.74 | 36.49 |
| 1187 | PA(18:0/22:4)-H | 9.97 | 4.69 | 10.47 |
| 1188 | PA(18:0/22:5)-H | 5.93 | 11.48 | 7 |
| 1189 | PA(18:0/22:6)-H | 15.78 | 7.2 | 5.68 |
| 1190 | PA(18:1/16:1)-H | NA | NA | NA |
| 1191 | PA(18:1/18:1)-H | 6.22 | 23.66 | 9.01 |
| 1192 | PA(18:1/18:2)-H | 23.1 | 8.29 | 9.21 |
| 1193 | PA(18:1/18:3)-H | 59.89 | NA | 58.59 |
| 1194 | PA(18:1/20:1)-H | 57.19 | NA | 49.89 |
| 1195 | PA(18:1/20:2)-H | 18.72 | 17.1 | NA |
| 1196 | PA(18:1/20:3)-H | 15.93 | 20.75 | 20.82 |
| 1197 | PA(18:1/20:4)-H | 21.71 | 30.9 | 23.82 |
| 1198 | PA(18:1/20:5)-H | 37.67 | NA | NA |
| 1199 | PA(18:1/22:4)-H | 44.82 | 83.57 | 33.86 |
| 1200 | PA(18:1/22:5)-H | 46.94 | 35.99 | 46.49 |
| 1201 | PA(18:1/22:6)-H | NA | 41.28 | NA |
| 1202 | PA(18:2/16:1)-H | 50.71 | NA | 34.08 |
| 1203 | PA(18:2/18:2)-H | 14.11 | NA | 6.68 |
| 1204 | PA(18:2/18:3)-H | NA | NA | NA |
| 1205 | PA(18:2/20:1)-H | NA | NA | 38.06 |
| 1206 | PA(18:2/20:2)-H | 75.61 | NA | 54.13 |
| 1207 | PA(18:2/20:3)-H | NA | NA | 48.6 |
| 1208 | PA(18:2/20:4)-H | 54.13 | 47.39 | 23.71 |
| 1209 | PA(18:2/20:5)-H | NA | NA | NA |
| 1210 | PA(18:2/22:4)-H | 72.04 | NA | 43.39 |
| 1211 | PA(18:2/22:5)-H | NA | NA | 85.31 |
| 1212 | PA(18:2/22:6)-H | NA | NA | 42.54 |
| 1213 | PA(20:0/16:1)-H | 49.49 | 21.22 | 9.64 |
| 1214 | PA(20:0/18:1)-H | 4.66 | 4.51 | 7.71 |
| 1215 | PA(20:0/18:2)-H | 10.63 | 9.32 | 9.73 |
| 1216 | PA(20:0/18:3)-H | 26.97 | 32.77 | 20.74 |
| 1217 | PA(20:0/20:1)-H | 49.52 | 51.25 | 15.5 |

| | | | | |
|---|---|---|---|---|
| 1218 | PA(20:0/20:2)-H | 28.52 | 39.04 | 25.42 |
| 1219 | PA(20:0/20:3)-H | 6.65 | 6.91 | 4.69 |
| 1220 | PA(20:0/20:4)-H | 14.76 | 4.99 | 4.45 |
| 1221 | PA(20:0/20:5)-H | 7.8 | 21.06 | 15.47 |
| 1222 | PA(20:0/22:4)-H | 13.66 | 7.14 | 11.05 |
| 1223 | PA(20:0/22:5)-H | 6.54 | 9.62 | 11.38 |
| 1224 | PA(20:0/22:6)-H | 8.22 | 6.87 | 8.56 |

Supplementary- table 8:

| S.No. | Sample Name | Fold Change (Low B12/ Normal B12 | p-value |
|---|---|---|---|
| 1 | SM | 0.979 | 0.819 |
| 2 | CE | 1.048 | 0.657 |
| 3 | CER | 1.014 | 0.921 |
| 4 | TAG | 1.064 | 0.454 |
| 5 | DAG | 1.023 | 0.818 |
| 6 | LPC | 1.016 | 0.779 |
| 7 | PC | 0.975 | 0.557 |
| 8 | LPE | 0.864 | 0.456 |
| 9 | PE | 0.858 | 0.192 |
| 10 | LPG | 0.720 | 0.262 |
| 11 | PG | 0.842 | 0.337 |
| 12 | PI | 0.978 | 0.834 |
| 13 | PS | 0.691 | 0.103 |
| 14 | PA | 0.812 | 0.233 |

Supplementary- table 9:

| S.No. | Sample Name | Fold change | P value |
|---|---|---|---|
| 1 | SM(14:0)+H | 0.9925 | 0.9194 |
| 2 | SM(16:0)+H | 1.0482 | 0.4044 |
| 3 | SM(18:0)+H | 1.0087 | 0.9035 |
| 4 | SM(18:1)+H | 1.0009 | 0.9910 |
| 5 | SM(20:0)+H | 0.8772 | 0.4504 |
| 6 | SM(20:1)+H | 1.0728 | 0.3430 |
| 7 | SM(22:0)+H | 0.7182 | 0.2729 |
| 8 | SM(22:1)+H | 0.9606 | 0.8547 |
| 9 | SM(24:0)+H | 1.0802 | 0.2437 |
| 10 | SM(24:1)+H | 0.9904 | 0.8961 |
| 11 | SM(26:0)+H | 1.2745 | 0.1346 |
| 12 | SM(26:1)+H | 1.0584 | 0.4467 |

| | | | |
|---|---|---|---|
| 13 | CE(24:0)+H | 0.9862 | 0.8498 |
| 14 | CE(22:6)+H | 1.1024 | 0.3808 |
| 15 | CE(20:0)+H | 1.0105 | 0.7356 |
| 16 | CE(20:1)+H | 0.9636 | 0.6606 |
| 17 | CE(18:2)+H | 1.0418 | 0.7137 |
| 18 | CE(18:3)+H | 0.9900 | 0.8682 |
| 19 | CE(20:3)+H | 0.9240 | 0.3333 |
| 20 | CE(20:4)+H | 0.9691 | 0.7852 |
| 21 | CE(20:5)+H | 0.9055 | 0.2595 |
| 22 | CE(22:2)+H | 0.8801 | 0.0746 |
| 23 | CE(22:4)+H | 0.9482 | 0.5122 |
| 24 | CER(18:0)+H | 0.8971 | 0.5878 |
| 25 | CER(20:0)+H | 0.9208 | 0.4321 |
| 26 | CER(22:0)+H | 1.0870 | 0.1778 |
| 27 | CER(22:1)+H | 1.0498 | 0.4814 |
| 28 | CER(24:0)+H | 1.0298 | 0.6803 |
| 29 | CER(24:1)+H | 0.9592 | 0.6241 |
| 30 | DCER(20:0)+H | 0.8046 | 0.0818 |
| 31 | DCER(22:0)+H | 1.1090 | 0.2441 |
| 32 | DCER(22:1)+H | 1.0677 | 0.4548 |
| 33 | DCER(24:0)+H | 1.0917 | 0.3327 |
| 34 | DCER(24:1)+H | 0.8986 | 0.4141 |
| 35 | HCER(16:0)+H | 0.7555 | 0.3729 |
| 36 | HCER(18:0)+H | 0.6226 | 0.1891 |
| 37 | HCER(18:1)+H | 0.9535 | 0.9216 |
| 38 | HCER(20:0)+H | 0.7568 | 0.3511 |
| 39 | HCER(20:1)+H | 0.9569 | 0.9222 |
| 40 | HCER(22:0)+H | 0.6401 | 0.1297 |
| 41 | HCER(22:1)+H | 1.1978 | 0.0935 |
| 42 | HCER(24:0)+H | 0.9469 | 0.7718 |
| 43 | HCER(24:1)+H | 0.6904 | 0.2209 |
| 44 | HCER(26:1)+H | 1.2373 | 0.6096 |
| 45 | HCER(d18:0/18:0)+H | 0.6744 | 0.0711 |
| 46 | HCER(d18:0/22:0)+H | 0.8548 | 0.5040 |
| 47 | HCER(d18:0/24:0)+H | 0.8850 | 0.5132 |
| 48 | HCER(d18:0/24:1)+H | 0.8208 | 0.3070 |
| 49 | HCER(d18:0/26:0)+H | 0.9125 | 0.5057 |
| 50 | HCER(d18:0/26:1)+H | 0.9776 | 0.9038 |
| 51 | LCER(14:0)+H | 1.2984 | 0.1317 |
| 52 | LCER(16:0)+H | 0.6957 | 0.0274 |
| 53 | LCER(18:0)+H | 1.0861 | 0.5634 |
| 54 | LCER(20:0)+H | 1.1409 | 0.5922 |
| 55 | LCER(24:1)+H | 0.8037 | 0.5541 |
| 56 | LCER(d18:0/18:0)+H | 0.8590 | 0.2800 |
| 57 | LCER(d18:0/20:0)+H | 0.9974 | 0.9894 |
| 58 | TAG(42:2/FA18:2)+NH4 | 1.1576 | 0.3560 |

| | | | |
|---|---|---|---|
| 59 | TAG(44:2/FA16:0)+NH4 | 1.3265 | 0.0202 |
| 60 | TAG(44:2/FA18:2)+NH4 | 1.2258 | 0.1461 |
| 61 | TAG(44:3/FA18:2)+NH4 | 1.0551 | 0.6954 |
| 62 | TAG(46:1/FA18:1)+NH4 | 0.8601 | 0.3761 |
| 63 | TAG(46:2/FA16:0)+NH4 | 1.0287 | 0.8009 |
| 64 | TAG(46:2/FA18:2)+NH4 | 1.1029 | 0.5737 |
| 65 | TAG(46:3/FA14:0)+NH4 | 0.9130 | 0.5849 |
| 66 | TAG(46:3/FA16:0)+NH4 | 1.3014 | 0.0079 |
| 67 | TAG(46:3/FA18:1)+NH4 | 1.1751 | 0.1626 |
| 68 | TAG(46:3/FA18:2)+NH4 | 1.3673 | 0.0059 |
| 69 | TAG(46:3/FA18:3)+NH4 | 1.0539 | 0.7363 |
| 70 | TAG(46:4/FA18:2)+NH4 | 1.6747 | 0.0050 |
| 71 | TAG(48:0/FA14:0)+NH4 | 0.7473 | 0.1097 |
| 72 | TAG(48:0/FA18:0)+NH4 | 0.6625 | 0.0293 |
| 73 | TAG(48:1/FA14:0)+NH4 | 0.8256 | 0.1451 |
| 74 | TAG(48:1/FA16:0)+NH4 | 0.8355 | 0.1933 |
| 75 | TAG(48:1/FA18:0)+NH4 | 0.7240 | 0.0448 |
| 76 | TAG(48:1/FA18:1)+NH4 | 0.7929 | 0.0734 |
| 77 | TAG(48:2/FA14:0)+NH4 | 0.9523 | 0.6522 |
| 78 | TAG(48:2/FA16:0)+NH4 | 0.8916 | 0.3187 |
| 79 | TAG(48:2/FA18:0)+NH4 | 1.0291 | 0.8309 |
| 80 | TAG(48:2/FA18:1)+NH4 | 0.7741 | 0.0389 |
| 81 | TAG(48:2/FA18:2)+NH4 | 1.0201 | 0.8559 |
| 82 | TAG(48:3/FA14:0)+NH4 | 0.9588 | 0.7055 |
| 83 | TAG(48:3/FA16:0)+NH4 | 1.0016 | 0.9873 |
| 84 | TAG(48:3/FA18:1)+NH4 | 1.0280 | 0.7809 |
| 85 | TAG(48:3/FA18:2)+NH4 | 1.0538 | 0.6312 |
| 86 | TAG(48:3/FA18:3)+NH4 | 0.9090 | 0.4472 |
| 87 | TAG(48:4/FA14:0)+NH4 | 1.0055 | 0.9703 |
| 88 | TAG(48:4/FA16:0)+NH4 | 1.1247 | 0.4861 |
| 89 | TAG(48:4/FA18:1)+NH4 | 1.1631 | 0.2176 |
| 90 | TAG(48:4/FA18:2)+NH4 | 1.4303 | 0.0090 |
| 91 | TAG(48:4/FA18:3)+NH4 | 0.9667 | 0.7784 |
| 92 | TAG(48:4/FA20:4)+NH4 | 0.8787 | 0.5472 |
| 93 | TAG(48:5/FA18:2)+NH4 | 1.5143 | 0.0118 |
| 94 | TAG(48:5/FA18:3)+NH4 | 1.3297 | 0.0169 |
| 95 | TAG(49:3/FA16:0)+NH4 | 1.0687 | 0.5111 |
| 96 | TAG(49:3/FA18:3)+NH4 | 1.0098 | 0.9278 |
| 97 | TAG(50:0/FA16:0)+NH4 | 0.7631 | 0.0435 |
| 98 | TAG(50:0/FA18:0)+NH4 | 0.7232 | 0.0468 |
| 99 | TAG(50:1/FA14:0)+NH4 | 0.7966 | 0.0492 |
| 100 | TAG(50:1/FA16:0)+NH4 | 0.8637 | 0.0908 |
| 101 | TAG(50:1/FA18:0)+NH4 | 0.7560 | 0.0435 |
| 102 | TAG(50:1/FA18:1)+NH4 | 0.8817 | 0.1300 |
| 103 | TAG(50:2/FA14:0)+NH4 | 0.8911 | 0.1274 |
| 104 | TAG(50:2/FA16:0)+NH4 | 0.9426 | 0.3370 |

| | | | |
|---|---|---|---|
| 105 | TAG(50:2/FA16:1)+NH4 | 0.7916 | 0.0138 |
| 106 | TAG(50:2/FA18:0)+NH4 | 0.9432 | 0.5868 |
| 107 | TAG(50:2/FA18:1)+NH4 | 0.8252 | 0.0094 |
| 108 | TAG(50:2/FA18:2)+NH4 | 1.1034 | 0.1332 |
| 109 | TAG(50:3/FA14:0)+NH4 | 1.1388 | 0.0613 |
| 110 | TAG(50:3/FA16:0)+NH4 | 0.9708 | 0.6475 |
| 111 | TAG(50:3/FA16:1)+NH4 | 0.8769 | 0.0899 |
| 112 | TAG(50:3/FA18:0)+NH4 | 0.9852 | 0.8920 |
| 113 | TAG(50:3/FA18:1)+NH4 | 1.0226 | 0.6860 |
| 114 | TAG(50:3/FA18:2)+NH4 | 1.0290 | 0.5780 |
| 115 | TAG(50:3/FA18:3)+NH4 | 0.9715 | 0.7373 |
| 116 | TAG(50:3/FA20:3)+NH4 | 0.8225 | 0.2913 |
| 117 | TAG(50:4/FA14:0)+NH4 | 1.5311 | 0.0011 |
| 118 | TAG(50:4/FA16:0)+NH4 | 0.8932 | 0.2234 |
| 119 | TAG(50:4/FA16:1)+NH4 | 0.8965 | 0.2816 |
| 120 | TAG(50:4/FA18:1)+NH4 | 1.0608 | 0.3863 |
| 121 | TAG(50:4/FA18:2)+NH4 | 1.3977 | 0.0034 |
| 122 | TAG(50:4/FA18:3)+NH4 | 0.8971 | 0.1441 |
| 123 | TAG(50:4/FA20:3)+NH4 | 0.8744 | 0.4354 |
| 124 | TAG(50:4/FA20:4)+NH4 | 0.8685 | 0.4338 |
| 125 | TAG(50:5/FA14:0)+NH4 | 1.5556 | 0.0167 |
| 126 | TAG(50:5/FA16:0)+NH4 | 0.9885 | 0.9232 |
| 127 | TAG(50:5/FA16:1)+NH4 | 0.9180 | 0.4595 |
| 128 | TAG(50:5/FA18:1)+NH4 | 1.0709 | 0.5546 |
| 129 | TAG(50:5/FA18:2)+NH4 | 1.4914 | 0.0026 |
| 130 | TAG(50:5/FA18:3)+NH4 | 1.2082 | 0.0609 |
| 131 | TAG(50:5/FA20:4)+NH4 | 0.7523 | 0.1460 |
| 132 | TAG(50:6/FA20:4)+NH4 | 1.0949 | 0.5914 |
| 133 | TAG(51:2/FA16:0)+NH4 | 0.8882 | 0.2920 |
| 134 | TAG(51:2/FA17:0)+NH4 | 0.7980 | 0.3799 |
| 135 | TAG(51:2/FA18:1)+NH4 | 0.7911 | 0.0717 |
| 136 | TAG(51:2/FA18:2)+NH4 | 1.0336 | 0.7130 |
| 137 | TAG(51:3/FA17:0)+NH4 | 0.9357 | 0.5054 |
| 138 | TAG(51:3/FA18:2)+NH4 | 1.0360 | 0.5969 |
| 139 | TAG(51:3/FA18:3)+NH4 | 0.9099 | 0.3747 |
| 140 | TAG(51:4/FA18:2)+NH4 | 1.3129 | 0.0146 |
| 141 | TAG(51:4/FA18:3)+NH4 | 0.8800 | 0.1771 |
| 142 | TAG(51:4/FA20:4)+NH4 | 0.8363 | 0.3075 |
| 143 | TAG(51:5/FA18:2)+NH4 | 1.1636 | 0.2659 |
| 144 | TAG(51:5/FA18:3)+NH4 | 1.1436 | 0.2567 |
| 145 | TAG(52:0/FA16:0)+NH4 | 0.8295 | 0.0862 |
| 146 | TAG(52:0/FA18:0)+NH4 | 0.7467 | 0.0345 |
| 147 | TAG(52:1/FA16:0)+NH4 | 0.8469 | 0.0122 |
| 148 | TAG(52:1/FA18:0)+NH4 | 0.8324 | 0.0300 |
| 149 | TAG(52:1/FA18:1)+NH4 | 0.8733 | 0.0266 |
| 150 | TAG(52:1/FA20:0)+NH4 | 0.9903 | 0.9205 |

| | | | |
|---|---|---|---|
| 151 | TAG(52:1/FA20:1)+NH4 | 0.8924 | 0.2414 |
| 152 | TAG(52:2/FA14:0)+NH4 | 0.8225 | 0.0949 |
| 153 | TAG(52:2/FA16:0)+NH4 | 0.9207 | 0.0528 |
| 154 | TAG(52:2/FA16:1)+NH4 | 0.7947 | 0.0018 |
| 155 | TAG(52:2/FA18:0)+NH4 | 0.9895 | 0.8670 |
| 156 | TAG(52:2/FA18:1)+NH4 | 0.9041 | 0.0318 |
| 157 | TAG(52:2/FA18:2)+NH4 | 1.0721 | 0.0812 |
| 158 | TAG(52:2/FA20:0)+NH4 | 1.3307 | 0.0098 |
| 159 | TAG(52:2/FA20:1)+NH4 | 0.8548 | 0.1183 |
| 160 | TAG(52:2/FA20:2)+NH4 | 0.9120 | 0.3174 |
| 161 | TAG(52:3/FA14:0)+NH4 | 1.0850 | 0.4114 |
| 162 | TAG(52:3/FA16:0)+NH4 | 1.1173 | 0.0053 |
| 163 | TAG(52:3/FA16:1)+NH4 | 0.8727 | 0.0400 |
| 164 | TAG(52:3/FA18:0)+NH4 | 1.0010 | 0.9839 |
| 165 | TAG(52:3/FA18:1)+NH4 | 1.0406 | 0.1809 |
| 166 | TAG(52:3/FA18:2)+NH4 | 1.0867 | 0.0326 |
| 167 | TAG(52:3/FA18:3)+NH4 | 0.9720 | 0.5650 |
| 168 | TAG(52:3/FA20:0)+NH4 | 1.2069 | 0.0169 |
| 169 | TAG(52:3/FA20:1)+NH4 | 1.1127 | 0.3122 |
| 170 | TAG(52:3/FA20:2)+NH4 | 0.8909 | 0.1451 |
| 171 | TAG(52:3/FA20:3)+NH4 | 0.8816 | 0.3420 |
| 172 | TAG(52:4/FA14:0)+NH4 | 1.1626 | 0.2288 |
| 173 | TAG(52:4/FA16:0)+NH4 | 1.2981 | 0.0010 |
| 174 | TAG(52:4/FA16:1)+NH4 | 1.0978 | 0.1514 |
| 175 | TAG(52:4/FA18:0)+NH4 | 0.9798 | 0.8175 |
| 176 | TAG(52:4/FA18:1)+NH4 | 1.0609 | 0.2542 |
| 177 | TAG(52:4/FA18:2)+NH4 | 1.3285 | 0.0004 |
| 178 | TAG(52:4/FA18:3)+NH4 | 1.0064 | 0.8857 |
| 179 | TAG(52:4/FA20:0)+NH4 | 1.3420 | 0.0033 |
| 180 | TAG(52:4/FA20:2)+NH4 | 1.0162 | 0.8597 |
| 181 | TAG(52:4/FA20:3)+NH4 | 0.8662 | 0.2430 |
| 182 | TAG(52:4/FA20:4)+NH4 | 0.8704 | 0.3586 |
| 183 | TAG(52:4/FA22:4)+NH4 | 0.8514 | 0.1805 |
| 184 | TAG(52:5/FA14:0)+NH4 | 1.2320 | 0.2425 |
| 185 | TAG(52:5/FA16:0)+NH4 | 1.2675 | 0.0231 |
| 186 | TAG(52:5/FA16:1)+NH4 | 1.2943 | 0.0206 |
| 187 | TAG(52:5/FA18:1)+NH4 | 1.0489 | 0.6026 |
| 188 | TAG(52:5/FA18:2)+NH4 | 1.3462 | 0.0048 |
| 189 | TAG(52:5/FA18:3)+NH4 | 1.1668 | 0.0531 |
| 190 | TAG(52:5/FA20:3)+NH4 | 1.1273 | 0.4244 |
| 191 | TAG(52:5/FA20:4)+NH4 | 0.8240 | 0.2464 |
| 192 | TAG(52:5/FA20:5)+NH4 | 0.8718 | 0.4256 |
| 193 | TAG(52:5/FA22:5)+NH4 | 1.1821 | 0.3239 |
| 194 | TAG(52:6/FA14:0)+NH4 | 1.1488 | 0.4520 |
| 195 | TAG(52:6/FA16:0)+NH4 | 1.0940 | 0.5303 |
| 196 | TAG(52:6/FA16:1)+NH4 | 1.1463 | 0.3104 |

| | | | |
|---|---|---|---|
| 197 | TAG(52:6/FA18:1)+NH4 | 1.0843 | 0.4876 |
| 198 | TAG(52:6/FA18:2)+NH4 | 1.3747 | 0.0218 |
| 199 | TAG(52:6/FA18:3)+NH4 | 1.1881 | 0.0977 |
| 200 | TAG(52:6/FA20:4)+NH4 | 1.1039 | 0.6129 |
| 201 | TAG(52:6/FA20:5)+NH4 | 0.6879 | 0.0331 |
| 202 | TAG(52:6/FA22:6)+NH4 | 1.0163 | 0.9147 |
| 203 | TAG(52:7/FA18:1)+NH4 | 0.8104 | 0.0883 |
| 204 | TAG(52:7/FA20:5)+NH4 | 0.9597 | 0.8603 |
| 205 | TAG(52:7/FA22:6)+NH4 | 1.0021 | 0.9892 |
| 206 | TAG(52:8/FA18:2)+NH4 | 1.0645 | 0.5171 |
| 207 | TAG(53:0/FA16:0)+NH4 | 0.9437 | 0.6547 |
| 208 | TAG(53:1/FA16:0)+NH4 | 0.8994 | 0.4063 |
| 209 | TAG(53:1/FA17:0)+NH4 | 0.7243 | 0.1187 |
| 210 | TAG(53:1/FA18:0)+NH4 | 0.7158 | 0.1248 |
| 211 | TAG(53:1/FA18:1)+NH4 | 0.8113 | 0.0978 |
| 212 | TAG(53:2/FA16:0)+NH4 | 0.9266 | 0.4617 |
| 213 | TAG(53:2/FA17:0)+NH4 | 0.8150 | 0.1691 |
| 214 | TAG(53:2/FA18:1)+NH4 | 0.8391 | 0.1191 |
| 215 | TAG(53:2/FA18:2)+NH4 | 1.0897 | 0.1803 |
| 216 | TAG(53:3/FA16:0)+NH4 | 1.0990 | 0.3157 |
| 217 | TAG(53:3/FA17:0)+NH4 | 1.1146 | 0.0480 |
| 218 | TAG(53:3/FA18:2)+NH4 | 1.0921 | 0.0945 |
| 219 | TAG(53:4/FA16:0)+NH4 | 1.2446 | 0.0762 |
| 220 | TAG(53:4/FA17:0)+NH4 | 1.3379 | 0.0025 |
| 221 | TAG(53:4/FA18:2)+NH4 | 1.2546 | 0.0218 |
| 222 | TAG(53:4/FA18:3)+NH4 | 0.9574 | 0.7054 |
| 223 | TAG(53:4/FA20:4)+NH4 | 0.8336 | 0.2706 |
| 224 | TAG(53:5/FA20:4)+NH4 | 0.7376 | 0.2257 |
| 225 | TAG(53:6/FA20:4)+NH4 | 0.7224 | 0.1547 |
| 226 | TAG(54:0/FA16:0)+NH4 | 0.9468 | 0.7224 |
| 227 | TAG(54:0/FA18:0)+NH4 | 0.8651 | 0.4600 |
| 228 | TAG(54:1/FA16:0)+NH4 | 0.9388 | 0.6390 |
| 229 | TAG(54:1/FA18:0)+NH4 | 0.8627 | 0.1073 |
| 230 | TAG(54:1/FA18:1)+NH4 | 0.8516 | 0.0129 |
| 231 | TAG(54:1/FA20:0)+NH4 | 1.0195 | 0.8503 |
| 232 | TAG(54:1/FA20:1)+NH4 | 0.9614 | 0.6385 |
| 233 | TAG(54:2/FA16:0)+NH4 | 1.0048 | 0.9588 |
| 234 | TAG(54:2/FA18:0)+NH4 | 0.9109 | 0.1023 |
| 235 | TAG(54:2/FA18:1)+NH4 | 0.9360 | 0.1653 |
| 236 | TAG(54:2/FA18:2)+NH4 | 1.1804 | 0.0021 |
| 237 | TAG(54:2/FA20:0)+NH4 | 1.2489 | 0.0601 |
| 238 | TAG(54:2/FA20:1)+NH4 | 0.9964 | 0.9660 |
| 239 | TAG(54:2/FA20:2)+NH4 | 0.9233 | 0.2168 |
| 240 | TAG(54:3/FA16:0)+NH4 | 1.1847 | 0.0661 |
| 241 | TAG(54:3/FA16:1)+NH4 | 0.9392 | 0.6006 |
| 242 | TAG(54:3/FA18:0)+NH4 | 1.1237 | 0.0198 |

| | | | |
|---|---|---|---|
| 243 | TAG(54:3/FA18:1)+NH4 | 0.9158 | 0.2344 |
| 244 | TAG(54:3/FA18:2)+NH4 | 1.1987 | 0.0018 |
| 245 | TAG(54:3/FA18:3)+NH4 | 1.1619 | 0.0487 |
| 246 | TAG(54:3/FA20:1)+NH4 | 1.1735 | 0.1015 |
| 247 | TAG(54:3/FA20:2)+NH4 | 0.9232 | 0.1059 |
| 248 | TAG(54:3/FA20:3)+NH4 | 0.8186 | 0.0610 |
| 249 | TAG(54:4/FA16:0)+NH4 | 1.0621 | 0.2806 |
| 250 | TAG(54:4/FA16:1)+NH4 | 1.1189 | 0.4053 |
| 251 | TAG(54:4/FA18:0)+NH4 | 1.4112 | 0.0008 |
| 252 | TAG(54:4/FA18:1)+NH4 | 1.1086 | 0.2018 |
| 253 | TAG(54:4/FA18:2)+NH4 | 1.1767 | 0.0405 |
| 254 | TAG(54:4/FA18:3)+NH4 | 1.0693 | 0.3020 |
| 255 | TAG(54:4/FA20:1)+NH4 | 1.2241 | 0.1128 |
| 256 | TAG(54:4/FA20:2)+NH4 | 1.0938 | 0.1383 |
| 257 | TAG(54:4/FA20:3)+NH4 | 0.8926 | 0.1691 |
| 258 | TAG(54:4/FA20:4)+NH4 | 0.8719 | 0.3079 |
| 259 | TAG(54:4/FA22:4)+NH4 | 0.9271 | 0.6143 |
| 260 | TAG(54:5/FA16:0)+NH4 | 0.9956 | 0.9695 |
| 261 | TAG(54:5/FA16:1)+NH4 | 1.0109 | 0.8965 |
| 262 | TAG(54:5/FA18:0)+NH4 | 1.2729 | 0.0718 |
| 263 | TAG(54:5/FA18:1)+NH4 | 1.2836 | 0.0643 |
| 264 | TAG(54:5/FA18:2)+NH4 | 1.3580 | 0.0313 |
| 265 | TAG(54:5/FA18:3)+NH4 | 1.0230 | 0.7855 |
| 266 | TAG(54:5/FA20:2)+NH4 | 1.0508 | 0.6170 |
| 267 | TAG(54:5/FA20:3)+NH4 | 1.0370 | 0.7146 |
| 268 | TAG(54:5/FA20:4)+NH4 | 0.8880 | 0.3551 |
| 269 | TAG(54:5/FA20:5)+NH4 | 0.8147 | 0.1270 |
| 270 | TAG(54:5/FA22:4)+NH4 | 0.7970 | 0.1258 |
| 271 | TAG(54:5/FA22:5)+NH4 | 1.0823 | 0.7194 |
| 272 | TAG(54:6/FA16:0)+NH4 | 1.0521 | 0.7386 |
| 273 | TAG(54:6/FA16:1)+NH4 | 0.8986 | 0.4365 |
| 274 | TAG(54:6/FA18:1)+NH4 | 1.2567 | 0.2435 |
| 275 | TAG(54:6/FA18:2)+NH4 | 1.6395 | 0.0166 |
| 276 | TAG(54:6/FA18:3)+NH4 | 1.3732 | 0.0257 |
| 277 | TAG(54:6/FA20:3)+NH4 | 0.9613 | 0.7714 |
| 278 | TAG(54:6/FA20:4)+NH4 | 1.0972 | 0.5793 |
| 279 | TAG(54:6/FA20:5)+NH4 | 0.7456 | 0.0165 |
| 280 | TAG(54:6/FA22:5)+NH4 | 1.0318 | 0.8564 |
| 281 | TAG(54:6/FA22:6)+NH4 | 1.0258 | 0.8583 |
| 282 | TAG(54:7/FA16:1)+NH4 | 0.7884 | 0.1820 |
| 283 | TAG(54:7/FA18:1)+NH4 | 1.0667 | 0.6613 |
| 284 | TAG(54:7/FA18:2)+NH4 | 1.8595 | 0.0157 |
| 285 | TAG(54:7/FA18:3)+NH4 | 1.5982 | 0.0117 |
| 286 | TAG(54:7/FA20:4)+NH4 | 0.9677 | 0.8635 |
| 287 | TAG(54:7/FA20:5)+NH4 | 0.8133 | 0.2605 |
| 288 | TAG(54:7/FA22:5)+NH4 | 1.3109 | 0.0886 |

| | | | |
|---|---|---|---|
| 289 | TAG(54:7/FA22:6)+NH4 | 0.9866 | 0.9305 |
| 290 | TAG(54:8/FA18:2)+NH4 | 1.2233 | 0.2149 |
| 291 | TAG(54:8/FA18:3)+NH4 | 1.3916 | 0.0904 |
| 292 | TAG(54:8/FA20:4)+NH4 | 0.9243 | 0.7145 |
| 293 | TAG(54:8/FA20:5)+NH4 | 0.8264 | 0.4115 |
| 294 | TAG(54:8/FA22:6)+NH4 | 1.2362 | 0.1592 |
| 295 | TAG(55:1/FA16:0)+NH4 | 1.0120 | 0.9045 |
| 296 | TAG(55:1/FA18:1)+NH4 | 0.8516 | 0.1868 |
| 297 | TAG(55:2/FA18:1)+NH4 | 0.7709 | 0.3176 |
| 298 | TAG(55:2/FA18:2)+NH4 | 1.1272 | 0.2662 |
| 299 | TAG(55:3/FA18:1)+NH4 | 0.6166 | 0.1910 |
| 300 | TAG(55:3/FA18:2)+NH4 | 1.1413 | 0.2274 |
| 301 | TAG(55:4/FA18:1)+NH4 | 1.0461 | 0.6318 |
| 302 | TAG(55:4/FA18:2)+NH4 | 1.0741 | 0.6680 |
| 303 | TAG(55:5/FA18:1)+NH4 | 1.2189 | 0.0881 |
| 304 | TAG(55:5/FA18:2)+NH4 | 1.3468 | 0.0147 |
| 305 | TAG(55:5/FA20:4)+NH4 | 0.7968 | 0.2305 |
| 306 | TAG(55:7/FA22:6)+NH4 | 1.0018 | 0.9897 |
| 307 | TAG(56:10/FA18:2)+NH4 | 1.1361 | 0.2592 |
| 308 | TAG(56:1/FA18:1)+NH4 | 0.9756 | 0.8543 |
| 309 | TAG(56:2/FA16:0)+NH4 | 1.0158 | 0.9445 |
| 310 | TAG(56:2/FA18:0)+NH4 | 1.0227 | 0.8296 |
| 311 | TAG(56:2/FA20:0)+NH4 | 1.0248 | 0.8107 |
| 312 | TAG(56:2/FA20:1)+NH4 | 1.0908 | 0.4653 |
| 313 | TAG(56:3/FA18:0)+NH4 | 1.2494 | 0.0759 |
| 314 | TAG(56:3/FA18:1)+NH4 | 0.9697 | 0.7984 |
| 315 | TAG(56:3/FA18:2)+NH4 | 1.1816 | 0.3084 |
| 316 | TAG(56:3/FA20:0)+NH4 | 1.3209 | 0.0346 |
| 317 | TAG(56:3/FA20:1)+NH4 | 1.0553 | 0.6245 |
| 318 | TAG(56:3/FA20:2)+NH4 | 1.0201 | 0.8062 |
| 319 | TAG(56:4/FA16:0)+NH4 | 1.0788 | 0.5935 |
| 320 | TAG(56:4/FA18:0)+NH4 | 1.0640 | 0.4339 |
| 321 | TAG(56:4/FA18:1)+NH4 | 1.1447 | 0.2655 |
| 322 | TAG(56:4/FA18:2)+NH4 | 1.1883 | 0.2137 |
| 323 | TAG(56:4/FA20:1)+NH4 | 1.2623 | 0.0996 |
| 324 | TAG(56:4/FA20:2)+NH4 | 0.9081 | 0.3115 |
| 325 | TAG(56:4/FA20:3)+NH4 | 0.8408 | 0.0824 |
| 326 | TAG(56:4/FA20:4)+NH4 | 0.8154 | 0.1614 |
| 327 | TAG(56:4/FA22:4)+NH4 | 0.8151 | 0.0719 |
| 328 | TAG(56:5/FA16:0)+NH4 | 0.9079 | 0.4051 |
| 329 | TAG(56:5/FA18:0)+NH4 | 1.1125 | 0.2932 |
| 330 | TAG(56:5/FA18:1)+NH4 | 1.0071 | 0.9459 |
| 331 | TAG(56:5/FA18:2)+NH4 | 1.4093 | 0.0444 |
| 332 | TAG(56:5/FA20:1)+NH4 | 1.4913 | 0.0328 |
| 333 | TAG(56:5/FA20:2)+NH4 | 1.0230 | 0.8630 |
| 334 | TAG(56:5/FA20:3)+NH4 | 0.8053 | 0.1152 |

| | | | |
|---|---|---|---|
| 335 | TAG(56:5/FA20:4)+NH4 | 0.9470 | 0.6703 |
| 336 | TAG(56:5/FA22:4)+NH4 | 0.8705 | 0.1507 |
| 337 | TAG(56:5/FA22:5)+NH4 | 1.4721 | 0.2589 |
| 338 | TAG(56:6/FA16:0)+NH4 | 0.8801 | 0.3779 |
| 339 | TAG(56:6/FA18:0)+NH4 | 0.9699 | 0.8588 |
| 340 | TAG(56:6/FA18:1)+NH4 | 0.8867 | 0.4245 |
| 341 | TAG(56:6/FA18:2)+NH4 | 1.3087 | 0.0846 |
| 342 | TAG(56:6/FA18:3)+NH4 | 1.3679 | 0.0652 |
| 343 | TAG(56:6/FA20:2)+NH4 | 1.2389 | 0.1970 |
| 344 | TAG(56:6/FA20:3)+NH4 | 0.8550 | 0.4417 |
| 345 | TAG(56:6/FA20:4)+NH4 | 0.8538 | 0.3498 |
| 346 | TAG(56:6/FA20:5)+NH4 | 0.7112 | 0.0537 |
| 347 | TAG(56:6/FA22:4)+NH4 | 1.0700 | 0.5893 |
| 348 | TAG(56:6/FA22:5)+NH4 | 1.1300 | 0.7467 |
| 349 | TAG(56:6/FA22:6)+NH4 | 1.0061 | 0.9621 |
| 350 | TAG(56:7/FA16:0)+NH4 | 0.9422 | 0.5921 |
| 351 | TAG(56:7/FA16:1)+NH4 | 0.7778 | 0.4286 |
| 352 | TAG(56:7/FA18:0)+NH4 | 0.9995 | 0.9979 |
| 353 | TAG(56:7/FA18:1)+NH4 | 0.8853 | 0.4164 |
| 354 | TAG(56:7/FA18:2)+NH4 | 1.0580 | 0.7781 |
| 355 | TAG(56:7/FA18:3)+NH4 | 1.3910 | 0.0243 |
| 356 | TAG(56:7/FA20:3)+NH4 | 1.0233 | 0.9293 |
| 357 | TAG(56:7/FA20:4)+NH4 | 1.0002 | 0.9992 |
| 358 | TAG(56:7/FA20:5)+NH4 | 0.5804 | 0.0165 |
| 359 | TAG(56:7/FA22:4)+NH4 | 1.1070 | 0.6592 |
| 360 | TAG(56:7/FA22:5)+NH4 | 1.2451 | 0.2247 |
| 361 | TAG(56:7/FA22:6)+NH4 | 1.1571 | 0.3195 |
| 362 | TAG(56:8/FA16:0)+NH4 | 1.0551 | 0.6189 |
| 363 | TAG(56:8/FA18:1)+NH4 | 0.8668 | 0.3417 |
| 364 | TAG(56:8/FA18:2)+NH4 | 1.2530 | 0.2717 |
| 365 | TAG(56:8/FA18:3)+NH4 | 0.9873 | 0.9506 |
| 366 | TAG(56:8/FA20:4)+NH4 | 1.1370 | 0.6510 |
| 367 | TAG(56:8/FA20:5)+NH4 | 0.6603 | 0.2427 |
| 368 | TAG(56:8/FA22:5)+NH4 | 1.0136 | 0.9301 |
| 369 | TAG(56:8/FA22:6)+NH4 | 1.2362 | 0.2367 |
| 370 | TAG(56:9/FA18:3)+NH4 | 0.8637 | 0.2742 |
| 371 | TAG(56:9/FA20:4)+NH4 | 1.0901 | 0.7242 |
| 372 | TAG(56:9/FA20:5)+NH4 | 1.3209 | 0.2501 |
| 373 | TAG(56:9/FA22:6)+NH4 | 1.3141 | 0.0588 |
| 374 | TAG(57:2/FA18:1)+NH4 | 0.9914 | 0.9409 |
| 375 | TAG(57:3/FA18:2)+NH4 | 1.0335 | 0.9078 |
| 376 | TAG(58:10/FA18:2)+NH4 | 0.7783 | 0.2923 |
| 377 | TAG(58:10/FA20:4)+NH4 | 0.9993 | 0.9990 |
| 378 | TAG(58:10/FA20:5)+NH4 | 0.8451 | 0.6022 |
| 379 | TAG(58:10/FA22:5)+NH4 | 1.5148 | 0.2116 |
| 380 | TAG(58:10/FA22:6)+NH4 | 1.6829 | 0.0064 |

| | | | |
|---|---|---|---|
| 381 | TAG(58:2/FA18:1)+NH4 | 0.8785 | 0.5234 |
| 382 | TAG(58:6/FA16:0)+NH4 | 1.0772 | 0.5289 |
| 383 | TAG(58:6/FA18:0)+NH4 | 0.9559 | 0.7861 |
| 384 | TAG(58:6/FA18:1)+NH4 | 0.8688 | 0.3518 |
| 385 | TAG(58:6/FA20:4)+NH4 | 0.7941 | 0.2652 |
| 386 | TAG(58:6/FA22:4)+NH4 | 0.8864 | 0.3767 |
| 387 | TAG(58:6/FA22:5)+NH4 | 1.1462 | 0.4733 |
| 388 | TAG(58:7/FA16:0)+NH4 | 1.1012 | 0.4186 |
| 389 | TAG(58:7/FA18:0)+NH4 | 0.9872 | 0.9419 |
| 390 | TAG(58:7/FA18:1)+NH4 | 0.6729 | 0.0810 |
| 391 | TAG(58:7/FA18:2)+NH4 | 1.0388 | 0.8211 |
| 392 | TAG(58:7/FA20:4)+NH4 | 0.5984 | 0.6060 |
| 393 | TAG(58:7/FA22:4)+NH4 | 1.0005 | 0.9977 |
| 394 | TAG(58:7/FA22:5)+NH4 | 1.2218 | 0.5934 |
| 395 | TAG(58:7/FA22:6)+NH4 | 1.1932 | 0.2743 |
| 396 | TAG(58:8/FA18:1)+NH4 | 0.9993 | 0.9968 |
| 397 | TAG(58:8/FA18:2)+NH4 | 0.9067 | 0.6092 |
| 398 | TAG(58:8/FA20:3)+NH4 | 0.8080 | 0.2590 |
| 399 | TAG(58:8/FA20:4)+NH4 | 1.3272 | 0.3128 |
| 400 | TAG(58:8/FA22:5)+NH4 | 2.5714 | 0.1242 |
| 401 | TAG(58:8/FA22:6)+NH4 | 1.1907 | 0.2292 |
| 402 | TAG(58:9/FA18:1)+NH4 | 0.8514 | 0.2841 |
| 403 | TAG(58:9/FA18:2)+NH4 | 1.1712 | 0.3253 |
| 404 | TAG(58:9/FA20:4)+NH4 | 1.0451 | 0.9035 |
| 405 | TAG(58:9/FA22:5)+NH4 | 1.0378 | 0.7880 |
| 406 | TAG(58:9/FA22:6)+NH4 | 1.4949 | 0.0141 |
| 407 | TAG(60:10/FA22:5)+NH4 | 1.7350 | 0.1006 |
| 408 | TAG(60:10/FA22:6)+NH4 | 1.2877 | 0.0984 |
| 409 | TAG(60:11/FA22:5)+NH4 | 1.0006 | 0.9974 |
| 410 | TAG(60:11/FA22:6)+NH4 | 1.1988 | 0.2094 |
| 411 | TAG(60:12/FA22:6)+NH4 | 1.2343 | 0.2062 |
| 412 | DAG(14:0/20:0)+NH4 | 1.1337 | 0.4858 |
| 413 | DAG(16:0/18:3)+NH4 | 1.3871 | 0.0333 |
| 414 | DAG(18:1/20:2)+NH4 | 0.9199 | 0.7080 |
| 415 | DAG(18:2/20:4)+NH4 | 2.5011 | 0.0522 |
| 416 | DAG(18:1/20:5)+NH4 | 0.7270 | 0.1598 |
| 417 | DAG(20:0/20:0)+NH4 | 0.9976 | 0.3987 |
| 418 | DAG(18:1/22:6)+NH4 | 0.9471 | 0.8240 |
| 419 | LPC(14:0)+AcO | 0.9334 | 0.4869 |
| 420 | LPC(16:0)+AcO | 0.9967 | 0.8989 |
| 421 | LPC(16:1)+AcO | 0.9250 | 0.2716 |
| 422 | LPC(18:0)+AcO | 1.0553 | 0.1066 |
| 423 | LPC(18:1)+AcO | 0.9891 | 0.8586 |
| 424 | LPC(18:2)+AcO | 0.9041 | 0.1178 |
| 425 | LPC(18:3)+AcO | 0.8781 | 0.2447 |
| 426 | LPC(20:0)+AcO | 1.1025 | 0.3049 |

| | | | |
|---|---|---|---|
| 427 | LPC(20:1)+AcO | 1.0311 | 0.8268 |
| 428 | LPC(20:2)+AcO | 1.0379 | 0.6060 |
| 429 | LPC(20:3)+AcO | 0.7975 | 0.1967 |
| 430 | LPC(20:4)+AcO | 0.8755 | 0.1079 |
| 431 | LPC(20:5)+AcO | 0.8973 | 0.7073 |
| 432 | LPC(22:4)+AcO | 1.0131 | 0.9161 |
| 433 | LPC(22:5)+AcO | 0.9320 | 0.6374 |
| 434 | LPC(22:6)+AcO | 0.6896 | 0.1293 |
| 435 | PC(14:0/14:0)+AcO | 0.8202 | 0.3291 |
| 436 | PC(14:0/18:1)+AcO | 0.9248 | 0.4904 |
| 437 | PC(14:0/18:2)+AcO | 1.1470 | 0.1231 |
| 438 | PC(14:0/18:3)+AcO | 1.0329 | 0.7704 |
| 439 | PC(14:0/20:2)+AcO | 1.0472 | 0.6479 |
| 440 | PC(14:0/20:3)+AcO | 1.0381 | 0.7573 |
| 441 | PC(14:0/20:4)+AcO | 0.9628 | 0.7551 |
| 442 | PC(14:0/20:5)+AcO | 1.0052 | 0.9659 |
| 443 | PC(14:0/22:4)+AcO | 1.0134 | 0.9207 |
| 444 | PC(14:0/22:5)+AcO | 1.0173 | 0.8851 |
| 445 | PC(14:0/22:6)+AcO | 0.9071 | 0.4980 |
| 446 | PC(16:0/14:0)+AcO | 0.8394 | 0.1189 |
| 447 | PC(16:0/16:0)+AcO | 1.0768 | 0.1285 |
| 448 | PC(16:0/16:1)+AcO | 0.8371 | 0.1960 |
| 449 | PC(16:0/18:0)+AcO | 1.1196 | 0.1357 |
| 450 | PC(16:0/18:1)+AcO | 0.9194 | 0.0305 |
| 451 | PC(16:0/18:2)+AcO | 1.0567 | 0.0807 |
| 452 | PC(16:0/18:3)+AcO | 0.8389 | 0.0403 |
| 453 | PC(16:0/20:1)+AcO | 0.9904 | 0.9290 |
| 454 | PC(16:0/20:2)+AcO | 0.9772 | 0.6777 |
| 455 | PC(16:0/20:3)+AcO | 0.9651 | 0.5201 |
| 456 | PC(16:0/20:4)+AcO | 0.9378 | 0.1924 |
| 457 | PC(16:0/20:5)+AcO | 0.6868 | 0.0062 |
| 458 | PC(16:0/22:4)+AcO | 1.0501 | 0.5370 |
| 459 | PC(16:0/22:5)+AcO | 0.9531 | 0.6087 |
| 460 | PC(16:1/18:1)+AcO | 1.0349 | 0.8060 |
| 461 | PC(16:1/18:2)+AcO | 0.9800 | 0.7581 |
| 462 | PC(16:0/22:6)+AcO | 0.8008 | 0.0774 |
| 463 | PC(18:0/14:0)+AcO | 0.8516 | 0.1070 |
| 464 | PC(18:0/16:1)+AcO | 0.8152 | 0.2601 |
| 465 | PC(18:0/18:0)+AcO | 0.9608 | 0.4686 |
| 466 | PC(18:0/18:1)+AcO | 0.8539 | 0.0046 |
| 467 | PC(18:0/18:2)+AcO | 1.0750 | 0.0060 |
| 468 | PC(18:0/18:3)+AcO | 0.8297 | 0.0333 |
| 469 | PC(18:0/20:0)+AcO | 1.2730 | 0.1206 |
| 470 | PC(18:0/20:1)+AcO | 0.8954 | 0.3639 |
| 471 | PC(18:0/20:2)+AcO | 0.9313 | 0.2699 |
| 472 | PC(18:0/20:3)+AcO | 1.0216 | 0.7150 |

| | | | |
|---|---|---|---|
| 473 | PC(18:0/20:4)+AcO | 0.9907 | 0.8655 |
| 474 | PC(18:0/20:5)+AcO | 0.7373 | 0.0171 |
| 475 | PC(18:0/22:4)+AcO | 1.0355 | 0.7410 |
| 476 | PC(18:0/22:5)+AcO | 0.9866 | 0.8894 |
| 477 | PC(18:0/22:6)+AcO | 0.8499 | 0.1784 |
| 478 | PC(18:1/16:1)+AcO | 0.9825 | 0.9088 |
| 479 | PC(18:1/18:1)+AcO | 0.9008 | 0.1249 |
| 480 | PC(18:1/18:2)+AcO | 1.0504 | 0.3565 |
| 481 | PC(18:1/18:3)+AcO | 0.8523 | 0.2324 |
| 482 | PC(18:1/20:1)+AcO | 0.8979 | 0.4554 |
| 483 | PC(18:1/20:2)+AcO | 0.9484 | 0.4997 |
| 484 | PC(18:1/20:3)+AcO | 1.0515 | 0.5082 |
| 485 | PC(18:1/20:4)+AcO | 0.9844 | 0.8657 |
| 486 | PC(18:1/20:5)+AcO | 0.7050 | 0.1883 |
| 487 | PC(18:1/22:4)+AcO | 1.1916 | 0.2118 |
| 488 | PC(18:1/22:5)+AcO | 0.9443 | 0.6938 |
| 489 | PC(18:1/22:6)+AcO | 0.9539 | 0.8286 |
| 490 | PC(18:2/16:1)+AcO | 1.0315 | 0.6596 |
| 491 | PC(18:2/18:2)+AcO | 1.0975 | 0.3251 |
| 492 | PC(18:2/18:3)+AcO | 1.0243 | 0.8754 |
| 493 | PC(18:2/20:1)+AcO | 1.0496 | 0.7976 |
| 494 | PC(18:2/20:2)+AcO | 1.0982 | 0.3918 |
| 495 | PC(18:2/20:3)+AcO | 1.1569 | 0.2020 |
| 496 | PC(18:2/20:4)+AcO | 1.1201 | 0.4035 |
| 497 | PC(18:2/20:5)+AcO | 0.6539 | 0.0594 |
| 498 | PC(18:2/22:4)+AcO | 1.0471 | 0.7669 |
| 499 | PC(18:2/22:5)+AcO | 1.0689 | 0.7166 |
| 500 | PC(18:2/22:6)+AcO | 1.0001 | 0.9997 |
| 501 | PC(20:0/16:1)+AcO | 1.1683 | 0.4679 |
| 502 | PC(20:0/18:1)+AcO | 1.1298 | 0.3053 |
| 503 | PC(20:0/18:3)+AcO | 0.7679 | 0.1167 |
| 504 | PC(20:0/20:1)+AcO | 1.1682 | 0.4738 |
| 505 | PC(20:0/20:3)+AcO | 0.8952 | 0.1597 |
| 506 | PC(20:0/20:4)+AcO | 0.9524 | 0.6210 |
| 507 | PC(20:0/20:5)+AcO | 0.6475 | 0.0322 |
| 508 | PC(20:0/22:4)+AcO | 1.0237 | 0.8817 |
| 509 | PC(20:0/22:6)+AcO | 1.0205 | 0.9194 |
| 510 | LPE(16:0)-H | 0.9769 | 0.8117 |
| 511 | LPE(18:0)-H | 1.0771 | 0.2033 |
| 512 | LPE(18:1)-H | 0.9938 | 0.9742 |
| 513 | LPE(18:2)-H | 0.8946 | 0.3514 |
| 514 | LPE(18:3)-H | 0.9126 | 0.6020 |
| 515 | LPE(20:0)-H | 1.9851 | 0.2500 |
| 516 | LPE(20:1)-H | 0.8625 | 0.8329 |
| 517 | LPE(20:2)-H | 1.4136 | 0.2885 |
| 518 | LPE(20:3)-H | 0.8231 | 0.1601 |

| | | | |
|---|---|---|---|
| 519 | LPE(20:4)-H | 1.2330 | 0.6539 |
| 520 | LPE(20:5)-H | 0.6912 | 0.0871 |
| 521 | LPE(22:4)-H | 1.2926 | 0.0548 |
| 522 | LPE(22:5)-H | 1.0167 | 0.9127 |
| 523 | LPE(22:6)-H | 0.7899 | 0.1938 |
| 524 | PE(14:0/14:0)-H | 0.4622 | 0.2987 |
| 525 | PE(14:0/18:1)-H | 0.6639 | 0.4018 |
| 526 | PE(14:0/18:2)-H | 1.0142 | 0.9266 |
| 527 | PE(14:0/20:3)-H | 1.2120 | 0.4346 |
| 528 | PE(14:0/20:4)-H | 0.9048 | 0.5733 |
| 529 | PE(14:0/22:4)-H | 0.8736 | 0.3359 |
| 530 | PE(14:0/22:5)-H | 1.0939 | 0.5552 |
| 531 | PE(14:0/22:6)-H | 0.8562 | 0.2390 |
| 532 | PE(16:0/14:0)-H | 0.3021 | 0.0533 |
| 533 | PE(16:0/16:0)-H | 0.9626 | 0.8204 |
| 534 | PE(16:0/18:1)-H | 1.0006 | 0.9984 |
| 535 | PE(16:0/18:2)-H | 1.0394 | 0.6427 |
| 536 | PE(16:0/18:3)-H | 0.9203 | 0.5940 |
| 537 | PE(16:0/20:1)-H | 0.9375 | 0.7458 |
| 538 | PE(16:0/20:2)-H | 1.0498 | 0.6516 |
| 539 | PE(16:0/20:3)-H | 0.8554 | 0.3332 |
| 540 | PE(16:0/20:4)-H | 0.8610 | 0.2129 |
| 541 | PE(16:0/20:5)-H | 0.5023 | 0.0572 |
| 542 | PE(16:0/22:4)-H | 0.9973 | 0.9867 |
| 543 | PE(16:0/22:5)-H | 0.8915 | 0.5051 |
| 544 | PE(16:0/22:6)-H | 0.7885 | 0.2317 |
| 545 | PE(18:0/14:0)-H | 1.2900 | 0.3789 |
| 546 | PE(18:0/16:0)-H | 0.9859 | 0.9404 |
| 547 | PE(18:0/16:1)-H | 0.9867 | 0.9763 |
| 548 | PE(18:0/18:0)-H | 0.9220 | 0.6139 |
| 549 | PE(18:0/18:1)-H | 0.9397 | 0.3858 |
| 550 | PE(18:0/18:2)-H | 1.1018 | 0.1493 |
| 551 | PE(18:0/18:3)-H | 0.8888 | 0.1954 |
| 552 | PE(18:0/20:1)-H | 0.9426 | 0.6386 |
| 553 | PE(18:0/20:2)-H | 1.0674 | 0.6972 |
| 554 | PE(18:0/20:3)-H | 1.0369 | 0.8309 |
| 555 | PE(18:0/20:4)-H | 1.0932 | 0.1551 |
| 556 | PE(18:0/20:5)-H | 0.6707 | 0.0182 |
| 557 | PE(18:0/22:4)-H | 1.0858 | 0.7395 |
| 558 | PE(18:0/22:5)-H | 1.0300 | 0.7986 |
| 559 | PE(18:0/22:6)-H | 0.9030 | 0.4494 |
| 560 | PE(18:1/18:1)-H | 0.7392 | 0.3234 |
| 561 | PE(18:1/18:2)-H | 1.0383 | 0.7439 |
| 562 | PE(18:1/18:3)-H | 0.8230 | 0.4194 |
| 563 | PE(18:1/20:1)-H | 0.9224 | 0.7139 |
| 564 | PE(18:1/20:2)-H | 0.8557 | 0.3847 |

| | | | |
|---|---|---|---|
| 565 | PE(18:1/20:3)-H | 0.8183 | 0.2854 |
| 566 | PE(18:1/20:4)-H | 0.9516 | 0.7231 |
| 567 | PE(18:1/20:5)-H | 0.4400 | 0.1355 |
| 568 | PE(18:1/22:4)-H | 1.0512 | 0.7840 |
| 569 | PE(18:1/22:5)-H | 0.9064 | 0.5279 |
| 570 | PE(18:1/22:6)-H | 0.7889 | 0.3155 |
| 571 | PE(18:2/16:1)-H | 0.8990 | 0.5477 |
| 572 | PE(18:2/18:2)-H | 1.3419 | 0.0872 |
| 573 | PE(18:2/18:3)-H | 0.9148 | 0.7819 |
| 574 | PE(18:2/20:1)-H | 1.2316 | 0.2898 |
| 575 | PE(18:2/20:2)-H | 1.2706 | 0.1330 |
| 576 | PE(18:2/20:3)-H | 1.1299 | 0.4758 |
| 577 | PE(18:2/20:4)-H | 1.3167 | 0.5184 |
| 578 | PE(18:2/20:5)-H | 1.1250 | 0.5613 |
| 579 | PE(18:2/22:4)-H | 1.2686 | 0.2500 |
| 580 | PE(18:2/22:5)-H | 1.4053 | 0.1305 |
| 581 | PE(18:2/22:6)-H | 1.1002 | 0.6618 |
| 582 | PE(O-16:0/18:0)-H | 0.8151 | 0.4620 |
| 583 | PE(O-16:0/18:1)-H | 0.8666 | 0.4373 |
| 584 | PE(O-16:0/18:2)-H | 1.0148 | 0.8535 |
| 585 | PE(O-16:0/18:3)-H | 0.7998 | 0.3673 |
| 586 | PE(O-16:0/20:1)-H | 0.8277 | 0.1355 |
| 587 | PE(O-16:0/20:2)-H | 1.1108 | 0.2996 |
| 588 | PE(O-16:0/20:3)-H | 0.8392 | 0.1068 |
| 589 | PE(O-16:0/20:4)-H | 0.9243 | 0.4698 |
| 590 | PE(O-16:0/20:5)-H | 0.8361 | 0.3754 |
| 591 | PE(O-16:0/22:4)-H | 1.1309 | 0.3343 |
| 592 | PE(O-16:0/22:5)-H | 0.8858 | 0.7271 |
| 593 | PE(O-16:0/22:6)-H | 0.9241 | 0.5829 |
| 594 | PE(O-18:0/16:0)-H | 1.0688 | 0.4258 |
| 595 | PE(O-18:0/16:1)-H | 0.8062 | 0.5040 |
| 596 | PE(O-18:0/18:0)-H | 0.8956 | 0.7543 |
| 597 | PE(O-18:0/18:1)-H | 0.7931 | 0.6349 |
| 598 | PE(O-18:0/18:2)-H | 0.9661 | 0.7159 |
| 599 | PE(O-18:0/18:3)-H | 0.6511 | 0.2295 |
| 600 | PE(O-18:0/20:1)-H | 0.7084 | 0.3598 |
| 601 | PE(O-18:0/20:2)-H | 0.6652 | 0.3218 |
| 602 | PE(O-18:0/20:3)-H | 0.7255 | 0.2553 |
| 603 | PE(O-18:0/20:4)-H | 0.9792 | 0.7563 |
| 604 | PE(O-18:0/20:5)-H | 0.4262 | 0.3073 |
| 605 | PE(O-18:0/22:4)-H | 1.1689 | 0.1073 |
| 606 | PE(O-18:0/22:5)-H | 0.8660 | 0.3178 |
| 607 | PE(O-18:0/22:6)-H | 0.8091 | 0.2033 |
| 608 | PE(P-14:0/18:1)-H | 1.4621 | 0.1153 |
| 609 | PE(P-16:0/16:0)-H | 1.1471 | 0.1864 |
| 610 | PE(P-16:0/16:1)-H | 0.8601 | 0.6027 |

| | | | |
|---|---|---|---|
| 611 | PE(P-16:0/18:0)-H | 1.0022 | 0.9776 |
| 612 | PE(P-16:0/18:1)-H | 0.9649 | 0.6209 |
| 613 | PE(P-16:0/18:2)-H | 1.0290 | 0.7246 |
| 614 | PE(P-16:0/18:3)-H | 1.0536 | 0.8010 |
| 615 | PE(P-16:0/20:1)-H | 0.9962 | 0.9731 |
| 616 | PE(P-16:0/20:2)-H | 0.9796 | 0.8247 |
| 617 | PE(P-16:0/20:3)-H | 0.9414 | 0.5511 |
| 618 | PE(P-16:0/20:4)-H | 1.2201 | 0.2574 |
| 619 | PE(P-16:0/20:5)-H | 0.9339 | 0.8974 |
| 620 | PE(P-16:0/22:4)-H | 1.2077 | 0.2777 |
| 621 | PE(P-16:0/22:5)-H | 1.0680 | 0.6779 |
| 622 | PE(P-16:0/22:6)-H | 0.8220 | 0.4576 |
| 623 | PE(P-16:1/18:1)-H | 1.4537 | 0.4157 |
| 624 | PE(P-18:0/16:0)-H | 1.2397 | 0.2508 |
| 625 | PE(P-18:0/16:1)-H | 1.0664 | 0.8673 |
| 626 | PE(P-18:0/18:0)-H | 2.5620 | 0.2949 |
| 627 | PE(P-18:0/18:1)-H | 0.8556 | 0.2414 |
| 628 | PE(P-18:0/18:2)-H | 0.9964 | 0.9630 |
| 629 | PE(P-18:0/18:3)-H | 0.9082 | 0.4821 |
| 630 | PE(P-18:0/20:1)-H | 0.8910 | 0.3104 |
| 631 | PE(P-18:0/20:2)-H | 0.9045 | 0.2240 |
| 632 | PE(P-18:0/20:3)-H | 0.8721 | 0.2517 |
| 633 | PE(P-18:0/20:4)-H | 1.0892 | 0.5179 |
| 634 | PE(P-18:0/20:5)-H | 0.5129 | 0.0999 |
| 635 | PE(P-18:0/22:4)-H | 1.3210 | 0.1755 |
| 636 | PE(P-18:0/22:5)-H | 1.0885 | 0.5837 |
| 637 | PE(P-18:0/22:6)-H | 0.8690 | 0.7189 |
| 638 | PE(P-18:1/16:0)-H | 1.0153 | 0.9308 |
| 639 | PE(P-18:1/16:1)-H | 1.0600 | 0.8146 |
| 640 | PE(P-18:1/18:0)-H | 0.9243 | 0.6011 |
| 641 | PE(P-18:1/18:1)-H | 0.9660 | 0.7019 |
| 642 | PE(P-18:1/18:2)-H | 0.9898 | 0.9228 |
| 643 | PE(P-18:1/20:1)-H | 0.5944 | 0.2410 |
| 644 | PE(P-18:1/20:2)-H | 0.9377 | 0.6827 |
| 645 | PE(P-18:1/20:3)-H | 0.9020 | 0.4607 |
| 646 | PE(P-18:1/20:4)-H | 0.9378 | 0.6668 |
| 647 | PE(P-18:1/20:5)-H | 0.6719 | 0.1142 |
| 648 | PE(P-18:1/22:4)-H | 1.0477 | 0.8035 |
| 649 | PE(P-18:1/22:5)-H | 0.8899 | 0.5466 |
| 650 | PE(P-18:1/22:6)-H | 0.8348 | 0.3464 |
| 651 | PE(P-18:2/18:2)-H | 1.3684 | 0.0202 |
| 652 | PE(P-18:2/20:4)-H | 1.1951 | 0.3291 |
| 653 | PE(P-18:2/22:6)-H | 1.0348 | 0.8290 |
| 654 | LPG(16:0)-H | 1.0734 | 0.6242 |
| 655 | LPG(18:0)-H | 1.1043 | 0.4039 |
| 656 | LPG(18:1)-H | 0.8995 | 0.4680 |

| | | | |
|---|---|---|---|
| 657 | LPG(18:2)-H | 1.0264 | 0.9129 |
| 658 | LPG(20:2)-H | 1.7135 | 0.0848 |
| 659 | PG(14:0/14:0)-H | 0.0485 | 0.3751 |
| 660 | PG(14:0/18:2)-H | 0.8647 | 0.4200 |
| 661 | PG(14:0/20:4)-H | 1.2220 | 0.5561 |
| 662 | PG(14:0/22:5)-H | 1.4067 | 0.2921 |
| 663 | PG(16:0/16:0)-H | 1.1060 | 0.4869 |
| 664 | PG(16:0/18:0)-H | 1.0518 | 0.6612 |
| 665 | PG(16:0/18:1)-H | 0.9354 | 0.6982 |
| 666 | PG(16:0/18:2)-H | 1.0931 | 0.4226 |
| 667 | PG(16:0/18:3)-H | 0.7354 | 0.2345 |
| 668 | PG(16:0/20:2)-H | 1.7868 | 0.0995 |
| 669 | PG(16:0/20:3)-H | 0.8532 | 0.3649 |
| 670 | PG(16:0/20:4)-H | 1.7675 | 0.1080 |
| 671 | PG(16:0/20:5)-H | 0.8388 | 0.4472 |
| 672 | PG(16:0/22:5)-H | 1.8303 | 0.1561 |
| 673 | PG(18:0/18:0)-H | 1.1734 | 0.6493 |
| 674 | PG(18:0/18:1)-H | 1.4485 | 0.1643 |
| 675 | PG(18:0/18:2)-H | 1.6226 | 0.1360 |
| 676 | PG(18:0/18:3)-H | 1.2947 | 0.5001 |
| 677 | PG(18:0/20:0)-H | 1.0192 | 0.9409 |
| 678 | PG(18:0/20:1)-H | 1.1154 | 0.7973 |
| 679 | PG(18:0/20:2)-H | 2.2511 | 0.0351 |
| 680 | PG(18:0/20:3)-H | 1.6346 | 0.1301 |
| 681 | PG(18:0/20:4)-H | 1.7941 | 0.0861 |
| 682 | PG(18:0/22:6)-H | 0.2027 | 0.2441 |
| 683 | PG(18:1/16:1)-H | 0.7938 | 0.1032 |
| 684 | PG(18:1/18:1)-H | 0.9243 | 0.6323 |
| 685 | PG(18:1/18:2)-H | 1.0785 | 0.7209 |
| 686 | PG(18:1/18:3)-H | 1.0614 | 0.8826 |
| 687 | PG(18:1/20:1)-H | 1.1079 | 0.7287 |
| 688 | PG(18:1/20:2)-H | 1.1689 | 0.6581 |
| 689 | PG(18:1/20:3)-H | 1.2774 | 0.2934 |
| 690 | PG(18:1/20:4)-H | 1.2200 | 0.4282 |
| 691 | PG(18:1/20:5)-H | 1.0380 | 0.8392 |
| 692 | PG(18:1/22:4)-H | 1.2574 | 0.1849 |
| 693 | PG(18:1/22:5)-H | 0.9567 | 0.7614 |
| 694 | PG(18:1/22:6)-H | 0.8770 | 0.4946 |
| 695 | PG(18:2/16:1)-H | 1.0167 | 0.9044 |
| 696 | PG(18:2/18:2)-H | 1.0859 | 0.5901 |
| 697 | PG(18:2/18:3)-H | 0.8826 | 0.5387 |
| 698 | PG(18:2/20:1)-H | 0.8604 | 0.5242 |
| 699 | PG(18:2/20:2)-H | 1.0162 | 0.9163 |
| 700 | PG(18:2/20:3)-H | 1.5356 | 0.0704 |
| 701 | PG(18:2/20:4)-H | 0.9516 | 0.7449 |
| 702 | PG(18:2/20:5)-H | 1.2327 | 0.7355 |

| | | | |
|---|---|---|---|
| 703 | PG(18:2/22:4)-H | 1.4017 | 0.2697 |
| 704 | PG(18:2/22:5)-H | 1.1295 | 0.5226 |
| 705 | PG(18:2/22:6)-H | 0.7171 | 0.1407 |
| 706 | PG(20:0/16:1)-H | 1.2137 | 0.5458 |
| 707 | PG(20:0/18:1)-H | 0.8481 | 0.2614 |
| 708 | PG(20:0/18:2)-H | 0.9786 | 0.9294 |
| 709 | PG(20:0/20:2)-H | 1.2304 | 0.2610 |
| 710 | PG(20:0/20:3)-H | 0.9288 | 0.4773 |
| 711 | PG(20:0/20:4)-H | 1.1905 | 0.5666 |
| 712 | PG(20:0/20:5)-H | 1.5807 | 0.2370 |
| 713 | PG(20:0/22:4)-H | 1.2034 | 0.5143 |
| 714 | PG(20:0/22:5)-H | 1.1398 | 0.5045 |
| 715 | PG(20:0/22:6)-H | 0.6880 | 0.3232 |
| 716 | PI(14:0/18:2)-H | 0.9466 | 0.7063 |
| 717 | PI(14:0/20:3)-H | 0.8548 | 0.4516 |
| 718 | PI(14:0/20:4)-H | 0.7453 | 0.2639 |
| 719 | PI(14:0/22:4)-H | 1.0573 | 0.7719 |
| 720 | PI(16:0/16:0)-H | 0.7957 | 0.1557 |
| 721 | PI(16:0/16:1)-H | 1.1420 | 0.7251 |
| 722 | PI(16:0/18:0)-H | 0.9485 | 0.6946 |
| 723 | PI(16:0/18:1)-H | 0.8953 | 0.3648 |
| 724 | PI(16:0/18:2)-H | 1.1300 | 0.0575 |
| 725 | PI(16:0/18:3)-H | 0.7964 | 0.6112 |
| 726 | PI(16:0/20:2)-H | 0.9805 | 0.8321 |
| 727 | PI(16:0/20:4)-H | 0.9004 | 0.2291 |
| 728 | PI(16:0/22:4)-H | 0.8220 | 0.1230 |
| 729 | PI(16:0/22:5)-H | 0.9830 | 0.9098 |
| 730 | PI(18:0/16:1)-H | 1.0478 | 0.8389 |
| 731 | PI(18:0/18:0)-H | 0.7693 | 0.2008 |
| 732 | PI(18:0/18:1)-H | 0.8858 | 0.2242 |
| 733 | PI(18:0/18:2)-H | 1.1665 | 0.0024 |
| 734 | PI(18:0/20:0)-H | 0.7103 | 0.0883 |
| 735 | PI(18:0/20:2)-H | 1.0137 | 0.8923 |
| 736 | PI(18:0/20:3)-H | 1.0200 | 0.7896 |
| 737 | PI(18:0/20:4)-H | 1.0418 | 0.3547 |
| 738 | PI(18:0/22:4)-H | 0.9484 | 0.5728 |
| 739 | PI(18:0/22:5)-H | 0.9856 | 0.8792 |
| 740 | PI(18:0/22:6)-H | 0.8145 | 0.1406 |
| 741 | PI(18:1/18:1)-H | 0.8452 | 0.2849 |
| 742 | PI(18:1/18:2)-H | 1.2419 | 0.0027 |
| 743 | PI(18:1/20:2)-H | 1.1530 | 0.3031 |
| 744 | PI(18:1/20:3)-H | 0.9991 | 0.9931 |
| 745 | PI(18:2/18:2)-H | 0.9712 | 0.8002 |
| 746 | PI(18:2/20:3)-H | 0.7830 | 0.4368 |
| 747 | PI(18:2/20:4)-H | 1.0314 | 0.8521 |
| 748 | PI(18:2/22:4)-H | 1.3980 | 0.2978 |

| | | | |
|---|---|---|---|
| 749 | PI(20:0/16:1)-H | 1.0046 | 0.9800 |
| 750 | PI(20:0/18:1)-H | 0.7915 | 0.1894 |
| 751 | PI(20:0/18:2)-H | 1.0683 | 0.7496 |
| 752 | PI(20:0/20:1)-H | 1.2753 | 0.5471 |
| 753 | PI(20:0/20:4)-H | 0.7143 | 0.1624 |
| 754 | PI(20:0/22:5)-H | 1.2590 | 0.5168 |
| 755 | PS(14:0/18:1)-H | 0.8669 | 0.6562 |
| 756 | PS(14:0/18:2)-H | 1.0415 | 0.7820 |
| 757 | PS(14:0/20:1)-H | 1.1157 | 0.7522 |
| 758 | PS(14:0/20:4)-H | 0.9072 | 0.4555 |
| 759 | PS(14:0/22:4)-H | 0.8225 | 0.6936 |
| 760 | PS(14:0/22:5)-H | 2.4907 | 0.0908 |
| 761 | PS(14:0/22:6)-H | 0.8231 | 0.6948 |
| 762 | PS(16:0/16:0)-H | 1.1205 | 0.6774 |
| 763 | PS(16:0/18:0)-H | 1.3181 | 0.1213 |
| 764 | PS(16:0/18:1)-H | 0.8438 | 0.6739 |
| 765 | PS(16:0/18:2)-H | 1.0025 | 0.9947 |
| 766 | PS(16:0/18:3)-H | 0.6580 | 0.0853 |
| 767 | PS(16:0/20:1)-H | 1.6753 | 0.2698 |
| 768 | PS(16:0/20:2)-H | 1.2496 | 0.6346 |
| 769 | PS(16:0/20:3)-H | 1.5942 | 0.0109 |
| 770 | PS(16:0/20:4)-H | 1.1104 | 0.5636 |
| 771 | PS(16:0/20:5)-H | 1.5634 | 0.1651 |
| 772 | PS(18:0/14:0)-H | 3.2652 | 0.0687 |
| 773 | PS(18:0/16:1)-H | 0.9977 | 0.9948 |
| 774 | PS(18:0/18:0)-H | 1.0407 | 0.8852 |
| 775 | PS(18:0/18:1)-H | 1.1821 | 0.6977 |
| 776 | PS(18:0/18:2)-H | 0.9903 | 0.9453 |
| 777 | PS(18:0/18:3)-H | 1.1110 | 0.7265 |
| 778 | PS(18:0/20:0)-H | 1.5118 | 0.0114 |
| 779 | PS(18:0/20:1)-H | 1.8178 | 0.1819 |
| 780 | PS(18:0/20:3)-H | 1.0825 | 0.6509 |
| 781 | PS(18:0/20:4)-H | 1.0764 | 0.5824 |
| 782 | PS(18:0/22:4)-H | 1.1669 | 0.7734 |
| 783 | PS(18:0/22:5)-H | 1.1257 | 0.5904 |
| 784 | PS(18:1/16:1)-H | 0.4956 | 0.0497 |
| 785 | PS(18:1/18:1)-H | 1.1876 | 0.3571 |
| 786 | PS(18:1/18:2)-H | 1.1188 | 0.4367 |
| 787 | PS(18:1/18:3)-H | 1.3811 | 0.2442 |
| 788 | PS(18:1/20:2)-H | 0.8702 | 0.6872 |
| 789 | PS(18:1/20:3)-H | 0.6931 | 0.0986 |
| 790 | PS(18:1/20:4)-H | 0.9712 | 0.9130 |
| 791 | PS(18:1/22:4)-H | 2.0075 | 0.0261 |
| 792 | PS(18:1/22:5)-H | 1.1980 | 0.3227 |
| 793 | PS(18:1/22:6)-H | 1.1264 | 0.6744 |
| 794 | PS(18:2/16:1)-H | 0.9263 | 0.8157 |

| | | | |
|---|---|---|---|
| 795 | PS(18:2/18:2)-H | 1.2847 | 0.1631 |
| 796 | PS(18:2/18:3)-H | 2.0181 | 0.0613 |
| 797 | PS(18:2/20:1)-H | 0.9957 | 0.9845 |
| 798 | PS(18:2/20:2)-H | 1.4274 | 0.1488 |
| 799 | PS(18:2/20:3)-H | 1.0843 | 0.6677 |
| 800 | PS(18:2/20:4)-H | 1.0059 | 0.9678 |
| 801 | PS(18:2/20:5)-H | 0.7792 | 0.3862 |
| 802 | PS(18:2/22:4)-H | 1.0690 | 0.7620 |
| 803 | PS(18:2/22:5)-H | 1.0309 | 0.8916 |
| 804 | PS(18:2/22:6)-H | 0.6384 | 0.2119 |
| 805 | PS(20:0/16:1)-H | 1.3059 | 0.5252 |
| 806 | PS(20:0/18:1)-H | 1.2685 | 0.4429 |
| 807 | PS(20:0/18:2)-H | 0.9019 | 0.7543 |
| 808 | PS(20:0/18:3)-H | 0.9001 | 0.6669 |
| 809 | PS(20:0/20:1)-H | 1.1516 | 0.5564 |
| 810 | PS(20:0/20:2)-H | 0.9483 | 0.8248 |
| 811 | PS(20:0/20:3)-H | 1.3422 | 0.2529 |
| 812 | PS(20:0/20:4)-H | 1.0402 | 0.8002 |
| 813 | PS(20:0/22:4)-H | 0.9765 | 0.8914 |
| 814 | PS(20:0/22:5)-H | 1.0091 | 0.9580 |
| 815 | PS(20:0/22:6)-H | 1.3502 | 0.2891 |
| 816 | PA(16:0/18:2)-H | 1.4479 | 0.2589 |
| 817 | PA(16:0/20:4)-H | 1.3121 | 0.4021 |
| 818 | PA(18:0/16:1)-H | 1.1386 | 0.6588 |
| 819 | PA(18:0/18:1)-H | 0.9594 | 0.6899 |
| 820 | PA(18:0/18:2)-H | 1.0022 | 0.9791 |
| 821 | PA(18:0/18:3)-H | 0.9126 | 0.5850 |
| 822 | PA(18:0/20:0)-H | 1.0343 | 0.9414 |
| 823 | PA(18:0/20:1)-H | 0.8677 | 0.3908 |
| 824 | PA(18:0/20:2)-H | 0.8374 | 0.6189 |
| 825 | PA(18:0/20:3)-H | 0.9392 | 0.8580 |
| 826 | PA(18:0/20:4)-H | 1.5789 | 0.1103 |
| 827 | PA(18:0/20:5)-H | 1.8584 | 0.1108 |
| 828 | PA(18:0/22:4)-H | 1.8013 | 0.1090 |
| 829 | PA(18:0/22:5)-H | 2.1849 | 0.0850 |
| 830 | PA(18:0/22:6)-H | 0.9689 | 0.8515 |
| 831 | PA(18:1/18:1)-H | 0.7505 | 0.3047 |
| 832 | PA(18:1/18:2)-H | 0.7955 | 0.5590 |
| 833 | PA(18:1/20:3)-H | 1.3378 | 0.2958 |
| 834 | PA(18:1/20:4)-H | 1.4698 | 0.1882 |
| 835 | PA(18:1/22:4)-H | 1.1375 | 0.6379 |
| 836 | PA(18:1/22:5)-H | 1.3987 | 0.0624 |
| 837 | PA(18:2/20:4)-H | 1.0861 | 0.8410 |
| 838 | PA(20:0/16:1)-H | 0.9882 | 0.9676 |
| 839 | PA(20:0/18:1)-H | 0.8211 | 0.2532 |
| 840 | PA(20:0/18:2)-H | 0.9853 | 0.8925 |

| | | | |
|---|---|---|---|
| 841 | PA(20:0/18:3)-H | 1.1348 | 0.2896 |
| 842 | PA(20:0/20:1)-H | 0.9318 | 0.6875 |
| 843 | PA(20:0/20:2)-H | 1.2430 | 0.4395 |
| 844 | PA(20:0/20:3)-H | 0.8656 | 0.1415 |
| 845 | PA(20:0/20:4)-H | 0.9414 | 0.6595 |
| 846 | PA(20:0/20:5)-H | 0.6801 | 0.0117 |
| 847 | PA(20:0/22:4)-H | 1.1098 | 0.6024 |
| 848 | PA(20:0/22:5)-H | 0.9996 | 0.9972 |
| 849 | PA(20:0/22:6)-H | 1.0334 | 0.8350 |
